# Environmentally Induced Sperm RNAs Transmit Cancer Susceptibility to Offspring in a Mouse Model

**DOI:** 10.1101/2020.03.23.004135

**Authors:** Raquel Santana da Cruz, Odalys Dominguez, Elaine Chen, Alexandra K. Gonsiewski, Apsra Nasir, M. Idalia Cruz, Xiaojun Zou, Susana Galli, Kepher Makambi, Matthew McCoy, Marcel O. Schmidt, Lu Jin, Ivana Peran, Sonia de Assis

## Abstract

DNA sequence accounts for the majority of disease heritability, including cancer. Yet, not all familial cancer cases can be explained by genetic factors. It is becoming clear that environmentally induced epigenetic inheritance occurs and that the progeny’s traits can be shaped by parental environmental experiences. In humans, epidemiological studies have implicated environmental toxicants, such as the pesticide DDT, in intergenerational cancer development, including breast and childhood tumors. Here, we show that the female progeny of males exposed to DDT in the pre-conception period have higher susceptibility to developing aggressive tumors in mouse models of breast cancer. Sperm of DDT-exposed males exhibited distinct patterns of small non-coding RNAs, with an increase in miRNAs and a specific surge in miRNA-10b levels. Remarkably, embryonic injection of the entire sperm RNA load of DDT-exposed males, or synthetic miRNA-10b, recapitulated the tumor phenotypes observed in DDT offspring. Mechanistically, miR-10b injection altered the transcriptional profile in early embryos with enrichment of genes associated with cell differentiation, tissue and immune system development. In adult DDT-derived progeny, transcriptional and protein analysis of mammary tumors revealed alterations in stromal and in immune system compartments. Our findings reveal a causal role for sperm RNAs in environmentally induced inheritance of cancer predisposition and, if confirmed in humans, this could help partially explain some of the “missing heritability” of breast, and other, malignancies.

## Introduction

Family history is one of the most important risk factors for cancer. A number of germline mutations are causally linked to familial cancer syndromes. Yet, not all familial cancer cases can be explained by high penetrance genes mutations. Genome-wide association studies (GWAS) of variants have been able to account for a fraction of familial cancer predisposition, but much of the “missing” cancer heritability remains unexplained.(1)

Accumulating evidence suggests that parental life experiences can affect the progeny’s predisposition to disease in an epigenetically inherited manner. (2,3) At conception, parents not only contribute their genome but also transmit a molecular memory of past environmental experiences to the offspring. (2,4) This environmentally-induced disease predisposition has been shown to be transmitted to the offspring via epigenetic mechanisms through both the female and male germ-lines. (5–7) Although most of the evidence for this mode of disease inheritance comes from maternal exposures in pregnancy, a number of independent studies have shown that paternal exposures in the pre-conception window are also important in determining disease outcomes in the offspring. (8–14)

Several recent published reports demonstrated that the sperm RNA load can transmit environmentally-induced phenotypes from fathers to offspring. (2,7,15,16) Some of these studies implicate specific classes of sperm small non-coding RNAs, such as miRNAs and tRNA fragments (tRFs), which can recapitulate the effect of specific paternal exposures in offspring when injected into normal embryos following fertilization. (15–19)

Population studies link parental exposures to environmental pesticides with both childhood and adult cancers in the progeny (20–29). For instance, the epidemiologic analysis using the Child Health and Development Studies cohort (CHDS) found that daughters of women exposed to high levels of dichlorodiphenyltrichloroethane (DDT) in pregnancy are more likely to have increased rates of breast cancer (20,30) and to be diagnosed with advanced stage tumors. (20)

Though the use of DDT has been restricted in many countries for decades, developing countries with endemic malaria, and other vector-borne diseases, continue to use it. (20,31–33) In countries where it has been banned, this pesticide is a persistent organic environmental pollutant and an endocrine disruptor. (31,34) Because of its low degradation rates and lipophilic properties, DDT bio-accumulates in the food chain, and can still be detected in fatty tissues and circulation of animals and humans in these countries. (31) In the U.S., the highest concentrations of DDT metabolites are found in minority populations and recent immigrants.(35,36)

Despite strong association between parental exposures to environmental toxins and cancer in the next generation, the precise mechanisms underlying this association remain unknown. Here, using mouse models of breast cancer, we tested the hypothesis that paternal exposure to DDT alters the sperm non-coding RNA load and increases cancer susceptibility in the next generation. We also examined whether paternal DDT-induced programming of breast cancer development in daughters is mechanistically linked to sperm non-coding RNAs and the underlying mechanisms.

## Results

### Paternal exposure to the environmental toxicant DDT leads to enhanced breast tumor growth in offspring

Parental exposure to environmental toxicants such as DDT and other pesticides has been associated to cancer development, including breast tumors, in the next generation. (20,22,28) To test whether we could replicate these findings in an animal model, adult male mice were either exposed to a vehicle-control (CO) or an environmentally relevant dose of DDT and, subsequently, mated with unexposed females to generate the progeny (**Fig. 1a**). Using the well-established DMBA-induced mouse model of breast cancer, (8,37) we examined mammary cancer development in the female progeny of CO and DDT exposed fathers. Tumor growth was significantly increased in female offspring of DDT exposed fathers (referred to as ‘DDT offspring’ from this point on) compared to CO (**Fig. 1b**), with significantly higher tumor burden and tumor weight at the end of the monitoring period (**Fig. 1c, Fig. S1**). Although not significantly different, the incidence and latency of mammary tumors was slightly higher and shorter, respectively, in DDT offspring compared to CO (**Fig. 1d-e**).

**Figure 1.**
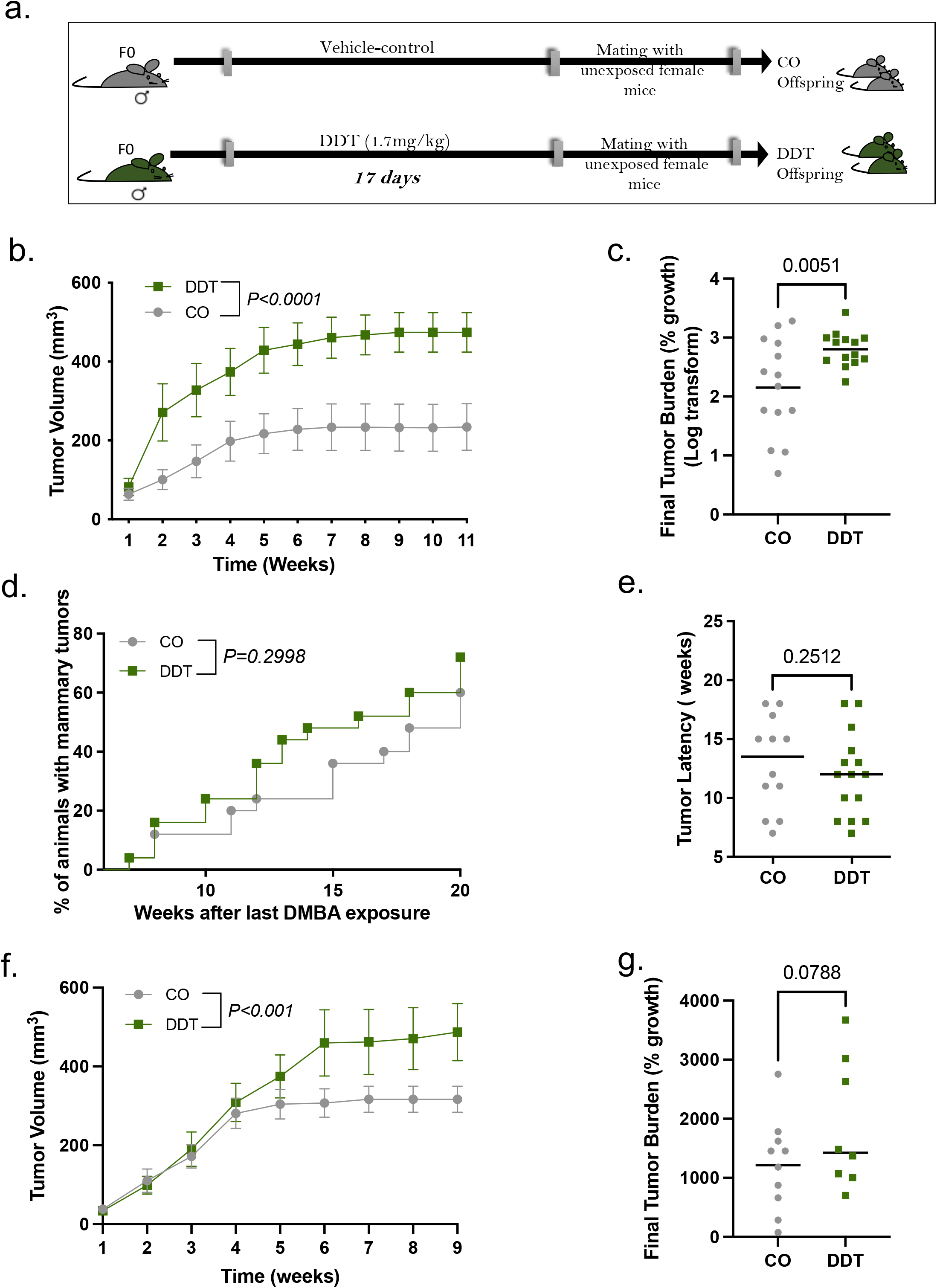
Paternal exposure to DDT leads to enhanced breast tumor growth in offspring. (a) Schematic representation of the experimental design: Eight week-old male mice were treated with a vehicle-control (CO) or a DDT solution (1.7 mg/kg) via oral gave for 17 days. CO and DDT males were subsequently mated to unexposed females to generate offspring. (b-e) Carcinogen-induced mammary tumorigenesis CO and DDT female offspring: (b) Mammary tumor volume (n=14), (c) tumor burden (percentage growth, n=14), (d) tumor incidence (n=25) and (e) tumor latency (n=12-15). (f-g) Orthotopic (EO771 cells) mammary tumorigenesis in CO and DDT female offspring: (f) Mammary tumor volume (n=9-11) and (g) tumor burden (percentage growth, n=8-10). Tumor incidence is shown as percentage of animals with tumors; All other data are shown as mean (horizontal bars in scatter plots) or mean± SEM (tumor growth curves). Tumor burden data does not include tumors with fewer than two measurements. Tumor latency includes both measurable and non-measurable tumors. Tumor volume curves were analyzed using two-away ANOVA (group, time – with repeated measures). Differences in tumor incidence were analyzed using Kaplan-Meier survival curves followed by the log-rank test. Differences in final tumor burden (% growth) and tumor latency were analyzed using unpaired t-test. P values are displayed in each figure panel.

To test whether the increase in carcinogen-induced mammary tumor growth observed in DDT offspring could be replicated in a different *in vivo* system, we used a syngeneic orthotopic mouse model of breast cancer. Consistent with our observations in the carcinogen-induced model, we found that EO771 murine breast cancer cells implanted in mammary fat pads of DDT offspring also resulted in significantly higher tumor growth compared to CO (**Fig. 1f-g**).

### Sperm RNAs contribute to DDT-induced paternal transmission of breast cancer predisposition to offspring

The sperm RNA load, which is abundant in small non-coding RNAs,(38) was previously thought to be a non-functional remnant of the spermatogenesis process. However, several reports show that sperm RNAs can alter early embryonic development, and they have been recently linked to transmission of phenotypes in a variety of disease models. (17–19,39)

To assess whether sperm RNAs are associated with the increased cancer development in DDT offspring, the RNA load extracted from mature sperm of either DDT-exposed or CO male mice was injected into normal mouse embryos at the zygote stage. Injected embryos were then transferred into surrogate dams to generate the DDT-sperm RNA and CO-sperm RNA offspring (**Fig.2a**). Remarkably, DMBA-induced mammary tumors grew significantly larger in DDT-sperm RNA female offspring than in the CO-sperm RNA group (**Fig. 2b-c**), recapitulating the original phenotype observed in DDT offspring. Consistent with results in DDT offspring, we also observed a non-significant increase in tumor incidence and shorter tumor latency in DDT-sperm RNA offspring (**Fig. 2d-e**).

**Figure 2.**
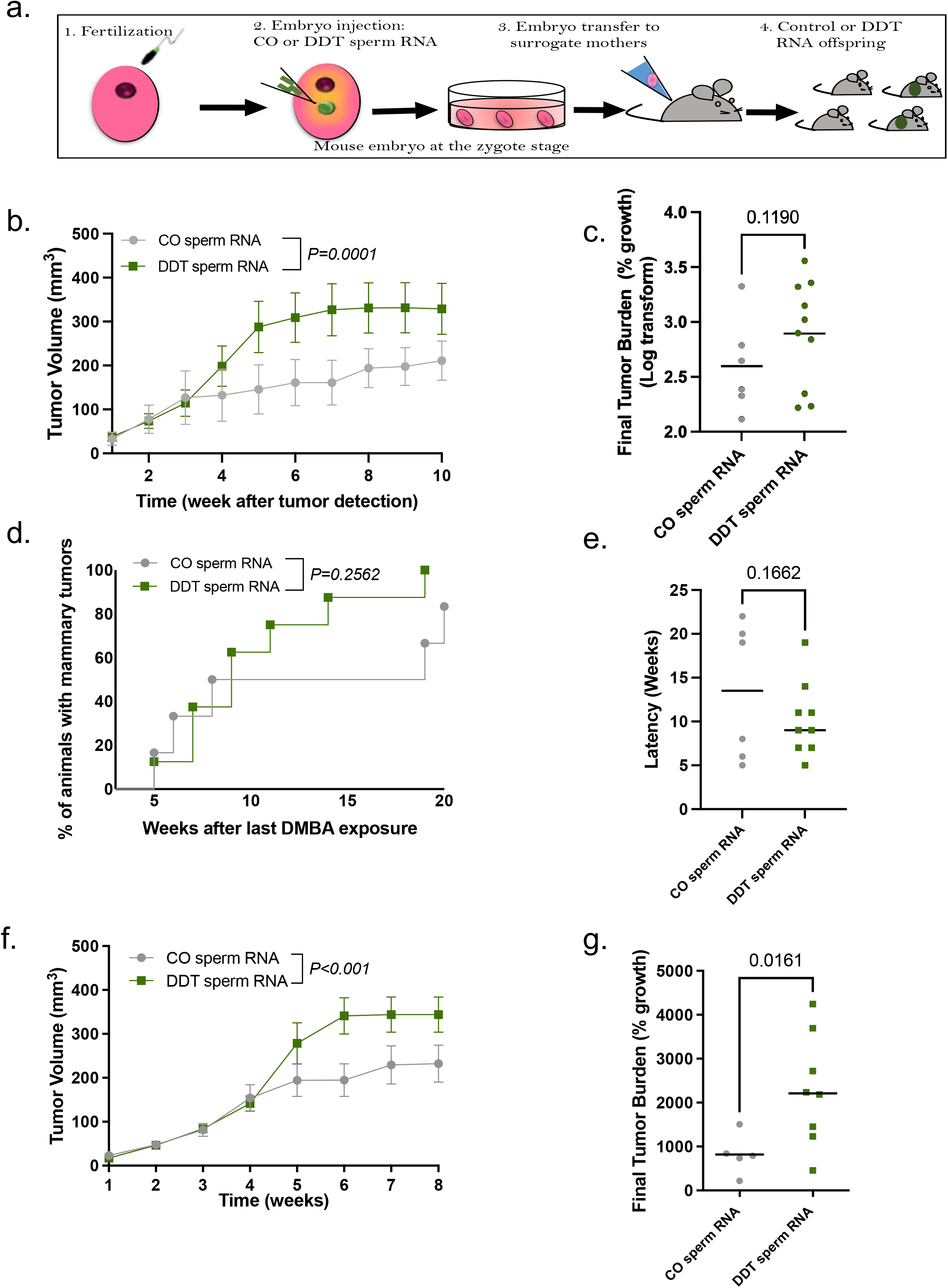
Sperm RNAs contribute to DDT-induced paternal transmission of breast cancer predisposition to offspring. (a) Schematic representation of the experimental design: Normal mouse embryos at the zygote stage were injected with the RNA load from sperm of CO or DDT-exposed males. Injected zygotes were transferred into recipient female mice to produce offspring. (b-e) Carcinogen-induced mammary tumorigenesis in CO-RNA and DDT-RNA female offspring: (b) Mammary tumor volume (n=6-10), (c) tumor burden (percentage growth, n=6-10), (d) tumor incidence(n=7-10) and (e) tumor latency (n=6-10). (f-g) Orthotopic (EO771 cells) mammary tumorigenesis CO-RNA and DDT-RNA female offspring: (f) Mammary tumor volume (n=5-8) and (g) tumor burden (percentage growth, n=5-8). Tumor incidence is shown as percentage of animals with tumors; All other data are shown as mean (scatter plots) or mean± SEM (tumor growth curves). Tumor burden data does not include tumors with fewer than two measurements. Tumor latency includes both measurable and non-measurable tumors. Tumor volume curves were analyzed using two-away ANOVA (group, time – with repeated measures). Differences in tumor incidence were analyzed using Kaplan-Meier survival curves followed by the log-rank test. Differences in final tumor burden (% growth) and tumor latency were analyzed using unpaired t-test. P values are displayed in each figure panel.

Our findings were further confirmed in an orthotopic mouse model of breast cancer. Mammary fat pad injection of EO771 cells resulted in increased mammary tumor growth in DDT-sperm RNA female offspring, which had significantly increased tumor volume over time with increased tumor percentage growth at the end of the monitoring period compared to CO-sperm RNA offspring (**Fig 2f-g**).

### Pre-conception paternal exposure to DDT alters the sperm non-coding RNA content

While mature sperm is transcriptionally inactive, recent studies show that, following testicular exit, the mammalian sperm acquires a load of small non-coding RNAs during transit in the epididymis,(40) the last stage of sperm maturation. Importantly, this non-coding RNA load is sensitive to environmental insults.(2,10,19) To determine the impact of DDT exposure on the mouse sperm RNA content, we performed RNA-seq analysis to profile small RNAs in sperm collected from CO and DDT-exposed mice (**Fig. 3a**). The distribution of the major small RNA species detected in the sperm is in line with previous reports, (18,39) with tRNA-derived fragments (tRFs) being the most abundant small RNA sub-type, followed by miRNAs and other RNAs in both groups (**Fig. 3b**).

**Figure 3.**
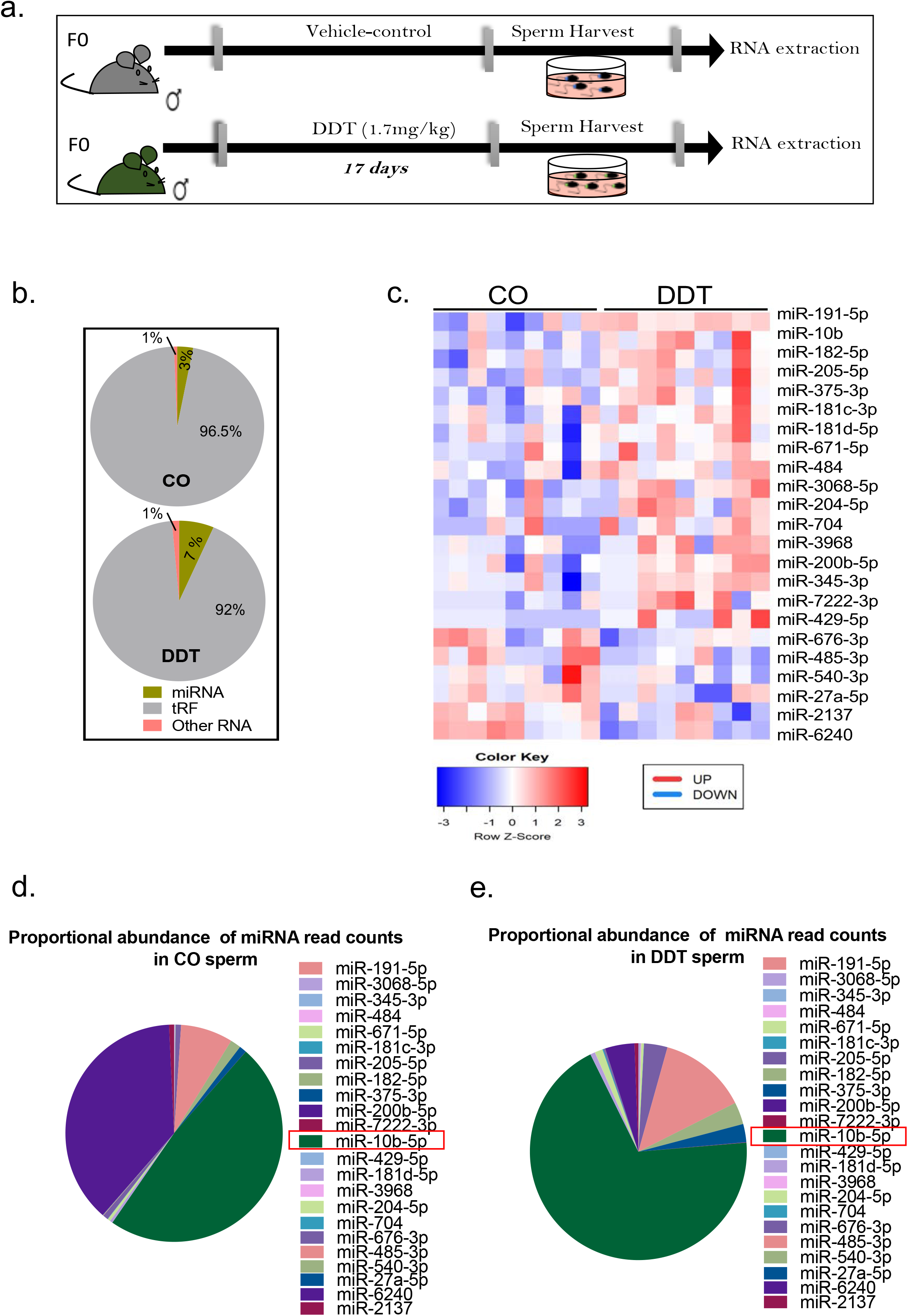
DDT exposure reprograms the sperm small non-coding RNA load. (a) Schematic representation of the experimental design: Eight week-old male mice were treated orally with a vehicle-control (CO) or a DDT solution (1.7 mg/kg) for 17 days. RNA extracted from sperm of CO and DDT-exposed males was analyzed via RNA-seq. (b) Small RNA subtype distribution (percentage of raw reads) and (c) heat-map showing differentially expressed miRNAs in sperm of CO and DDT-exposed males (n=9). (d-e) Proportional abundance of differentially expressed miRNAs (DESeq2 normalized counts) in sperm of (d) CO and (e) DDT-exposed males (n=9).

We found that sperm of DDT-exposed males showed higher abundance of miRNAs reads (7%) compared to the CO group (3%) (**Fig. 3b**). A total of 23 miRNAs were differentially expressed (six down- and seventeen upregulated) in sperm of DDT-exposed mice compared to CO (**Fig. 3c**). Among the upregulated miRNAs, miRNA-10b had the highest surge, representing close to three quarters of the differentially expressed miRNA normalized counts in sperm of DDT-exposed males compared to about half of the reads in the CO group (**Fig. 3d-e**). This is consistent with a recent report showing that miRNA-10b is one the main miRNAs acquired by sperm cells during epididymal transit. (40)

To establish whether regulation of miRNAs in sperm was, in fact, induced by DDT exposure, we treated DDT-exposed mice with phenobarbital (PB, **Fig.4a**), a drug known to promote hepatic enzymes activation and accelerate DDT metabolism and excretion in humans (41,42) and animals. (43–45) Indeed, we observed a sharp increase in the hepatic phase I enzyme, Cyp2B10, mRNA levels (**Fig.4b**) and a reduction in DDT main metabolites in DDT-exposed males also treated with PB (**Fig.4c**).

**Figure 4.**
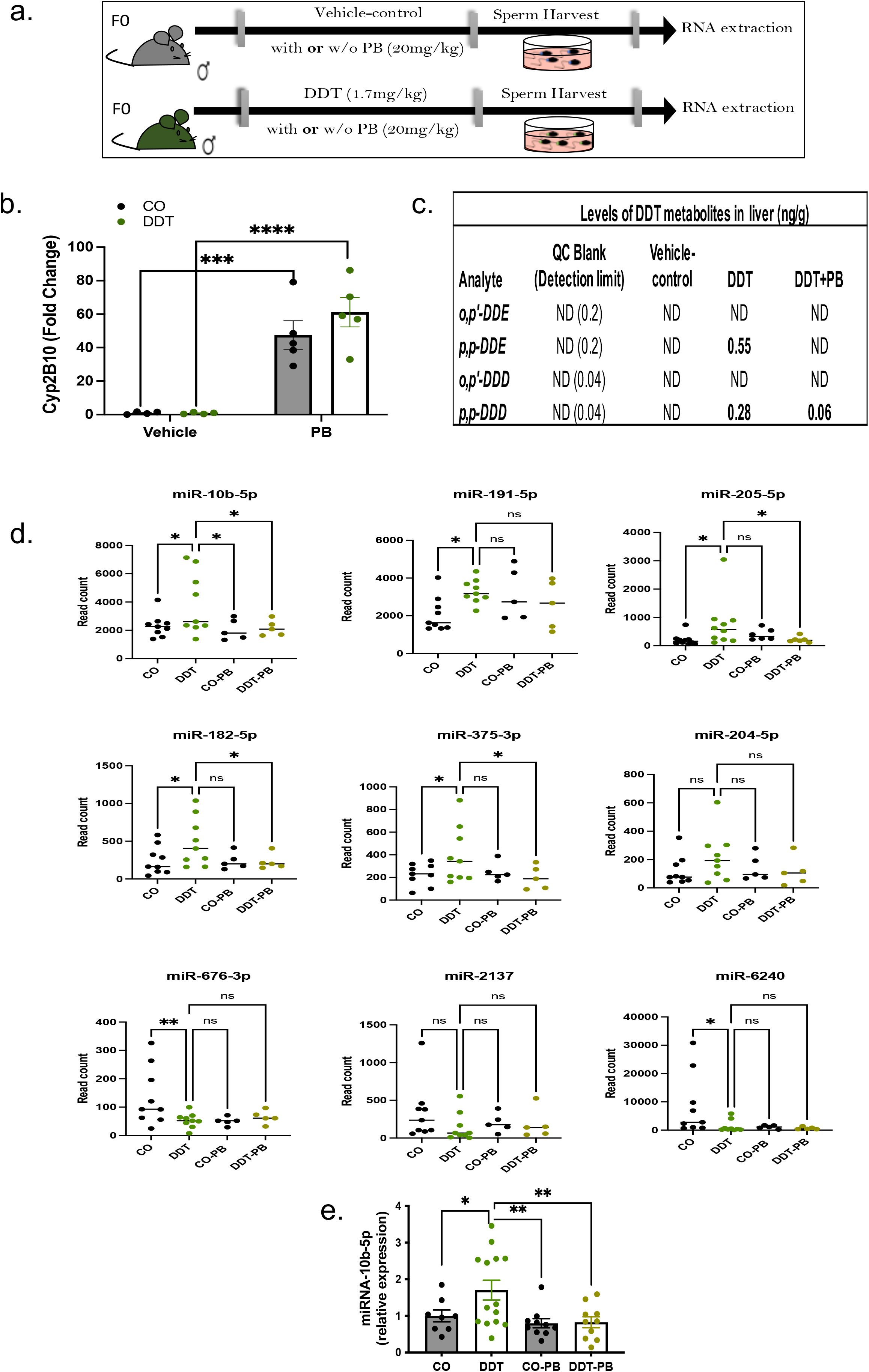
Treatment with the hepatic enzyme inducer, phenobarbital, leads to reduction of DDT metabolites and normalization of sperm miRNA-10b levels in DDT-exposed males. (a) Schematic representation of the experimental design: Eight week-old male mice were treated orally with a vehicle-control (CO) or a DDT solution (1.7 mg/kg) with or without i.p. injection of phenobarbital (PB, 20mg/kg) treatment. (b) Expression levels of *Cyp2B10*, assessed by q-PCR, in liver tissues of CO and DDT-exposed male mice treated with PB or vehicle injection (n=4-5). (c) Levels of DDT metabolites in liver tissues of CO and DDT-exposed male mice treated with PB or vehicle injection (n=3 pools). (d) miRNA expression levels (DESeq2 normalized counts), assessed by RNA-seq, in sperm of CO and DDT-exposed male mice treated with PB or vehicle injection (n=5-9). (e) Expression levels of miRNA-10b, assessed by q-PCR, in sperm of CO and DDT-exposed male mice treated with PB or vehicle injection (n=8-16). Expression levels of *Cyp2B10* and miRNA-10b are relative to GAPDH and miR-26a, respectively. Data are shown as mean (horizontal bars in scatter plots) or mean± SEM (bar graphs). ns, nonsignificant, *p<0.05, **p<0.01, ***p<0.001, ****p<0.0001. Expression data was analyzed using one-way ANOVA.

Next, we investigated the effects of PB treatment on miRNAs that were differentially expressed in sperm of DDT-exposed mice. For greater reliability, our analysis focused on the nine differentially expressed miRNAs, which had an average of at least 100 normalized read counts in our original RNA-seq data.

Compared to mice in the DDT-exposed group, miRNA-10b levels were significantly reversed in sperm of mice treated with both DDT and PB (DDT+PB) with levels returning to those observed in controls (CO and CO+PB) both by RNA-seq and q-PCR (**Fig. 4d-e**).

In addition to miRNA-10b, levels of miRNA-182-5p, miRNA-205-5p, and miRNA-375-3p were also reversed by PB treatment in sperm of DDT-exposed males in the RNA-seq analysis, though these findings could not be confirmed by q-PCR (**Fig. S3**). The expression levels of the five remaining miRNAs— including miR-6240, which had the sharpest decrease—were not reversed by PB treatment (**Fig. 4d** and **Fig. S3**), suggesting that they may not be DDT-specific.

Because tRFs have also been linked to intergenerational transmission of phenotypes, (15,16) we also investigated these small RNAs in sperm of DDT-exposed males via RNA-seq analysis. We found nine tRFs differentially expressed (one upregulated and eight downregulated) in sperm of DDT-exposed mice. However, these differentially expressed tRFs are likely not DDT-specific as they were not reversed by PB treatment and could not be confirmed in the follow-up analysis (**Fig. S4**).

### miRNA-10b embryonic injection recapitulates tumor phenotypes observed in offspring of DDT-exposed fathers

To further validate the functional importance of miRNA-10b to phenotypes observed in DDT offspring, we injected synthetic miR-10b or controls (scramble-miR or vehicle solution) into normal mouse zygotes. Injected embryos were then transferred into surrogate dams to generate the miR-10b and CO offspring (**Fig. 5a**). The resulting miR-10b female offspring showed increased orthotopic (EO771 cells) mammary tumor development, with higher tumor growth compared to both control groups (vehicle injection or scrambled miR, **Fig. 5b**). The miR-10b females also had higher tumor burden and weight compared to the scrambled group (**Fig. 5c, Fig. S5**), at the end of the monitoring period.

**Figure 5.**
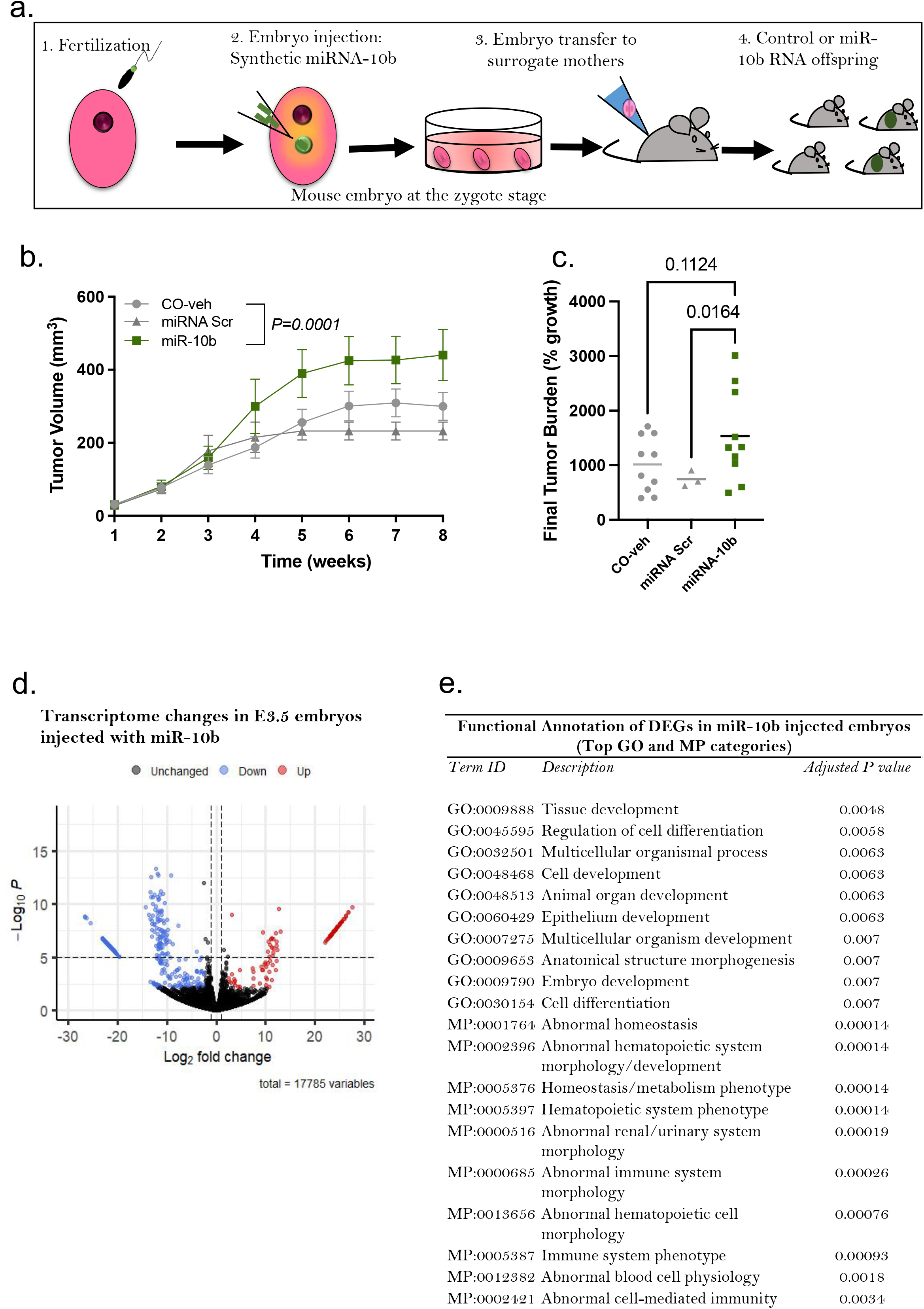
miRNA-10b embryonic injection recapitulates tumor phenotypes observed in offspring of DDT-exposed fathers. (a) Schematic representation of the experimental design: Normal mouse embryos at the zygote stage were injected with synthetic miRNA-10b, scramble miRNA (miR-Src) or vehicle-control (CO-veh). Injected zygotes were culture to the blastocyst stage or transferred into recipient female mice to produce offspring. (b-c) Orthotopic (EO771 cells) mammary tumorigenesis in controls and miR-10b female offspring: (b) Mammary tumor volume (n=3-10) and (c) tumor burden (percentage growth, n=3-10). (d) Volcano plot showing differentially expressed genes (DEG) in miR-10b-injected E3.5 embryos (n=3 pools). (e) Functional annotation of DEG in miR-10b-injected E3.5 embryos. Data are shown as mean (horizontal bars in scatter plots) or mean± SEM (tumor growth curves). Tumor volume curves were analyzed using two-away ANOVA (group, time-with repeated measures). Differences in final tumor burden (% growth) were analyzed using one-way ANOVA. P values are displayed in each figure panel.

Because small RNAs driven alterations in early embryonic development can have long lasting consequences and affect adult phenotype(19), we next examined whether embryonic injection of miR-10b at the zygote stage reshaped the mouse transcriptome profile at the blastocyst stage (E3.5), using bulk RNA-sequencing (**Fig. 5d, Table S1**). We first assessed whether differentially expressed genes (DEG) were potentially regulated by miR-10b. In total, 63 genes down-regulated in miR-10b injected embryos are predicted targets of miR-10b (**Table S2**). Next, we performed a functional annotation analysis to identify the biological signatures most strongly associated with DEGs in miR-10b injected embryos compared to CO. We found an enrichment for terms related to embryonic, cell, tissue and organ development and cell differentiation as well as abnormal hematopoietic and immune system development in miRNA-10b injected embryos (**Fig. 5e)**. Among DEGs involved in immune response several are predicted targets of miR-10b, including *Ifng, Mmp25, Igsf23, Lrrc8b, Mtcp1and Nfam*.

### Tumors of DDT-derived offspring have distinct signaling programs and alterations in the stromal and immune compartments

To define the molecular changes underlying the increased cancer growth phenotype in DDT, DDT-sperm RNA and miR-10b offspring (collectively referred to as DDT-derived offspring from this point on), we performed a bulk RNA-sequencing analysis, followed by a gene set enrichment analysis (GSEA) of DEG, in their tumors. The GSEA revealed several significantly downregulated pathways in DDT-derived offspring tumors compared to CO, including epithelial mesenchymal transition (EMT), interferon gamma (IFN-γ) response and allograft rejection pathways (**Fig. 6a, Fig. S6, Tables S3A-3C**).

**Figure 6.**
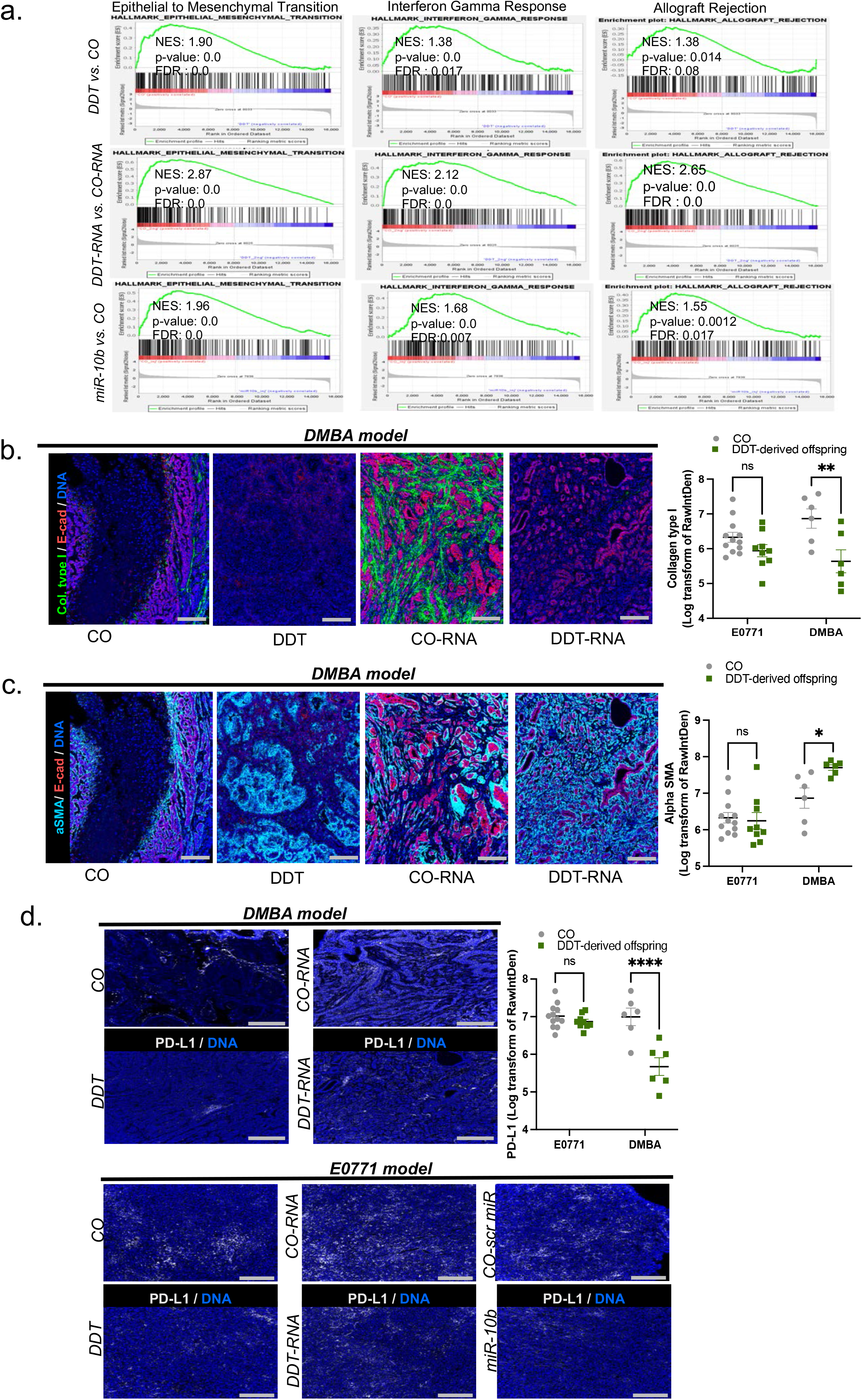
Tumors of DDT-derived offspring have distinct signaling programs and alterations in the stromal and immune compartments. (a) GSEA analysis of DEG in mammary tumors of DDT-derived offspring (DDT, DDT-RNA and miRNA-10b, n=3), showing significantly down-regulated pathways: Epithelial Mesenchymal Transition, Interferon γ Response, and Allograft Rejection. (b) IMC staining for collagen type I (green), E-cadherin (red), DNA (blue, iridium intercalator) and quantification of collagen type I from IMC data in orthotopic (EO771 cells, n=9-12) and carcinogen-induced mammary tumors of DDT-derived offspring (n=6). Scale bar equals to 200 μm. (c) IMC staining for α-SMA (cyan), E-cadherin (red), DNA (blue, iridium intercalator) and quantification of α-SMA from IMC data in in orthotopic (EO771 cells, n=9-12) and carcinogen-induced mammary tumors of DDT-derived offspring (n=6). Scale bar equals to 200 μm. (d) IMC staining for PD-L1(white), DNA (blue, iridium intercalator) and quantification of PD-L1 from IMC data in orthotopic (EO771 cells, n=9-12) and carcinogen-induced mammary tumors of DDT-derived offspring (n=6). Scale bar equals to 200 μm. Horizontal bars in scatter plots represent the mean in b-d. ns, non-significant, *p<0.05, **p<0.01, ****p<0.0001. IMC quantification data was analyzed using two-way ANOVA(group, tumor model).

Our imaging mass cytometry (IMC) analysis confirmed alterations in EMT-associated proteins with a decrease in the extracellular matrix protein collagen type I expression (**Fig. 6b, Fig. S6**) and an increase in α-Smooth Muscle Actin (α-SMA; **Fig. 6c, Fig. S6**) in carcinogen-induced tumors of DDT-derived offspring. Orthotopic tumors generally have a less organized stromal architecture, and although we observed a reduction in collagen type I in EO771 tumors of DDT-derived offspring, the results were not statistically significant.

To confirm the downregulation of immune-related pathways in tumors of DDT-derived offspring, we next investigated whether there were differences in the tumor immune cell infiltrates between the groups, but no consistent differences were detected. Because the IFN-γ signaling has been shown to be a dominant driver of PD-L1 expression in many tumor types,(46) we then examined the levels of this protein in tumors of DDT-derived offspring compared to CO. In line with the downregulation of the IFN-γ response pathway in the transcriptome analysis, we found a significant decrease in PD-L1 protein expression levels in carcinogen-induced tumors—and a trend towards decrease in orthotopic tumors—of DDT-derived offspring compared to CO (**Fig. 6d**).

## Discussion

Family history is one of the most important risk factors for cancer. Yet, not all familial cancer cases can be explained by germline genetic mutations.(1) While there is agreement that the interplay between the environment and genetic factors could also contribute to cancer predisposition, much of the “missing” cancer heritability remains unexplained.

Growing evidence suggests that environmentally induced epigenetic inheritance occurs in mammals and that the progeny’s traits can be shaped by parental environmental experiences.(17,18,39) The findings presented here lend support to this concept: Our study showed that pre-conception paternal exposure to an environmentally relevant dose of the pesticide DDT modulates the sperm non-coding RNA load, particularly miRNAs, and programs female offspring’s breast cancer development and growth in a carcinogen-induced mouse model. These results were substantiated in an orthotopic mouse model of breast cancer. We also showed that sperm RNAs are functionally linked to the cancer phenotypes observed in offspring and demonstrated that embryo injection of total sperm RNA from DDT-exposed males or synthetic miRNA-10b, the miRNA with the highest increase in sperm of DDT-exposed fathers, recapitulated the mammary tumor phenotypes observed in DDT daughters.

Our findings —that adult DDT exposure alters the non-coding small RNA load in sperm— are consistent with other reports showing that environmental insults can modify sperm RNA cargo.(7,17,19) While the proposed mechanisms are still being investigated, recent reports suggest that sperm cells acquire their small RNA load as they transit through the reproductive system and that extra cellular vesicles produced by epididymal cells play a role in this process.(39,47) Furthermore, it has been proposed that during the normal fertilization process, sperm small RNAs are delivered to the oocyte and modulate the transcriptome during the first few cell divisions, setting a signaling cascade that can continue to impact embryonic development. (48) This ripple effect disturbs regulatory networks in the developing embryo which can result in permanent alterations in specific organs.(2,19) In line with this, embryonic injection of sperm RNAs from DDT-exposed male or synthetic miRNA-10b replicates the major tumor phenotypes observed in DDT daughters and transcriptional changes can be detected in pre-implantation embryos injected with miR-10b.

Though we focused on the male germline in our study, it is important to note that environmentally-induced epigenetic alterations likely also occur in the female germline. However, it is technically challenging to directly examine maternal germline effects due to the scarcity of eggs in both rodent and human females as well as the potential confounding of pregnancy exposures.

In our mouse model, DDT offspring appear normal under basal conditions but become more vulnerable to cancer development upon being challenged with a carcinogen or implanted with tumorigenic cells. This suggests that ancestrally acquired traits manifest upon further post-natal insults, acting in synergy to increase cancer susceptibility in offspring. It is possible that the mammary tumor phenotype in DDT-derived offspring results from alterations in immune response (down regulation of genes involved in IFN-γ signaling and reduced PD-L1 levels) and in the mammary stroma (reduced collagen type I). It has been reported that loss of collagen type I is functionally linked to rapid tumor growth and immune suppression. (49) Furthermore, PD-L1 levels has been show to correlate with EMT markers in different solid tumors types.(46) Nonetheless, how, and at what stage of development, these underlying molecular events take place in DDT-derived offspring needs to be further examined. Our data indicates that these alterations may begin in early embryonic development as downregulation of *Ifng* and other immune associated genes are detected in miRNA-10b injected embryos. However, it remains to be determined whether reduction in IFN-γ signaling is maintained throughout development and whether functionally impacts outcomes in vivo.

Our study revealed that miRNAs constituted the main small RNA subtype altered by DDT exposure, with a specific DDT-induced surge in sperm miRNA-10b. This miRNA, along with other members of the miRNA-10 family, plays a critical role in embryonic development and cell differentiation through the regulation of Homeobox *(Hox)* and other genes. (50) Interestingly, studies examining different paternal environmental exposures including obesity and malnutrition report an upregulation of miRNA-10b and other members of the miR-10 family in sperm. (9,10,18,51) It has also been reported that both sperm miRNA-10a and miRNA-10b are conserved across species including humans.(52) These miRNAs are acquired by sperm cells during epididymal transit in mice,(40) suggesting that alterations in their expression could potentially act as a “sensor” of paternal environments.

While life-style factors such as obesity have been shown to alter sperm non-coding RNAs in men,(53) no human studies have directly investigated the link between environmental toxicants such as DDT and germline RNA changes to the best of our knowledge; though several reports in animal models exist.(54–56) Nonetheless, a number of epidemiological studies link parental environmental toxins and cancer in the progeny. For instance, an association between paternal exposure to pesticides and a number of childhood cancers in offspring has been reported. (22–29) Furthermore, published reports using the CHDS cohort showed that maternal exposure to DDT increases rates of breast cancer in daughters who are also more likely to be diagnosed with advanced-stage tumors. (20) Another recent population study using a historical cohort showed that paternal grandfathers’ nutrition leads to a transgenerational increase in cancer mortality rates in grandsons.(57)

In summary, our study shows that the environmentally induced alterations in paternal sperm non-coding RNAs are functionally linked to increased cancer susceptibility in the progeny. Though our results are intriguing and suggest that cancer predisposition could be determined via epigenetic inheritance, many questions still remain: For instance, how sperm non-coding RNAs and miR-10b specifically alter early embryonic development to disrupt offspring’s systemic and mammary specific development needs to be further examined. Although epidemiologic studies strongly suggest it, whether this phenomenon is restricted to breast tumors or whether it applies to other malignancies needs confirmation in additional experimental models.

According to published studies and the Center for Disease Control and Prevention (CDC), minorities such as African Americans and Mexican-Americans and recent immigrants have the highest circulating concentrations of DDT metabolites and other environmental toxins in the U.S. (35,36,58). Thus, our findings are likely relevant to other environmental pollutants which, like DDT, have endocrine disrupting properties(59–61) and could have important implications for cancer disparities.

## Materials and Methods

### DDT exposures and breeding

The *c57bl/6* strain of mice was used in all experiments. Adult male mice (8 weeks of age) were either treated daily with a DDT solution (1.7 mg/kg) or vehicle-control solution (peanut oil) via oral gavage for 17 days. Duration of exposures was chosen in order to encompass the entire period of sperm transit in the epididymis (which takes about 15 days in mammals) and when the sperm acquire its mature non-coding RNA load (62). The DDT solution used mimics the formulation of DDT before its ban in the U.S.: 77.2% p,p’-DDT and 22.8% o,p’-DDT as described before (63). Mice body weight was monitored weekly. At the end of the exposure period, DDT-exposed and control male mice were mated to unexposed females to generate the female offspring. Mice were kept on a standard chow diet during the breeding period, for the extent of pregnancy (21 days) and after birth. To avoid litter-effect, pups were cross-fostered one day after birth. Pups from 2-3 dams were pooled and housed in a litter of 8-10 pups per nursing dam. All pups were weaned on postnatal day (PND) 21. The female offspring of control or DDT-exposed fathers were used to study mammary tumorigenesis and for tissue collection as described in the following sections. All animal procedures were approved by the Georgetown University Animal Care and Use Committee, and the experiments were performed following the National Institutes of Health guidelines for the proper and humane use of animals in biomedical research. All exposures were performed blindly.

### DDT and Phenobarbital (PB) treatment

PB treatment is known to promote hepatic enzymes activation, accelerating DDT metabolism and excretion in humans (41,42) and animals (43–45). A sub-set of control-vehicle (CO) or DDT exposed male mice (as above) were concomitantly treated with phenobarbital (20 mg/kg of body weight, i.p.) or saline and used for sperm harvesting.

### Mature spermatozoa collection and purification

CO and DDT-exposed male mice as well as those concomitantly treated with PB were euthanized and caudal epididymis dissected for sperm collection. The epididymis was collected, punctured, and transferred to a tissue culture dish containing M2 media (M2 Medium-with HEPES, without penicillin and streptomycin, liquid, sterile-filtered, suitable for mouse embryo, SIGMA, product #M7167) where it was incubated for 1 hour at 37°C. Sperm samples were isolated and purified from somatic cells. Briefly, the samples were washed with PBS, and then incubated with SCLB (somatic cell lysis buffer, 0.1% SDS, 0.5% TX-100 in Diethylpyrocarbonate water) for 1 hour. SCLB was rinsed off with 2 washes of PBS and the somatic cell-free purified spermatozoa sample pelleted and used for RNA extraction.

### Measurement of DDT metabolites

Liver tissues from CO, DDT and DDT-PB treated males, collected within 24 h after last dose, were pooled (3 samples/group) and used to determine the levels of DDT’s main metabolites. Measurements were performed commercially by Pacific Rim Labs, a fully accredited laboratory providing analysis of DDT and other persistent organic pollutants. Briefly, samples were fortified with each of the six 13C-DDT isotopes and then extracted with hexane. The extract was columned on Florisil and analyzed by GC-MS/MS. The instrument was calibrated with a five-point calibration covering the range of 0.2-400 ng/g or mL.

### Sperm Small RNA Sequencing

Total RNA was isolated from purified sperm using Qiagen’s miRNeasy extraction kit, according to the manufacturer’s instructions. RNA integrity and quality was examined by Bioanalyzer 2100 (Agilent Technologies). Small non-coding RNA transcript libraries were constructed according to Illumina TrueSeq Small RNA Pre-Kit. After quality control, the library preparations were sequenced on an Illumina platform and reads generated. Raw data quality were checked using FastQC (v0.11.9), and adapter trimming on raw data were performed using Cutadapt (v3.5). Reads with low quality (quality score < 25, error rate > 10%) or reduced length after trimming (<15 bp) were removed before alignment. Single end trimmed reads was aligned against mouse reference genome downloaded from Ensembl GRCm38 release 101 using STAR (v2.7.9a). Small RNA tags were annotated with miRNA and t-RNA sequence fragments (tRFs) obtained from miRBase, GtRNAdb, Ref-seq, GenBank and Rfam databases using featureCounts (v2.0.3). DESeq2 R package (v1.36.0) was used for group wise statistical analysis. To allow for broad pattern identification of small RNAs, significance was set at p-value < 0.05, with a fold change threshold at >1.5.

### Quantitative real time PCR validation (qRT-PCR)

cDNA was synthesized from total RNA samples using the TaqMan^™^ Advanced cDNA Synthesis Kit (Applied Biosystems). PCR products were amplified from cDNA using TaqMan® Fast Advanced Master Mix and sequence-specific primers from TaqMan Assays (Applied Biosystems). Fold change was calculated from Ct values and the expression levels of specific miRNAs or genes were determined by normalizing these values with the fold change values for appropriate endogenous controls (miR-26a or *GAPDH*, respectively).

### Embryonic RNA microinjections

To induce super-ovulation, female mice (*c57bl/6*) received intra-peritoneal injections of 5 IU of pregnant mare’s serum gonadotropin (PMSG). About 48 hours after the PMSG injection, mice were injected with 5 IU of human chorionic gonadotropin (hCG). Super-ovulated females were mated with *c57bl/6* male mice. About 16 hours later, fertilized eggs were harvested. Zygotes were injected with the RNA load from sperm of CO and DDT-exposed males (sperm RNA isolated from 5 mice per group was pooled prior to injection). Each zygote received 10 pL injection of the sperm RNA load (the amount of sperm RNA injected is the equivalent to RNA from 10 sperm cells (7,18,19) or approximately 20-100 femtograms per injection). Microinjections were confirmed by observing the swelling of the male pro-nucleus.

Injected zygotes were transferred into recipient female mice to produce offspring as described before (17). Briefly, recipient females were mated with vasectomized males overnight and identified by copulation plug before the zygote transfer. Each recipient female was implanted with 25-30 zygotes. Dams were housed in groups of two with free access to food and water. Female offspring were used to study tumor growth or for tissue collection.

Another subset of zygotes was injected with synthetic miRNA**-**10b or scramble miR (at 20 femtograms per injection; *mirVana®*, ThermoFisher) diluted in TE solution (10mM Tris-HCl, pH7.4, 0.25 EDTA) or TE solution only (to control for microinjection). Synthetic miRNAs are double-stranded RNA oligonucleotides that mimic a specific endogenous mature miRNA. Injected zygotes were transferred into recipient female mice as described above for term delivery. Female offspring were used to study tumor growth or for tissue collection.

A subset of the of zygotes injected with synthetic miRNA-10b or controls were cultured and harvested at the blastocyst stage (E3.5) and used for RNA-seq analysis. Briefly, injected one-cell zygotes were cultured in Advanced KSOM (Millipore, MR-101-D) at 37°C in 5% CO2 until collected. Embryos at the blastocyst stage were collected at approximately 72 hours after miRNA microinjection (~E3.5) and placed in M2 (Millipore, MR-015-D) and stored in −80°C.

### Carcinogen-induced mammary tumorigenesis

Mammary tumors were induced in CO and DDT-derived female offspring (CO, DDT, CO-RNA, DDT-RNA) by administration of MPA (medroxyprogesterone acetate, 15 mg, s.c.) at 6 weeks of age, followed by three weekly doses of 1mg of 7,12-dimethylbenz[a]anthracene (DMBA) (Sigma, St. Louis, MO) dissolved in peanut oil by oral gavage. This established model of breast cancer has been used by us and others (8,37). Mice were examined for mammary tumors by palpation once per week, starting on week two after the last dose of DMBA and continue for up to 20 weeks after tumor detection. During follow-up, those animals in which tumor burden approximated 10% of total body weight were euthanized, as required by our institution. Tumor growth was measured using a caliper and the width and height of each tumor were recorded. Other end-points for data analysis were latency to tumor appearance and tumor incidence (percentage of animals with tumor in a given group). Histopathology of tumors was determined by a pathologist (S.G.). All tumors included in our analyses were classified as mammary carcinomas.

### Syngeneic orthotopic tumor implantation

Syngeneic mammary tumors were induced in CO and DDT-derived female offspring [CO, DDT, CO-RNA, DDT-RNA, miR-10b and controls (scramble-miR or vehicle solution)]. Mouse EO771 breast cancer cells were cultured in complete medium [Dulbecco’s Modified Eagle’s Medium (DMEM) supplemented with 10% (v/v) FBS and 1% L-glutamine, at 37°C in 5% CO2 (v/v)] to 80% confluence, trypsinized, counted, checked for viability and re-suspended in PBS for injection into both 4^th^ inguinal mammary glands. A total of 1×10^6^cells were injected into each of the 4^th^ inguinal mammary gland fat pads. Tumor growth was monitored for ten weeks. During this period, those animals in which tumor burden approximated 10% of total body weight were euthanized, as required by our institution. Histopathology of tumors was determined by a pathologist (S.G.). All tumors included in our analyses were classified as mammary carcinomas.

### Whole tissue RNA sequencing

Total RNA was isolated from snap frozen mouse tissues (mammary tumors and embryos at the blastocyst stage) using Qiagen’s RNeasy extraction kit, according to the manufacturer’s instructions. RNA integrity and quality was examined by Bioanalyzer 2100 (Agilent Technologies). Transcript libraries were constructed using the TruSeq mRNA Library Prep Kit (Illumina). After cluster generation, the library preparations were sequenced on an Illumina platform and paired-end reads were generated. RNAseq raw data quality was checked using FastQC (v0.11.09) and adapter trimming on raw data was performed using Cutadapt (v3.5). Reads with low quality (quality score < 33, error rate > 10%) or reduced length after trimming (<25 bp) were removed before alignment. We used the reference genome downloaded from Ensembl GRCm38 release 101, and the reference index was built using Star (v2.7.9a). Paired end trimmed read alignment and raw read count calculation was performed using RSEM software (v1.3.1), and an additional bioinformatic workflow utilized Hisat2 v2.0.5 for alignment to the GRCm38 genome and quantified to the Gencode vM25 transcriptome annotation using Stringtie 2.2.1. For both workflows, differential expression analysis of two conditions/groups was performed using the DESeq2 R package (v1.36.0), and visualized using the Enhanced Volcano R package (v3.16). We considered the genes with q-value <0.1 to be differentially expressed (DEG) in embryos and genes with a p-value <0.05 to be DEG in mammary tumors to account for tissue heterogeneity. DEGs in mammary tumors were used as input for Gene Set Enrichment Analysis (GSEA) (v4.2.3, Broad Institute). Both KEGG and HALLMARK gene sets were selected for enrichment score calculation. DEGs in E3.5 embryos were used as input for Gene Ontology (GO) functional characterization and pathway enrichment analysis. miR-10b target prediction in E3.5 embryos was determined using TargetScan and miRBase.

### *Imaging mass cytometry (IMC):* Staining Procedures

After baking for 2 hours at 60 °C, FFPE sections were dewaxed and rehydrated through a graded alcohol series. Heat-induced epitope retrieval was conducted in a steamer at 96 °C in antigen retrieval AR9 buffer for 30 min. After cooling, the sections were blocked with 3% BSA in PBS for 45 min at room temperature. Samples were incubated overnight at 4 °C with metal-conjugated antibody cocktails diluted in PBS/0.5% BSA (See **Table S4** for antibody list and dilutions). Samples were then washed twice with PBS/0.5% Tween and twice with PBS and exposed to 1:400 Ir-Intercalator (Fluidigm) for 30 min at RT for nuclei staining. The samples were rinsed in distilled water and air-dried before image acquisition.

### Image Acquisition

IMC data acquisition and visualization were performed as previously described(64,65). Briefly, stained and air-dried tumor slides were inserted into the ablation chamber of the Hyperion™ Imaging System (Fluidigm). The tissue was then scanned by a pulsed 200 Hz laser focused to a 1 μm spot and applied over a user-defined regions of interest (ROIs), ablating adjacent spots in 1 μm steps as the slide moves under the laser beam. Tissue from each ablation spot was vaporized and streamed by inert gas into the inductively coupled plasma ion source for analysis by the mass cytometer. Images of each mass channel were reconstructed by plotting the laser shot signals in the order in which they were recorded, line scan by line scan. Multi-channel images were overlaid for desired channel combinations in MCD™ Viewer v1.0.560.2 (Fluidigm).

### Statistical analysis

Statistical analyses were performed using GraphPad Prism (GraphPad Software, San Diego, CA, USA). Normal probability plots were used to ascertain normality, which were confirmed by Anderson-Darling, D’Agostino-Pearson, Kolmogorov-Smirnov and Shapiro-Wilk tests. Where data failed the normality test, log transformation was performed prior to statistical analysis. Two-way ANOVA was used to analyze tumor growth curves (group, time – with repeated measures) and IMC quantification data (group, tumor model) followed by post-hoc analysis using Fisher’s LSD multiple comparison tests. Differences in tumor incidence (time to tumor appearance) were analyzed using Kaplan-Meier survival curves followed by the log-rank test. Differences in tumor latency (DMBA model only) and final tumor burden (% growth) were analyzed using unpaired t-test (for two groups) or Welch one-way ANOVA (for three groups), followed by Welch’s t test. Gene and miRNA expression data was analyzed using one-way ANOVA, followed by Fisher’s LSD multiple comparison test. Differences were considered statistically significant at P< 0.05. Unless indicated, *n* corresponds to the number of animals used in each experiment.

## Acknowledgments

We thank the following Lombardi Cancer Center Shared Resources (SR) for their assistance: Animal Model SR, Histopathology & Tissue SR, Microscopy and Imaging SR and Genomics & Epigenomics SR. We also thank Mr. Chip Hawkins from the Johns Hopkins Transgenic Core Facility for technical support with embryos injections. We thank Dr. Sandra Jablonski and Dr. Louis Weiner for donating the murine EO771 cells.

## Data availability

RNA-seq data (small RNAs and whole transcriptome) has been deposited in GEO (Gene Expression Omnibus) database under accession codes GSE221649 and GSE222357.

## Author contributions

S.D.A. conceived the study, oversaw the research and wrote the manuscript; R.S.D.C. conceived the study, supervised the animal work and tissue collection, performed molecular analysis and wrote the manuscript; O.D., E.C., A.N., A.K.G. and X.Z. assisted with animal studies, processed tissues and/or helped perform molecular analysis; I.P. performed IMC analysis, M.I.C. assisted with animal studies; S.G. performed histopathological classification of tumors; L.J., M.S. and M.M. performed RNA-seq data and enrichment analysis; K.M. assisted with statistical analysis.

