## Supplemental Table 1 for "Environmentally Induced Sperm RNAs Transmit Cancer Susceptibility to Offspring in a Mouse Model"

**Table S1-Differentially expressed genes in E3.5 embryos injected with miR-10b**

| | log2FoldCh | pvalue | padj |
|---|---|---|---|
| 1110051M20Rik | -10.9994 | 1.55E-05 | 0.000286 |
| 1110065P20Rik | 2.050278 | 0.005477 | 0.083929 |
| 1600010M07Rik | -11.2637 | 6.06E-12 | 1.47E-08 |
| 1700013H16Rik | -20.8538 | 2.26E-06 | 4.90E-05 |
| 1700030K09Rik | -20.7961 | 2.41E-06 | 5.03E-05 |
| 1700042O10Rik | -22.3785 | 3.85E-07 | 3.21E-05 |
| 1700056N10Rik | -21.2795 | 1.39E-06 | 3.90E-05 |
| 1700126G02Rik | 8.792007 | 0.0031 | 0.050036 |
| 1810024B03Rik | -21.5879 | 9.76E-07 | 3.60E-05 |
| 1810026B05Rik | -3.62343 | 0.00094 | 0.015913 |
| 2210010C04Rik | -20.4644 | 3.47E-06 | 6.73E-05 |
| 2210039B01Rik | -21.2107 | 1.51E-06 | 4.02E-05 |
| 2210408I21Rik | -9.35592 | 1.06E-05 | 0.000198 |
| 2210409E12Rik | -9.20541 | 2.11E-13 | 8.98E-10 |
| 2310030G06Rik | -12.6157 | 7.29E-09 | 2.00E-06 |
| 2310068J16Rik | -21.7593 | 8.00E-07 | 3.50E-05 |
| 2500002B13Rik | -12.0234 | 5.55E-06 | 0.000105 |
| 2610002M06Rik | -21.4587 | 1.13E-06 | 3.65E-05 |
| 2610206C17Rik | -22.0875 | 5.44E-07 | 3.21E-05 |
| 2810029C07Rik | 25.24267 | 5.47E-09 | 1.69E-06 |
| 2810030D12Rik | -21.4931 | 1.09E-06 | 3.65E-05 |
| 2810408A11Rik | -20.9158 | 2.11E-06 | 4.70E-05 |
| 2810429I04Rik | -20.7444 | 2.55E-06 | 5.12E-05 |
| 3110040M04Rik | -21.1143 | 1.68E-06 | 4.23E-05 |
| 3110045C21Rik | -22.5333 | 3.19E-07 | 2.97E-05 |
| 4632411P08Rik | -20.8017 | 2.39E-06 | 5.03E-05 |
| 4732471J01Rik | -21.5409 | 1.03E-06 | 3.65E-05 |
| 4930466K18Rik | -21.355 | 1.28E-06 | 3.78E-05 |
| 4930500L23Rik | 24.34617 | 1.86E-08 | 4.09E-06 |
| 4930507D05Rik | -21.1135 | 1.68E-06 | 4.23E-05 |
| 4930509G22Rik | 23.22052 | 8.12E-08 | 1.17E-05 |
| 4930513N10Rik | -21.0717 | 1.76E-06 | 4.27E-05 |
| 4930548H24Rik | -8.53184 | 0.005835 | 0.088761 |
| 4930556M19Rik | -22.0414 | 5.74E-07 | 3.21E-05 |
| 4930578M01Rik | -21.3651 | 1.26E-06 | 3.78E-05 |
| 4930579G18Rik | -22.0024 | 6.01E-07 | 3.21E-05 |
| 4930581F22Rik | 9.436802 | 0.000162 | 0.002876 |
| 4930590J08Rik | -20.9509 | 2.03E-06 | 4.62E-05 |
| 4933401D09Rik | 23.29401 | 7.39E-08 | 1.09E-05 |
| 4933421O10Rik | -20.3584 | 3.89E-06 | 7.47E-05 |
| 4933427D06Rik | -20.8219 | 2.34E-06 | 5.01E-05 |
| 4933440M02Rik | -21.3502 | 1.28E-06 | 3.78E-05 |
| 4933440N22Rik | -22.3909 | 3.79E-07 | 3.18E-05 |
| 5033403F01Rik | -20.8017 | 2.39E-06 | 5.03E-05 |
| 5430414B19Rik | -21.2795 | 1.39E-06 | 3.90E-05 |
| 5530601H04Rik | -11.1532 | 1.03E-09 | 5.31E-07 |

| | | | |
|---|---|---|---|
| 5730409E04Rik | -11.1881 | 4.94E-07 | 3.21E-05 |
| 5930430L01Rik | -21.4829 | 1.10E-06 | 3.65E-05 |
| 6030400A10Rik | -20.9742 | 1.97E-06 | 4.57E-05 |
| 6330403L08Rik | -11.3907 | 2.45E-05 | 0.000448 |
| 9030625G05Rik | 7.535073 | 0.000241 | 0.004227 |
| 9330160F10Rik | 7.278485 | 0.00011 | 0.001964 |
| 9430064I24Rik | 10.87875 | 6.51E-07 | 3.26E-05 |
| 9530059O14Rik | -22.0667 | 5.57E-07 | 3.21E-05 |
| 9530082P21Rik | -11.0659 | 0.000346 | 0.006014 |
| 9630013K17Rik | -20.6196 | 2.92E-06 | 5.76E-05 |
| 9930004E17Rik | -21.4931 | 1.09E-06 | 3.65E-05 |
| A230020J21Rik | -9.46408 | 3.48E-07 | 3.09E-05 |
| A330009N23Rik | -22.423 | 3.65E-07 | 3.15E-05 |
| A530084C06Rik | -21.6778 | 8.79E-07 | 3.55E-05 |
| A930012O16Rik | -21.436 | 1.16E-06 | 3.69E-05 |
| A930029G22Rik | -22.1793 | 4.88E-07 | 3.21E-05 |
| AA386476 | -21.2442 | 1.45E-06 | 4.00E-05 |
| Abca13 | -21.0755 | 1.76E-06 | 4.27E-05 |
| Abcc2 | -21.1041 | 1.70E-06 | 4.23E-05 |
| Abcc3 | 23.65089 | 4.66E-08 | 7.97E-06 |
| Abcg1 | -10.4458 | 4.74E-05 | 0.000858 |
| Abhd14b | -12.1862 | 8.90E-07 | 3.57E-05 |
| Acvr1b | -1.65863 | 0.002514 | 0.041 |
| Acvrl1 | 23.11911 | 9.25E-08 | 1.21E-05 |
| Adamts18 | -22.0875 | 5.44E-07 | 3.21E-05 |
| Adap2os | -21.4015 | 1.21E-06 | 3.75E-05 |
| Adat3 | -11.0982 | 0.006496 | 0.097164 |
| Adck2 | -23.0194 | 1.77E-07 | 1.97E-05 |
| Adcyap1r1 | -21.011 | 1.89E-06 | 4.46E-05 |
| Add1 | -1.13797 | 0.006505 | 0.097207 |
| Adgrd2-ps | -21.6096 | 9.52E-07 | 3.60E-05 |
| Adgrg6 | -22.0667 | 5.57E-07 | 3.21E-05 |
| Adh1 | -14.3088 | 2.01E-10 | 1.63E-07 |
| AF067061 | -22.1223 | 5.22E-07 | 3.21E-05 |
| Afdn | -1.85194 | 1.50E-05 | 0.000279 |
| Aga | -21.1041 | 1.70E-06 | 4.23E-05 |
| Ahsg | -21.3331 | 1.31E-06 | 3.80E-05 |
| Akap5 | -21.9138 | 6.67E-07 | 3.26E-05 |
| Akap9 | -7.51503 | 0.002324 | 0.03794 |
| Akna | -20.8219 | 2.34E-06 | 5.01E-05 |
| Akr1d1 | -21.559 | 1.01E-06 | 3.65E-05 |
| Aldh1a1 | -20.9395 | 2.05E-06 | 4.64E-05 |
| Amigo2 | -21.4427 | 1.15E-06 | 3.67E-05 |
| Amigo3 | -11.6421 | 7.09E-06 | 0.000134 |
| Ankrd13d | -21.0213 | 1.86E-06 | 4.42E-05 |
| Ano1 | -22.0085 | 5.97E-07 | 3.21E-05 |
| Anxa1 | -22.2353 | 4.56E-07 | 3.21E-05 |
| Anxa5 | -11.1182 | 0.006207 | 0.093747 |

| | | | |
|---|---|---|---|
| Ap3m2 | -22.0024 | 6.01E-07 | 3.21E-05 |
| Apba1 | -22.5827 | 3.01E-07 | 2.83E-05 |
| Apc-ps1 | -8.33043 | 0.004162 | 0.065734 |
| Apc2 | -21.3587 | 1.27E-06 | 3.78E-05 |
| Apoc2 | -21.519 | 1.06E-06 | 3.65E-05 |
| Apold1 | -22.0337 | 5.79E-07 | 3.21E-05 |
| Apom | -21.0734 | 1.76E-06 | 4.27E-05 |
| Aqp11 | -20.8017 | 2.39E-06 | 5.03E-05 |
| Arhgap30 | -1.60148 | 0.000228 | 0.004008 |
| Arhgef10l | -21.4015 | 1.21E-06 | 3.75E-05 |
| Arhgef15 | 24.5138 | 1.48E-08 | 3.55E-06 |
| Arhgef40 | -21.0398 | 1.83E-06 | 4.36E-05 |
| Arl2bp | -22.0085 | 5.97E-07 | 3.21E-05 |
| Armc9 | -22.0626 | 5.60E-07 | 3.21E-05 |
| Armh4 | -21.3416 | 1.30E-06 | 3.78E-05 |
| Arntl2 | -11.986 | 6.77E-08 | 1.03E-05 |
| Asb4 | -21.6999 | 8.57E-07 | 3.55E-05 |
| Ascl2 | -21.0398 | 1.83E-06 | 4.36E-05 |
| Aspa | -11.5529 | 2.95E-11 | 3.33E-08 |
| Atat1 | -20.1092 | 5.12E-06 | 9.70E-05 |
| Atcay | -20.9742 | 1.97E-06 | 4.57E-05 |
| Atp10d | 12.06125 | 5.80E-08 | 9.34E-06 |
| Atp2b1 | -1.31314 | 0.006359 | 0.095607 |
| Awat2 | 23.04629 | 1.02E-07 | 1.27E-05 |
| Axin1 | -1.62666 | 7.18E-05 | 0.001296 |
| B130055M24Rik | -20.7444 | 2.55E-06 | 5.12E-05 |
| B3gnt5 | -4.0487 | 0.00021 | 0.0037 |
| B4galt2 | -21.5782 | 9.87E-07 | 3.60E-05 |
| B9d2 | -21.436 | 1.16E-06 | 3.69E-05 |
| BB287469 | -9.46328 | 0.004332 | 0.068226 |
| Bbs1 | -20.9158 | 2.11E-06 | 4.70E-05 |
| BC028528 | 24.50053 | 1.51E-08 | 3.55E-06 |
| BC029722 | -20.7755 | 2.47E-06 | 5.08E-05 |
| BC037039 | 10.57955 | 3.16E-07 | 2.96E-05 |
| BC051665 | -13.023 | 7.33E-09 | 2.00E-06 |
| BC080696 | -21.9504 | 6.39E-07 | 3.26E-05 |
| Bean1 | -22.2928 | 4.26E-07 | 3.21E-05 |
| Bfsp2 | -20.8989 | 2.15E-06 | 4.75E-05 |
| Bloc1s5 | -3.20944 | 0.004828 | 0.0752 |
| Blvra | -12.1686 | 5.03E-14 | 8.53E-10 |
| Bmp8a | -22.0044 | 6.00E-07 | 3.21E-05 |
| Boc | -22.176 | 4.90E-07 | 3.21E-05 |
| Btn1a1 | -22.0024 | 6.01E-07 | 3.21E-05 |
| Btn2a2 | -21.1041 | 1.70E-06 | 4.23E-05 |
| C030015A19Rik | -21.7712 | 7.89E-07 | 3.47E-05 |
| C1rl | -21.393 | 1.22E-06 | 3.76E-05 |
| C230014O12Rik | -21.0398 | 1.83E-06 | 4.36E-05 |
| C230096K16Rik | 23.1716 | 8.65E-08 | 1.20E-05 |

| | | | |
|---|---|---|---|
| C430049B03Rik | -11.7156 | 7.71E-07 | 3.47E-05 |
| C5ar2 | -21.355 | 1.28E-06 | 3.78E-05 |
| C920009B18Rik | -20.8795 | 2.20E-06 | 4.81E-05 |
| Cachd1 | -21.5116 | 1.07E-06 | 3.65E-05 |
| Cacna1b | -21.6374 | 9.22E-07 | 3.57E-05 |
| Cand2 | -21.4587 | 1.13E-06 | 3.65E-05 |
| Capn12 | -21.3845 | 1.23E-06 | 3.76E-05 |
| Car7 | -22.0801 | 5.48E-07 | 3.21E-05 |
| Caskin1 | -1.61021 | 0.001625 | 0.026914 |
| Cavin4 | -21.8883 | 6.87E-07 | 3.27E-05 |
| Cbr3 | -20.7332 | 2.59E-06 | 5.15E-05 |
| Ccdc120 | -21.011 | 1.89E-06 | 4.46E-05 |
| Ccdc125 | 11.25248 | 1.13E-06 | 3.65E-05 |
| Ccdc155 | -21.1527 | 1.60E-06 | 4.16E-05 |
| Ccdc169 | -21.5953 | 9.68E-07 | 3.60E-05 |
| Ccdc173 | -21.011 | 1.89E-06 | 4.46E-05 |
| Ccdc18 | -12.7335 | 0.00298 | 0.048139 |
| Ccdc189 | -20.9379 | 2.06E-06 | 4.64E-05 |
| Ccdc194 | -21.3587 | 1.27E-06 | 3.78E-05 |
| Ccdc24 | -20.8989 | 2.15E-06 | 4.75E-05 |
| Ccdc32 | -11.81 | 0.005471 | 0.083906 |
| Ccdc38 | 23.12681 | 9.16E-08 | 1.21E-05 |
| Ccdc69 | -22.3517 | 3.97E-07 | 3.21E-05 |
| Ccdc71l | -10.1577 | 2.04E-05 | 0.000375 |
| Ccdc82 | -22.1661 | 4.96E-07 | 3.21E-05 |
| Ccdc87 | -21.5315 | 1.04E-06 | 3.65E-05 |
| Cd164l2 | -21.0516 | 1.81E-06 | 4.34E-05 |
| Cd47 | -11.7771 | 0.006533 | 0.09746 |
| Cd55b | -20.7444 | 2.55E-06 | 5.12E-05 |
| Cd79b | -22.1246 | 5.20E-07 | 3.21E-05 |
| Cdh18 | -21.2212 | 1.49E-06 | 4.01E-05 |
| Cdh24 | -22.1708 | 4.92E-07 | 3.21E-05 |
| Cdh5 | -20.4855 | 3.37E-06 | 6.57E-05 |
| Cdkl3 | -22.1756 | 4.90E-07 | 3.21E-05 |
| Cdkl5 | -21.9864 | 6.13E-07 | 3.23E-05 |
| Cdr2l | -20.7624 | 2.50E-06 | 5.08E-05 |
| Cdx1 | -10.8413 | 4.92E-08 | 8.34E-06 |
| Celf2 | -13.2216 | 2.39E-11 | 2.92E-08 |
| Celsr3 | -21.7712 | 7.89E-07 | 3.47E-05 |
| Cep112 | -22.8428 | 2.19E-07 | 2.25E-05 |
| Cep128 | -11.3088 | 0.005688 | 0.086612 |
| Cercam | 23.42772 | 6.22E-08 | 9.72E-06 |
| Cfap299 | -22.5892 | 2.98E-07 | 2.83E-05 |
| Cfap44 | -20.4644 | 3.47E-06 | 6.73E-05 |
| Cfap46 | -22.0903 | 5.42E-07 | 3.21E-05 |
| Cfc1 | -21.1909 | 1.54E-06 | 4.04E-05 |
| Chrnb1 | -22.4233 | 3.64E-07 | 3.15E-05 |
| Chst4 | 11.9701 | 0.00015 | 0.002676 |

| | | | |
|---|---|---|---|
| Chsy1 | -11.8777 | 2.67E-09 | 1.03E-06 |
| Ciart | -21.5879 | 9.76E-07 | 3.60E-05 |
| Clcn5 | -1.78232 | 0.005604 | 0.085643 |
| Cldn13 | -22.2147 | 4.68E-07 | 3.21E-05 |
| Cldn20 | -22.63 | 2.84E-07 | 2.75E-05 |
| Clhc1 | -21.5715 | 9.95E-07 | 3.60E-05 |
| Cln3 | 1.992138 | 2.93E-05 | 0.000535 |
| Clspn | -1.33615 | 0.000179 | 0.003165 |
| Clu | -21.7328 | 8.25E-07 | 3.55E-05 |
| Cmc4 | -20.7961 | 2.41E-06 | 5.03E-05 |
| Cnga1 | -20.32 | 4.06E-06 | 7.73E-05 |
| Cnksr1 | -11.2964 | 0.005668 | 0.086381 |
| Cnn2 | -26.659 | 1.46E-09 | 6.51E-07 |
| Cnot7 | -1.85755 | 9.68E-05 | 0.001737 |
| Cnpy1 | -11.566 | 0.005413 | 0.083313 |
| Coprs | 1.520996 | 1.92E-06 | 4.49E-05 |
| Coq10b | -2.15434 | 1.96E-07 | 2.06E-05 |
| Coq8a | 9.7546 | 4.15E-05 | 0.000753 |
| Cpeb2 | -1.39884 | 0.003221 | 0.051794 |
| Cpeb4 | -11.8068 | 4.41E-10 | 2.99E-07 |
| Crppa | -21.3479 | 1.29E-06 | 3.78E-05 |
| Ctsd | 1.231318 | 0.000234 | 0.004116 |
| Cttnbp2 | -21.7457 | 8.12E-07 | 3.51E-05 |
| Cxcl10 | -21.3143 | 1.34E-06 | 3.83E-05 |
| Cyfip2 | -22.1096 | 5.30E-07 | 3.21E-05 |
| Cyp1a1 | -22.0506 | 5.68E-07 | 3.21E-05 |
| Cyp26b1 | -20.9158 | 2.11E-06 | 4.70E-05 |
| Cyp2c67 | -21.0516 | 1.81E-06 | 4.34E-05 |
| Cyp2d9 | 23.00607 | 1.07E-07 | 1.28E-05 |
| Cyp2s1 | -3.96653 | 2.98E-06 | 5.85E-05 |
| Cyp4f13 | -11.2288 | 0.006428 | 0.096236 |
| D030028A08Rik | -21.1135 | 1.68E-06 | 4.23E-05 |
| D130017N08Rik | -20.7755 | 2.47E-06 | 5.08E-05 |
| D630023O14Rik | -20.807 | 2.38E-06 | 5.03E-05 |
| D730003I15Rik | -21.2957 | 1.37E-06 | 3.87E-05 |
| D930028M14Rik | -21.0808 | 1.75E-06 | 4.27E-05 |
| Dapk1 | -2.00309 | 0.000814 | 0.013821 |
| Dcst1 | -21.4829 | 1.10E-06 | 3.65E-05 |
| Ddx43 | -21.9724 | 6.23E-07 | 3.26E-05 |
| Ddx60 | -21.6387 | 9.20E-07 | 3.57E-05 |
| Defb42 | -21.2228 | 1.49E-06 | 4.01E-05 |
| Degs2 | -21.1254 | 1.66E-06 | 4.23E-05 |
| Derl3 | -21.3416 | 1.30E-06 | 3.78E-05 |
| Dglucy | -6.88612 | 0.005589 | 0.08549 |
| Dhh | -20.7444 | 2.55E-06 | 5.12E-05 |
| Dhrs4 | -11.7628 | 0.004754 | 0.074182 |
| Diras1 | -20.9509 | 2.03E-06 | 4.62E-05 |
| Dkk1 | -12.9638 | 3.33E-09 | 1.18E-06 |

| | | | |
|---|---|---|---|
| Dlgap2 | -21.2544 | 1.43E-06 | 3.98E-05 |
| Dmkn | -7.30647 | 0.000529 | 0.009104 |
| Dmtf1 | 1.356652 | 0.000343 | 0.005962 |
| Dnah10 | 25.31595 | 4.94E-09 | 1.55E-06 |
| Dnajc4 | -11.0674 | 2.33E-05 | 0.000427 |
| Dnhd1 | -11.8153 | 1.56E-06 | 4.07E-05 |
| Dnm3 | -20.7444 | 2.55E-06 | 5.12E-05 |
| Dnmt3b | -1.79004 | 4.45E-05 | 0.000807 |
| Doc2a | -21.7712 | 7.89E-07 | 3.47E-05 |
| Dock2 | -21.9222 | 6.61E-07 | 3.26E-05 |
| Dock3 | -20.7755 | 2.47E-06 | 5.08E-05 |
| Dock4 | -3.40127 | 1.67E-05 | 0.000309 |
| Dqx1 | -22.0553 | 5.65E-07 | 3.21E-05 |
| Dsel | -21.6311 | 9.29E-07 | 3.58E-05 |
| Dstyk | -21.011 | 1.89E-06 | 4.46E-05 |
| Dtx3 | -21.5575 | 1.01E-06 | 3.65E-05 |
| DXBay18 | -2.99087 | 0.000226 | 0.003976 |
| E030044B06Rik | -20.8795 | 2.20E-06 | 4.81E-05 |
| E130215H24Rik | -22.1397 | 5.11E-07 | 3.21E-05 |
| E330014E10Rik | 25.39997 | 4.40E-09 | 1.43E-06 |
| Ebag9 | -11.9882 | 0.003639 | 0.057911 |
| Ebf4 | -10.6456 | 0.000125 | 0.002232 |
| Ech1 | -2.17655 | 0.005273 | 0.081381 |
| Echdc3 | -21.7775 | 7.83E-07 | 3.47E-05 |
| Eci3 | -21.9994 | 6.03E-07 | 3.21E-05 |
| Edaradd | -21.4748 | 1.11E-06 | 3.65E-05 |
| Eepd1 | -10.9461 | 3.99E-06 | 7.63E-05 |
| Efcab12 | -19.5919 | 8.88E-06 | 0.000166 |
| Efhb | -21.6865 | 8.71E-07 | 3.55E-05 |
| Ehd2 | -21.6865 | 8.71E-07 | 3.55E-05 |
| Ehd3 | -21.9816 | 6.16E-07 | 3.24E-05 |
| Eid1 | -10.2608 | 1.19E-07 | 1.39E-05 |
| Eif2ak2 | -20.3584 | 3.89E-06 | 7.47E-05 |
| Eif2ak4 | -12.5292 | 0.003298 | 0.052881 |
| Eif5a2 | -21.6865 | 8.71E-07 | 3.55E-05 |
| Elavl3 | -22.0334 | 5.79E-07 | 3.21E-05 |
| Elf5 | -13.0084 | 1.11E-07 | 1.32E-05 |
| Elmod1 | -22.0593 | 5.62E-07 | 3.21E-05 |
| Emc3 | 1.202358 | 0.002756 | 0.044647 |
| Entpd1 | -10.3521 | 2.68E-07 | 2.67E-05 |
| Entpd3 | 11.54731 | 3.71E-06 | 7.16E-05 |
| Ephb2 | -11.5991 | 3.45E-09 | 1.19E-06 |
| Ephb6 | -21.0516 | 1.81E-06 | 4.34E-05 |
| Epn1 | -7.75859 | 1.05E-08 | 2.67E-06 |
| Erich3 | -20.9509 | 2.03E-06 | 4.62E-05 |
| Esr1 | -21.0015 | 1.91E-06 | 4.49E-05 |
| Etohd2 | 22.54947 | 1.89E-07 | 2.06E-05 |
| Ets1 | -21.9676 | 6.26E-07 | 3.26E-05 |

| | | | |
|---|---|---|---|
| F3 | -21.2049 | 1.52E-06 | 4.02E-05 |
| F730035P03Rik | -21.3673 | 1.26E-06 | 3.78E-05 |
| Fads2 | -21.9994 | 6.03E-07 | 3.21E-05 |
| Fads3 | -22.0414 | 5.74E-07 | 3.21E-05 |
| Fam126b | -11.4138 | 1.04E-07 | 1.27E-05 |
| Fam129c | 23.25291 | 7.79E-08 | 1.13E-05 |
| Fam131c | 25.01929 | 7.45E-09 | 2.00E-06 |
| Fam151b | -21.3759 | 1.25E-06 | 3.76E-05 |
| Fam160a1 | -11.8786 | 2.12E-11 | 2.92E-08 |
| Fam189a2 | -21.4983 | 1.08E-06 | 3.65E-05 |
| Fam43a | -6.64034 | 0.000802 | 0.013657 |
| Fam71e1 | -21.7747 | 7.85E-07 | 3.47E-05 |
| Fam71f2 | 24.95153 | 8.17E-09 | 2.10E-06 |
| Fam81a | -21.3245 | 1.32E-06 | 3.81E-05 |
| Fbxl8 | -22.6429 | 2.80E-07 | 2.74E-05 |
| Fbxo10 | -21.5416 | 1.03E-06 | 3.65E-05 |
| Fbxo17 | -10.9883 | 1.23E-06 | 3.76E-05 |
| Fcor | -21.6961 | 8.61E-07 | 3.55E-05 |
| Fdps | 1.783248 | 0.000541 | 0.009287 |
| Fgf2 | -21.2212 | 1.49E-06 | 4.01E-05 |
| Fgf7 | -22.2142 | 4.68E-07 | 3.21E-05 |
| Fjx1 | -9.99151 | 7.81E-08 | 1.13E-05 |
| Fkbp1b | -22.1756 | 4.90E-07 | 3.21E-05 |
| Flywch2 | -20.7863 | 2.44E-06 | 5.06E-05 |
| Fmnl3 | -1.96873 | 0.002939 | 0.047563 |
| Fnbp1l | -1.38239 | 2.25E-05 | 0.000413 |
| Foxf1 | -21.6999 | 8.57E-07 | 3.55E-05 |
| Fpgt | -11.1554 | 0.006066 | 0.092028 |
| Fthl17b | 7.481958 | 0.00493 | 0.076578 |
| Fut2 | -20.7961 | 2.41E-06 | 5.03E-05 |
| Fut4 | -20.7755 | 2.47E-06 | 5.08E-05 |
| Fxyd6 | -11.92 | 1.44E-09 | 6.51E-07 |
| Fyn | -22.8834 | 2.09E-07 | 2.16E-05 |
| Fzd6 | -21.519 | 1.06E-06 | 3.65E-05 |
| Gab3 | 23.16072 | 8.77E-08 | 1.20E-05 |
| Gak | -1.21518 | 0.001168 | 0.019558 |
| Galnt10 | -10.7369 | 3.34E-09 | 1.18E-06 |
| Gata2 | -6.45113 | 0.004787 | 0.074633 |
| Gbe1 | -21.9035 | 6.75E-07 | 3.26E-05 |
| Gc | -21.0757 | 1.76E-06 | 4.27E-05 |
| Gda | -21.603 | 9.59E-07 | 3.60E-05 |
| Gjc1 | -21.4983 | 1.08E-06 | 3.65E-05 |
| Gk | -11.4172 | 0.005384 | 0.082948 |
| Glrx | -4.03332 | 0.004684 | 0.073225 |
| Gm10353 | -22.3078 | 4.18E-07 | 3.21E-05 |
| Gm10509 | -21.3143 | 1.34E-06 | 3.83E-05 |
| Gm10651 | -6.76321 | 0.001355 | 0.022608 |
| Gm10718 | -21.8248 | 7.41E-07 | 3.44E-05 |

| | | | |
|---|---|---|---|
| Gm10800 | -21.1143 | 1.68E-06 | 4.23E-05 |
| Gm10873 | 24.49989 | 1.51E-08 | 3.55E-06 |
| Gm11222 | 2.907884 | 0.002551 | 0.041531 |
| Gm11236 | -22.176 | 4.90E-07 | 3.21E-05 |
| Gm11237 | -21.6642 | 8.93E-07 | 3.57E-05 |
| Gm11238 | -21.5514 | 1.02E-06 | 3.65E-05 |
| Gm11239 | -21.6932 | 8.64E-07 | 3.55E-05 |
| Gm11510 | -22.2229 | 4.63E-07 | 3.21E-05 |
| Gm11543 | -20.5873 | 3.04E-06 | 5.94E-05 |
| Gm11935 | -21.2107 | 1.51E-06 | 4.02E-05 |
| Gm12169 | -5.01838 | 0.00088 | 0.014935 |
| Gm12184 | 1.345714 | 0.000551 | 0.009448 |
| Gm12569 | -21.8446 | 7.24E-07 | 3.40E-05 |
| Gm12576 | -22.0198 | 5.89E-07 | 3.21E-05 |
| Gm12594 | 11.10325 | 2.05E-07 | 2.13E-05 |
| Gm12596 | -21.4748 | 1.11E-06 | 3.65E-05 |
| Gm12616 | 3.842361 | 0.004092 | 0.064684 |
| Gm12621 | 1.687421 | 0.001477 | 0.024511 |
| Gm12625 | 1.314502 | 0.001211 | 0.020246 |
| Gm12758 | -11.3831 | 2.36E-07 | 2.39E-05 |
| Gm12766 | -20.6196 | 2.92E-06 | 5.76E-05 |
| Gm12992 | -13.3275 | 0.001906 | 0.031368 |
| Gm12996 | -20.8017 | 2.39E-06 | 5.03E-05 |
| Gm13032 | -19.5919 | 8.88E-06 | 0.000166 |
| Gm13068 | -21.5511 | 1.02E-06 | 3.65E-05 |
| Gm13101 | -21.3845 | 1.23E-06 | 3.76E-05 |
| Gm13110 | -21.4587 | 1.13E-06 | 3.65E-05 |
| Gm13228 | -22.2937 | 4.26E-07 | 3.21E-05 |
| Gm13360 | -21.4829 | 1.10E-06 | 3.65E-05 |
| Gm13548 | -20.9509 | 2.03E-06 | 4.62E-05 |
| Gm13611 | 1.235172 | 0.001749 | 0.028862 |
| Gm13830 | -20.3584 | 3.89E-06 | 7.47E-05 |
| Gm1401 | -21.1995 | 1.53E-06 | 4.04E-05 |
| Gm14127 | -21.6932 | 8.64E-07 | 3.55E-05 |
| Gm14383 | 3.07747 | 0.00041 | 0.0071 |
| Gm14411 | -7.87597 | 0.000149 | 0.002659 |
| Gm14451 | -22.6575 | 2.75E-07 | 2.71E-05 |
| Gm14452 | -22.3252 | 4.09E-07 | 3.21E-05 |
| Gm14549 | -21.4931 | 1.09E-06 | 3.65E-05 |
| Gm15401 | -21.6999 | 8.57E-07 | 3.55E-05 |
| Gm15408 | 25.0463 | 7.17E-09 | 2.00E-06 |
| Gm15442 | -21.9041 | 6.75E-07 | 3.26E-05 |
| Gm15455 | -4.0588 | 0.000205 | 0.003624 |
| Gm15583 | -1.53234 | 0.000145 | 0.002589 |
| Gm15608 | -21.9144 | 6.67E-07 | 3.26E-05 |
| Gm15720 | 4.443704 | 0.006405 | 0.096054 |
| Gm15728 | -21.9887 | 6.11E-07 | 3.23E-05 |
| Gm15803 | -10.4655 | 3.89E-08 | 7.02E-06 |

| | | | |
|---|---|---|---|
| Gm15893 | -22.775 | 2.38E-07 | 2.39E-05 |
| Gm15903 | -21.0927 | 1.72E-06 | 4.27E-05 |
| Gm15964 | 3.143956 | 9.79E-10 | 5.18E-07 |
| Gm15998 | -22.3722 | 3.87E-07 | 3.21E-05 |
| Gm16001 | -21.2049 | 1.52E-06 | 4.02E-05 |
| Gm16089 | 2.567495 | 0.001016 | 0.017092 |
| Gm16133 | -20.7444 | 2.55E-06 | 5.12E-05 |
| Gm16153 | -21.393 | 1.22E-06 | 3.76E-05 |
| Gm16201 | -21.3433 | 1.29E-06 | 3.78E-05 |
| Gm16332 | -21.0757 | 1.76E-06 | 4.27E-05 |
| Gm17041 | -20.8483 | 2.27E-06 | 4.93E-05 |
| Gm17103 | -21.4015 | 1.21E-06 | 3.75E-05 |
| Gm17200 | -22.1189 | 5.24E-07 | 3.21E-05 |
| Gm17809 | -22.3517 | 3.97E-07 | 3.21E-05 |
| Gm17981 | 24.68242 | 1.18E-08 | 2.94E-06 |
| Gm18273 | -22.2769 | 4.34E-07 | 3.21E-05 |
| Gm18646 | -3.2959 | 0.003358 | 0.053738 |
| Gm18760 | -21.4587 | 1.13E-06 | 3.65E-05 |
| Gm18914 | -21.0734 | 1.76E-06 | 4.27E-05 |
| Gm19040 | -20.6881 | 2.71E-06 | 5.37E-05 |
| Gm19299 | -21.2348 | 1.47E-06 | 4.01E-05 |
| Gm19426 | 23.68152 | 4.47E-08 | 7.74E-06 |
| Gm19637 | -20.8378 | 2.30E-06 | 4.98E-05 |
| Gm2022 | -8.79385 | 0.006421 | 0.096219 |
| Gm20234 | -5.14989 | 0.003663 | 0.058241 |
| Gm20324 | -21.0734 | 1.76E-06 | 4.27E-05 |
| Gm20457 | 23.42381 | 6.25E-08 | 9.72E-06 |
| Gm20498 | -12.0686 | 1.23E-05 | 0.000228 |
| Gm20517 | -20.9158 | 2.11E-06 | 4.70E-05 |
| Gm20518 | -21.1135 | 1.68E-06 | 4.23E-05 |
| Gm20611 | -20.8219 | 2.34E-06 | 5.01E-05 |
| Gm20625 | 2.598364 | 0.002071 | 0.033979 |
| Gm20683 | 12.7094 | 2.89E-10 | 2.22E-07 |
| Gm20731 | -9.60731 | 0.000719 | 0.012299 |
| Gm21742 | -20.8538 | 2.26E-06 | 4.90E-05 |
| Gm21885 | -21.72 | 8.37E-07 | 3.55E-05 |
| Gm21986 | -20.789 | 2.43E-06 | 5.05E-05 |
| Gm22009 | -21.1079 | 1.69E-06 | 4.23E-05 |
| Gm22204 | -21.0475 | 1.81E-06 | 4.36E-05 |
| Gm22770 | 23.16072 | 8.77E-08 | 1.20E-05 |
| Gm23246 | 23.83537 | 3.66E-08 | 6.74E-06 |
| Gm23969 | 10.69374 | 9.12E-06 | 0.00017 |
| Gm2415 | 9.973696 | 3.31E-05 | 0.000604 |
| Gm24201 | 10.69983 | 0.00023 | 0.004047 |
| Gm24411 | -21.1254 | 1.66E-06 | 4.23E-05 |
| Gm24507 | -21.355 | 1.28E-06 | 3.78E-05 |
| Gm24523 | -21.9222 | 6.61E-07 | 3.26E-05 |
| Gm24644 | -21.7478 | 8.11E-07 | 3.51E-05 |

| | | | |
|---|---|---|---|
| Gm25580 | 24.36076 | 1.82E-08 | 4.06E-06 |
| Gm25835 | 10.06225 | 0.000752 | 0.012848 |
| Gm25894 | 23.92887 | 3.23E-08 | 6.22E-06 |
| Gm26254 | 24.39212 | 1.75E-08 | 3.94E-06 |
| Gm26397 | -19.5919 | 8.88E-06 | 0.000166 |
| Gm26616 | -10.9004 | 0.000154 | 0.002742 |
| Gm26660 | -21.3502 | 1.28E-06 | 3.78E-05 |
| Gm26770 | -21.8111 | 7.53E-07 | 3.44E-05 |
| Gm26789 | -22.1708 | 4.92E-07 | 3.21E-05 |
| Gm26800 | -22.418 | 3.66E-07 | 3.15E-05 |
| Gm26811 | -9.02662 | 0.000416 | 0.007187 |
| Gm26877 | -20.7961 | 2.41E-06 | 5.03E-05 |
| Gm26943 | -22.0667 | 5.57E-07 | 3.21E-05 |
| Gm27017 | -20.8219 | 2.34E-06 | 5.01E-05 |
| Gm27042 | -21.4507 | 1.14E-06 | 3.66E-05 |
| Gm27861 | -20.9742 | 1.97E-06 | 4.57E-05 |
| Gm28535 | -21.1041 | 1.70E-06 | 4.23E-05 |
| Gm28551 | 27.60117 | 1.80E-10 | 1.53E-07 |
| Gm29284 | 10.43061 | 2.95E-06 | 5.81E-05 |
| Gm29290 | -20.4263 | 3.60E-06 | 6.96E-05 |
| Gm29642 | -22.4699 | 3.45E-07 | 3.09E-05 |
| Gm29695 | -20.8678 | 2.22E-06 | 4.86E-05 |
| Gm2a | -1.7818 | 0.001444 | 0.023995 |
| Gm30414 | -21.7393 | 8.19E-07 | 3.53E-05 |
| Gm31503 | -20.7631 | 2.50E-06 | 5.08E-05 |
| Gm32341 | -21.6244 | 9.36E-07 | 3.58E-05 |
| Gm33280 | -21.6416 | 9.17E-07 | 3.57E-05 |
| Gm35167 | -21.1041 | 1.70E-06 | 4.23E-05 |
| Gm35256 | -21.4587 | 1.13E-06 | 3.65E-05 |
| Gm3650 | -21.3331 | 1.31E-06 | 3.80E-05 |
| Gm36638 | -20.6196 | 2.92E-06 | 5.76E-05 |
| Gm36933 | -21.2544 | 1.43E-06 | 3.98E-05 |
| Gm37008 | -20.7961 | 2.41E-06 | 5.03E-05 |
| Gm37189 | -20.7755 | 2.47E-06 | 5.08E-05 |
| Gm37411 | -21.8081 | 7.55E-07 | 3.44E-05 |
| Gm37436 | -21.6709 | 8.87E-07 | 3.56E-05 |
| Gm37850 | -20.9216 | 2.09E-06 | 4.70E-05 |
| Gm37933 | 23.04217 | 1.02E-07 | 1.27E-05 |
| Gm37939 | -11.0938 | 1.15E-07 | 1.36E-05 |
| Gm38114 | -21.2874 | 1.38E-06 | 3.87E-05 |
| Gm38253 | -21.603 | 9.59E-07 | 3.60E-05 |
| Gm38329 | -21.2319 | 1.47E-06 | 4.01E-05 |
| Gm38372 | -11.2667 | 3.72E-05 | 0.000678 |
| Gm38375 | -21.0808 | 1.75E-06 | 4.27E-05 |
| Gm38387 | -21.2544 | 1.43E-06 | 3.98E-05 |
| Gm38403 | -21.7131 | 8.44E-07 | 3.55E-05 |
| Gm39090 | -21.1532 | 1.61E-06 | 4.16E-05 |
| Gm39271 | -23.0084 | 1.79E-07 | 1.98E-05 |

| | | | |
|---|---|---|---|
| Gm4013 | -21.6387 | 9.20E-07 | 3.57E-05 |
| Gm40557 | -20.32 | 4.06E-06 | 7.73E-05 |
| Gm41609 | -22.1527 | 5.03E-07 | 3.21E-05 |
| Gm42432 | -21.6416 | 9.17E-07 | 3.57E-05 |
| Gm42572 | -22.0001 | 6.03E-07 | 3.21E-05 |
| Gm42658 | -20.7624 | 2.50E-06 | 5.08E-05 |
| Gm42662 | -21.6387 | 9.20E-07 | 3.57E-05 |
| Gm42671 | -20.7961 | 2.41E-06 | 5.03E-05 |
| Gm42690 | -21.4667 | 1.12E-06 | 3.65E-05 |
| Gm42717 | -21.2242 | 1.48E-06 | 4.01E-05 |
| Gm428 | -21.5116 | 1.07E-06 | 3.65E-05 |
| Gm42951 | -22.503 | 3.31E-07 | 3.03E-05 |
| Gm43075 | -20.8947 | 2.16E-06 | 4.76E-05 |
| Gm43254 | -21.2412 | 1.46E-06 | 4.00E-05 |
| Gm43511 | -7.5748 | 0.000107 | 0.001923 |
| Gm43513 | -21.2891 | 1.38E-06 | 3.87E-05 |
| Gm43661 | -21.2957 | 1.37E-06 | 3.87E-05 |
| Gm44008 | -21.393 | 1.22E-06 | 3.76E-05 |
| Gm44037 | -22.9458 | 1.93E-07 | 2.06E-05 |
| Gm44116 | -21.6416 | 9.17E-07 | 3.57E-05 |
| Gm44165 | -20.32 | 4.06E-06 | 7.73E-05 |
| Gm44198 | -20.9158 | 2.11E-06 | 4.70E-05 |
| Gm44260 | 25.32043 | 4.91E-09 | 1.55E-06 |
| Gm44415 | -21.4057 | 1.21E-06 | 3.75E-05 |
| Gm44567 | -20.7332 | 2.59E-06 | 5.15E-05 |
| Gm44597 | -12.1192 | 0.004837 | 0.075277 |
| Gm44616 | -22.0701 | 5.55E-07 | 3.21E-05 |
| Gm44709 | -21.2891 | 1.38E-06 | 3.87E-05 |
| Gm44736 | -5.73338 | 0.001895 | 0.031215 |
| Gm44751 | 25.87 | 2.27E-09 | 9.38E-07 |
| Gm44867 | -21.3845 | 1.23E-06 | 3.76E-05 |
| Gm45051 | -21.7131 | 8.44E-07 | 3.55E-05 |
| Gm45058 | -21.6594 | 8.98E-07 | 3.57E-05 |
| Gm45203 | -20.8678 | 2.22E-06 | 4.86E-05 |
| Gm45293 | -21.712 | 8.45E-07 | 3.55E-05 |
| Gm45412 | -22.0658 | 5.58E-07 | 3.21E-05 |
| Gm45494 | -21.4587 | 1.13E-06 | 3.65E-05 |
| Gm45518 | -21.5879 | 9.76E-07 | 3.60E-05 |
| Gm45592 | -21.9041 | 6.75E-07 | 3.26E-05 |
| Gm45718 | -22.0188 | 5.90E-07 | 3.21E-05 |
| Gm45809 | -22.7743 | 2.39E-07 | 2.39E-05 |
| Gm45845 | -22.6027 | 2.94E-07 | 2.81E-05 |
| Gm45861 | -21.5744 | 9.87E-07 | 3.60E-05 |
| Gm45864 | -21.4427 | 1.15E-06 | 3.67E-05 |
| Gm45894 | -10.7081 | 7.12E-08 | 1.07E-05 |
| Gm46555 | 23.59313 | 5.02E-08 | 8.42E-06 |
| Gm46560 | -21.899 | 6.79E-07 | 3.27E-05 |
| Gm4675 | -22.1114 | 5.28E-07 | 3.21E-05 |

| | | | |
|---|---|---|---|
| Gm46996 | -20.9509 | 2.03E-06 | 4.62E-05 |
| Gm47198 | -21.4507 | 1.14E-06 | 3.66E-05 |
| Gm47233 | -22.4656 | 3.46E-07 | 3.09E-05 |
| Gm47251 | -21.5782 | 9.87E-07 | 3.60E-05 |
| Gm47321 | -21.4983 | 1.08E-06 | 3.65E-05 |
| Gm47468 | -21.0547 | 1.80E-06 | 4.34E-05 |
| Gm47585 | -22.1314 | 5.16E-07 | 3.21E-05 |
| Gm47640 | -22.1232 | 5.21E-07 | 3.21E-05 |
| Gm47995 | -20.8989 | 2.15E-06 | 4.75E-05 |
| Gm47996 | -21.6374 | 9.22E-07 | 3.57E-05 |
| Gm48353 | -22.3 | 4.22E-07 | 3.21E-05 |
| Gm48420 | -21.7823 | 7.78E-07 | 3.47E-05 |
| Gm48478 | -21.0398 | 1.83E-06 | 4.36E-05 |
| Gm48551 | -22.249 | 4.49E-07 | 3.21E-05 |
| Gm48609 | -21.2348 | 1.47E-06 | 4.01E-05 |
| Gm48641 | -21.1259 | 1.66E-06 | 4.23E-05 |
| Gm48826 | -20.7755 | 2.47E-06 | 5.08E-05 |
| Gm49027 | -22.2928 | 4.26E-07 | 3.21E-05 |
| Gm49052 | -21.7775 | 7.83E-07 | 3.47E-05 |
| Gm49085 | 12.20842 | 8.74E-07 | 3.55E-05 |
| Gm49132 | -22.4032 | 3.73E-07 | 3.16E-05 |
| Gm49156 | -21.011 | 1.89E-06 | 4.46E-05 |
| Gm49204 | -21.2212 | 1.49E-06 | 4.01E-05 |
| Gm49284 | -20.1092 | 5.12E-06 | 9.70E-05 |
| Gm4930 | -20.9509 | 2.03E-06 | 4.62E-05 |
| Gm49327 | -20.8122 | 2.36E-06 | 5.03E-05 |
| Gm49357 | -20.7332 | 2.59E-06 | 5.15E-05 |
| Gm49417 | -21.1532 | 1.61E-06 | 4.16E-05 |
| Gm49455 | -21.1254 | 1.66E-06 | 4.23E-05 |
| Gm49486 | 13.03051 | 3.89E-08 | 7.02E-06 |
| Gm49673 | 26.2935 | 1.24E-09 | 6.00E-07 |
| Gm49721 | 26.23389 | 1.35E-09 | 6.35E-07 |
| Gm49767 | -21.5803 | 9.81E-07 | 3.60E-05 |
| Gm4982 | -21.3143 | 1.34E-06 | 3.83E-05 |
| Gm49870 | -21.3245 | 1.32E-06 | 3.81E-05 |
| Gm49909 | 26.81028 | 5.85E-10 | 3.33E-07 |
| Gm49932 | -21.4748 | 1.11E-06 | 3.65E-05 |
| Gm49937 | -21.2172 | 1.50E-06 | 4.02E-05 |
| Gm49960 | -22.5986 | 2.95E-07 | 2.81E-05 |
| Gm49987 | -21.7478 | 8.11E-07 | 3.51E-05 |
| Gm50041 | -5.2893 | 0.00547 | 0.083906 |
| Gm50053 | -20.9379 | 2.06E-06 | 4.64E-05 |
| Gm50092 | -22.074 | 5.52E-07 | 3.21E-05 |
| Gm50115 | -8.86884 | 5.37E-07 | 3.21E-05 |
| Gm50179 | -20.8017 | 2.39E-06 | 5.03E-05 |
| Gm50180 | -21.1909 | 1.54E-06 | 4.04E-05 |
| Gm50281 | -21.0015 | 1.91E-06 | 4.49E-05 |
| Gm5049 | -20.9742 | 1.97E-06 | 4.57E-05 |

| | | | |
|---|---|---|---|
| Gm5127 | -22.0446 | 5.72E-07 | 3.21E-05 |
| Gm5251 | -22.1096 | 5.30E-07 | 3.21E-05 |
| Gm5302 | 3.560404 | 0.0012 | 0.020069 |
| Gm5485 | 25.75864 | 2.66E-09 | 1.03E-06 |
| Gm5616 | 2.204475 | 0.004674 | 0.073145 |
| Gm5791 | 1.796106 | 0.004847 | 0.075357 |
| Gm5834 | -21.2957 | 1.37E-06 | 3.87E-05 |
| Gm5890 | -21.5416 | 1.03E-06 | 3.65E-05 |
| Gm5917 | -2.1556 | 0.001718 | 0.028369 |
| Gm5951 | -21.6532 | 9.05E-07 | 3.57E-05 |
| Gm6087 | -3.87677 | 0.002144 | 0.035107 |
| Gm6312 | -20.7624 | 2.50E-06 | 5.08E-05 |
| Gm6351 | -21.1532 | 1.61E-06 | 4.16E-05 |
| Gm6436 | -9.93735 | 3.13E-09 | 1.15E-06 |
| Gm6509 | -21.4265 | 1.18E-06 | 3.69E-05 |
| Gm6899 | 10.73482 | 0.006086 | 0.092174 |
| Gm6967 | 10.30697 | 0.001106 | 0.018602 |
| Gm7008 | -21.0398 | 1.83E-06 | 4.36E-05 |
| Gm7244 | -20.7332 | 2.59E-06 | 5.15E-05 |
| Gm7265 | -22.3316 | 4.07E-07 | 3.21E-05 |
| Gm7539 | -21.1909 | 1.54E-06 | 4.04E-05 |
| Gm7600 | 2.308333 | 8.06E-06 | 0.000152 |
| Gm7647 | -22.219 | 4.65E-07 | 3.21E-05 |
| Gm7663 | -22.0231 | 5.87E-07 | 3.21E-05 |
| Gm7682 | -21.1254 | 1.66E-06 | 4.23E-05 |
| Gm7889 | -20.8017 | 2.39E-06 | 5.03E-05 |
| Gm7896 | 3.790069 | 0.001652 | 0.027337 |
| Gm7939 | -21.2874 | 1.38E-06 | 3.87E-05 |
| Gm7942 | -20.5364 | 3.19E-06 | 6.22E-05 |
| Gm7982 | -20.9277 | 2.08E-06 | 4.67E-05 |
| Gm807 | -22.1246 | 5.20E-07 | 3.21E-05 |
| Gm8689 | -21.8116 | 7.52E-07 | 3.44E-05 |
| Gm8701 | -21.1909 | 1.54E-06 | 4.04E-05 |
| Gm8775 | -5.46441 | 0.000796 | 0.013577 |
| Gm8909 | 23.11762 | 9.27E-08 | 1.21E-05 |
| Gm9025 | -21.1154 | 1.68E-06 | 4.23E-05 |
| Gm9117 | -22.0188 | 5.90E-07 | 3.21E-05 |
| Gm9260 | 3.596926 | 0.002317 | 0.037864 |
| Gm9285 | -20.7961 | 2.41E-06 | 5.03E-05 |
| Gm9519 | -21.6999 | 8.57E-07 | 3.55E-05 |
| Gm9531 | 1.829098 | 0.001976 | 0.032455 |
| Gm9621 | -21.519 | 1.06E-06 | 3.65E-05 |
| Gm9732 | -20.7624 | 2.50E-06 | 5.08E-05 |
| Gm9785 | -22.0701 | 5.55E-07 | 3.21E-05 |
| Gm9908 | 24.16568 | 2.36E-08 | 4.99E-06 |
| Gm9978 | -21.4265 | 1.18E-06 | 3.69E-05 |
| Gorasp2 | -2.99061 | 1.07E-05 | 0.000198 |
| Gpld1 | -21.1759 | 1.57E-06 | 4.10E-05 |

| | | | |
|---|---|---|---|
| Gpr137c | -21.8484 | 7.21E-07 | 3.39E-05 |
| Gpr180 | -11.4752 | 0.006227 | 0.09389 |
| Gpr87 | -21.476 | 1.11E-06 | 3.65E-05 |
| Gprasp1 | -2.2444 | 0.000261 | 0.00457 |
| Gprin2 | -21.9222 | 6.61E-07 | 3.26E-05 |
| Gpsm3 | -22.4705 | 3.44E-07 | 3.09E-05 |
| Gramd1a | -21.4587 | 1.13E-06 | 3.65E-05 |
| Gramd1c | -21.8049 | 7.58E-07 | 3.44E-05 |
| Gramd3 | -21.8037 | 7.59E-07 | 3.44E-05 |
| Grhl1 | -11.6784 | 2.12E-13 | 8.98E-10 |
| Grik3 | -9.88252 | 4.32E-06 | 8.21E-05 |
| Grin3a | -11.5597 | 4.85E-07 | 3.21E-05 |
| Grm4 | -11.287 | 0.006315 | 0.095048 |
| Gsta4 | -1.99115 | 0.005439 | 0.083646 |
| Gstk1 | -12.7907 | 0.002637 | 0.042854 |
| Gtf2ird1 | -1.4193 | 0.00464 | 0.072667 |
| Gulp1 | -21.537 | 1.04E-06 | 3.65E-05 |
| H19 | -21.4845 | 1.10E-06 | 3.65E-05 |
| H2-T22 | -22.1232 | 5.21E-07 | 3.21E-05 |
| H2bc21 | -21.4182 | 1.19E-06 | 3.71E-05 |
| H3c2 | -8.25743 | 0.00069 | 0.011803 |
| H3c4 | -8.39503 | 0.000239 | 0.004187 |
| Haghl | -20.8538 | 2.26E-06 | 4.90E-05 |
| Haus8 | -2.55525 | 0.00041 | 0.007102 |
| Hcrtr1 | -21.9041 | 6.75E-07 | 3.26E-05 |
| Hdc | 23.02954 | 1.04E-07 | 1.27E-05 |
| Hddc3 | 23.54588 | 5.34E-08 | 8.87E-06 |
| Hdhd2 | -12.6668 | 0.002546 | 0.041493 |
| Herpud1 | -12.902 | 1.20E-09 | 6.00E-07 |
| Hmgcll1 | -21.3845 | 1.23E-06 | 3.76E-05 |
| Hoxa13 | -20.8989 | 2.15E-06 | 4.75E-05 |
| Hoxb13 | 23.31671 | 7.18E-08 | 1.07E-05 |
| Hpse | -20.5364 | 3.19E-06 | 6.22E-05 |
| Hs3st5 | 24.56734 | 1.38E-08 | 3.39E-06 |
| Hs6st3 | -22.2353 | 4.56E-07 | 3.21E-05 |
| Hspa13 | -9.91356 | 2.35E-07 | 2.39E-05 |
| Hspb8 | 1.756231 | 0.003219 | 0.051794 |
| Hspb9 | -20.8342 | 2.31E-06 | 4.99E-05 |
| Htr1b | -21.0808 | 1.75E-06 | 4.27E-05 |
| Hus1b | -8.04404 | 0.000158 | 0.002817 |
| Hyal3 | -20.7631 | 2.50E-06 | 5.08E-05 |
| Icam4 | 24.1195 | 2.51E-08 | 5.01E-06 |
| Id4 | -22.4809 | 3.40E-07 | 3.08E-05 |
| Ifng | -21.3245 | 1.32E-06 | 3.81E-05 |
| Ift122 | -10.5761 | 3.48E-08 | 6.49E-06 |
| Ift140 | -12.5966 | 1.19E-11 | 2.01E-08 |
| Igf1r | -10.4894 | 0.002277 | 0.037249 |
| Igf2 | -7.44496 | 0.0038 | 0.060347 |

| | | | |
|---|---|---|---|
| Igll1 | -20.8219 | 2.34E-06 | 5.01E-05 |
| Igsf23 | -21.1909 | 1.54E-06 | 4.04E-05 |
| Ikzf3 | -21.1254 | 1.66E-06 | 4.23E-05 |
| Ikzf4 | 24.1195 | 2.51E-08 | 5.01E-06 |
| Ilk | -2.08665 | 0.001371 | 0.022831 |
| Inpp5b | -4.41349 | 0.002108 | 0.034556 |
| Insr | -11.8673 | 4.06E-09 | 1.35E-06 |
| Invs | -20.7332 | 2.59E-06 | 5.15E-05 |
| Iqub | -21.6932 | 8.64E-07 | 3.55E-05 |
| Irf5 | -22.0543 | 5.65E-07 | 3.21E-05 |
| Irgc1 | -21.4909 | 1.09E-06 | 3.65E-05 |
| Irs3 | -21.0054 | 1.90E-06 | 4.48E-05 |
| Irx2 | -21.1176 | 1.68E-06 | 4.23E-05 |
| Irx3 | -10.2072 | 5.24E-06 | 9.91E-05 |
| Isg20 | -20.9395 | 2.05E-06 | 4.64E-05 |
| Islr2 | -9.89583 | 5.02E-05 | 0.000908 |
| Isoc1 | -1.06993 | 0.001523 | 0.025249 |
| Itgb3bp | 23.18766 | 8.47E-08 | 1.20E-05 |
| Itih5 | -9.87426 | 2.41E-08 | 4.99E-06 |
| Jade2 | -21.7197 | 8.38E-07 | 3.55E-05 |
| Kcnb1 | 23.52547 | 5.48E-08 | 8.93E-06 |
| Kcnj4 | -21.2107 | 1.51E-06 | 4.02E-05 |
| Kcp | 23.14234 | 8.98E-08 | 1.21E-05 |
| Kdm4dl | -21.2544 | 1.43E-06 | 3.98E-05 |
| Kdm5d | -5.91942 | 0.000979 | 0.016532 |
| Klf1 | -20.9277 | 2.08E-06 | 4.67E-05 |
| Klf7 | -19.5919 | 8.88E-06 | 0.000166 |
| Klhdc1 | -10.4854 | 5.81E-06 | 0.00011 |
| Klhl13 | -22.4466 | 3.54E-07 | 3.11E-05 |
| Klhl14 | -21.9222 | 6.61E-07 | 3.26E-05 |
| Klhl41 | -20.8538 | 2.26E-06 | 4.90E-05 |
| Klhl5 | -9.36438 | 0.000959 | 0.016214 |
| Klk7 | -13.6971 | 9.41E-10 | 5.14E-07 |
| Klk9 | -22.1817 | 4.86E-07 | 3.21E-05 |
| Krt14 | -21.5416 | 1.03E-06 | 3.65E-05 |
| Krt19 | -11.1742 | 5.99E-07 | 3.21E-05 |
| Krt8 | -2.51151 | 0.003961 | 0.062795 |
| Krt87 | -22.5067 | 3.29E-07 | 3.03E-05 |
| Krtdap | -21.2874 | 1.38E-06 | 3.87E-05 |
| Lat2 | -21.4845 | 1.10E-06 | 3.65E-05 |
| Ldhal6b | -20.1092 | 5.12E-06 | 9.70E-05 |
| Lgals3bp | -20.9625 | 2.00E-06 | 4.60E-05 |
| Limch1 | -3.73843 | 0.005034 | 0.078046 |
| Lipo4 | -22.432 | 3.61E-07 | 3.15E-05 |
| Lmntd2 | -20.4263 | 3.60E-06 | 6.96E-05 |
| Lmx1a | -20.4221 | 3.62E-06 | 6.98E-05 |
| Lncpint | -20.7961 | 2.41E-06 | 5.03E-05 |
| Lpar6 | -21.4991 | 1.08E-06 | 3.65E-05 |

| | | | |
|---|---|---|---|
| Lpcat2 | -21.9994 | 6.03E-07 | 3.21E-05 |
| Lrig3 | -21.9144 | 6.67E-07 | 3.26E-05 |
| Lrp12 | -22.0694 | 5.55E-07 | 3.21E-05 |
| Lrp4 | -22.2308 | 4.59E-07 | 3.21E-05 |
| Lrrc8b | -11.4608 | 1.35E-11 | 2.08E-08 |
| Lst1 | 21.99896 | 3.73E-07 | 3.16E-05 |
| Lypd3 | -21.2228 | 1.49E-06 | 4.01E-05 |
| Lyrm7 | -11.6015 | 2.09E-08 | 4.48E-06 |
| Lyst | -11.7539 | 0.005508 | 0.084319 |
| Maats1 | -21.7457 | 8.12E-07 | 3.51E-05 |
| Maged2 | -6.08762 | 0.00096 | 0.016214 |
| Mageh1 | -22.0543 | 5.65E-07 | 3.21E-05 |
| Maml2 | 23.1802 | 8.56E-08 | 1.20E-05 |
| Manba | -21.8519 | 7.18E-07 | 3.39E-05 |
| Maoa | -11.9635 | 0.003949 | 0.062665 |
| Map3k6 | -11.136 | 0.006218 | 0.093834 |
| Map6d1 | -19.5919 | 8.88E-06 | 0.000166 |
| Mblac1 | 23.03796 | 1.03E-07 | 1.27E-05 |
| Mblac2 | 25.16068 | 6.13E-09 | 1.79E-06 |
| Mdfic | -11.6851 | 1.65E-06 | 4.23E-05 |
| Med24 | -6.3199 | 0.001138 | 0.019102 |
| Meioc | -21.6932 | 8.64E-07 | 3.55E-05 |
| Memo1 | -1.39549 | 8.51E-05 | 0.00153 |
| Mettl7b | -21.0734 | 1.76E-06 | 4.27E-05 |
| Mfng | -21.1532 | 1.61E-06 | 4.16E-05 |
| Mfsd4a | -21.9035 | 6.75E-07 | 3.26E-05 |
| Mgat1 | -5.58265 | 0.005358 | 0.082617 |
| Micall2 | -21.8116 | 7.52E-07 | 3.44E-05 |
| Milr1 | -22.2716 | 4.37E-07 | 3.21E-05 |
| Mindy2 | -21.0755 | 1.76E-06 | 4.27E-05 |
| Mir1199 | -20.6881 | 2.71E-06 | 5.37E-05 |
| Mir503 | -22.0907 | 5.42E-07 | 3.21E-05 |
| Mir6516 | -21.3753 | 1.25E-06 | 3.76E-05 |
| Mir7670 | 23.75514 | 4.06E-08 | 7.17E-06 |
| Mir8116 | -21.6798 | 8.74E-07 | 3.55E-05 |
| Mitf | -11.3134 | 1.35E-06 | 3.87E-05 |
| Mmgt2 | -11.9588 | 1.47E-13 | 8.98E-10 |
| Mmp14 | -21.3479 | 1.29E-06 | 3.78E-05 |
| Mmp25 | -22.0044 | 6.00E-07 | 3.21E-05 |
| Mmp9 | -20.5873 | 3.04E-06 | 5.94E-05 |
| Mpp4 | 24.41071 | 1.70E-08 | 3.94E-06 |
| Mrpl22 | 1.228835 | 0.001707 | 0.028222 |
| Mrps27 | -12.0562 | 0.003492 | 0.055675 |
| Mslnl | -21.3622 | 1.26E-06 | 3.78E-05 |
| Msn | -2.52154 | 0.000377 | 0.006533 |
| Mtcp1 | -21.2442 | 1.45E-06 | 4.00E-05 |
| Mterf1b | -21.4983 | 1.08E-06 | 3.65E-05 |
| Mterf2 | -10.466 | 8.22E-11 | 8.20E-08 |

| | | | |
|---|---|---|---|
| Muc13 | 10.09794 | 7.84E-05 | 0.001413 |
| Mycn | -13.3453 | 2.54E-09 | 1.03E-06 |
| Myh10 | -1.77069 | 4.04E-07 | 3.21E-05 |
| Myh9 | -1.38765 | 0.003972 | 0.062916 |
| Myl9 | -11.8333 | 1.91E-07 | 2.06E-05 |
| Myof | -12.575 | 0.003253 | 0.052262 |
| Mzf1 | -11.0864 | 3.86E-05 | 0.000701 |
| Naaladl1 | -10.0459 | 3.79E-07 | 3.18E-05 |
| Nav3 | -21.2228 | 1.49E-06 | 4.01E-05 |
| Nceh1 | -12.0626 | 1.97E-08 | 4.28E-06 |
| Nckap1l | -21.8248 | 7.41E-07 | 3.44E-05 |
| Nckap5l | -21.2442 | 1.45E-06 | 4.00E-05 |
| Ncoa7 | -11.3662 | 5.55E-10 | 3.33E-07 |
| Ndrg4 | 22.58484 | 1.81E-07 | 1.99E-05 |
| Necap2 | -6.33052 | 0.001895 | 0.031215 |
| Nedd9 | -21.0734 | 1.76E-06 | 4.27E-05 |
| Neurl2 | -20.9625 | 2.00E-06 | 4.60E-05 |
| Nfam1 | -21.9335 | 6.52E-07 | 3.26E-05 |
| Nfatc2 | -11.7912 | 1.50E-05 | 0.000278 |
| Nfil3 | -21.519 | 1.06E-06 | 3.65E-05 |
| Nfix | -21.4667 | 1.12E-06 | 3.65E-05 |
| Nfkbiz | -21.3759 | 1.25E-06 | 3.76E-05 |
| Ngb | -21.2852 | 1.38E-06 | 3.87E-05 |
| Nkd2 | -20.9277 | 2.08E-06 | 4.67E-05 |
| Nkx2-3 | -21.8081 | 7.55E-07 | 3.44E-05 |
| Nlrx1 | -12.927 | 2.41E-11 | 2.92E-08 |
| Nmnat3 | -20.5965 | 2.98E-06 | 5.85E-05 |
| Nnat | -20.7624 | 2.50E-06 | 5.08E-05 |
| Nod1 | -20.7624 | 2.50E-06 | 5.08E-05 |
| Nodal | -21.3416 | 1.30E-06 | 3.78E-05 |
| Nova2 | -22.3078 | 4.18E-07 | 3.21E-05 |
| Npr2 | -12.6087 | 3.96E-08 | 7.07E-06 |
| Nqo1 | -21.4748 | 1.11E-06 | 3.65E-05 |
| Nqo2 | -1.36966 | 0.002597 | 0.042233 |
| Nr1d1 | 11.6234 | 1.96E-07 | 2.06E-05 |
| Nt5dc2 | 10.88502 | 2.17E-05 | 0.000399 |
| Ntf3 | -21.3845 | 1.23E-06 | 3.76E-05 |
| Nub1 | 1.771568 | 0.000507 | 0.008741 |
| Nudt12 | -1.57322 | 0.000268 | 0.004685 |
| Nudt8 | -22.1845 | 4.85E-07 | 3.21E-05 |
| Nxf2 | -21.6416 | 9.17E-07 | 3.57E-05 |
| Oaf | -21.1041 | 1.70E-06 | 4.23E-05 |
| Oas1c | 23.35826 | 6.81E-08 | 1.03E-05 |
| Oas1f | -21.3759 | 1.25E-06 | 3.76E-05 |
| Oas1g | 23.39858 | 6.46E-08 | 9.95E-06 |
| Oat | -2.97322 | 0.0002 | 0.003533 |
| Oaz3 | 10.14246 | 0.004532 | 0.071122 |
| Obox3-ps2 | -20.9742 | 1.97E-06 | 4.57E-05 |

| | | | |
|---|---|---|---|
| Obox3-ps6 | -21.9724 | 6.23E-07 | 3.26E-05 |
| Obox4-ps15 | -21.8612 | 7.10E-07 | 3.36E-05 |
| Obox4-ps16 | -21.4909 | 1.09E-06 | 3.65E-05 |
| Obox4-ps17 | -20.6196 | 2.92E-06 | 5.76E-05 |
| Obox4-ps18 | -21.2212 | 1.49E-06 | 4.01E-05 |
| Obox4-ps21 | -21.7197 | 8.38E-07 | 3.55E-05 |
| Obox4-ps33 | -21.6797 | 8.77E-07 | 3.55E-05 |
| Olfr106-ps | -22.2928 | 4.26E-07 | 3.21E-05 |
| Olfr787 | -21.2228 | 1.49E-06 | 4.01E-05 |
| Olig1 | -21.4558 | 1.14E-06 | 3.66E-05 |
| Oscp1 | -22.1382 | 5.12E-07 | 3.21E-05 |
| Otub2 | -21.2874 | 1.38E-06 | 3.87E-05 |
| Ovol1 | 23.21331 | 8.20E-08 | 1.17E-05 |
| P2ry1 | -20.7444 | 2.55E-06 | 5.12E-05 |
| P2ry14 | -22.9768 | 1.86E-07 | 2.04E-05 |
| P3h3 | -10.9137 | 3.85E-06 | 7.42E-05 |
| Paqr3 | -25.5084 | 5.71E-09 | 1.70E-06 |
| Paqr4 | -12.2746 | 1.66E-10 | 1.48E-07 |
| Paqr7 | -21.8883 | 6.87E-07 | 3.27E-05 |
| Parp14 | 23.03135 | 1.03E-07 | 1.27E-05 |
| Paxx | -21.0808 | 1.75E-06 | 4.27E-05 |
| Pbdc1 | 1.204354 | 0.000319 | 0.005549 |
| Pbld1 | 9.182175 | 8.10E-05 | 0.001459 |
| Pbx1 | -22.0532 | 5.66E-07 | 3.21E-05 |
| Pcbp3 | -21.7521 | 8.06E-07 | 3.51E-05 |
| Pcdh19 | 24.1404 | 2.44E-08 | 4.99E-06 |
| Pcdhgc5 | -22.1793 | 4.88E-07 | 3.21E-05 |
| Pde1b | -11.7842 | 0.004733 | 0.073922 |
| Pear1 | -23.1008 | 1.60E-07 | 1.81E-05 |
| Peg10 | -2.55922 | 0.006364 | 0.095607 |
| Peg13 | -22.2398 | 4.54E-07 | 3.21E-05 |
| Per3 | -21.2544 | 1.43E-06 | 3.98E-05 |
| Perm1 | -20.9395 | 2.05E-06 | 4.64E-05 |
| Pex11b | -11.1576 | 5.05E-10 | 3.29E-07 |
| Phf11a | -22.1824 | 4.85E-07 | 3.21E-05 |
| Phyhipl | -20.7863 | 2.44E-06 | 5.06E-05 |
| Pisd-ps2 | -22.0667 | 5.57E-07 | 3.21E-05 |
| Pitpnm1 | -21.6416 | 9.17E-07 | 3.57E-05 |
| Pkhd1 | -20.32 | 4.06E-06 | 7.73E-05 |
| Pla2g2f | -21.1259 | 1.66E-06 | 4.23E-05 |
| Plac1 | -13.0698 | 2.75E-08 | 5.43E-06 |
| Plat | -13.375 | 7.04E-12 | 1.49E-08 |
| Platr23 | -21.9335 | 6.52E-07 | 3.26E-05 |
| Platr28 | -21.7775 | 7.83E-07 | 3.47E-05 |
| Platr31 | -19.5919 | 8.88E-06 | 0.000166 |
| Plcxd2 | -21.628 | 9.32E-07 | 3.58E-05 |
| Plekhh2 | -22.0098 | 5.96E-07 | 3.21E-05 |
| Plekhh3 | -12.4295 | 3.26E-10 | 2.40E-07 |

| | | | |
|---|---|---|---|
| Plin4 | -21.6387 | 9.20E-07 | 3.57E-05 |
| Plscr1 | 23.12936 | 9.13E-08 | 1.21E-05 |
| Plvap | -21.3331 | 1.31E-06 | 3.80E-05 |
| Pmm1 | 2.053853 | 3.77E-05 | 0.000687 |
| Pogk | -26.3016 | 2.15E-09 | 9.12E-07 |
| Poli | -11.3333 | 0.00663 | 0.098815 |
| Poln | -20.5873 | 3.04E-06 | 5.94E-05 |
| Polr2j | 1.151438 | 0.005147 | 0.079661 |
| Ppfia3 | -21.3331 | 1.31E-06 | 3.80E-05 |
| Ppp1r14d | -12.1601 | 0.004227 | 0.0667 |
| Ppp1r2-ps4 | -3.75599 | 0.005103 | 0.079044 |
| Ppp1r3c | -21.4265 | 1.18E-06 | 3.69E-05 |
| Ppp2r2b | -21.5879 | 9.76E-07 | 3.60E-05 |
| Prag1 | -23.1051 | 1.59E-07 | 1.81E-05 |
| Prkch | -22.2796 | 4.32E-07 | 3.21E-05 |
| Prmt2 | -10.3286 | 5.31E-07 | 3.21E-05 |
| Prn | -20.8122 | 2.36E-06 | 5.03E-05 |
| Prox2 | -20.9742 | 1.97E-06 | 4.57E-05 |
| Prr15 | -20.9379 | 2.06E-06 | 4.64E-05 |
| Prr23a2 | 23.12773 | 9.15E-08 | 1.21E-05 |
| Prss32 | -12.4855 | 8.06E-09 | 2.10E-06 |
| Prss45 | 25.00829 | 7.56E-09 | 2.00E-06 |
| Prtn3 | 11.54628 | 1.44E-06 | 3.99E-05 |
| Psd3 | -20.7624 | 2.50E-06 | 5.08E-05 |
| Ptbp3 | -1.93002 | 0.005446 | 0.08367 |
| Ptch2 | -21.393 | 1.22E-06 | 3.76E-05 |
| Ptges3l | -22.4037 | 3.73E-07 | 3.16E-05 |
| Pth1r | -11.4489 | 1.17E-06 | 3.69E-05 |
| Ptpn9 | -11.4277 | 0.005647 | 0.08622 |
| Pus10 | -1.42457 | 0.00454 | 0.071172 |
| Pus7l | -21.4427 | 1.15E-06 | 3.67E-05 |
| Pyy | -20.8017 | 2.39E-06 | 5.03E-05 |
| Rab11fip5 | -1.90785 | 0.002697 | 0.043745 |
| Rab33b | -1.4707 | 0.006393 | 0.095968 |
| Rab43 | -21.5715 | 9.95E-07 | 3.60E-05 |
| Rad51b | -20.7444 | 2.55E-06 | 5.12E-05 |
| Ramp2 | 26.80448 | 5.90E-10 | 3.33E-07 |
| Rangrf | 1.978322 | 9.07E-05 | 0.001631 |
| Rasgrp1 | -20.4557 | 3.51E-06 | 6.80E-05 |
| Rassf10 | -21.1041 | 1.70E-06 | 4.23E-05 |
| Raver2 | -21.2049 | 1.52E-06 | 4.02E-05 |
| Rbks | -22.1137 | 5.27E-07 | 3.21E-05 |
| Rbm25 | -11.3108 | 1.85E-05 | 0.000341 |
| Rftn1 | -20.7961 | 2.41E-06 | 5.03E-05 |
| Rfx3 | -21.7775 | 7.83E-07 | 3.47E-05 |
| Rhbdf1 | -11.4226 | 0.005663 | 0.08638 |
| Rhbg | -21.5715 | 9.95E-07 | 3.60E-05 |
| Rhd | -21.4587 | 1.13E-06 | 3.65E-05 |

| | | | |
|---|---|---|---|
| Rhobtb3 | -20.7624 | 2.50E-06 | 5.08E-05 |
| Rhoc | -12.8509 | 9.19E-11 | 8.65E-08 |
| Rhox1 | -21.5953 | 9.68E-07 | 3.60E-05 |
| Rhox9 | 12.21488 | 3.21E-07 | 2.97E-05 |
| Rilp | -21.5803 | 9.81E-07 | 3.60E-05 |
| Ripk3 | -21.9559 | 6.35E-07 | 3.26E-05 |
| Ripor3 | -21.3759 | 1.25E-06 | 3.76E-05 |
| Rlbp1 | -21.9994 | 6.03E-07 | 3.21E-05 |
| Rnd2 | 2.097964 | 0.003124 | 0.050374 |
| Rnf224 | -21.2319 | 1.47E-06 | 4.01E-05 |
| Rpgrip1 | 1.401502 | 0.00054 | 0.009287 |
| Rpl13-ps2 | -22.5986 | 2.95E-07 | 2.81E-05 |
| Rpl36a | 1.158332 | 0.003389 | 0.054181 |
| Rpp38 | 10.49318 | 3.84E-05 | 0.000698 |
| Rps6ka2 | -20.8989 | 2.15E-06 | 4.75E-05 |
| Rtn1 | -11.6036 | 0.00524 | 0.080955 |
| Rufy1 | -11.5148 | 0.004983 | 0.077339 |
| Rufy4 | -21.9335 | 6.52E-07 | 3.26E-05 |
| Runx1 | -13.1757 | 2.71E-13 | 9.20E-10 |
| Runx2 | -21.9144 | 6.67E-07 | 3.26E-05 |
| Rwdd2a | -21.2874 | 1.38E-06 | 3.87E-05 |
| Ryr2 | -22.0694 | 5.55E-07 | 3.21E-05 |
| S100a3 | 9.38606 | 4.21E-08 | 7.35E-06 |
| Sacs | -20.7624 | 2.50E-06 | 5.08E-05 |
| Sall2 | 22.25989 | 2.71E-07 | 2.69E-05 |
| Sash3 | -20.3584 | 3.89E-06 | 7.47E-05 |
| Sat2 | -21.8182 | 7.46E-07 | 3.44E-05 |
| Sdc3 | -21.9222 | 6.61E-07 | 3.26E-05 |
| Sdcbp2 | -22.2142 | 4.68E-07 | 3.21E-05 |
| Sec31b | -21.6244 | 9.36E-07 | 3.58E-05 |
| Selenom | -21.4909 | 1.09E-06 | 3.65E-05 |
| Sema3f | -21.9504 | 6.39E-07 | 3.26E-05 |
| Sema4c | 12.27954 | 2.21E-06 | 4.84E-05 |
| Serinc4 | -9.53193 | 0.000996 | 0.016802 |
| Serpinb6b | -22.4517 | 3.52E-07 | 3.11E-05 |
| Serpinb9 | -21.3416 | 1.30E-06 | 3.78E-05 |
| Serping1 | -21.0398 | 1.83E-06 | 4.36E-05 |
| Sertad3 | -21.0717 | 1.76E-06 | 4.27E-05 |
| Sestd1 | -22.1708 | 4.92E-07 | 3.21E-05 |
| Sh2b1 | -1.51137 | 0.000814 | 0.013821 |
| Sh3bp5l | -11.6465 | 0.004464 | 0.070176 |
| Shisa7 | 23.06225 | 9.95E-08 | 1.27E-05 |
| Shisa8 | -21.8124 | 7.52E-07 | 3.44E-05 |
| Slc15a2 | -7.26479 | 0.003271 | 0.052501 |
| Slc16a3 | -20.8989 | 2.15E-06 | 4.75E-05 |
| Slc16a4 | -20.4845 | 3.38E-06 | 6.57E-05 |
| Slc25a18 | -21.011 | 1.89E-06 | 4.46E-05 |
| Slc25a45 | -21.6387 | 9.20E-07 | 3.57E-05 |

| | | | |
|---|---|---|---|
| Slc26a3 | -21.7775 | 7.83E-07 | 3.47E-05 |
| Slc2a4 | -21.8124 | 7.52E-07 | 3.44E-05 |
| Slc35f3 | -21.2049 | 1.52E-06 | 4.02E-05 |
| Slc36a3os | -22.4986 | 3.33E-07 | 3.03E-05 |
| Slc38a4 | -26.602 | 1.58E-09 | 6.88E-07 |
| Slc39a2 | -11.9265 | 0.004347 | 0.068409 |
| Slc48a1 | -21.0734 | 1.76E-06 | 4.27E-05 |
| Slc51b | -22.2395 | 4.54E-07 | 3.21E-05 |
| Slc5a10 | -20.8795 | 2.20E-06 | 4.81E-05 |
| Slc5a4b | -21.5715 | 9.95E-07 | 3.60E-05 |
| Slc6a12 | -21.4931 | 1.09E-06 | 3.65E-05 |
| Slc6a7 | -21.5801 | 9.85E-07 | 3.60E-05 |
| Slc7a14 | -20.9625 | 2.00E-06 | 4.60E-05 |
| Snhg7os | 23.47219 | 5.87E-08 | 9.34E-06 |
| Snora16a | 11.17111 | 0.001358 | 0.022623 |
| Snord15b | -21.1545 | 1.61E-06 | 4.16E-05 |
| Snord17 | -21.0927 | 1.72E-06 | 4.27E-05 |
| Snta1 | -21.3245 | 1.32E-06 | 3.81E-05 |
| Snx4 | -3.62062 | 0.003165 | 0.050978 |
| Socs7 | -11.7974 | 0.004297 | 0.067738 |
| Sod2 | 4.791319 | 0.006146 | 0.09292 |
| Sox9 | 24.01442 | 2.89E-08 | 5.62E-06 |
| Spata31d1a | 22.99944 | 1.08E-07 | 1.29E-05 |
| Spdya | 23.01803 | 1.05E-07 | 1.27E-05 |
| Speer4cos | -21.4286 | 1.17E-06 | 3.69E-05 |
| Sphk1 | -21.1909 | 1.54E-06 | 4.04E-05 |
| Srp9 | 1.132573 | 0.001162 | 0.019483 |
| Ssmem1 | -20.7332 | 2.59E-06 | 5.15E-05 |
| St3gal2 | -11.0979 | 2.43E-08 | 4.99E-06 |
| St8sia1 | -21.4507 | 1.14E-06 | 3.66E-05 |
| Stam | -2.47344 | 1.01E-12 | 2.86E-09 |
| Stard5 | -6.39843 | 0.000358 | 0.00622 |
| Stard7 | -1.93255 | 1.12E-05 | 0.000209 |
| Stat1 | -11.2901 | 0.005907 | 0.089779 |
| Stau1 | -1.34462 | 0.001125 | 0.018896 |
| Steap3 | -1.84316 | 0.003467 | 0.055323 |
| Sting1 | -20.9742 | 1.97E-06 | 4.57E-05 |
| Stk32c | -21.603 | 9.59E-07 | 3.60E-05 |
| Stra8 | -21.2049 | 1.52E-06 | 4.02E-05 |
| Stx19 | -21.5511 | 1.02E-06 | 3.65E-05 |
| Stx2 | -22.1626 | 4.97E-07 | 3.21E-05 |
| Sun2 | -11.7627 | 3.54E-11 | 3.75E-08 |
| Susd1 | 23.10076 | 9.47E-08 | 1.23E-05 |
| Susd6 | -1.55515 | 0.006531 | 0.09746 |
| Syce1l | -21.6709 | 8.87E-07 | 3.56E-05 |
| Sycn | 2.005851 | 0.000143 | 0.002564 |
| Synpr | -20.7332 | 2.59E-06 | 5.15E-05 |
| Syt1 | -20.32 | 4.06E-06 | 7.73E-05 |

| | | | |
|---|---|---|---|
| Syt11 | -21.4667 | 1.12E-06 | 3.65E-05 |
| Tacr3 | -21.6865 | 8.71E-07 | 3.55E-05 |
| Tacstd2 | -3.16565 | 0.002646 | 0.042945 |
| Taf7l2 | -20.4263 | 3.60E-06 | 6.96E-05 |
| Tagln | -21.1041 | 1.70E-06 | 4.23E-05 |
| Tapbpl | 11.33581 | 7.78E-06 | 0.000146 |
| Tbc1d22bos | -21.1532 | 1.61E-06 | 4.16E-05 |
| Tbc1d32 | -21.4909 | 1.09E-06 | 3.65E-05 |
| Tbx1 | -21.4099 | 1.20E-06 | 3.74E-05 |
| Tbx18 | -21.5953 | 9.68E-07 | 3.60E-05 |
| Tc2n | -21.3637 | 1.26E-06 | 3.78E-05 |
| Tceal1 | -21.9614 | 6.31E-07 | 3.26E-05 |
| Tcf24 | -20.8795 | 2.20E-06 | 4.81E-05 |
| Tcp11l2 | -22.0658 | 5.58E-07 | 3.21E-05 |
| Tdpoz3 | -22.1434 | 5.09E-07 | 3.21E-05 |
| Tdpoz9 | -22.1537 | 5.03E-07 | 3.21E-05 |
| Terb2 | 11.23407 | 1.69E-07 | 1.90E-05 |
| Tfpi | -10.8213 | 1.17E-11 | 2.01E-08 |
| Thap1 | 1.822481 | 0.000418 | 0.007215 |
| Thbs3 | -21.9957 | 6.06E-07 | 3.22E-05 |
| Them6 | -11.3587 | 1.45E-07 | 1.67E-05 |
| Tifab | -11.8714 | 5.73E-10 | 3.33E-07 |
| Timeless | -1.40429 | 0.002959 | 0.047849 |
| Timp3 | -21.8385 | 7.29E-07 | 3.41E-05 |
| Tle6 | -11.2898 | 0.006131 | 0.092764 |
| Tln1 | -1.44488 | 0.000271 | 0.004724 |
| Tm4sf5 | -20.8219 | 2.34E-06 | 5.01E-05 |
| Tmco2 | -21.1041 | 1.70E-06 | 4.23E-05 |
| Tmem241 | -20.7863 | 2.44E-06 | 5.06E-05 |
| Tmem260 | -5.95661 | 0.000449 | 0.007755 |
| Tmem54 | -22.1284 | 5.18E-07 | 3.21E-05 |
| Tmem59l | -21.0516 | 1.81E-06 | 4.34E-05 |
| Tmprss9 | -20.835 | 2.31E-06 | 4.99E-05 |
| Tmtc4 | -11.6255 | 1.06E-05 | 0.000197 |
| Tns2 | -21.9559 | 6.35E-07 | 3.26E-05 |
| Tpcn1 | -7.53907 | 1.68E-05 | 0.00031 |
| Tpi-rs4 | -20.8122 | 2.36E-06 | 5.03E-05 |
| Tpi1 | -3.25475 | 0.000287 | 0.004999 |
| Trim16 | -22.2716 | 4.37E-07 | 3.21E-05 |
| Trim63 | 23.46917 | 5.90E-08 | 9.34E-06 |
| Trp53cor1 | -20.8989 | 2.15E-06 | 4.75E-05 |
| Trp53tg5 | -21.5116 | 1.07E-06 | 3.65E-05 |
| Trpm3 | -21.4182 | 1.19E-06 | 3.71E-05 |
| Trpv2 | -21.1143 | 1.68E-06 | 4.23E-05 |
| Tsc22d3 | -21.2412 | 1.46E-06 | 4.00E-05 |
| Tsga10 | 23.52547 | 5.48E-08 | 8.93E-06 |
| Tspan8 | -3.99559 | 0.000197 | 0.003481 |
| Tspo | -7.96769 | 0.005238 | 0.080955 |

| | | | |
|---|---|---|---|
| Tssk1 | -20.7332 | 2.59E-06 | 5.15E-05 |
| Ttc12 | -21.6797 | 8.77E-07 | 3.55E-05 |
| Ttc16 | -20.6881 | 2.71E-06 | 5.37E-05 |
| Ttc23 | -20.1092 | 5.12E-06 | 9.70E-05 |
| Ttc41 | -22.1845 | 4.85E-07 | 3.21E-05 |
| Ttc8 | -20.984 | 1.95E-06 | 4.57E-05 |
| Ttn | -21.5416 | 1.03E-06 | 3.65E-05 |
| Tuba3b | -21.6244 | 9.36E-07 | 3.58E-05 |
| Tubb3 | 12.26721 | 2.05E-05 | 0.000378 |
| Tvp23b | -12.1126 | 0.003541 | 0.056396 |
| Twf2 | -1.70557 | 0.003344 | 0.05357 |
| Twnk | 1.931214 | 0.001945 | 0.031963 |
| Txlnb | 9.36386 | 0.00017 | 0.003014 |
| Txnip | -6.2264 | 1.55E-05 | 0.000286 |
| Ubap1l | -20.9625 | 2.00E-06 | 4.60E-05 |
| Ucn | 24.40371 | 1.72E-08 | 3.94E-06 |
| Uevld | -1.37315 | 0.000804 | 0.013688 |
| Ulk3 | -10.6546 | 6.39E-09 | 1.83E-06 |
| Unc5a | -20.9395 | 2.05E-06 | 4.64E-05 |
| Unc93a | -11.329 | 0.006714 | 0.099979 |
| Unc93a2 | -11.3939 | 4.63E-07 | 3.21E-05 |
| Usp17la | -22.1845 | 4.85E-07 | 3.21E-05 |
| Usp24 | -1.0341 | 0.000769 | 0.013118 |
| Usp26 | -21.5116 | 1.07E-06 | 3.65E-05 |
| Vasn | -21.8883 | 6.87E-07 | 3.27E-05 |
| Vasp | -1.69229 | 0.001399 | 0.023264 |
| Vegfa | -10.4049 | 0.000258 | 0.004513 |
| Vegfc | -20.6196 | 2.92E-06 | 5.76E-05 |
| Vgll1 | -11.2115 | 0.005989 | 0.090945 |
| Vim | -11.3175 | 2.86E-09 | 1.08E-06 |
| Vldlr | -22.3298 | 4.08E-07 | 3.21E-05 |
| Vmn1r-ps32 | -22.9046 | 2.03E-07 | 2.13E-05 |
| Vmn1r139 | 25.21159 | 5.71E-09 | 1.70E-06 |
| Vmn2r-ps106 | 7.429013 | 0.001015 | 0.017092 |
| Vmn2r29 | -20.9564 | 2.01E-06 | 4.62E-05 |
| Vpreb1 | -21.3416 | 1.30E-06 | 3.78E-05 |
| Vps37d | 23.88648 | 3.42E-08 | 6.44E-06 |
| Vps52 | -1.30255 | 0.000654 | 0.011214 |
| Vps9d1 | -10.6913 | 1.56E-07 | 1.79E-05 |
| Was | 23.03796 | 1.03E-07 | 1.27E-05 |
| Wasf3 | -20.8538 | 2.26E-06 | 4.90E-05 |
| Wbp2nl | -21.5953 | 9.68E-07 | 3.60E-05 |
| Wdr19 | -22.9458 | 1.93E-07 | 2.06E-05 |
| Wdr27 | 11.33328 | 5.10E-05 | 0.000921 |
| Wdr5b | -11.3636 | 3.20E-05 | 0.000585 |
| Wdr72 | -11.1185 | 3.84E-10 | 2.71E-07 |
| Wdr95 | -21.5879 | 9.76E-07 | 3.60E-05 |
| Wls | -11.7224 | 6.46E-07 | 3.26E-05 |

| | | | |
|---|---|---|---|
| Wnk4 | -21.7649 | 7.94E-07 | 3.49E-05 |
| Wnt2b | -21.8766 | 6.97E-07 | 3.31E-05 |
| Wnt4 | -20.9742 | 1.97E-06 | 4.57E-05 |
| Xdh | -6.04714 | 0.004514 | 0.070903 |
| Xist | 4.719308 | 0.000171 | 0.003026 |
| Xlr4e-ps | -7.48782 | 0.006074 | 0.092077 |
| Zeb1 | -21.494 | 1.09E-06 | 3.65E-05 |
| Zfp114 | -21.9041 | 6.75E-07 | 3.26E-05 |
| Zfp119b | -22.63 | 2.84E-07 | 2.75E-05 |
| Zfp13 | -20.6196 | 2.92E-06 | 5.76E-05 |
| Zfp148 | -1.0119 | 0.00093 | 0.01576 |
| Zfp202 | -21.6932 | 8.64E-07 | 3.55E-05 |
| Zfp354b | -21.2049 | 1.52E-06 | 4.02E-05 |
| Zfp36l3 | -20.7631 | 2.50E-06 | 5.08E-05 |
| Zfp422 | -20.9277 | 2.08E-06 | 4.67E-05 |
| Zfp46 | -11.2373 | 0.006296 | 0.094846 |
| Zfp563 | -21.4909 | 1.09E-06 | 3.65E-05 |
| Zfp597 | -22.2353 | 4.56E-07 | 3.21E-05 |
| Zfp605 | -21.931 | 6.54E-07 | 3.26E-05 |
| Zfp647 | -20.7624 | 2.50E-06 | 5.08E-05 |
| Zfp783 | -21.8883 | 6.87E-07 | 3.27E-05 |
| Zfp786 | -20.8219 | 2.34E-06 | 5.01E-05 |
| Zfp810 | -9.85537 | 0.000536 | 0.009219 |
| Zfp93 | -21.5879 | 9.76E-07 | 3.60E-05 |
| Zfp937 | -21.4909 | 1.09E-06 | 3.65E-05 |
| Zfp938 | -11.2873 | 3.36E-08 | 6.39E-06 |
| Zfp964 | -20.8017 | 2.39E-06 | 5.03E-05 |
| Zfp974 | 25.48208 | 3.92E-09 | 1.33E-06 |
| Zfy1 | -22.0085 | 5.97E-07 | 3.21E-05 |
| Zic2 | -20.8795 | 2.20E-06 | 4.81E-05 |
| Zic5 | -21.5822 | 9.83E-07 | 3.60E-05 |
| Zim3 | 22.81259 | 1.36E-07 | 1.58E-05 |
| Zscan18 | -21.7197 | 8.38E-07 | 3.55E-05 |
