## Supplemental Table 2 for "Environmentally Induced Sperm RNAs Transmit Cancer Susceptibility to Offspring in a Mouse Model"

**Table S2- Predicted targets of miR-10b which are down-regulted in miR-10b injected E3.5 embryos**

| Data base | Gene | log2FC | pvalue | padj | miRNA Name |
|---|---|---|---|---|---|
| miRBase | Abcg1 | -10.445775 | 4.74E-05 | 8.58E-04 | mmu-miR-10b-5p |
| miRBase | Akap5 | -21.913765 | 6.67E-07 | 3.26E-05 | mmu-miR-10b-5p |
| miRBase/Target Scan | Dlgap2 | -21.254448 | 1.43E-06 | 3.98E-05 | mmu-miR-10b-5p |
| miRBase/Target Scan | Elavl3 | -22.033425 | 5.79E-07 | 3.21E-05 | mmu-miR-10b-5p |
| Target Scan | Fads3 | -22.041393 | 5.74E-07 | 3.21E-05 | mmu-miR-10b-5p |
| miRBase/Target Scan | Fnbp1l | -1.382392 | 2.25E-05 | 4.13E-04 | mmu-miR-10b-5p |
| miRBase/Target Scan | Grin3a | -11.559716 | 4.85E-07 | 3.21E-05 | mmu-miR-10b-5p |
| miRBase | Id4 | -22.480901 | 3.40E-07 | 3.08E-05 | mmu-miR-10b-5p |
| miRBase | Serpinb6b | -22.4517 | 3.52E-07 | 3.11E-05 | mmu-miR-10b-5p |
| miRBase/Target Scan | Snx4 | -3.6206167 | 0.00316453 | 0.05097835 | mmu-miR-10b-5p |
| miRBase | Susd6 | -1.5551483 | 0.00653052 | 0.0974599 | mmu-miR-10b-5p |
| miRBase | Zfp563 | -21.490918 | 1.09E-06 | 3.65E-05 | mmu-miR-10b-5p |
| Target Scan | Ccdc71l | -10.157681 | 2.04E-05 | 3.75E-04 | mmu-miR-10b-5p |
| Target Scan | Nfix | -21.466706 | 1.12E-06 | 3.65E-05 | mmu-miR-10b-5p |
| Target Scan | Abcc2 | -21.104134 | 1.70E-06 | 4.23E-05 | mmu-miR-10b-3p |
| Target Scan | Akap5 | -21.913765 | 6.67E-07 | 3.26E-05 | mmu-miR-10b-3p |
| Target Scan | Ano1 | -22.008544 | 5.97E-07 | 3.21E-05 | mmu-miR-10b-3p |
| Target Scan | Atp2b1 | -1.3131448 | 0.00635891 | 0.09560708 | mmu-miR-10b-3p |
| Target Scan | Coq10b | -2.154341 | 1.96E-07 | 2.06E-05 | mmu-miR-10b-3p |
| Target Scan | Cpeb2 | -1.3988442 | 0.00322124 | 0.05179355 | mmu-miR-10b-3p |
| Target Scan | Efhb | -21.68645 | 8.71E-07 | 3.55E-05 | mmu-miR-10b-3p |
| Target Scan | Eid1 | -10.260812 | 1.19E-07 | 1.39E-05 | mmu-miR-10b-3p |
| Target Scan | Eif5a2 | -21.68645 | 8.71E-07 | 3.55E-05 | mmu-miR-10b-3p |
| Target Scan | Fam126b | -11.413839 | 1.04E-07 | 1.27E-05 | mmu-miR-10b-3p |
| Target Scan | Fgf7 | -22.214217 | 4.68E-07 | 3.21E-05 | mmu-miR-10b-3p |
| Target Scan | Gda | -21.603008 | 9.59E-07 | 3.60E-05 | mmu-miR-10b-3p |
| Target Scan | Gramd1a | -21.458666 | 1.13E-06 | 3.65E-05 | mmu-miR-10b-3p |
| Target Scan | Grik3 | -9.8825209 | 4.32E-06 | 8.21E-05 | mmu-miR-10b-3p |
| Target Scan | Hoxa13 | -20.898893 | 2.15E-06 | 4.75E-05 | mmu-miR-10b-3p |
| Target Scan | Hspa13 | -9.9135648 | 2.35E-07 | 2.39E-05 | mmu-miR-10b-3p |
| Target Scan | Ifng | -21.324502 | 1.32E-06 | 3.81E-05 | mmu-miR-10b-3p |
| Target Scan | Igsf23 | -21.190879 | 1.54E-06 | 4.04E-05 | mmu-miR-10b-3p |
| Target Scan | Itih5 | -9.8742581 | 2.41E-08 | 4.99E-06 | mmu-miR-10b-3p |
| Target Scan | Kcnj4 | -21.21073 | 1.51E-06 | 4.02E-05 | mmu-miR-10b-3p |
| Target Scan | Klf7 | -19.591937 | 8.88E-06 | 1.66E-04 | mmu-miR-10b-3p |
| Target Scan | Lrrc8b | -11.460779 | 1.35E-11 | 2.08E-08 | mmu-miR-10b-3p |
| Target Scan | Mfng | -21.153212 | 1.61E-06 | 4.16E-05 | mmu-miR-10b-3p |
| Target Scan | Mgat1 | -5.5826543 | 0.00535766 | 0.08261713 | mmu-miR-10b-3p |
| Target Scan | Mmp25 | -22.004396 | 6.00E-07 | 3.21E-05 | mmu-miR-10b-3p |
| Target Scan | Mtcp1 | -21.244157 | 1.45E-06 | 4.00E-05 | mmu-miR-10b-3p |
| miRBase/Target Scan | Myh9 | -1.3876537 | 0.00397242 | 0.06291645 | mmu-miR-10b-3p |

| | | | | | |
|---|---|---|---|---|---|
| Target Scan | Nav3 | -21.222819 | 1.49E-06 | 4.01E-05 | mmu-miR-10b-3p |
| Target Scan | Nfam1 | -21.933472 | 6.52E-07 | 3.26E-05 | mmu-miR-10b-3p |
| Target Scan | Nfix | -21.466706 | 1.12E-06 | 3.65E-05 | mmu-miR-10b-3p |
| Target Scan | Nod1 | -20.762434 | 2.50E-06 | 5.08E-05 | mmu-miR-10b-3p |
| Target Scan | Nqo1 | -21.474793 | 1.11E-06 | 3.65E-05 | mmu-miR-10b-3p |
| Target Scan | Oat | -2.9732222 | 2.00E-04 | 0.00353344 | mmu-miR-10b-3p |
| Target Scan | Plcxd2 | -21.627974 | 9.32E-07 | 3.58E-05 | mmu-miR-10b-3p |
| Target Scan | Psd3 | -20.762434 | 2.50E-06 | 5.08E-05 | mmu-miR-10b-3p |
| Target Scan | Ptbp3 | -1.9300208 | 0.00544566 | 0.08366963 | mmu-miR-10b-3p |
| Target Scan | Raver2 | -21.204941 | 1.52E-06 | 4.02E-05 | mmu-miR-10b-3p |
| Target Scan | Slc38a4 | -26.602025 | 1.58E-09 | 6.88E-07 | mmu-miR-10b-3p |
| Target Scan | St3gal2 | -11.097895 | 2.43E-08 | 4.99E-06 | mmu-miR-10b-3p |
| Target Scan | St8sia1 | -21.450685 | 1.14E-06 | 3.66E-05 | mmu-miR-10b-3p |
| Target Scan | Stx19 | -21.551094 | 1.02E-06 | 3.65E-05 | mmu-miR-10b-3p |
| Target Scan | Tacstd2 | -3.1656519 | 0.00264557 | 0.04294489 | mmu-miR-10b-3p |
| Target Scan | Tbx18 | -21.595314 | 9.68E-07 | 3.60E-05 | mmu-miR-10b-3p |
| Target Scan | Vasp | -1.6922905 | 0.00139885 | 0.02326428 | mmu-miR-10b-3p |
| Target Scan | Wasf3 | -20.853832 | 2.26E-06 | 4.90E-05 | mmu-miR-10b-3p |
| Target Scan | Wbp2nl | -21.595313 | 9.68E-07 | 3.60E-05 | mmu-miR-10b-3p |
| Target Scan | Wnt2b | -21.876608 | 6.97E-07 | 3.31E-05 | mmu-miR-10b-3p |
| Target Scan | Zfp114 | -21.904075 | 6.75E-07 | 3.26E-05 | mmu-miR-10b-3p |
| Target Scan | Zfp148 | -1.011897 | 9.30E-04 | 0.01575986 | mmu-miR-10b-3p |
