## Supplemental Table 3-A for "Environmentally Induced Sperm RNAs Transmit Cancer Susceptibility to Offspring in a Mouse Model"

**Table S3.A-Differentially expressed genes in DDT offspring's mammary tumors**

| Name | gene_symbol | FC | pvalue | padj |
|---|---|---|---|---|
| ENSMUSG00000023391 | Dlx2 | -199480.6088 | 1.52E-15 | 3.86E-11 |
| ENSMUSG00000027401 | Tgm3 | -86871.21435 | 1.72E-09 | 2.19E-05 |
| ENSMUSG00000042216 | Sgsm1 | 2.509942065 | 2.28E-07 | 0.00193333 |
| ENSMUSG00000097694 | G730013B05Rik | -19.45114123 | 7.45E-07 | 0.00473604 |
| ENSMUSG00000084803 | 5830444B04Rik | -6.049590811 | 1.11E-05 | 0.05658161 |
| ENSMUSG00000030111 | A2m | 28.34185271 | 1.47E-05 | 0.06242168 |
| ENSMUSG00000029054 | Gabrd | -5.041554233 | 0.00011335 | 0.41149176 |
| ENSMUSG00000085541 | Gm16010 | 12212.49851 | 0.000162426 | 0.43712492 |
| ENSMUSG00000103032 | Gm37121 | -53.43649836 | 0.00018248 | 0.43712492 |
| ENSMUSG00000031125 | 3830403N18Rik | 11.11443911 | 0.000189242 | 0.43712492 |
| ENSMUSG00000103865 | Gm37416 | -25.79354962 | 0.000199947 | 0.43712492 |
| ENSMUSG00000029019 | Nppb | 22.62640202 | 0.000250369 | 0.43712492 |
| ENSMUSG00000024561 | Mbd1 | 1.374785083 | 0.000266744 | 0.43712492 |
| ENSMUSG00000065999 | Zfp985 | -23.94360511 | 0.000269199 | 0.43712492 |
| ENSMUSG00000105703 | Gm43305 | -3.493127845 | 0.000272939 | 0.43712492 |
| ENSMUSG00000057000 | Nxf3 | -23.94009096 | 0.000280514 | 0.43712492 |
| ENSMUSG00000005640 | Insrr | 56.94388272 | 0.000292426 | 0.43712492 |
| ENSMUSG00000028341 | Nr4a3 | -5.600227325 | 0.000333318 | 0.47057142 |
| ENSMUSG00000001497 | Pax9 | 20.67116391 | 0.000511146 | 0.63408707 |
| ENSMUSG00000021614 | Vcan | -2.682277522 | 0.000521191 | 0.63408707 |
| ENSMUSG00000050875 | Minar2 | -16.47980717 | 0.000523998 | 0.63408707 |
| ENSMUSG00000025475 | Adgra1 | -8.030044577 | 0.000578568 | 0.65610621 |
| ENSMUSG00000023443 | Esx1 | -20.25881244 | 0.000593831 | 0.65610621 |
| ENSMUSG00000053297 | AI854703 | 7.378751159 | 0.000645392 | 0.670623 |
| ENSMUSG00000066632 | Pgk1-rs7 | -12828.90925 | 0.00065975 | 0.670623 |
| ENSMUSG00000027082 | Tfpi | 2.213526599 | 0.000809711 | 0.71633464 |
| ENSMUSG00000031465 | Angpt2 | 3.35153119 | 0.000825515 | 0.71633464 |
| ENSMUSG00000024084 | Qpct | -2.137614082 | 0.000861819 | 0.71633464 |
| ENSMUSG00000046186 | Cd109 | -2.297149119 | 0.000866173 | 0.71633464 |
| ENSMUSG00000085791 | Rpl30-ps9 | -160.3908606 | 0.000912319 | 0.71633464 |
| ENSMUSG00000092624 | Gm3654 | 49.95817829 | 0.000953376 | 0.71633464 |
| ENSMUSG00000078490 | Cfap74 | -4.461635223 | 0.000982832 | 0.71633464 |
| ENSMUSG00000013663 | Pten | 1.55275795 | 0.001005172 | 0.71633464 |
| ENSMUSG00000115302 | Gm49394 | -3113935.673 | 0.001007595 | 0.71633464 |
| ENSMUSG00000026725 | Tnn | -13.73442359 | 0.00101148 | 0.71633464 |
| ENSMUSG00000027994 | Mcub | -1.82402405 | 0.001014798 | 0.71633464 |
| ENSMUSG00000071230 | Npw | -21.75256825 | 0.001056248 | 0.72544282 |
| ENSMUSG00000003352 | Cacnb3 | 1.552778001 | 0.001167905 | 0.75205594 |
| ENSMUSG00000027254 | Map1a | -2.591546254 | 0.001168 | 0.75205594 |
| ENSMUSG00000027810 | Eif2a | -1.312233763 | 0.001183781 | 0.75205594 |
| ENSMUSG00000020728 | Cep112 | -1.468619667 | 0.001346886 | 0.8348067 |
| ENSMUSG00000081895 | Rpl17-ps10 | 3.784048167 | 0.001474618 | 0.87551033 |
| ENSMUSG00000034768 | Asb16 | 2.529794385 | 0.001481463 | 0.87551033 |

| | | | | |
|---|---|---|---|---|
| ENSMUSG00000036172 | Cd200r3 | 31.75283568 | 0.001622857 | 0.93208817 |
| ENSMUSG00000116165 | Pdxp | 1.686577731 | 0.001650558 | 0.93208817 |
| ENSMUSG00000033498 | Strc | -7.808266889 | 0.001829042 | 1 |
| ENSMUSG00000074311 | Vmn1r139 | 24.5820524 | 0.002022779 | 1 |
| ENSMUSG00000056758 | Hmga2 | -17.00285243 | 0.002132324 | 1 |
| ENSMUSG00000111202 | Gm48275 | 4.792477794 | 0.002182144 | 1 |
| ENSMUSG00000025198 | Erlin1 | -1.336694522 | 0.002231226 | 1 |
| ENSMUSG00000028602 | Tnfrsf8 | 3.509983927 | 0.002259283 | 1 |
| ENSMUSG00000081965 | Gm11620 | 22.92790119 | 0.002387276 | 1 |
| ENSMUSG00000081604 | Gm11518 | -2.192678927 | 0.002477283 | 1 |
| ENSMUSG00000026751 | Nr5a1 | -14.73571141 | 0.002603572 | 1 |
| ENSMUSG00000071398 | 2410004P03Rik | -5.919735583 | 0.002653304 | 1 |
| ENSMUSG00000024862 | Klc2 | -1.312753268 | 0.002776656 | 1 |
| ENSMUSG00000026740 | Dnajc1 | 1.655185287 | 0.002783358 | 1 |
| ENSMUSG00000039337 | Tex19.2 | -14.73283975 | 0.002800962 | 1 |
| ENSMUSG00000039126 | Prune2 | -1.956917782 | 0.002979977 | 1 |
| ENSMUSG00000117231 | Gm41609 | -27.63779587 | 0.003015348 | 1 |
| ENSMUSG00000032495 | Lrrc2 | -33.23129377 | 0.003042261 | 1 |
| ENSMUSG00000086717 | Gm15655 | -82.87568278 | 0.003408471 | 1 |
| ENSMUSG00000033458 | Fan1 | -1.380732001 | 0.003583931 | 1 |
| ENSMUSG00000107121 | 1110006O24Rik | -3.667612883 | 0.003604938 | 1 |
| ENSMUSG00000085912 | Trp53cor1 | 4.656129168 | 0.003830551 | 1 |
| ENSMUSG00000027863 | Cd2 | 3.602147331 | 0.003896092 | 1 |
| ENSMUSG00000036545 | Adamts2 | -1.889186569 | 0.003962902 | 1 |
| ENSMUSG00000040078 | Ptges3-ps | 1.895770703 | 0.004038709 | 1 |
| ENSMUSG00000074731 | Zfp345 | 6.384060909 | 0.004046817 | 1 |
| ENSMUSG00000110344 | Gm45716 | -3.288419743 | 0.004083403 | 1 |
| ENSMUSG00000028214 | Gem | -2.825070713 | 0.00411262 | 1 |
| ENSMUSG00000041684 | Bivm | 1.247634929 | 0.004313576 | 1 |
| ENSMUSG00000118012 | Gm46620 | -1.733577908 | 0.004317982 | 1 |
| ENSMUSG00000067017 | Capza1-ps1 | 1.672721329 | 0.004347799 | 1 |
| ENSMUSG00000025538 | Sumf2 | -1.331732288 | 0.004453332 | 1 |
| ENSMUSG00000028717 | Tal1 | 2.483313859 | 0.004462695 | 1 |
| ENSMUSG00000100954 | Gm10138 | -2.210480153 | 0.004570582 | 1 |
| ENSMUSG00000079190 | AC133103.1 | 12.33346457 | 0.004725344 | 1 |
| ENSMUSG00000020272 | Stk10 | -1.385966711 | 0.004876265 | 1 |
| ENSMUSG00000068240 | Gm11808 | -1.811389389 | 0.005016106 | 1 |
| ENSMUSG00000052396 | Mogat2 | 3.709449218 | 0.005170955 | 1 |
| ENSMUSG00000056973 | Ces1d | -4.598265095 | 0.005278189 | 1 |
| ENSMUSG00000030154 | Klrb1f | 16.92033844 | 0.005431459 | 1 |
| ENSMUSG00000079387 | Luzp4 | 12.50102236 | 0.005513022 | 1 |
| ENSMUSG00000047986 | Palm3 | -7.490004015 | 0.005516899 | 1 |
| ENSMUSG00000003545 | Fosb | -5.025496946 | 0.005585701 | 1 |
| ENSMUSG00000040037 | Negr1 | -6.537875135 | 0.005591062 | 1 |
| ENSMUSG00000022385 | Gtse1 | 1.487626 | 0.005781073 | 1 |

| | | | | |
|---|---|---|---|---|
| ENSMUSG00000108298 | Gm44226 | 5.963102154 | 0.005807348 | 1 |
| ENSMUSG00000117292 | E330032C10Rik | 12.43388281 | 0.005822617 | 1 |
| ENSMUSG00000055110 | A630012P03Rik | -20.27989165 | 0.005908371 | 1 |
| ENSMUSG00000046694 | Tent5b | -1.615925845 | 0.005919198 | 1 |
| ENSMUSG00000068735 | Trp53i11 | 1.682748994 | 0.005928965 | 1 |
| ENSMUSG00000081855 | Rpl17-ps5 | 3.227741052 | 0.006213064 | 1 |
| ENSMUSG00000024268 | Celf4 | 1.490595969 | 0.006269025 | 1 |
| ENSMUSG00000019932 | Kera | -261581.7047 | 0.006269324 | 1 |
| ENSMUSG00000046897 | Zfp740 | 1.182700414 | 0.006536727 | 1 |
| ENSMUSG00000031576 | Kcnu1 | -4.035040698 | 0.006699295 | 1 |
| ENSMUSG00000032271 | Nnmt | -2.321755903 | 0.006780138 | 1 |
| ENSMUSG00000062762 | Ei24 | 1.173355579 | 0.006799763 | 1 |
| ENSMUSG00000104168 | Gm38250 | 6.625506092 | 0.006861457 | 1 |
| ENSMUSG00000104346 | Pcdhga3 | -2.779293323 | 0.006865815 | 1 |
| ENSMUSG00000115685 | D730044K07Rik | -6.104854482 | 0.00699051 | 1 |
| ENSMUSG00000113786 | Gm49384 | -5.129703312 | 0.007142738 | 1 |
| ENSMUSG00000039405 | Prss23 | -2.014035148 | 0.007321602 | 1 |
| ENSMUSG00000044258 | Ctla2a | 2.160940088 | 0.007362202 | 1 |
| ENSMUSG00000092312 | Zfp419 | 21.70247607 | 0.008241428 | 1 |
| ENSMUSG00000027403 | Tgm6 | -171518.5373 | 0.008396644 | 1 |
| ENSMUSG00000075463 | 4930594M22Rik | -4.450039689 | 0.008592178 | 1 |
| ENSMUSG00000091212 | Krtap11-1 | -163416.2369 | 0.00866115 | 1 |
| ENSMUSG00000057457 | Phex | -5.776153587 | 0.008719187 | 1 |
| ENSMUSG00000104876 | Trdc | -5.314202231 | 0.008739735 | 1 |
| ENSMUSG00000036446 | Lum | -3.58848655 | 0.008776939 | 1 |
| ENSMUSG00000000440 | Pparg | -2.207406866 | 0.008816441 | 1 |
| ENSMUSG00000054057 | A930004D18Rik | 1.530755367 | 0.00892826 | 1 |
| ENSMUSG00000025094 | Slc18a2 | 1.968975994 | 0.009008152 | 1 |
| ENSMUSG00000038612 | Mcl1 | -1.200016578 | 0.009118293 | 1 |
| ENSMUSG00000111692 | Gm49373 | 6.174387363 | 0.009178479 | 1 |
| ENSMUSG00000006574 | Slc4a1 | 20.06356711 | 0.009352826 | 1 |
| ENSMUSG00000079808 | AC168977.1 | 9.638846337 | 0.009474602 | 1 |
| ENSMUSG00000044043 | Pcdhb14 | -1.984341175 | 0.00947805 | 1 |
| ENSMUSG00000038009 | Dnajc22 | 2.944474417 | 0.009576332 | 1 |
| ENSMUSG00000116640 | Gm8670 | -16.56645552 | 0.009595474 | 1 |
| ENSMUSG00000032127 | Vps11 | -1.166186483 | 0.009634709 | 1 |
| ENSMUSG00000053475 | Tnfaip6 | -2.878298172 | 0.010000357 | 1 |
| ENSMUSG00000016619 | Nup50 | 1.170336053 | 0.010112295 | 1 |
| ENSMUSG00000085355 | 3010003L21Rik | 2.185968703 | 0.01057369 | 1 |
| ENSMUSG00000032038 | St3gal4 | -1.428330217 | 0.010668346 | 1 |
| ENSMUSG00000030088 | Aldh1l1 | -2.36820118 | 0.010728742 | 1 |
| ENSMUSG00000102246 | 9430037O13Rik | 14.58984799 | 0.010971562 | 1 |
| ENSMUSG00000035112 | Wnk4 | 2.652673936 | 0.011005235 | 1 |
| ENSMUSG00000056270 | Prr9 | 14669.6762 | 0.011257903 | 1 |
| ENSMUSG00000026475 | Rgs16 | -4.360057627 | 0.011341278 | 1 |

| | | | | |
|---|---|---|---|---|
| ENSMUSG00000022992 | Kansl2 | 1.280849893 | 0.011587353 | 1 |
| ENSMUSG00000060882 | Kcnd2 | -7.731281946 | 0.011697349 | 1 |
| ENSMUSG00000031952 | Chst5 | -2.099280907 | 0.011864411 | 1 |
| ENSMUSG00000118394 | Gm50475 | 3.353857261 | 0.012128095 | 1 |
| ENSMUSG00000091594 | Gm17067 | 10.74458018 | 0.012235728 | 1 |
| ENSMUSG00000039883 | Lrrc17 | -3.145089669 | 0.012338904 | 1 |
| ENSMUSG00000071113 | Mboat4 | -20.22654795 | 0.012489451 | 1 |
| ENSMUSG00000004535 | Tax1bp1 | 1.213122294 | 0.012579807 | 1 |
| ENSMUSG00000001804 | Dsg4 | -95652.04495 | 0.012666578 | 1 |
| ENSMUSG00000112393 | Gm48655 | 23.73260098 | 0.012702945 | 1 |
| ENSMUSG00000061728 | Btnl7-ps | 8.13001624 | 0.012886322 | 1 |
| ENSMUSG00000098369 | Gm2274 | 206.2437813 | 0.013080819 | 1 |
| ENSMUSG00000020583 | Matn3 | 4.81938255 | 0.013087769 | 1 |
| ENSMUSG00000094915 | AC168977.2 | -12.90229695 | 0.013089352 | 1 |
| ENSMUSG00000032199 | Polr2m | 1.165544044 | 0.013261249 | 1 |
| ENSMUSG00000022208 | Jph4 | 2.372721889 | 0.013567788 | 1 |
| ENSMUSG00000050345 | 4930486L24Rik | 17.26762529 | 0.013647561 | 1 |
| ENSMUSG00000016344 | Ppdpf | 1.261204179 | 0.013727754 | 1 |
| ENSMUSG00000027655 | Dhx35 | 1.277911623 | 0.013829726 | 1 |
| ENSMUSG00000035179 | Ppp1r32 | 18.79172158 | 0.01389572 | 1 |
| ENSMUSG00000020990 | Cdkl1 | 3.467196228 | 0.013919929 | 1 |
| ENSMUSG00000109695 | Gm31166 | 12.25545431 | 0.014195685 | 1 |
| ENSMUSG00000022184 | Fbxo4 | 1.440260127 | 0.014348467 | 1 |
| ENSMUSG00000039956 | Mrap | -3.283796081 | 0.014395727 | 1 |
| ENSMUSG00000097643 | A130051J06Rik | 2.562268716 | 0.014400217 | 1 |
| ENSMUSG00000001014 | Icam4 | -11.06009776 | 0.014433893 | 1 |
| ENSMUSG00000067614 | Krt86 | -70754.02223 | 0.014804953 | 1 |
| ENSMUSG00000044854 | 1700056E22Rik | 2.859168209 | 0.015115707 | 1 |
| ENSMUSG00000116220 | A430088P11Rik | 3.114916828 | 0.015406688 | 1 |
| ENSMUSG00000023034 | Nr4a1 | -2.623319296 | 0.015437805 | 1 |
| ENSMUSG00000022893 | Adamts1 | -2.720231337 | 0.015442851 | 1 |
| ENSMUSG00000020009 | Ifngr1 | -1.293264271 | 0.015590491 | 1 |
| ENSMUSG00000004994 | Ccdc130 | -1.350946865 | 0.01562126 | 1 |
| ENSMUSG00000044786 | Zfp36 | -1.700237575 | 0.015733308 | 1 |
| ENSMUSG00000031444 | F10 | 3.448291158 | 0.015759087 | 1 |
| ENSMUSG00000041737 | Tmem45b | -4.74980445 | 0.01585086 | 1 |
| ENSMUSG00000020381 | Mrnip | 1.795525694 | 0.015945718 | 1 |
| ENSMUSG00000087497 | 2810001G20Rik | -1.641069589 | 0.016490399 | 1 |
| ENSMUSG00000045259 | Klhdc9 | -13.14230587 | 0.016622542 | 1 |
| ENSMUSG00000087166 | L1td1 | 56275.06199 | 0.016715545 | 1 |
| ENSMUSG00000053644 | Aldh7a1 | -1.298810537 | 0.016874083 | 1 |
| ENSMUSG00000062515 | Fabp4 | -2.348546039 | 0.017075521 | 1 |
| ENSMUSG00000048583 | Igf2 | -5.563733225 | 0.017077746 | 1 |
| ENSMUSG00000038668 | Lpar1 | -1.902630257 | 0.017165554 | 1 |
| ENSMUSG00000034460 | Six4 | 1.562902466 | 0.017326386 | 1 |

| | | | | |
|---|---|---|---|---|
| ENSMUSG00000069227 | Gprin1 | 3.95493458 | 0.017502638 | 1 |
| ENSMUSG00000026405 | C4bp | 50888.9556 | 0.017612142 | 1 |
| ENSMUSG00000112239 | Gm17823 | 33.59163884 | 0.017658141 | 1 |
| ENSMUSG00000045917 | Tmem268 | 1.284462916 | 0.017739742 | 1 |
| ENSMUSG00000030159 | Clec1b | -4.665148119 | 0.017763581 | 1 |
| ENSMUSG00000072258 | Taf1a | -1.325099297 | 0.017823572 | 1 |
| ENSMUSG00000034783 | Cd207 | -5.739319995 | 0.017981183 | 1 |
| ENSMUSG00000073486 | Gm10518 | -16.57142265 | 0.017981669 | 1 |
| ENSMUSG00000020248 | Nfyb | 1.229114458 | 0.017985058 | 1 |
| ENSMUSG00000005371 | Fbxo11 | 1.175582235 | 0.018018497 | 1 |
| ENSMUSG00000015568 | Lpl | -2.172989324 | 0.018104895 | 1 |
| ENSMUSG00000028693 | Nasp | -1.365494188 | 0.018196506 | 1 |
| ENSMUSG00000033184 | Tmed7 | 1.18108512 | 0.018306515 | 1 |
| ENSMUSG00000057729 | Prtn3 | 3.120186986 | 0.018313332 | 1 |
| ENSMUSG00000038085 | Cnbd2 | -2.08750842 | 0.018371894 | 1 |
| ENSMUSG00000058291 | Zfp68 | 1.158515789 | 0.018537741 | 1 |
| ENSMUSG00000079685 | Ulbp1 | 1.354639169 | 0.018703784 | 1 |
| ENSMUSG00000005045 | Chd5 | -2.358328239 | 0.01871892 | 1 |
| ENSMUSG00000001119 | Col6a1 | -1.815026783 | 0.01873735 | 1 |
| ENSMUSG00000026628 | Atf3 | -2.991058657 | 0.018787219 | 1 |
| ENSMUSG00000022362 | Gm29394 | 7.936184985 | 0.019107162 | 1 |
| ENSMUSG00000043346 | Gm6741 | 66.99146592 | 0.019213202 | 1 |
| ENSMUSG00000032852 | Rspo4 | -49629.68004 | 0.0192178 | 1 |
| ENSMUSG00000000031 | H19 | 4.528498472 | 0.019277581 | 1 |
| ENSMUSG00000086050 | Gm16045 | 16.76256417 | 0.019279027 | 1 |
| ENSMUSG00000059325 | Hopx | -2.289601958 | 0.019315936 | 1 |
| ENSMUSG00000035105 | Egln3 | 2.336122086 | 0.019531813 | 1 |
| ENSMUSG00000081179 | Gm13136 | 5.996456332 | 0.019644034 | 1 |
| ENSMUSG00000063171 | Rps4l | -2.444046041 | 0.019697225 | 1 |
| ENSMUSG00000075551 | Cyp3a41a | 41128.02174 | 0.019736104 | 1 |
| ENSMUSG00000026676 | Ccdc3 | -3.719379425 | 0.019787228 | 1 |
| ENSMUSG00000017588 | Krt27 | 40465.07235 | 0.020076875 | 1 |
| ENSMUSG00000028386 | Slc46a2 | -4.169320239 | 0.020134642 | 1 |
| ENSMUSG00000020875 | Hoxb9 | 3.510838237 | 0.02025213 | 1 |
| ENSMUSG00000117670 | Gm50270 | -11.34551707 | 0.020411893 | 1 |
| ENSMUSG00000117069 | Gm49894 | -3.604100505 | 0.020559984 | 1 |
| ENSMUSG00000028195 | Ccn1 | -2.237828748 | 0.020598732 | 1 |
| ENSMUSG00000059256 | Gzmd | 17.58752868 | 0.020624405 | 1 |
| ENSMUSG00000027763 | Mbnl1 | -1.33715044 | 0.020731113 | 1 |
| ENSMUSG00000075304 | Sp5 | 38954.58853 | 0.020746713 | 1 |
| ENSMUSG00000022255 | Mtdh | 1.196055805 | 0.020949679 | 1 |
| ENSMUSG00000040396 | Abhd13 | 1.183989834 | 0.021153173 | 1 |
| ENSMUSG00000055148 | Klf2 | -1.842486704 | 0.021238198 | 1 |
| ENSMUSG00000026527 | Rgs7 | -9.032468575 | 0.021283953 | 1 |
| ENSMUSG00000101597 | Gm5621 | 14.15348348 | 0.021413706 | 1 |

| | | | | |
|---|---|---|---|---|
| ENSMUSG00000111840 | Gm48832 | -8.725170206 | 0.021452448 | 1 |
| ENSMUSG00000031487 | Brf2 | 1.294996254 | 0.021500879 | 1 |
| ENSMUSG00000011751 | Sptbn4 | 2.430418774 | 0.021507089 | 1 |
| ENSMUSG00000057836 | Xlr3a | 5.841187004 | 0.021550357 | 1 |
| ENSMUSG00000097039 | Pvt1 | 1.443246562 | 0.021675322 | 1 |
| ENSMUSG00000113769 | 5033406O09Rik | 1.813127706 | 0.021682034 | 1 |
| ENSMUSG00000090098 | Alms1-ps2 | 4.655706898 | 0.021704195 | 1 |
| ENSMUSG00000025479 | Cyp2e1 | -4.635920162 | 0.02174928 | 1 |
| ENSMUSG00000014846 | Tppp3 | 1.925682855 | 0.021816883 | 1 |
| ENSMUSG00000118495 | AC161757.1 | -12.8990665 | 0.021998814 | 1 |
| ENSMUSG00000062110 | Scfd2 | -1.382710224 | 0.022025582 | 1 |
| ENSMUSG00000028339 | Col15a1 | -2.194263688 | 0.022034413 | 1 |
| ENSMUSG00000023828 | Slc22a3 | -3.930041017 | 0.022050292 | 1 |
| ENSMUSG00000106831 | Ube2n-ps1 | 1.393812539 | 0.022145253 | 1 |
| ENSMUSG00000039725 | Trp53rka | 1.268849186 | 0.022251479 | 1 |
| ENSMUSG00000052477 | C130026I21Rik | 2.94638328 | 0.022257721 | 1 |
| ENSMUSG00000032839 | Trpc1 | 1.827358817 | 0.022479593 | 1 |
| ENSMUSG00000103731 | Gm17530 | 4.390561578 | 0.023002978 | 1 |
| ENSMUSG00000032818 | Loxhd1 | -9.569558778 | 0.023030517 | 1 |
| ENSMUSG00000028563 | Tm2d1 | 1.322413297 | 0.023205579 | 1 |
| ENSMUSG00000024990 | Rbp4 | -2.534179006 | 0.023389019 | 1 |
| ENSMUSG00000040740 | Slc25a34 | 7.570028705 | 0.023413637 | 1 |
| ENSMUSG00000025738 | Fbxl16 | -2.556348682 | 0.023496049 | 1 |
| ENSMUSG00000113047 | Gm47469 | 1.592777152 | 0.023563744 | 1 |
| ENSMUSG00000073400 | Trim10 | -8.505905331 | 0.023567939 | 1 |
| ENSMUSG00000083853 | Gm15696 | -17.98710958 | 0.02378401 | 1 |
| ENSMUSG00000064225 | Paqr9 | -6.440178285 | 0.023809165 | 1 |
| ENSMUSG00000024190 | Dusp1 | -2.381532757 | 0.023922065 | 1 |
| ENSMUSG00000028885 | Smpdl3b | -1.474402315 | 0.023973671 | 1 |
| ENSMUSG00000020564 | Atxn7l1 | -1.285751922 | 0.024030467 | 1 |
| ENSMUSG00000060572 | Mfap2 | -2.399089919 | 0.024538962 | 1 |
| ENSMUSG00000037447 | Arid5a | -1.483864253 | 0.024699026 | 1 |
| ENSMUSG00000103199 | Gm37648 | -14.72057408 | 0.025047505 | 1 |
| ENSMUSG00000044957 | Pp2d1 | -11.36063212 | 0.025350532 | 1 |
| ENSMUSG00000069310 | H3c3 | 3.025614843 | 0.025453012 | 1 |
| ENSMUSG00000039828 | Wdr70 | 1.25238405 | 0.025573194 | 1 |
| ENSMUSG00000068744 | Psrc1 | 1.890940578 | 0.025791165 | 1 |
| ENSMUSG00000028517 | Plpp3 | -1.368549337 | 0.026071265 | 1 |
| ENSMUSG00000062590 | Armc9 | -1.234496815 | 0.026172964 | 1 |
| ENSMUSG00000025451 | Paip1 | 1.130501303 | 0.026204536 | 1 |
| ENSMUSG00000036766 | Dner | -12.26970516 | 0.026366534 | 1 |
| ENSMUSG00000017466 | Timp2 | -1.283704374 | 0.026607563 | 1 |
| ENSMUSG00000025537 | Phkg1 | -2.544023621 | 0.026874541 | 1 |
| ENSMUSG00000027380 | Acoxl | 15.23029452 | 0.02728139 | 1 |
| ENSMUSG00000040093 | Bmf | 1.500374875 | 0.027412913 | 1 |

| | | | | |
|---|---|---|---|---|
| ENSMUSG00000027573 | Gid8 | 1.17966232 | 0.027427451 | 1 |
| ENSMUSG00000069721 | Krtap3-2 | 24571.66557 | 0.027524693 | 1 |
| ENSMUSG00000014932 | Yes1 | 1.263105405 | 0.027718808 | 1 |
| ENSMUSG00000102759 | Gm10463 | 13.26639476 | 0.02780843 | 1 |
| ENSMUSG00000029819 | Npy | -57.50884724 | 0.027892444 | 1 |
| ENSMUSG00000020241 | Col6a2 | -1.877867539 | 0.027969424 | 1 |
| ENSMUSG00000020326 | Ccng1 | 1.34047737 | 0.028052577 | 1 |
| ENSMUSG00000073008 | Gpr174 | 4.374997173 | 0.028219547 | 1 |
| ENSMUSG00000028825 | Rhd | -9.209353967 | 0.028241837 | 1 |
| ENSMUSG00000085023 | Gm12744 | -2.625174509 | 0.028471338 | 1 |
| ENSMUSG00000027465 | Tbc1d20 | 1.13318584 | 0.028564588 | 1 |
| ENSMUSG00000024937 | Ehbp1l1 | -1.280199259 | 0.028622711 | 1 |
| ENSMUSG00000025418 | Bsnd | 22733.79859 | 0.028737944 | 1 |
| ENSMUSG00000066677 | Ifi208 | -2.880809093 | 0.028740851 | 1 |
| ENSMUSG00000019997 | Ccn2 | -2.024916487 | 0.02896261 | 1 |
| ENSMUSG00000023931 | Efhb | 2.53871159 | 0.029030556 | 1 |
| ENSMUSG00000044361 | BC024139 | -18.44960395 | 0.029250349 | 1 |
| ENSMUSG00000027859 | Ngf | -2.623552688 | 0.029356928 | 1 |
| ENSMUSG00000036611 | Eepd1 | -1.895852568 | 0.02936685 | 1 |
| ENSMUSG00000057069 | Ero1lb | -1.221762428 | 0.02944294 | 1 |
| ENSMUSG00000025461 | Cd163l1 | -5.700727128 | 0.029832944 | 1 |
| ENSMUSG00000090121 | Abhd12b | 20844.41553 | 0.029873989 | 1 |
| ENSMUSG00000040212 | Emp3 | -1.732798791 | 0.030157685 | 1 |
| ENSMUSG00000095180 | Rhox5 | -3.647627479 | 0.030272331 | 1 |
| ENSMUSG00000074256 | Gm10655 | -19.85076608 | 0.030442509 | 1 |
| ENSMUSG00000083044 | Gm12416 | 14.07290866 | 0.030755761 | 1 |
| ENSMUSG00000024513 | Mbd2 | 1.209830862 | 0.030816034 | 1 |
| ENSMUSG00000081669 | Npm3-ps1 | -7.327611912 | 0.030833233 | 1 |
| ENSMUSG00000020251 | Glt8d2 | -1.838279545 | 0.030873656 | 1 |
| ENSMUSG00000032487 | Ptgs2 | -6.19511827 | 0.030898829 | 1 |
| ENSMUSG00000091971 | Hspa1a | -3.736361663 | 0.030922259 | 1 |
| ENSMUSG00000104830 | 5830487J09Rik | 3.060333831 | 0.030925165 | 1 |
| ENSMUSG00000029227 | Fip1l1 | 1.153689527 | 0.030925889 | 1 |
| ENSMUSG00000045362 | Tnfrsf26 | 2.763253973 | 0.031016677 | 1 |
| ENSMUSG00000111929 | Gm48780 | 27.67070965 | 0.0310341 | 1 |
| ENSMUSG00000101995 | Gm29480 | 13.25436783 | 0.031153197 | 1 |
| ENSMUSG00000025825 | Iscu | -1.189451132 | 0.031185062 | 1 |
| ENSMUSG00000107896 | Gm8719 | 29.73010079 | 0.031263616 | 1 |
| ENSMUSG00000058173 | Smco4 | -1.619088888 | 0.031353086 | 1 |
| ENSMUSG00000051495 | Irf2bp2 | -1.251033867 | 0.031523927 | 1 |
| ENSMUSG00000026200 | Glb1l | -1.454022247 | 0.031615189 | 1 |
| ENSMUSG00000022878 | Adipoq | -3.22265544 | 0.031799011 | 1 |
| ENSMUSG00000109713 | Pvrig | 2.1539891 | 0.031847958 | 1 |
| ENSMUSG00000116780 | Gm3417 | 20.84439253 | 0.031963235 | 1 |
| ENSMUSG00000076615 | Ighg3 | 61.29205346 | 0.032102324 | 1 |

| | | | | |
|---|---|---|---|---|
| ENSMUSG00000061780 | Cfd | -3.35357079 | 0.032110148 | 1 |
| ENSMUSG00000042515 | Pwwp3b | 2.405790123 | 0.032154772 | 1 |
| ENSMUSG00000020264 | Slc36a2 | -3.266705169 | 0.0323472 | 1 |
| ENSMUSG00000067750 | Khdc1a | 15.55702094 | 0.03255657 | 1 |
| ENSMUSG00000032322 | Pstpip1 | -1.663670539 | 0.032582423 | 1 |
| ENSMUSG00000096433 | Zfp994 | 1.353228396 | 0.03266674 | 1 |
| ENSMUSG00000025185 | Loxl4 | -2.063996781 | 0.032791613 | 1 |
| ENSMUSG00000050967 | Creg2 | 1.878943018 | 0.03285518 | 1 |
| ENSMUSG00000113019 | Gm47467 | -5.665075431 | 0.032891191 | 1 |
| ENSMUSG00000042379 | Esm1 | 2.198591364 | 0.032958023 | 1 |
| ENSMUSG00000085766 | 2810430I11Rik | -2.859584962 | 0.033339715 | 1 |
| ENSMUSG00000096986 | 4930509E16Rik | 8.830186631 | 0.033347562 | 1 |
| ENSMUSG00000045515 | Pou3f3 | 16905.49086 | 0.033557984 | 1 |
| ENSMUSG00000104682 | Gm42636 | 2.522702787 | 0.033629656 | 1 |
| ENSMUSG00000003348 | Mob3a | -1.367419546 | 0.033635697 | 1 |
| ENSMUSG00000038936 | Sccpdh | 1.267285097 | 0.033847981 | 1 |
| ENSMUSG00000001661 | Hoxc6 | -2.175410431 | 0.03399998 | 1 |
| ENSMUSG00000028108 | Ecm1 | -1.899874851 | 0.034287336 | 1 |
| ENSMUSG00000106946 | Gm42856 | 15.28919404 | 0.034348157 | 1 |
| ENSMUSG00000046442 | Ppm1e | -3.955273736 | 0.034378249 | 1 |
| ENSMUSG00000061769 | Klra6 | 11.86588731 | 0.034413163 | 1 |
| ENSMUSG00000054309 | Cpsf3 | -1.17539214 | 0.034540989 | 1 |
| ENSMUSG00000029314 | Gpat3 | -1.6970311 | 0.034552946 | 1 |
| ENSMUSG00000022526 | Zfp251 | 1.402794701 | 0.034591256 | 1 |
| ENSMUSG00000090248 | Gm14027 | 22.80193108 | 0.034741816 | 1 |
| ENSMUSG00000049287 | Iba57 | 1.248385909 | 0.034826593 | 1 |
| ENSMUSG00000032492 | Pth1r | 3.17059223 | 0.034875383 | 1 |
| ENSMUSG00000029776 | Hibadh | 1.190331576 | 0.035196475 | 1 |
| ENSMUSG00000102748 | Pcdhgb2 | -2.81779951 | 0.035240187 | 1 |
| ENSMUSG00000047511 | Olfr1396 | -11.35063337 | 0.035438252 | 1 |
| ENSMUSG00000045838 | Ccdc9b | 1.433267824 | 0.035448865 | 1 |
| ENSMUSG00000104052 | Gm38125 | 36.0331667 | 0.035453333 | 1 |
| ENSMUSG00000030498 | Gas2 | 1.472709097 | 0.0358342 | 1 |
| ENSMUSG00000032307 | Ube2q2 | 1.239144028 | 0.035889113 | 1 |
| ENSMUSG00000096917 | 2500002B13Rik | 2.902815908 | 0.036063221 | 1 |
| ENSMUSG00000022607 | Ptk2 | 1.162704707 | 0.036162504 | 1 |
| ENSMUSG00000082588 | Gm15443 | -17.99410905 | 0.036247668 | 1 |
| ENSMUSG00000037214 | Thap1 | 1.235684327 | 0.036266347 | 1 |
| ENSMUSG00000025509 | Pnpla2 | -1.412062236 | 0.036590568 | 1 |
| ENSMUSG00000027204 | Fbn1 | -1.725505612 | 0.036620896 | 1 |
| ENSMUSG00000039841 | Zfp800 | -1.286067719 | 0.036649656 | 1 |
| ENSMUSG00000100865 | Gm9320 | -12.91278687 | 0.036843824 | 1 |
| ENSMUSG00000084880 | Tomm6os | -2.916342909 | 0.036852581 | 1 |
| ENSMUSG00000029092 | D5Ertd615e | 7.510685666 | 0.036866443 | 1 |
| ENSMUSG00000031443 | F7 | 3.484184452 | 0.036888778 | 1 |

| | | | | |
|---|---|---|---|---|
| ENSMUSG00000118210 | Gm50394 | -22.07076839 | 0.036890074 | 1 |
| ENSMUSG00000098934 | Gm18853 | -4.523635538 | 0.036899534 | 1 |
| ENSMUSG00000117294 | Gm49839 | 10.6087554 | 0.036917076 | 1 |
| ENSMUSG00000029658 | Wdr95 | 13.03704565 | 0.037076703 | 1 |
| ENSMUSG00000029830 | Svopl | 4.564859154 | 0.037807031 | 1 |
| ENSMUSG00000024797 | Vps51 | -1.225268178 | 0.03788183 | 1 |
| ENSMUSG00000024642 | Tle4 | -1.351219498 | 0.037881962 | 1 |
| ENSMUSG00000082480 | Gm11687 | 13.83599834 | 0.038059688 | 1 |
| ENSMUSG00000086416 | Gm14002 | 9.195222071 | 0.038067023 | 1 |
| ENSMUSG00000103065 | Gm20236 | 10.79689889 | 0.038122143 | 1 |
| ENSMUSG00000103720 | Gm37094 | 10.79689889 | 0.038122143 | 1 |
| ENSMUSG00000032549 | Rab6b | -1.442750548 | 0.038147119 | 1 |
| ENSMUSG00000074715 | Ccl28 | 7.591504317 | 0.038192056 | 1 |
| ENSMUSG00000037010 | Apln | 1.949315617 | 0.038203954 | 1 |
| ENSMUSG00000001436 | Slc19a1 | 1.284015843 | 0.038248779 | 1 |
| ENSMUSG00000028788 | Ptp4a2 | 1.131604108 | 0.038425725 | 1 |
| ENSMUSG00000002265 | Peg3 | 2.629313688 | 0.038440996 | 1 |
| ENSMUSG00000002944 | Cd36 | -2.409014029 | 0.038519334 | 1 |
| ENSMUSG00000106296 | 4632404M16Rik | -3.131983241 | 0.038541759 | 1 |
| ENSMUSG00000099759 | 1700030C10Rik | 5.174412754 | 0.03857882 | 1 |
| ENSMUSG00000097111 | Peak1os | -6.06726848 | 0.038750309 | 1 |
| ENSMUSG00000107577 | Gm44103 | -2.023251063 | 0.038872601 | 1 |
| ENSMUSG00000022577 | Ly6h | -27.66805681 | 0.038961023 | 1 |
| ENSMUSG00000009108 | Gnat2 | 2.422185517 | 0.038988755 | 1 |
| ENSMUSG00000029661 | Col1a2 | -2.164906831 | 0.039149909 | 1 |
| ENSMUSG00000046312 | Myorg | -1.796979495 | 0.039227763 | 1 |
| ENSMUSG00000020101 | Vsir | 1.290732749 | 0.039270634 | 1 |
| ENSMUSG00000026814 | Eng | 1.433275988 | 0.039303887 | 1 |
| ENSMUSG00000028072 | Ntrk1 | -7.367326966 | 0.039389504 | 1 |
| ENSMUSG00000026785 | Pkn3 | -1.34570315 | 0.039648341 | 1 |
| ENSMUSG00000091230 | H1f11-ps | 15.97763294 | 0.03973827 | 1 |
| ENSMUSG00000070867 | Trabd2b | -1.862669711 | 0.039889273 | 1 |
| ENSMUSG00000107017 | Gm43196 | -6.614049146 | 0.040005286 | 1 |
| ENSMUSG00000026841 | Fibcd1 | -8.499055523 | 0.040005706 | 1 |
| ENSMUSG00000112160 | BC024063 | 1.675555207 | 0.04005435 | 1 |
| ENSMUSG00000061544 | Zfp229 | 1.263140422 | 0.040138352 | 1 |
| ENSMUSG00000075704 | Txnrd2 | -1.353489731 | 0.040181371 | 1 |
| ENSMUSG00000095872 | Gm15128 | -6.619474936 | 0.040371922 | 1 |
| ENSMUSG00000106818 | Gm43790 | 10.17955638 | 0.040391806 | 1 |
| ENSMUSG00000040405 | Havcr1 | 16.11233382 | 0.040400551 | 1 |
| ENSMUSG00000020422 | Tns3 | -1.61095259 | 0.040401507 | 1 |
| ENSMUSG00000059895 | Ptp4a3 | 1.587145546 | 0.04047773 | 1 |
| ENSMUSG00000039639 | Kcne1 | 20.86554091 | 0.040500605 | 1 |
| ENSMUSG00000027559 | Car3 | -3.099195024 | 0.040581054 | 1 |
| ENSMUSG00000046546 | Fam43a | 1.550448613 | 0.040903574 | 1 |

| | | | | |
|---|---|---|---|---|
| ENSMUSG00000115220 | Gm49768 | -2.677199682 | 0.041016784 | 1 |
| ENSMUSG00000116207 | Nnt | -179.725212 | 0.041062643 | 1 |
| ENSMUSG00000023132 | Gzma | -2.616236064 | 0.04110662 | 1 |
| ENSMUSG00000026043 | Col3a1 | -1.976888941 | 0.041247561 | 1 |
| ENSMUSG00000026012 | Cd28 | 3.325130374 | 0.041264762 | 1 |
| ENSMUSG00000085971 | Gm15411 | 4.399746681 | 0.041295221 | 1 |
| ENSMUSG00000102712 | Gm37758 | 2.367993567 | 0.041532532 | 1 |
| ENSMUSG00000033287 | Kctd17 | 1.526333986 | 0.041635119 | 1 |
| ENSMUSG00000000938 | Hoxa10 | -4.745808091 | 0.041905238 | 1 |
| ENSMUSG00000022270 | Retreg1 | 1.626272824 | 0.041910737 | 1 |
| ENSMUSG00000117810 | Gm8934 | -10.40536504 | 0.041914232 | 1 |
| ENSMUSG00000027663 | Zmat3 | 1.389139284 | 0.041922461 | 1 |
| ENSMUSG00000100514 | Gm12960 | 2.753206222 | 0.042067357 | 1 |
| ENSMUSG00000102691 | Gm37780 | -3.116163302 | 0.042086381 | 1 |
| ENSMUSG00000005907 | Pex1 | -1.259599246 | 0.042127962 | 1 |
| ENSMUSG00000043336 | Filip1l | -1.358268224 | 0.042142423 | 1 |
| ENSMUSG00000091055 | Siglec15 | -5.966776221 | 0.04242091 | 1 |
| ENSMUSG00000028049 | Scamp3 | -1.151166536 | 0.042697817 | 1 |
| ENSMUSG00000040302 | Rbm48 | -1.29903064 | 0.04294466 | 1 |
| ENSMUSG00000111028 | Gm5922 | -23.94843979 | 0.042996024 | 1 |
| ENSMUSG00000027423 | Snx5 | 1.146860517 | 0.043076966 | 1 |
| ENSMUSG00000040640 | Erc2 | 1.801149886 | 0.043220472 | 1 |
| ENSMUSG00000066178 | 6030445D17Rik | -3.859824558 | 0.043510209 | 1 |
| ENSMUSG00000039754 | Alkbh4 | -1.374390896 | 0.043527164 | 1 |
| ENSMUSG00000115546 | Gm49077 | -38.66591663 | 0.043681091 | 1 |
| ENSMUSG00000103162 | Gm38147 | 10.86327436 | 0.043815262 | 1 |
| ENSMUSG00000003355 | Fkbp11 | 1.420386419 | 0.044356561 | 1 |
| ENSMUSG00000011382 | Dhdh | -1.575133852 | 0.044440348 | 1 |
| ENSMUSG00000068923 | Syt11 | -1.565158084 | 0.044871619 | 1 |
| ENSMUSG00000027947 | Il6ra | -1.363476893 | 0.044878435 | 1 |
| ENSMUSG00000006906 | Stambp | -1.200246235 | 0.045155449 | 1 |
| ENSMUSG00000107944 | Gm44280 | -3.588799712 | 0.045263732 | 1 |
| ENSMUSG00000103094 | Gm37558 | 13.36844086 | 0.045303381 | 1 |
| ENSMUSG00000046908 | Ltb4r1 | 1.972184123 | 0.04540165 | 1 |
| ENSMUSG00000033589 | Reep4 | 1.3329931 | 0.04543704 | 1 |
| ENSMUSG00000086631 | Gm12784 | 7.170866432 | 0.04577341 | 1 |
| ENSMUSG00000001334 | Fndc5 | 3.247113027 | 0.045790894 | 1 |
| ENSMUSG00000025497 | Cdhr5 | 6.807214377 | 0.045826801 | 1 |
| ENSMUSG00000048445 | Ccdc57 | -1.484508661 | 0.045840223 | 1 |
| ENSMUSG00000046585 | Cfap58 | -7.364662126 | 0.045866566 | 1 |
| ENSMUSG00000024912 | Fosl1 | -2.111509148 | 0.045957674 | 1 |
| ENSMUSG00000106341 | Gm43330 | 11.07556835 | 0.045982228 | 1 |
| ENSMUSG00000108308 | Gm45218 | -23.96986217 | 0.046103622 | 1 |
| ENSMUSG00000002324 | Rec8 | -3.751910637 | 0.047032967 | 1 |
| ENSMUSG00000028730 | Cfap57 | 8.063818471 | 0.047068267 | 1 |

| | | | | |
|---|---|---|---|---|
| ENSMUSG00000085957 | Syna | 4.798978501 | 0.047090622 | 1 |
| ENSMUSG00000090610 | Gm3571 | -9.208703438 | 0.04712013 | 1 |
| ENSMUSG00000030609 | Aen | 1.214378952 | 0.047147907 | 1 |
| ENSMUSG00000027427 | Polr3f | 1.222322083 | 0.047170317 | 1 |
| ENSMUSG00000107937 | Gm44200 | 18.85759848 | 0.047174644 | 1 |
| ENSMUSG00000067787 | Blcap | 1.219283906 | 0.047218378 | 1 |
| ENSMUSG00000099465 | Gm3830 | 8.841835442 | 0.047299174 | 1 |
| ENSMUSG00000050075 | Gpr171 | -1.949538092 | 0.047378057 | 1 |
| ENSMUSG00000027905 | Ddx20 | -1.169885819 | 0.047514297 | 1 |
| ENSMUSG00000031535 | Dkk4 | 9389.39747 | 0.047620193 | 1 |
| ENSMUSG00000019804 | Snx3 | 1.171761417 | 0.047699547 | 1 |
| ENSMUSG00000026131 | Dst | -1.365144545 | 0.047847879 | 1 |
| ENSMUSG00000022313 | Utp23 | 1.154567061 | 0.047897292 | 1 |
| ENSMUSG00000097245 | Gm5421 | -3.0936528 | 0.047983312 | 1 |
| ENSMUSG00000089998 | Phtf1os | 2.71520839 | 0.048000881 | 1 |
| ENSMUSG00000061731 | Ext1 | -1.234981987 | 0.048072409 | 1 |
| ENSMUSG00000086191 | Zfp652os | -9.458468766 | 0.048119849 | 1 |
| ENSMUSG00000010830 | Kdelr3 | 1.830253318 | 0.048352502 | 1 |
| ENSMUSG00000066361 | Serpina3c | -3.987128527 | 0.048590962 | 1 |
| ENSMUSG00000032911 | Cspg4 | -1.931538157 | 0.048714861 | 1 |
| ENSMUSG00000083282 | Ctsf | -1.511292515 | 0.048793723 | 1 |
| ENSMUSG00000116048 | Septin2 | 1.455351691 | 0.048820214 | 1 |
| ENSMUSG00000070335 | Krtap9-1 | 8933.483422 | 0.048842487 | 1 |
| ENSMUSG00000038179 | Slamf7 | -1.397933725 | 0.049207756 | 1 |
| ENSMUSG00000072952 | Gm5878 | -7.367076164 | 0.049209405 | 1 |
| ENSMUSG00000079177 | Fam228a | -3.174659263 | 0.049267814 | 1 |
| ENSMUSG00000044991 | Shld1 | -1.483968158 | 0.049450933 | 1 |
| ENSMUSG00000049928 | Glp2r | -7.975042239 | 0.049753642 | 1 |
| ENSMUSG00000079022 | Col22a1 | -3.762354094 | 0.049756443 | 1 |
