## Supplemental Table 3-B for "Environmentally Induced Sperm RNAs Transmit Cancer Susceptibility to Offspring in a Mouse Model"

**Table S3.B-Differentially expressed genes in DDT-RNA offspring's mammary tumors**

| Name | gene_symbol | FC | pvalue | padj |
|---|---|---|---|---|
| ENSMUSG00000116579 | Gm49701 | 10819091 | 1.10E-29 | 2.81E-25 |
| ENSMUSG00000030205 | Gprc5d | -8.471E+10 | 4.98E-12 | 6.33E-08 |
| ENSMUSG00000038943 | Prc1 | 2.41942929 | 8.69E-12 | 7.36E-08 |
| ENSMUSG00000021091 | Serpina3n | -10.676195 | 7.58E-11 | 4.82E-07 |
| ENSMUSG00000038252 | Ncapd2 | 2.16003932 | 1.44E-10 | 6.52E-07 |
| ENSMUSG00000033581 | Igf2bp2 | -4.6782643 | 1.54E-10 | 6.52E-07 |
| ENSMUSG00000015354 | Pcolce2 | -5.9220488 | 2.81E-10 | 1.02E-06 |
| ENSMUSG00000039252 | Lgi2 | -13.415616 | 5.18E-10 | 1.65E-06 |
| ENSMUSG00000035683 | Melk | 2.68723111 | 6.72E-10 | 1.90E-06 |
| ENSMUSG00000047878 | A4galt | -3.1894518 | 1.54E-09 | 3.92E-06 |
| ENSMUSG00000020598 | Nrcam | -12.595114 | 1.81E-09 | 4.17E-06 |
| ENSMUSG00000051048 | P4ha3 | -5.415633 | 2.35E-09 | 4.97E-06 |
| ENSMUSG00000073530 | Pappa2 | -69.928247 | 3.44E-09 | 6.71E-06 |
| ENSMUSG00000072941 | Sod3 | -7.0492189 | 3.75E-09 | 6.81E-06 |
| ENSMUSG00000062380 | Tubb3 | -16.725706 | 4.12E-09 | 6.98E-06 |
| ENSMUSG00000035365 | Parpbp | 2.5940479 | 5.21E-09 | 8.27E-06 |
| ENSMUSG00000045165 | AI467606 | -2.3278585 | 9.41E-09 | 1.38E-05 |
| ENSMUSG00000050578 | Mmp13 | -7.1256968 | 9.79E-09 | 1.38E-05 |
| ENSMUSG00000027360 | Hdc | -39.924478 | 1.33E-08 | 1.77E-05 |
| ENSMUSG00000036356 | Csgalnact1 | -4.1760116 | 1.61E-08 | 2.05E-05 |
| ENSMUSG00000005470 | Asf1b | 1.98458624 | 2.22E-08 | 2.68E-05 |
| ENSMUSG00000027994 | Mcub | -2.5802971 | 3.03E-08 | 3.50E-05 |
| ENSMUSG00000055612 | Cdca7 | 3.00588275 | 3.18E-08 | 3.51E-05 |
| ENSMUSG00000006014 | Prg4 | -5.8284297 | 3.64E-08 | 3.74E-05 |
| ENSMUSG00000031906 | Smpd3 | -17.410573 | 3.68E-08 | 3.74E-05 |
| ENSMUSG00000029771 | Irf5 | -1.8712799 | 3.96E-08 | 3.87E-05 |
| ENSMUSG00000062515 | Fabp4 | -7.107977 | 4.49E-08 | 4.13E-05 |
| ENSMUSG00000031425 | Plp1 | -8.1898961 | 4.55E-08 | 4.13E-05 |
| ENSMUSG00000023015 | Racgap1 | 2.10233641 | 5.00E-08 | 4.38E-05 |
| ENSMUSG00000002289 | Angptl4 | -2.6227925 | 5.68E-08 | 4.59E-05 |
| ENSMUSG00000037725 | Ckap2 | 2.22870267 | 5.77E-08 | 4.59E-05 |
| ENSMUSG00000009687 | Fxyd5 | -2.5264222 | 5.78E-08 | 4.59E-05 |
| ENSMUSG00000022034 | Esco2 | 2.74335698 | 6.25E-08 | 4.81E-05 |
| ENSMUSG00000001131 | Timp1 | -5.0194859 | 8.43E-08 | 6.30E-05 |
| ENSMUSG00000053797 | Krt16 | -93.06501 | 9.59E-08 | 6.85E-05 |
| ENSMUSG00000079685 | Ulbp1 | 2.00580086 | 9.93E-08 | 6.85E-05 |
| ENSMUSG00000034220 | Gpc1 | -2.6352538 | 9.98E-08 | 6.85E-05 |
| ENSMUSG00000097789 | Gm2115 | -13.786682 | 1.05E-07 | 7.05E-05 |
| ENSMUSG00000016496 | Cd274 | -2.2252168 | 1.08E-07 | 7.05E-05 |
| ENSMUSG00000035769 | Xylb | 2.2128852 | 1.30E-07 | 8.14E-05 |
| ENSMUSG00000059456 | Ptk2b | -1.7104405 | 1.31E-07 | 8.14E-05 |
| ENSMUSG00000048905 | Bnip5 | 4.21315715 | 1.38E-07 | 8.18E-05 |
| ENSMUSG00000031391 | L1cam | -4.453976 | 1.41E-07 | 8.18E-05 |

| | | | | |
|---|---|---|---|---|
| ENSMUSG00000020101 | Vsir | -1.9156982 | 1.42E-07 | 8.18E-05 |
| ENSMUSG00000021714 | Cenpk | 2.29082613 | 1.63E-07 | 9.20E-05 |
| ENSMUSG00000049723 | Mmp12 | -36.082689 | 1.82E-07 | 0.00010043 |
| ENSMUSG00000019813 | Cep57l1 | 1.85154594 | 1.86E-07 | 0.00010043 |
| ENSMUSG00000015568 | Lpl | -5.5263465 | 1.93E-07 | 0.0001024 |
| ENSMUSG00000026656 | Fcgr2b | -3.2536164 | 1.99E-07 | 0.00010341 |
| ENSMUSG00000029306 | Ibsp | -156.5943 | 2.17E-07 | 0.00011045 |
| ENSMUSG00000063415 | Cyp26b1 | -7.8108697 | 2.51E-07 | 0.00012393 |
| ENSMUSG00000002221 | Paxip1 | 1.52308793 | 2.59E-07 | 0.00012393 |
| ENSMUSG00000034906 | Ncaph | 2.27633335 | 2.63E-07 | 0.00012393 |
| ENSMUSG00000026509 | Capn2 | -1.3693826 | 2.63E-07 | 0.00012393 |
| ENSMUSG00000029223 | Uchl1 | -5.579565 | 2.81E-07 | 0.00012961 |
| ENSMUSG00000115338 | Pnp | -1.7606843 | 2.97E-07 | 0.00013494 |
| ENSMUSG00000046169 | Adamts6 | -3.2074842 | 3.07E-07 | 0.00013706 |
| ENSMUSG00000002944 | Cd36 | -8.6826388 | 3.66E-07 | 0.00016042 |
| ENSMUSG00000071847 | Apcdd1 | -7.4989562 | 3.85E-07 | 0.00016311 |
| ENSMUSG00000032815 | Fanca | 2.13426346 | 3.85E-07 | 0.00016311 |
| ENSMUSG00000028068 | Iqgap3 | 2.85352895 | 4.67E-07 | 0.0001947 |
| ENSMUSG00000027654 | Fam83d | 2.33918227 | 5.29E-07 | 0.00021702 |
| ENSMUSG00000029816 | Gpnmb | -39.147378 | 5.47E-07 | 0.00021855 |
| ENSMUSG00000031262 | Cenpi | 2.69933988 | 5.51E-07 | 0.00021855 |
| ENSMUSG00000023224 | Serping1 | -3.9138221 | 5.59E-07 | 0.00021855 |
| ENSMUSG00000072980 | Oip5 | 2.37105969 | 5.89E-07 | 0.00022662 |
| ENSMUSG00000058290 | Espl1 | 1.87975334 | 6.59E-07 | 0.00024977 |
| ENSMUSG00000025912 | Mybl1 | 2.5376782 | 7.07E-07 | 0.00026168 |
| ENSMUSG00000048865 | Arhgap30 | -1.7477185 | 7.12E-07 | 0.00026168 |
| ENSMUSG00000022488 | Nckap1l | -1.7821874 | 7.21E-07 | 0.00026168 |
| ENSMUSG00000024912 | Fosl1 | -12.336524 | 7.39E-07 | 0.00026434 |
| ENSMUSG00000062345 | Serpinb2 | -216.1795 | 7.54E-07 | 0.00026626 |
| ENSMUSG00000001642 | Akr1b3 | -2.1022605 | 7.94E-07 | 0.00027629 |
| ENSMUSG00000000627 | Sema4f | -11.266435 | 8.15E-07 | 0.00027998 |
| ENSMUSG00000039396 | Neil3 | 2.2233982 | 8.48E-07 | 0.00028728 |
| ENSMUSG00000034349 | Smc4 | 1.89600224 | 9.73E-07 | 0.0003255 |
| ENSMUSG00000021319 | Sfrp4 | -8.1603933 | 1.02E-06 | 0.00033554 |
| ENSMUSG00000025355 | Mmp19 | -7.8590419 | 1.06E-06 | 0.00034434 |
| ENSMUSG00000020099 | Unc5b | -2.3295806 | 1.08E-06 | 0.00034434 |
| ENSMUSG00000097156 | Gm3764 | 41.2496468 | 1.08E-06 | 0.00034434 |
| ENSMUSG00000057522 | Spop | -1.4253509 | 1.14E-06 | 0.00035718 |
| ENSMUSG00000026274 | Pask | 2.00992932 | 1.17E-06 | 0.00035954 |
| ENSMUSG00000049932 | H2ax | 2.33934169 | 1.17E-06 | 0.00035954 |
| ENSMUSG00000057836 | Xlr3a | -52.839027 | 1.20E-06 | 0.00036006 |
| ENSMUSG00000015880 | Ncapg | 2.27506803 | 1.20E-06 | 0.00036006 |
| ENSMUSG00000028175 | Depdc1a | 2.52559315 | 1.30E-06 | 0.00038388 |
| ENSMUSG00000074195 | Clca4b | 3495064564 | 1.34E-06 | 0.00039239 |
| ENSMUSG00000008843 | Cldn13 | 3446117580 | 1.37E-06 | 0.00039469 |

| | | | | |
|---|---|---|---|---|
| ENSMUSG00000031443 | F7 | -13.919722 | 1.39E-06 | 0.00039558 |
| ENSMUSG00000036545 | Adamts2 | -2.8891638 | 1.55E-06 | 0.00043884 |
| ENSMUSG00000021260 | Hhipl1 | -3.528662 | 1.63E-06 | 0.00045408 |
| ENSMUSG00000040747 | Cd53 | -2.6703527 | 1.76E-06 | 0.00048546 |
| ENSMUSG00000023505 | Cdca3 | 1.97284381 | 1.79E-06 | 0.0004902 |
| ENSMUSG00000029103 | Lrpap1 | -1.6177887 | 1.88E-06 | 0.0005069 |
| ENSMUSG00000027326 | Knl1 | 2.53519397 | 1.97E-06 | 0.00052576 |
| ENSMUSG00000060131 | Atp8b4 | -3.369825 | 2.04E-06 | 0.00053429 |
| ENSMUSG00000003779 | Kif20a | 2.23457956 | 2.04E-06 | 0.00053429 |
| ENSMUSG00000022098 | Bmp1 | -2.0124667 | 2.07E-06 | 0.0005362 |
| ENSMUSG00000001517 | Foxm1 | 2.1658945 | 2.27E-06 | 0.00058382 |
| ENSMUSG00000027715 | Ccna2 | 2.26641793 | 2.35E-06 | 0.00059668 |
| ENSMUSG00000051235 | Gen1 | 1.96022382 | 2.41E-06 | 0.00060712 |
| ENSMUSG00000026039 | Sgo2a | 2.31891503 | 2.51E-06 | 0.00062585 |
| ENSMUSG00000041351 | Rap1gap | 2.51765865 | 2.66E-06 | 0.00065176 |
| ENSMUSG00000027306 | Nusap1 | 2.20374318 | 2.67E-06 | 0.00065176 |
| ENSMUSG00000027408 | Cpxm1 | -3.6565851 | 2.73E-06 | 0.00065987 |
| ENSMUSG00000031004 | Mki67 | 2.51497907 | 2.82E-06 | 0.00067146 |
| ENSMUSG00000033952 | Aspm | 2.64555785 | 2.83E-06 | 0.00067146 |
| ENSMUSG00000031444 | F10 | -8.7599657 | 2.90E-06 | 0.00068194 |
| ENSMUSG00000061838 | Suclg2 | 1.39572218 | 2.99E-06 | 0.00069723 |
| ENSMUSG00000020901 | Pik3r5 | -2.7537722 | 3.06E-06 | 0.00070601 |
| ENSMUSG00000021939 | Ctsb | -2.1161058 | 3.27E-06 | 0.0007431 |
| ENSMUSG00000052316 | Lrrc15 | -22.652043 | 3.28E-06 | 0.0007431 |
| ENSMUSG00000027699 | Ect2 | 2.22564556 | 3.45E-06 | 0.00077235 |
| ENSMUSG00000036825 | Ssx2ip | 1.44070213 | 3.48E-06 | 0.00077235 |
| ENSMUSG00000029414 | Kntc1 | 2.22452959 | 3.52E-06 | 0.00077235 |
| ENSMUSG00000036256 | Igfbp7 | -2.2811859 | 3.53E-06 | 0.00077235 |
| ENSMUSG00000034883 | Lrr1 | 3.52774118 | 3.58E-06 | 0.00077235 |
| ENSMUSG00000033033 | Calhm2 | -1.831413 | 3.59E-06 | 0.00077235 |
| ENSMUSG00000027358 | Bmp2 | -17.141963 | 3.87E-06 | 0.00082708 |
| ENSMUSG00000035493 | Tgfbi | -1.6526883 | 4.25E-06 | 0.00089951 |
| ENSMUSG00000042489 | Clspn | 1.98153386 | 4.41E-06 | 0.00092045 |
| ENSMUSG00000005124 | Ccn4 | -3.5906517 | 4.42E-06 | 0.00092045 |
| ENSMUSG00000104806 | Gm42566 | 20.8889951 | 4.46E-06 | 0.00092112 |
| ENSMUSG00000038910 | Plcl2 | -1.7260758 | 4.59E-06 | 0.00094046 |
| ENSMUSG00000001021 | S100a3 | -18.764932 | 4.73E-06 | 0.00096243 |
| ENSMUSG00000026915 | Strbp | 1.70031547 | 4.83E-06 | 0.00097409 |
| ENSMUSG00000020185 | E2f7 | 2.71095143 | 5.33E-06 | 0.00106569 |
| ENSMUSG00000023940 | Sgo1 | 2.33950089 | 5.39E-06 | 0.0010708 |
| ENSMUSG00000012443 | Kif11 | 2.49763117 | 5.51E-06 | 0.00108552 |
| ENSMUSG00000006398 | Cdc20 | 2.2538683 | 5.56E-06 | 0.00108763 |
| ENSMUSG00000044934 | Zfp367 | 1.61856747 | 5.62E-06 | 0.00108944 |
| ENSMUSG00000050721 | Plekho2 | -2.1106251 | 5.79E-06 | 0.00111438 |
| ENSMUSG00000038179 | Slamf7 | -2.1114247 | 6.09E-06 | 0.00116281 |

| | | | | |
|---|---|---|---|---|
| ENSMUSG00000026872 | Zeb2 | -2.6440099 | 6.39E-06 | 0.00120518 |
| ENSMUSG00000017499 | Cdc6 | 1.92065939 | 6.40E-06 | 0.00120518 |
| ENSMUSG00000032254 | Kif23 | 2.33228192 | 6.68E-06 | 0.00124506 |
| ENSMUSG00000030978 | Rrm1 | 1.68481205 | 6.75E-06 | 0.00124506 |
| ENSMUSG00000028435 | Aqp3 | -35.595636 | 6.80E-06 | 0.00124506 |
| ENSMUSG00000106379 | Lhfpl3 | -56.63899 | 6.81E-06 | 0.00124506 |
| ENSMUSG00000026786 | Apbb1ip | -1.8193253 | 7.08E-06 | 0.00128435 |
| ENSMUSG00000087598 | Zfp111 | 1.39866375 | 7.14E-06 | 0.00128645 |
| ENSMUSG00000002844 | Adprh | -1.5260268 | 7.85E-06 | 0.00139731 |
| ENSMUSG00000015947 | Fcgr1 | -2.2093673 | 7.92E-06 | 0.00139731 |
| ENSMUSG00000034023 | Fancd2 | 1.92884368 | 7.92E-06 | 0.00139731 |
| ENSMUSG00000040084 | Bub1b | 2.20684678 | 8.02E-06 | 0.00140137 |
| ENSMUSG00000032591 | Mst1 | 20.6081991 | 8.05E-06 | 0.00140137 |
| ENSMUSG00000079057 | Cyp4v3 | -1.8906064 | 9.27E-06 | 0.00159961 |
| ENSMUSG00000097328 | Tnfsf12 | -1.7075687 | 9.32E-06 | 0.00159961 |
| ENSMUSG00000026405 | C4bp | 550937408 | 9.54E-06 | 0.00162161 |
| ENSMUSG00000054115 | Skp2 | 2.23111309 | 9.57E-06 | 0.00162161 |
| ENSMUSG00000039748 | Exo1 | 2.47988562 | 9.71E-06 | 0.00162896 |
| ENSMUSG00000040026 | Saa3 | -12.74113 | 9.78E-06 | 0.00162896 |
| ENSMUSG00000052688 | Rab7b | -1.5139922 | 9.81E-06 | 0.00162896 |
| ENSMUSG00000070713 | Gm10282 | 1.69204839 | 1.01E-05 | 0.00166894 |
| ENSMUSG00000020682 | Mmp28 | -3.1854243 | 1.07E-05 | 0.00176078 |
| ENSMUSG00000029716 | Tfr2 | -5.6203722 | 1.09E-05 | 0.00177165 |
| ENSMUSG00000024891 | Slc29a2 | 2.7319349 | 1.14E-05 | 0.00183826 |
| ENSMUSG00000059089 | Fcgr4 | -3.2546535 | 1.15E-05 | 0.00185722 |
| ENSMUSG00000007891 | Ctsd | -2.0962627 | 1.18E-05 | 0.00188978 |
| ENSMUSG00000019564 | Arid3a | -1.9664238 | 1.24E-05 | 0.0019683 |
| ENSMUSG00000034107 | Ano7 | 5.93745813 | 1.29E-05 | 0.00202368 |
| ENSMUSG00000036560 | Lgi4 | -15.323335 | 1.29E-05 | 0.00202368 |
| ENSMUSG00000024795 | Kif20b | 2.16728346 | 1.31E-05 | 0.00203318 |
| ENSMUSG00000079186 | Gzmc | -8.5336727 | 1.31E-05 | 0.00203318 |
| ENSMUSG00000033350 | Chst2 | -2.926537 | 1.32E-05 | 0.00203318 |
| ENSMUSG00000054196 | Cthrc1 | -3.0003548 | 1.35E-05 | 0.00206281 |
| ENSMUSG00000024644 | Cndp2 | -1.842432 | 1.37E-05 | 0.00206934 |
| ENSMUSG00000026204 | Ptprn | -7.2099935 | 1.37E-05 | 0.00206934 |
| ENSMUSG00000043613 | Mmp3 | -7.2169107 | 1.40E-05 | 0.00211093 |
| ENSMUSG00000028599 | Tnfrsf1b | -1.8448114 | 1.47E-05 | 0.00219796 |
| ENSMUSG00000029019 | Nppb | -60.326459 | 1.48E-05 | 0.0022026 |
| ENSMUSG00000026605 | Cenpf | 2.19491887 | 1.55E-05 | 0.0022838 |
| ENSMUSG00000030678 | Maz | 1.34733939 | 1.60E-05 | 0.00234845 |
| ENSMUSG00000002835 | Chaf1a | 1.79407548 | 1.61E-05 | 0.00234845 |
| ENSMUSG00000048012 | Zfp473 | 3.15427886 | 1.69E-05 | 0.00246125 |
| ENSMUSG00000031375 | Bgn | -2.7570174 | 1.78E-05 | 0.00257488 |
| ENSMUSG00000000028 | Cdc45 | 2.15273658 | 1.83E-05 | 0.00262059 |
| ENSMUSG00000021298 | Gpr132 | -2.8982321 | 1.84E-05 | 0.00262139 |

| | | | | |
|---|---|---|---|---|
| ENSMUSG00000027737 | Slc7a11 | -10.827931 | 1.89E-05 | 0.00267086 |
| ENSMUSG00000036298 | Slc2a13 | -4.4571247 | 1.89E-05 | 0.00267086 |
| ENSMUSG00000003882 | Il7r | -3.9841335 | 1.91E-05 | 0.00268581 |
| ENSMUSG00000048583 | Igf2 | -21.371988 | 1.94E-05 | 0.00271065 |
| ENSMUSG00000027323 | Rad51 | 1.94896717 | 1.96E-05 | 0.0027161 |
| ENSMUSG00000026196 | Bard1 | 2.02701283 | 1.97E-05 | 0.00271614 |
| ENSMUSG00000025758 | Plk4 | 2.12144477 | 1.98E-05 | 0.00271614 |
| ENSMUSG00000022322 | Shcbp1 | 2.17115756 | 2.06E-05 | 0.00281345 |
| ENSMUSG00000038736 | Nudcd1 | 1.45822556 | 2.08E-05 | 0.00282563 |
| ENSMUSG00000059323 | Tonsl | 1.98047432 | 2.13E-05 | 0.00288269 |
| ENSMUSG00000025473 | Adam8 | -5.6951037 | 2.15E-05 | 0.00288863 |
| ENSMUSG00000087166 | L1td1 | 240545187 | 2.19E-05 | 0.00291283 |
| ENSMUSG00000031070 | Mrgprf | -9.1535312 | 2.20E-05 | 0.00291283 |
| ENSMUSG00000046259 | Sprr2h | -239161730 | 2.20E-05 | 0.00291283 |
| ENSMUSG00000027533 | Fabp5 | -4.3005565 | 2.24E-05 | 0.00293205 |
| ENSMUSG00000092624 | Gm3654 | 145.434758 | 2.25E-05 | 0.00293205 |
| ENSMUSG00000105703 | Gm43305 | 4.30364539 | 2.25E-05 | 0.00293205 |
| ENSMUSG00000048109 | Rbm15 | 1.58452753 | 2.30E-05 | 0.00298803 |
| ENSMUSG00000029254 | Stap1 | -2.8939436 | 2.33E-05 | 0.0030045 |
| ENSMUSG00000032271 | Nnmt | -3.7237435 | 2.38E-05 | 0.00305315 |
| ENSMUSG00000037362 | Ccn3 | -4.9964369 | 2.40E-05 | 0.00305315 |
| ENSMUSG00000078131 | Krtap1-3 | -217643575 | 2.41E-05 | 0.00305315 |
| ENSMUSG00000021190 | Lgmn | -1.9733431 | 2.42E-05 | 0.00305315 |
| ENSMUSG00000025479 | Cyp2e1 | -17.684386 | 2.43E-05 | 0.00305315 |
| ENSMUSG00000030867 | Plk1 | 2.1563808 | 2.44E-05 | 0.00305315 |
| ENSMUSG00000063767 | S100a7a | -23.779945 | 2.49E-05 | 0.00310159 |
| ENSMUSG00000045328 | Cenpe | 2.14575902 | 2.52E-05 | 0.0031228 |
| ENSMUSG00000044783 | Hjurp | 1.70966142 | 2.54E-05 | 0.00313107 |
| ENSMUSG00000057406 | Nsd2 | 1.65639001 | 2.56E-05 | 0.00313681 |
| ENSMUSG00000028364 | Tnc | -3.8308424 | 2.59E-05 | 0.00314363 |
| ENSMUSG00000032586 | Traip | 2.1210642 | 2.60E-05 | 0.00314363 |
| ENSMUSG00000023903 | Mmp25 | -2.6345945 | 2.60E-05 | 0.00314363 |
| ENSMUSG00000022426 | Josd1 | 1.40253793 | 2.63E-05 | 0.00316196 |
| ENSMUSG00000026608 | Kctd3 | 1.35011771 | 2.64E-05 | 0.00316196 |
| ENSMUSG00000009281 | Rarres2 | -4.230567 | 2.67E-05 | 0.00318904 |
| ENSMUSG00000032322 | Pstpip1 | -2.6771724 | 2.70E-05 | 0.00320581 |
| ENSMUSG00000039206 | Daglb | -1.8019452 | 2.72E-05 | 0.00321382 |
| ENSMUSG00000024301 | Kifc5b | 2.13343501 | 2.77E-05 | 0.00325816 |
| ENSMUSG00000085843 | Kank4os | 3.27973241 | 2.83E-05 | 0.003314 |
| ENSMUSG00000034641 | Cd300ld | -3.8517573 | 2.89E-05 | 0.00336312 |
| ENSMUSG00000019773 | Fbxo5 | 2.00501658 | 3.00E-05 | 0.00347307 |
| ENSMUSG00000030515 | Tarsl2 | 1.83268932 | 3.01E-05 | 0.00347307 |
| ENSMUSG00000029304 | Spp1 | -11.851991 | 3.11E-05 | 0.00356088 |
| ENSMUSG00000026837 | Col5a1 | -2.9083785 | 3.11E-05 | 0.00356088 |
| ENSMUSG00000010122 | Slc47a1 | 6.96786368 | 3.12E-05 | 0.00356088 |

| | | | | |
|---|---|---|---|---|
| ENSMUSG00000032218 | Ccnb2 | 1.97973852 | 3.18E-05 | 0.00360632 |
| ENSMUSG00000027469 | Tpx2 | 2.04251115 | 3.20E-05 | 0.00361915 |
| ENSMUSG00000022055 | Nefl | -99.981817 | 3.36E-05 | 0.00378181 |
| ENSMUSG00000030798 | Cd37 | -2.3123066 | 3.39E-05 | 0.00379766 |
| ENSMUSG00000028933 | Xrcc2 | 1.52108724 | 3.57E-05 | 0.00397986 |
| ENSMUSG00000024008 | Cpne5 | -22.413128 | 3.65E-05 | 0.00403539 |
| ENSMUSG00000006435 | Neurl1a | -3.1929338 | 3.65E-05 | 0.00403539 |
| ENSMUSG00000051220 | Ercc6l | 2.04876964 | 3.74E-05 | 0.00411438 |
| ENSMUSG00000025232 | Hexa | -1.8552892 | 3.77E-05 | 0.00411438 |
| ENSMUSG00000050410 | Tcf19 | 1.73623704 | 3.77E-05 | 0.00411438 |
| ENSMUSG00000045109 | Krtap4-7 | -130261452 | 3.96E-05 | 0.00429027 |
| ENSMUSG00000021477 | Ctsl | -3.0740313 | 3.98E-05 | 0.00429027 |
| ENSMUSG00000064165 | Krt39 | -128947039 | 3.99E-05 | 0.00429027 |
| ENSMUSG00000098557 | Kctd12 | -2.0859319 | 4.00E-05 | 0.00429027 |
| ENSMUSG00000026177 | Slc11a1 | -2.9701472 | 4.05E-05 | 0.00431914 |
| ENSMUSG00000000290 | Itgb2 | -2.6912598 | 4.06E-05 | 0.00431914 |
| ENSMUSG00000025185 | Loxl4 | -3.9464751 | 4.10E-05 | 0.00434124 |
| ENSMUSG00000060594 | Layn | -3.7499493 | 4.13E-05 | 0.00434251 |
| ENSMUSG00000110628 | Gm37797 | -124312462 | 4.14E-05 | 0.00434251 |
| ENSMUSG00000003518 | Dusp3 | -1.6935709 | 4.20E-05 | 0.00438928 |
| ENSMUSG00000024056 | Ndc80 | 1.9982846 | 4.27E-05 | 0.00441331 |
| ENSMUSG00000097177 | 9330159M07 | 3.25891164 | 4.27E-05 | 0.00441331 |
| ENSMUSG00000039187 | Fanci | 2.0590707 | 4.30E-05 | 0.00441331 |
| ENSMUSG00000003032 | Klf4 | -3.7919634 | 4.30E-05 | 0.00441331 |
| ENSMUSG00000058725 | Gm11937 | -119104736 | 4.31E-05 | 0.00441331 |
| ENSMUSG00000063626 | Unc5d | -117548854 | 4.36E-05 | 0.00443841 |
| ENSMUSG00000030748 | Il4ra | -1.7540759 | 4.37E-05 | 0.00443841 |
| ENSMUSG00000051378 | Kif18b | 1.9273273 | 4.41E-05 | 0.00446627 |
| ENSMUSG00000075555 | Gm10863 | 2.74526302 | 4.46E-05 | 0.00449462 |
| ENSMUSG00000039518 | Cdsn | -10.615236 | 4.57E-05 | 0.00458964 |
| ENSMUSG00000029591 | Ung | 2.15952057 | 4.66E-05 | 0.00466314 |
| ENSMUSG00000024843 | Chka | 1.94360292 | 4.71E-05 | 0.00468578 |
| ENSMUSG00000015053 | Gata2 | -7.1446945 | 4.73E-05 | 0.00468578 |
| ENSMUSG00000028197 | Col24a1 | -24.200675 | 4.74E-05 | 0.00468578 |
| ENSMUSG00000056394 | Lig1 | 1.82032602 | 4.80E-05 | 0.00471502 |
| ENSMUSG00000046186 | Cd109 | -2.7641244 | 4.83E-05 | 0.00471502 |
| ENSMUSG00000036223 | Ska1 | 2.08609104 | 4.84E-05 | 0.00471502 |
| ENSMUSG00000048327 | Ckap2l | 1.99205409 | 4.84E-05 | 0.00471502 |
| ENSMUSG00000068078 | 2310034C09I | -104558183 | 4.87E-05 | 0.00472637 |
| ENSMUSG00000083161 | Gm11427 | -3.2946037 | 4.96E-05 | 0.00477232 |
| ENSMUSG00000001175 | Calm1 | -1.439161 | 4.97E-05 | 0.00477232 |
| ENSMUSG00000025877 | Hk3 | -3.8238874 | 5.01E-05 | 0.00477232 |
| ENSMUSG00000040552 | C3ar1 | -2.6928281 | 5.02E-05 | 0.00477232 |
| ENSMUSG00000035403 | Crb2 | 23.7705272 | 5.02E-05 | 0.00477232 |
| ENSMUSG00000041431 | Ccnb1 | 2.13857248 | 5.03E-05 | 0.00477232 |

| | | | | |
|---|---|---|---|---|
| ENSMUSG00000038463 | Olfml2b | -3.8216167 | 5.15E-05 | 0.00483603 |
| ENSMUSG00000078262 | Krtap4-9 | -98091636 | 5.17E-05 | 0.00483603 |
| ENSMUSG00000020658 | Efr3b | -2.7937618 | 5.21E-05 | 0.00483603 |
| ENSMUSG00000055095 | Spink6 | -97368665 | 5.21E-05 | 0.00483603 |
| ENSMUSG00000071858 | Gm94 | -97432949 | 5.21E-05 | 0.00483603 |
| ENSMUSG00000043740 | B430306N03 | -5.3098759 | 5.23E-05 | 0.00483603 |
| ENSMUSG00000021403 | Serpinb9b | -5.9226361 | 5.23E-05 | 0.00483603 |
| ENSMUSG00000020077 | Srgn | -2.3114892 | 5.26E-05 | 0.00484018 |
| ENSMUSG00000020354 | Sgcd | -4.2964293 | 5.29E-05 | 0.00485328 |
| ENSMUSG00000021822 | Plau | -2.1070876 | 5.41E-05 | 0.00494603 |
| ENSMUSG00000000440 | Pparg | -3.3881402 | 5.50E-05 | 0.00501236 |
| ENSMUSG00000026728 | Vim | -2.6211135 | 5.62E-05 | 0.00507828 |
| ENSMUSG00000069114 | Zbtb10 | 1.66938502 | 5.62E-05 | 0.00507828 |
| ENSMUSG00000075567 | Krtap1-4 | -89570581 | 5.64E-05 | 0.00507828 |
| ENSMUSG00000050075 | Gpr171 | -3.2239011 | 5.72E-05 | 0.00513252 |
| ENSMUSG00000031379 | Pir | -5.7955394 | 5.74E-05 | 0.00513683 |
| ENSMUSG00000026683 | Nuf2 | 1.86417242 | 5.82E-05 | 0.00519117 |
| ENSMUSG00000085773 | Gm16233 | 84942980.2 | 5.92E-05 | 0.00525988 |
| ENSMUSG00000023885 | Thbs2 | -3.6266424 | 5.94E-05 | 0.00525988 |
| ENSMUSG00000032802 | Srxn1 | -2.4652182 | 6.03E-05 | 0.00532144 |
| ENSMUSG00000012282 | Wnt8a | 142.394775 | 6.07E-05 | 0.00533325 |
| ENSMUSG00000007080 | Pole | 1.9041939 | 6.27E-05 | 0.00549185 |
| ENSMUSG00000029175 | Slc35f6 | -1.4693332 | 6.29E-05 | 0.00549185 |
| ENSMUSG00000030393 | Zik1 | 2.13048486 | 6.37E-05 | 0.00554278 |
| ENSMUSG00000030677 | Kif22 | 2.08358086 | 6.55E-05 | 0.00568092 |
| ENSMUSG00000079227 | Ccr5 | -2.0726159 | 6.65E-05 | 0.00574718 |
| ENSMUSG00000050224 | Krtap13 | -73728049 | 6.76E-05 | 0.00581935 |
| ENSMUSG00000028581 | Laptm5 | -1.8506572 | 6.83E-05 | 0.00586695 |
| ENSMUSG00000021457 | Syk | -1.9376078 | 6.87E-05 | 0.00587403 |
| ENSMUSG00000056758 | Hmga2 | -36.537728 | 6.89E-05 | 0.00587587 |
| ENSMUSG00000050069 | Grem2 | -11.954875 | 6.94E-05 | 0.00589602 |
| ENSMUSG00000032400 | Zwilch | 1.6351206 | 6.96E-05 | 0.00589865 |
| ENSMUSG00000062433 | Krtap6-2 | -70974170 | 7.00E-05 | 0.00590941 |
| ENSMUSG00000001228 | Uhrf1 | 2.08562016 | 7.07E-05 | 0.00595162 |
| ENSMUSG00000002603 | Tgfb1 | -1.6737453 | 7.10E-05 | 0.00595592 |
| ENSMUSG00000021148 | Gm9745 | -68956386 | 7.19E-05 | 0.00600967 |
| ENSMUSG00000024989 | Cep55 | 1.99481408 | 7.25E-05 | 0.0060416 |
| ENSMUSG00000092312 | Zfp419 | 98.0196811 | 7.29E-05 | 0.00605293 |
| ENSMUSG00000036768 | Kif15 | 2.28219683 | 7.46E-05 | 0.00617185 |
| ENSMUSG00000039196 | Orm1 | -14.024399 | 7.55E-05 | 0.00622739 |
| ENSMUSG00000028278 | Rragd | 3.14425216 | 7.59E-05 | 0.00624145 |
| ENSMUSG00000052298 | Cdc42se2 | -1.2991853 | 7.61E-05 | 0.00624145 |
| ENSMUSG00000031657 | Heatr3 | 1.33244311 | 7.65E-05 | 0.00624801 |
| ENSMUSG00000002808 | Epdr1 | -1.8966506 | 7.69E-05 | 0.00626009 |
| ENSMUSG00000039126 | Prune2 | -2.4341521 | 7.88E-05 | 0.00638891 |

| | | | | |
|---|---|---|---|---|
| ENSMUSG00000074476 | Spc24 | 2.07914042 | 7.89E-05 | 0.00638891 |
| ENSMUSG00000031756 | Cenpn | 2.00598917 | 8.04E-05 | 0.00645344 |
| ENSMUSG00000067613 | Krt83 | -148.61836 | 8.04E-05 | 0.00645344 |
| ENSMUSG00000055799 | Tcf7l1 | 2.23118922 | 8.05E-05 | 0.00645344 |
| ENSMUSG00000031278 | Acsl4 | -1.8195383 | 8.12E-05 | 0.0064755 |
| ENSMUSG00000045331 | 2310079G19 | -60351707 | 8.13E-05 | 0.0064755 |
| ENSMUSG00000078771 | Evi2a | -1.5758895 | 8.19E-05 | 0.00650092 |
| ENSMUSG00000008153 | Clstn3 | 6.36574145 | 8.28E-05 | 0.00655374 |
| ENSMUSG00000043323 | Fbrsl1 | 1.46640335 | 8.33E-05 | 0.00657272 |
| ENSMUSG00000036777 | Anln | 1.83675748 | 8.36E-05 | 0.00657868 |
| ENSMUSG00000034022 | Cpsf1 | 1.31173052 | 8.44E-05 | 0.00660481 |
| ENSMUSG00000058368 | Krtap21-1 | -57879381 | 8.45E-05 | 0.00660481 |
| ENSMUSG00000040212 | Emp3 | -2.7013565 | 8.62E-05 | 0.00671805 |
| ENSMUSG00000029165 | Agbl5 | 1.53123795 | 8.67E-05 | 0.00673124 |
| ENSMUSG00000043263 | Ifi209 | -2.410106 | 8.69E-05 | 0.00673124 |
| ENSMUSG00000079037 | Prnp | -2.0018254 | 8.95E-05 | 0.00691496 |
| ENSMUSG00000028702 | Rad54l | 1.91608464 | 8.98E-05 | 0.00691741 |
| ENSMUSG00000029762 | Akr1b8 | -2.547414 | 9.05E-05 | 0.00694918 |
| ENSMUSG00000024165 | Jpt2 | 1.52927754 | 9.08E-05 | 0.00694969 |
| ENSMUSG00000020808 | Pimreg | 1.95672489 | 9.11E-05 | 0.00695375 |
| ENSMUSG00000016194 | Hsd11b1 | -3.0509634 | 9.15E-05 | 0.00696363 |
| ENSMUSG00000071068 | Treml2 | -3.5371099 | 9.44E-05 | 0.00714221 |
| ENSMUSG00000048830 | 2310057N15 | -51095811 | 9.47E-05 | 0.00714221 |
| ENSMUSG00000046223 | Plaur | -3.3063458 | 9.47E-05 | 0.00714221 |
| ENSMUSG00000004207 | Psap | -1.3873128 | 9.59E-05 | 0.00721127 |
| ENSMUSG00000021678 | F2rl1 | 3.02827064 | 9.62E-05 | 0.00721229 |
| ENSMUSG00000015396 | Cd83 | -2.4112659 | 9.81E-05 | 0.00732013 |
| ENSMUSG00000060703 | Cd302 | -1.9085585 | 9.82E-05 | 0.00732013 |
| ENSMUSG00000113136 | Gm19951 | -12.899292 | 9.90E-05 | 0.00735225 |
| ENSMUSG00000030142 | Clec4e | -9.9549208 | 9.92E-05 | 0.00735225 |
| ENSMUSG00000072258 | Taf1a | 1.59563711 | 0.00010101 | 0.00746167 |
| ENSMUSG00000028312 | Smc2 | 1.80028329 | 0.00010137 | 0.0074669 |
| ENSMUSG00000038644 | Pold1 | 1.75572174 | 0.00010209 | 0.0074978 |
| ENSMUSG00000028832 | Stmn1 | 2.2877725 | 0.00010314 | 0.00755312 |
| ENSMUSG00000056832 | Ttc26 | 1.75880003 | 0.00010519 | 0.00766777 |
| ENSMUSG00000053205 | Styx | 1.68397601 | 0.00010531 | 0.00766777 |
| ENSMUSG00000017760 | Ctsa | -1.5773016 | 0.00010665 | 0.00774368 |
| ENSMUSG00000054203 | Ifi205 | -2.9387197 | 0.00011106 | 0.00804097 |
| ENSMUSG00000028031 | Dkk2 | -4.6884888 | 0.0001116 | 0.00805671 |
| ENSMUSG00000019988 | Nedd1 | 1.56825411 | 0.00011234 | 0.00808166 |
| ENSMUSG00000032997 | Chpf | -1.8365979 | 0.00011258 | 0.00808166 |
| ENSMUSG00000019796 | Lrp11 | -2.0945607 | 0.0001139 | 0.00815311 |
| ENSMUSG00000022483 | Col2a1 | -7.6553747 | 0.00011431 | 0.00815972 |
| ENSMUSG00000020914 | Top2a | 1.80282843 | 0.0001149 | 0.0081787 |
| ENSMUSG00000032783 | Troap | 2.1213556 | 0.00011567 | 0.00819413 |

| | | | | |
|---|---|---|---|---|
| ENSMUSG00000058818 | Pirb | -2.3337411 | 0.00011576 | 0.00819413 |
| ENSMUSG00000044548 | Dact1 | -4.4653307 | 0.00011619 | 0.00820149 |
| ENSMUSG00000022180 | Slc7a8 | -2.3629986 | 0.00011703 | 0.00823848 |
| ENSMUSG00000030144 | Clec4d | -8.1247126 | 0.00011827 | 0.00830263 |
| ENSMUSG00000038523 | 1700003F12I | 8.34967011 | 0.00011963 | 0.00837469 |
| ENSMUSG00000048163 | Selplg | -1.8497771 | 0.00012043 | 0.0084076 |
| ENSMUSG00000030223 | Ptpro | -2.0823742 | 0.00012154 | 0.00846174 |
| ENSMUSG00000030726 | Pold3 | 1.43660695 | 0.00012276 | 0.00852355 |
| ENSMUSG00000041498 | Kif14 | 2.12566348 | 0.0001234 | 0.00854441 |
| ENSMUSG00000047415 | Gpr68 | -2.5738702 | 0.00012447 | 0.00859522 |
| ENSMUSG00000007029 | Vars | 1.45956994 | 0.00012593 | 0.00865855 |
| ENSMUSG00000034311 | Kif4 | 2.22897517 | 0.00012625 | 0.00865855 |
| ENSMUSG00000014592 | Camta1 | 2.47022259 | 0.00012668 | 0.00865855 |
| ENSMUSG00000048922 | Cdca2 | 2.18584008 | 0.00012704 | 0.00865855 |
| ENSMUSG00000025089 | Gfra1 | -5.386082 | 0.00012709 | 0.00865855 |
| ENSMUSG00000030653 | Gm45837 | 8939143.54 | 0.00012764 | 0.00867297 |
| ENSMUSG00000009047 | Gm5965 | -36385036 | 0.00012871 | 0.008722 |
| ENSMUSG00000075266 | Cenpw | 2.04128727 | 0.00013004 | 0.00878851 |
| ENSMUSG00000028238 | Atp6v0d2 | -13.107636 | 0.00013091 | 0.00882377 |
| ENSMUSG00000078763 | Slfn1 | -2.4687021 | 0.0001315 | 0.00884038 |
| ENSMUSG00000038342 | Mlxip | 1.5539422 | 0.00013337 | 0.00893728 |
| ENSMUSG00000024885 | Aldh3b1 | -1.7286878 | 0.00013374 | 0.00893728 |
| ENSMUSG00000029413 | Naaa | -1.8162649 | 0.00013406 | 0.00893728 |
| ENSMUSG00000036523 | Greb1 | -3.472052 | 0.00013436 | 0.00893728 |
| ENSMUSG00000031367 | Ap1s2 | -1.7135647 | 0.0001347 | 0.00893728 |
| ENSMUSG00000017765 | Slc12a4 | -1.8050941 | 0.00013515 | 0.00894408 |
| ENSMUSG00000026390 | Marco | -40.208005 | 0.00013638 | 0.00900153 |
| ENSMUSG00000063193 | Cd300lb | -5.2505279 | 0.00013814 | 0.00909464 |
| ENSMUSG00000033220 | Rac2 | -2.0396032 | 0.00013867 | 0.00910542 |
| ENSMUSG00000024990 | Rbp4 | -5.3400411 | 0.00013902 | 0.00910542 |
| ENSMUSG00000047786 | Lix1 | -3.7235546 | 0.00014054 | 0.00918111 |
| ENSMUSG00000021504 | B4galt7 | -1.4585704 | 0.00014162 | 0.00922805 |
| ENSMUSG00000025980 | Hspd1 | 1.63713325 | 0.00014325 | 0.00930926 |
| ENSMUSG00000020330 | Hmmr | 2.04063885 | 0.00014393 | 0.00930926 |
| ENSMUSG00000079625 | Tm4sf19 | -6.9092524 | 0.00014469 | 0.00930926 |
| ENSMUSG00000035697 | Arhgap45 | -1.6806814 | 0.00014473 | 0.00930926 |
| ENSMUSG00000037613 | Tnfrsf23 | -2.8912476 | 0.00014505 | 0.00930926 |
| ENSMUSG00000011958 | Bnip2 | -1.2467781 | 0.00014509 | 0.00930926 |
| ENSMUSG00000031903 | Pla2g15 | -1.772663 | 0.00014569 | 0.00930926 |
| ENSMUSG00000043832 | Clec4a3 | -2.4994861 | 0.0001458 | 0.00930926 |
| ENSMUSG00000024232 | Bambi | -5.9833852 | 0.00014665 | 0.00934018 |
| ENSMUSG00000058624 | Gda | -2.3116358 | 0.00015048 | 0.00954115 |
| ENSMUSG00000068075 | Gm10229 | -30491295 | 0.00015056 | 0.00954115 |
| ENSMUSG00000026737 | Pip4k2a | -1.5562333 | 0.00015303 | 0.00967351 |
| ENSMUSG00000030101 | Sumf1 | -1.4901508 | 0.00015449 | 0.00974156 |

| | | | | |
|---|---|---|---|---|
| ENSMUSG00000027379 | Bub1 | 1.89246734 | 0.00015804 | 0.00994065 |
| ENSMUSG00000026622 | Nek2 | 2.09933666 | 0.00016136 | 0.01010941 |
| ENSMUSG00000019992 | Mtfr2 | 1.9055637 | 0.00016152 | 0.01010941 |
| ENSMUSG00000034206 | Polq | 2.19322737 | 0.00016305 | 0.0101804 |
| ENSMUSG00000025383 | Il23a | 6.44932631 | 0.00016526 | 0.01028051 |
| ENSMUSG00000062510 | Nsl1 | 1.71210322 | 0.00016546 | 0.01028051 |
| ENSMUSG00000028439 | Fam219a | -1.6044254 | 0.00016647 | 0.01028756 |
| ENSMUSG00000021391 | Cenpp | 2.32804843 | 0.00016652 | 0.01028756 |
| ENSMUSG00000030541 | Idh2 | 1.51852724 | 0.00016679 | 0.01028756 |
| ENSMUSG00000031629 | Cenpu | 1.84491651 | 0.00016775 | 0.01032196 |
| ENSMUSG00000027490 | E2f1 | 1.97590708 | 0.0001727 | 0.01057991 |
| ENSMUSG00000004707 | Ly9 | -1.9182278 | 0.00017304 | 0.01057991 |
| ENSMUSG00000057174 | Krtap19-9b | -26036087 | 0.0001732 | 0.01057991 |
| ENSMUSG00000053219 | Raet1e | 4.33145325 | 0.00017515 | 0.01064921 |
| ENSMUSG00000022586 | Ly6i | 10.0926949 | 0.00017548 | 0.01064921 |
| ENSMUSG00000062168 | Ppef1 | 158.55507 | 0.00017567 | 0.01064921 |
| ENSMUSG00000039058 | Ak5 | -5.963666 | 0.00017631 | 0.01064921 |
| ENSMUSG00000031513 | Leprotl1 | -1.4122815 | 0.00017648 | 0.01064921 |
| ENSMUSG00000019942 | Cdk1 | 1.97322687 | 0.00017692 | 0.01064921 |
| ENSMUSG00000039330 | Tsga10ip | 231.07422 | 0.00017726 | 0.01064921 |
| ENSMUSG00000051748 | Wfdc21 | -8.9315152 | 0.0001785 | 0.01068902 |
| ENSMUSG00000085766 | 2810430I11R | -7.6880772 | 0.00017877 | 0.01068902 |
| ENSMUSG00000076617 | Ighm | -10.044896 | 0.00017979 | 0.0107252 |
| ENSMUSG00000047104 | Pbp2 | -20.956408 | 0.00018117 | 0.01078218 |
| ENSMUSG00000079629 | Rhox2g | 33.4237815 | 0.00018199 | 0.01080552 |
| ENSMUSG00000021944 | Gata4 | 37.7209845 | 0.00018262 | 0.01081748 |
| ENSMUSG00000074447 | Defa21 | -23459595 | 0.0001898 | 0.0112026 |
| ENSMUSG00000020897 | Aurkb | 1.87322401 | 0.00019 | 0.0112026 |
| ENSMUSG00000023991 | Foxp4 | 1.2978622 | 0.00019128 | 0.0112518 |
| ENSMUSG00000090515 | Krtap27-1 | -23143213 | 0.00019204 | 0.01127067 |
| ENSMUSG00000045326 | Fndc7 | 25.3105853 | 0.0001933 | 0.01131833 |
| ENSMUSG00000097910 | 5033428I22R | -88.931168 | 0.00019388 | 0.01132639 |
| ENSMUSG00000035891 | Cerk | -2.0992121 | 0.00019736 | 0.01150307 |
| ENSMUSG00000005506 | Celf1 | 1.30777931 | 0.0001983 | 0.01153141 |
| ENSMUSG00000028423 | Nfx1 | 1.29067002 | 0.00019986 | 0.0115953 |
| ENSMUSG00000029307 | Dmp1 | -16.759372 | 0.00020227 | 0.01170879 |
| ENSMUSG00000056481 | Cd248 | -2.2429534 | 0.00020406 | 0.01178516 |
| ENSMUSG00000029217 | Tec | -1.625709 | 0.00020568 | 0.01184808 |
| ENSMUSG00000036273 | Lrrk2 | -2.0887618 | 0.00020608 | 0.01184808 |
| ENSMUSG00000040972 | Igsf21 | -94.951928 | 0.00020698 | 0.0118535 |
| ENSMUSG00000085148 | Mir22hg | -1.8451812 | 0.00020722 | 0.0118535 |
| ENSMUSG00000036381 | P2ry14 | -2.7016603 | 0.00020757 | 0.0118535 |
| ENSMUSG00000041219 | Arhgap11a | 1.69251397 | 0.00020952 | 0.0119137 |
| ENSMUSG00000026790 | Odf2 | 1.33591539 | 0.00020956 | 0.0119137 |
| ENSMUSG00000103440 | Gm37131 | 31.4011061 | 0.00021553 | 0.01222565 |

| | | | | |
|---|---|---|---|---|
| ENSMUSG00000037217 | Syn1 | -3.065891 | 0.00021654 | 0.01225556 |
| ENSMUSG00000042734 | Ttc9 | -3.777754 | 0.000218 | 0.01228737 |
| ENSMUSG00000053477 | Tcf4 | -2.2562159 | 0.00021807 | 0.01228737 |
| ENSMUSG00000022033 | Pbk | 2.11017962 | 0.00021982 | 0.01233234 |
| ENSMUSG00000004952 | Rasa4 | -1.5841649 | 0.00021984 | 0.01233234 |
| ENSMUSG00000098112 | Bin2 | -1.6879705 | 0.00022052 | 0.01233553 |
| ENSMUSG00000031242 | 2610002M06 | 1.64804816 | 0.00022087 | 0.01233553 |
| ENSMUSG00000062593 | Gm49339 | -2.8692614 | 0.00022419 | 0.01247304 |
| ENSMUSG00000023004 | Tuba1b | 1.56149956 | 0.00022431 | 0.01247304 |
| ENSMUSG00000028048 | Gba | -1.4898592 | 0.00022631 | 0.01255678 |
| ENSMUSG00000040725 | Hnrnpul1 | 1.36606774 | 0.00022717 | 0.01256421 |
| ENSMUSG00000019838 | Slc16a10 | -3.1030129 | 0.00022743 | 0.01256421 |
| ENSMUSG00000042745 | Id1 | -4.4261471 | 0.00022887 | 0.01261626 |
| ENSMUSG00000046591 | Ticrr | 2.43728828 | 0.00023212 | 0.01276739 |
| ENSMUSG00000021242 | Npc2 | -1.5610291 | 0.00023287 | 0.01278117 |
| ENSMUSG00000079553 | Kifc1 | 1.73197445 | 0.00023428 | 0.01283072 |
| ENSMUSG00000010142 | Tnfrsf13b | -2.3384097 | 0.0002352 | 0.01283127 |
| ENSMUSG00000049265 | Kcnk3 | -12.229018 | 0.00023539 | 0.01283127 |
| ENSMUSG00000028885 | Smpdl3b | 1.88319733 | 0.00023626 | 0.01283127 |
| ENSMUSG00000002870 | Mcm2 | 1.62214425 | 0.00023631 | 0.01283127 |
| ENSMUSG00000014846 | Tppp3 | -2.8815679 | 0.00023892 | 0.01294391 |
| ENSMUSG00000026779 | Mastl | 1.83061486 | 0.0002394 | 0.01294391 |
| ENSMUSG00000029379 | Cxcl3 | -96.474849 | 0.00024009 | 0.01295347 |
| ENSMUSG00000026981 | Il1rn | -4.4576056 | 0.00024141 | 0.01299732 |
| ENSMUSG00000048450 | Msx1 | -3.2042578 | 0.00024313 | 0.01306233 |
| ENSMUSG00000002204 | Napsa | -2.4175242 | 0.00024797 | 0.01329405 |
| ENSMUSG00000026676 | Ccdc3 | -7.7745711 | 0.00025276 | 0.01352231 |
| ENSMUSG00000038740 | Mvb12b | -1.6665016 | 0.00025418 | 0.01356957 |
| ENSMUSG00000040653 | Ppp1r14c | -7.6408125 | 0.00025494 | 0.01358185 |
| ENSMUSG00000060459 | Kng2 | -33.971585 | 0.00025777 | 0.01369281 |
| ENSMUSG00000045362 | Tnfrsf26 | -5.402069 | 0.0002581 | 0.01369281 |
| ENSMUSG00000075707 | Dio3 | -9.1074103 | 0.00026041 | 0.01373657 |
| ENSMUSG00000022766 | Serpind1 | -12.554547 | 0.00026079 | 0.01373657 |
| ENSMUSG00000039217 | Il18 | -2.1617642 | 0.00026084 | 0.01373657 |
| ENSMUSG00000071714 | Csf2rb2 | -2.1328688 | 0.00026167 | 0.01373657 |
| ENSMUSG00000023781 | Hes7 | 6.65082345 | 0.00026179 | 0.01373657 |
| ENSMUSG00000030199 | Etv6 | 1.42352922 | 0.00026217 | 0.01373657 |
| ENSMUSG00000004880 | Lbr | 1.67651281 | 0.00026549 | 0.01388215 |
| ENSMUSG00000044221 | Grsf1 | 1.27286052 | 0.00026901 | 0.01403708 |
| ENSMUSG00000032122 | Slc37a2 | -2.9374007 | 0.00027103 | 0.01411341 |
| ENSMUSG00000019230 | Lhx9 | -41.504827 | 0.00027295 | 0.01418423 |
| ENSMUSG00000028256 | Odf2l | 1.58252795 | 0.00027376 | 0.01419759 |
| ENSMUSG00000031959 | Wdr59 | 1.42231948 | 0.00027773 | 0.01437215 |
| ENSMUSG00000017493 | Igfbp4 | -2.9950748 | 0.00027851 | 0.01437215 |
| ENSMUSG00000049037 | Clec4a1 | -2.7104879 | 0.00027945 | 0.01437215 |

| | | | | |
|---|---|---|---|---|
| ENSMUSG00000000374 | Trappc10 | 1.40318904 | 0.00027982 | 0.01437215 |
| ENSMUSG00000018819 | Lsp1 | -2.0551084 | 0.00027995 | 0.01437215 |
| ENSMUSG00000037313 | Tacc3 | 1.75059584 | 0.00028078 | 0.01438531 |
| ENSMUSG00000061780 | Cfd | -7.7745122 | 0.00028321 | 0.01448069 |
| ENSMUSG00000024095 | Hnrnpll | 1.26693046 | 0.00028624 | 0.01460606 |
| ENSMUSG00000025747 | Tyms | 1.55476033 | 0.00029081 | 0.01479546 |
| ENSMUSG00000031922 | Cep57 | 1.51494053 | 0.00029111 | 0.01479546 |
| ENSMUSG00000042759 | Apobr | 3.29047738 | 0.00029289 | 0.01485597 |
| ENSMUSG00000028873 | Cdca8 | 2.01339547 | 0.00029775 | 0.0150725 |
| ENSMUSG00000071713 | Csf2rb | -2.2851405 | 0.00030296 | 0.01530583 |
| ENSMUSG00000040249 | Lrp1 | -2.1545428 | 0.00030806 | 0.01553246 |
| ENSMUSG00000072082 | Ccnf | 1.75715083 | 0.00030993 | 0.01559397 |
| ENSMUSG00000056054 | S100a8 | -13.791188 | 0.0003105 | 0.01559397 |
| ENSMUSG00000030786 | Itgam | -3.1773693 | 0.0003116 | 0.01561786 |
| ENSMUSG00000028809 | Srrm1 | 1.36303683 | 0.00031553 | 0.01569783 |
| ENSMUSG00000020340 | Cyfip2 | -2.5333616 | 0.00031579 | 0.01569783 |
| ENSMUSG00000097203 | 4732419C18I | 16.5670967 | 0.00031583 | 0.01569783 |
| ENSMUSG00000018548 | Trim37 | 1.54638277 | 0.00031619 | 0.01569783 |
| ENSMUSG00000038271 | Iffo1 | -1.9934111 | 0.00031628 | 0.01569783 |
| ENSMUSG00000096965 | 3300005D01 | -4.7211991 | 0.00031976 | 0.01583959 |
| ENSMUSG00000006724 | Cyp27b1 | 4.91926923 | 0.00032474 | 0.01605512 |
| ENSMUSG00000028718 | Stil | 1.97519248 | 0.00032562 | 0.01606715 |
| ENSMUSG00000027367 | Stard7 | 1.3264131 | 0.00032722 | 0.01611506 |
| ENSMUSG00000018008 | Cyth4 | -1.7267743 | 0.00033283 | 0.01635748 |
| ENSMUSG00000090946 | Ccdc71l | -1.9986236 | 0.00033343 | 0.01635748 |
| ENSMUSG00000054717 | Hmgb2 | 1.93482382 | 0.00034211 | 0.0167508 |
| ENSMUSG00000028896 | Rcc1 | 1.66689081 | 0.00034308 | 0.01676182 |
| ENSMUSG00000039462 | Col10a1 | -16.508477 | 0.00034413 | 0.01676182 |
| ENSMUSG00000020288 | Ahsa2 | -1.3455078 | 0.00034462 | 0.01676182 |
| ENSMUSG00000015112 | Slc25a13 | 1.44220814 | 0.00034506 | 0.01676182 |
| ENSMUSG00000021732 | Fgf10 | -8.9780505 | 0.00034563 | 0.01676182 |
| ENSMUSG00000016529 | Il10 | -6.6363231 | 0.00034633 | 0.01676384 |
| ENSMUSG00000029516 | Cit | 2.15960027 | 0.00034755 | 0.01676918 |
| ENSMUSG00000036913 | Trim67 | -44.617391 | 0.00034781 | 0.01676918 |
| ENSMUSG00000056267 | Cep70 | 1.70317531 | 0.00034842 | 0.01676918 |
| ENSMUSG00000003038 | Hmgn2 | 1.52753816 | 0.00034927 | 0.01677827 |
| ENSMUSG00000037474 | Dtl | 1.79662892 | 0.00035208 | 0.01688134 |
| ENSMUSG00000028681 | Ptch2 | -6.9685283 | 0.00035507 | 0.016979 |
| ENSMUSG00000069793 | Slfn9 | 1.84011121 | 0.00035546 | 0.016979 |
| ENSMUSG00000062432 | Cyp26c1 | -39.852533 | 0.0003603 | 0.01717821 |
| ENSMUSG00000006678 | Pola1 | 1.7961246 | 0.00036242 | 0.01724687 |
| ENSMUSG00000022021 | Diaph3 | 2.01814999 | 0.0003647 | 0.01731569 |
| ENSMUSG00000073830 | Mup14 | -26.192049 | 0.00036523 | 0.01731569 |
| ENSMUSG00000037315 | Jade3 | 1.86310964 | 0.00037205 | 0.01760607 |
| ENSMUSG00000032498 | Mlh1 | 1.50002409 | 0.00037296 | 0.0176165 |

| | | | | |
|---|---|---|---|---|
| ENSMUSG00000033016 | Nfatc1 | -1.4759422 | 0.00037414 | 0.01763951 |
| ENSMUSG00000112023 | Lilr4b | -3.6761274 | 0.00037867 | 0.01782008 |
| ENSMUSG00000028108 | Ecm1 | -2.9368299 | 0.00037953 | 0.01782751 |
| ENSMUSG00000028339 | Col15a1 | -3.3839489 | 0.00038099 | 0.01785103 |
| ENSMUSG00000029860 | Zyx | -1.5204516 | 0.00038144 | 0.01785103 |
| ENSMUSG00000040528 | Milr1 | -1.9500686 | 0.00038529 | 0.01799799 |
| ENSMUSG00000037544 | Dlgap5 | 2.02362353 | 0.00039053 | 0.0181927 |
| ENSMUSG00000019961 | Tmpo | 1.59774041 | 0.00039104 | 0.0181927 |
| ENSMUSG00000021697 | Depdc1b | 2.20782796 | 0.0003916 | 0.0181927 |
| ENSMUSG00000027513 | Pck1 | -8.0339453 | 0.00040025 | 0.01851209 |
| ENSMUSG00000020143 | Dock2 | -1.8075829 | 0.00040182 | 0.01851209 |
| ENSMUSG00000038259 | Gdf5 | 6.37375811 | 0.00040183 | 0.01851209 |
| ENSMUSG00000021194 | Chga | -10.06277 | 0.00040197 | 0.01851209 |
| ENSMUSG00000041911 | Dlx1 | -23.83454 | 0.00040247 | 0.01851209 |
| ENSMUSG00000026463 | Atp2b4 | -2.169796 | 0.00040285 | 0.01851209 |
| ENSMUSG00000034591 | Slc41a2 | -1.9266078 | 0.00041171 | 0.01888502 |
| ENSMUSG00000012705 | Retn | -9.2596816 | 0.00041412 | 0.0189615 |
| ENSMUSG00000030089 | Slc41a3 | -1.7550108 | 0.000418 | 0.01908669 |
| ENSMUSG00000059923 | Grb2 | -1.2961628 | 0.00041836 | 0.01908669 |
| ENSMUSG00000044676 | Zfp612 | 1.51731438 | 0.00042025 | 0.01911338 |
| ENSMUSG00000037579 | Kcnh3 | 9.07730812 | 0.00042045 | 0.01911338 |
| ENSMUSG00000020120 | Plek | -2.1751327 | 0.00042156 | 0.01911899 |
| ENSMUSG00000040713 | Creg1 | -2.0690505 | 0.00042207 | 0.01911899 |
| ENSMUSG00000046179 | E2f8 | 1.86358384 | 0.00042571 | 0.01924937 |
| ENSMUSG00000024679 | Ms4a6d | -2.2052984 | 0.00042749 | 0.01928783 |
| ENSMUSG00000017466 | Timp2 | -1.4869116 | 0.00042808 | 0.01928783 |
| ENSMUSG00000036882 | Arhgap33 | 2.84785779 | 0.0004291 | 0.01929951 |
| ENSMUSG00000030546 | Plin1 | -7.0131311 | 0.00043213 | 0.01939999 |
| ENSMUSG00000024486 | Hbegf | -3.438618 | 0.00043357 | 0.01939999 |
| ENSMUSG00000030752 | Kdm8 | 1.62168422 | 0.00043362 | 0.01939999 |
| ENSMUSG00000027959 | Sass6 | 1.47764054 | 0.00043591 | 0.01946787 |
| ENSMUSG00000041147 | Brca2 | 1.67234273 | 0.00043727 | 0.01948435 |
| ENSMUSG00000034707 | Gns | -1.4411105 | 0.00043781 | 0.01948435 |
| ENSMUSG00000118171 | Gm50390 | -61.087144 | 0.00043968 | 0.01953354 |
| ENSMUSG00000022878 | Adipoq | -6.8402364 | 0.00044145 | 0.01956306 |
| ENSMUSG00000027635 | Dsn1 | 1.80978542 | 0.00044189 | 0.01956306 |
| ENSMUSG00000030254 | Rad18 | 1.83875394 | 0.00045085 | 0.01992531 |
| ENSMUSG00000021868 | Ppif | 1.66206136 | 0.00045492 | 0.0200701 |
| ENSMUSG00000005102 | Eif2ak4 | 1.45196422 | 0.00045685 | 0.02010502 |
| ENSMUSG00000046295 | Ankle1 | 3.27549449 | 0.00045729 | 0.02010502 |
| ENSMUSG00000022831 | Hcls1 | -1.7568812 | 0.00045809 | 0.02010522 |
| ENSMUSG00000036770 | Stpg3 | -53.090871 | 0.00046612 | 0.02042265 |
| ENSMUSG00000034612 | Chst11 | -2.2622847 | 0.00046699 | 0.02042537 |
| ENSMUSG00000040751 | Lat2 | -2.105612 | 0.00046885 | 0.02047151 |
| ENSMUSG00000032698 | Lmo2 | -2.2609682 | 0.0004728 | 0.02060837 |

| | | | | |
|---|---|---|---|---|
| ENSMUSG00000059811 | Atl2 | 1.26411113 | 0.00047525 | 0.02067973 |
| ENSMUSG00000042821 | Snai1 | -2.9847886 | 0.00047642 | 0.02069548 |
| ENSMUSG00000005410 | Mcm5 | 1.59968069 | 0.0004793 | 0.02078481 |
| ENSMUSG00000024535 | Snx24 | -1.788987 | 0.00048188 | 0.02086131 |
| ENSMUSG00000018774 | Cd68 | -2.4046083 | 0.00048545 | 0.02097992 |
| ENSMUSG00000035373 | Ccl7 | -2.9675556 | 0.00048675 | 0.02099589 |
| ENSMUSG00000027809 | Etfdh | 1.36945107 | 0.00048747 | 0.02099589 |
| ENSMUSG00000007946 | Phox2a | 4.28994223 | 0.00048872 | 0.02101426 |
| ENSMUSG00000027160 | Ccdc34 | 1.37226285 | 0.00049009 | 0.02103755 |
| ENSMUSG00000058440 | Nrf1 | 1.29393689 | 0.0005011 | 0.02147379 |
| ENSMUSG00000018593 | Sparc | -2.0154634 | 0.00050279 | 0.02150991 |
| ENSMUSG00000015745 | Plekho1 | -1.632262 | 0.00051531 | 0.02200143 |
| ENSMUSG00000070867 | Trabd2b | -2.857232 | 0.00051601 | 0.02200143 |
| ENSMUSG00000101174 | Hoxd4 | -5.455952 | 0.00051783 | 0.02204206 |
| ENSMUSG00000022150 | Dab2 | -1.8003308 | 0.00052173 | 0.02215078 |
| ENSMUSG00000032422 | Snx14 | 1.25587673 | 0.00052213 | 0.02215078 |
| ENSMUSG00000018459 | Slc13a3 | -2.2816277 | 0.00052673 | 0.02230897 |
| ENSMUSG00000027242 | Wdr76 | 1.65357359 | 0.00053167 | 0.02245607 |
| ENSMUSG00000027489 | Necab3 | 14.3802186 | 0.00053198 | 0.02245607 |
| ENSMUSG00000035964 | Tmem59l | 20.0296787 | 0.00053402 | 0.02250469 |
| ENSMUSG00000038379 | Ttk | 2.21507752 | 0.0005353 | 0.02250469 |
| ENSMUSG00000045629 | Sh3tc2 | 1.70268766 | 0.00053578 | 0.02250469 |
| ENSMUSG00000095567 | Noc2l | 1.51823402 | 0.00053937 | 0.02261776 |
| ENSMUSG00000022360 | Atad2 | 1.67391231 | 0.00055391 | 0.02318941 |
| ENSMUSG00000038155 | Gstp2 | 3.95519852 | 0.00055546 | 0.02321608 |
| ENSMUSG00000024798 | Htr7 | -6.2859681 | 0.00055672 | 0.0232306 |
| ENSMUSG00000028479 | Gne | 1.68335249 | 0.00056049 | 0.02333059 |
| ENSMUSG00000029521 | Chek2 | 1.54157151 | 0.00056116 | 0.02333059 |
| ENSMUSG00000020312 | Shc2 | -2.7734617 | 0.00056221 | 0.02333059 |
| ENSMUSG00000056515 | Rab31 | -1.8271274 | 0.00056325 | 0.02333059 |
| ENSMUSG00000061689 | Dlgap4 | -1.4261275 | 0.00056371 | 0.02333059 |
| ENSMUSG00000020235 | Fzr1 | 1.35626018 | 0.00056493 | 0.02334314 |
| ENSMUSG00000048031 | Fcrl5 | -13.404824 | 0.00057158 | 0.02357968 |
| ENSMUSG00000073792 | Alg6 | 1.60122072 | 0.00057583 | 0.02371622 |
| ENSMUSG00000060044 | Tmem26 | -3.6305311 | 0.0005901 | 0.02426458 |
| ENSMUSG00000053046 | Brsk2 | -8.5189291 | 0.00059347 | 0.02432781 |
| ENSMUSG00000061306 | Slc38a10 | -1.3600595 | 0.00059421 | 0.02432781 |
| ENSMUSG00000028600 | Podn | -3.9740639 | 0.00059451 | 0.02432781 |
| ENSMUSG00000038295 | Atg9b | -4.7732709 | 0.00060197 | 0.02457783 |
| ENSMUSG00000001403 | Ube2c | 2.00404543 | 0.00060255 | 0.02457783 |
| ENSMUSG00000026725 | Tnn | -16.199182 | 0.0006061 | 0.02462114 |
| ENSMUSG00000079697 | Gm5751 | 101.164736 | 0.00060622 | 0.02462114 |
| ENSMUSG00000025001 | Hells | 1.93713039 | 0.00060652 | 0.02462114 |
| ENSMUSG00000025372 | Baiap2 | 1.55241633 | 0.0006079 | 0.02462335 |
| ENSMUSG00000008398 | Elk3 | -1.589572 | 0.00060872 | 0.02462335 |

| | | | | |
|---|---|---|---|---|
| ENSMUSG00000068196 | Col8a1 | -3.547425 | 0.00060972 | 0.02462335 |
| ENSMUSG00000028906 | Epb41 | 1.31753905 | 0.00061067 | 0.02462335 |
| ENSMUSG00000024696 | Lpxn | -2.0323435 | 0.00061142 | 0.02462335 |
| ENSMUSG00000096035 | Odaph | -7.4859053 | 0.00061274 | 0.02463763 |
| ENSMUSG00000023092 | Fhl1 | -3.2121425 | 0.00061819 | 0.02481744 |
| ENSMUSG00000003379 | Cd79a | -6.5280368 | 0.0006201 | 0.02485499 |
| ENSMUSG00000032589 | Bsn | -2.9784036 | 0.00062132 | 0.0248644 |
| ENSMUSG00000032334 | Loxl1 | -2.6077727 | 0.00062576 | 0.02500269 |
| ENSMUSG00000052752 | Traf7 | 1.21056688 | 0.00062815 | 0.02502213 |
| ENSMUSG00000035455 | Fignl1 | 1.66062935 | 0.00062821 | 0.02502213 |
| ENSMUSG00000017929 | B4galt5 | -2.1641916 | 0.00063306 | 0.02517574 |
| ENSMUSG00000032741 | Tpcn1 | 1.50104159 | 0.0006373 | 0.02530466 |
| ENSMUSG00000072674 | Plac9b | -2.1399183 | 0.00064096 | 0.02538115 |
| ENSMUSG00000087107 | AI662270 | -1.5801382 | 0.00064122 | 0.02538115 |
| ENSMUSG00000020437 | Myo1g | -1.9664607 | 0.00064838 | 0.02562456 |
| ENSMUSG00000028990 | Lzic | 1.55757258 | 0.00065093 | 0.02565186 |
| ENSMUSG00000059832 | Kprp | -5397487.8 | 0.00065109 | 0.02565186 |
| ENSMUSG00000005087 | Cd44 | -2.1258603 | 0.00066555 | 0.02613445 |
| ENSMUSG00000029191 | Rfc1 | 1.25269157 | 0.00066572 | 0.02613445 |
| ENSMUSG00000021835 | Bmp4 | -16.722455 | 0.00066642 | 0.02613445 |
| ENSMUSG00000047989 | Ino80c | -1.4058992 | 0.00067047 | 0.02625269 |
| ENSMUSG00000022123 | Scel | -14.890164 | 0.00067998 | 0.02658407 |
| ENSMUSG00000041515 | Irf8 | -1.5534522 | 0.00068133 | 0.02659589 |
| ENSMUSG00000027746 | Ufm1 | -1.2525883 | 0.00068505 | 0.02670025 |
| ENSMUSG00000033364 | Usp37 | 1.5829811 | 0.00068645 | 0.0267138 |
| ENSMUSG00000022698 | Naa50 | 1.33413727 | 0.00069827 | 0.02713063 |
| ENSMUSG00000052087 | Rgs14 | -1.8309946 | 0.00070079 | 0.02713063 |
| ENSMUSG00000026574 | Dpt | -3.1043275 | 0.00070095 | 0.02713063 |
| ENSMUSG00000041324 | Inhba | -4.6478878 | 0.00070143 | 0.02713063 |
| ENSMUSG00000038668 | Lpar1 | -2.4881289 | 0.00071292 | 0.02753312 |
| ENSMUSG00000026383 | Epb41l5 | 1.53623248 | 0.00071422 | 0.02754144 |
| ENSMUSG00000028560 | Usp1 | 1.54867725 | 0.00072157 | 0.02778281 |
| ENSMUSG00000086552 | Dlx4os | -3.3495369 | 0.00072737 | 0.02796357 |
| ENSMUSG00000033777 | Tlr13 | -2.1199703 | 0.00072892 | 0.02798095 |
| ENSMUSG00000028033 | Kcnq5 | 3.60686515 | 0.00073376 | 0.02812407 |
| ENSMUSG00000051495 | Irf2bp2 | 1.42165443 | 0.00073842 | 0.02822855 |
| ENSMUSG00000032035 | Ets1 | -1.5234977 | 0.00073871 | 0.02822855 |
| ENSMUSG00000025644 | Gm7628 | 4.88981279 | 0.00074054 | 0.02825629 |
| ENSMUSG00000026918 | Brd3 | 1.37218814 | 0.00074222 | 0.0282777 |
| ENSMUSG00000027276 | Jag1 | -1.6935632 | 0.00074498 | 0.0283403 |
| ENSMUSG00000055407 | Map6 | -1.9632276 | 0.00074688 | 0.02837013 |
| ENSMUSG00000030409 | Dmpk | -3.0095322 | 0.00074882 | 0.02840161 |
| ENSMUSG00000020695 | Mrc2 | -2.4135662 | 0.00075761 | 0.02869192 |
| ENSMUSG00000029999 | Tgfa | -5.3937936 | 0.00075888 | 0.02869735 |
| ENSMUSG00000031284 | Pak3 | 1.84073465 | 0.00076416 | 0.02885408 |

| | | | | |
|---|---|---|---|---|
| ENSMUSG00000048707 | Tprn | 1.83449694 | 0.0007659 | 0.02887686 |
| ENSMUSG00000035310 | Lin54 | 1.47383937 | 0.00076864 | 0.02888838 |
| ENSMUSG00000026580 | Selp | -3.9643468 | 0.00076999 | 0.02888838 |
| ENSMUSG00000028381 | Ugcg | -1.8686733 | 0.00077064 | 0.02888838 |
| ENSMUSG00000032549 | Rab6b | -1.8171835 | 0.00077135 | 0.02888838 |
| ENSMUSG00000039385 | Cdh6 | -33.880268 | 0.00077189 | 0.02888838 |
| ENSMUSG00000024660 | Incenp | 1.70127078 | 0.00077636 | 0.02901301 |
| ENSMUSG00000004098 | Col5a3 | -2.7075643 | 0.00078261 | 0.02916529 |
| ENSMUSG00000026879 | Gsn | -2.1007061 | 0.00078273 | 0.02916529 |
| ENSMUSG00000039747 | Orai2 | -1.4966482 | 0.00078723 | 0.02925702 |
| ENSMUSG00000037960 | Card19 | -1.6961512 | 0.00078749 | 0.02925702 |
| ENSMUSG00000021461 | Fancc | 1.40998427 | 0.00079017 | 0.02931362 |
| ENSMUSG00000059498 | Fcgr3 | -2.0425008 | 0.00079316 | 0.02938148 |
| ENSMUSG00000047220 | Ccdc36 | 18.2320098 | 0.00079928 | 0.02956519 |
| ENSMUSG00000032014 | Oaf | -2.4029645 | 0.0008017 | 0.02958451 |
| ENSMUSG00000004099 | Dnmt1 | 1.4457056 | 0.00080274 | 0.02958451 |
| ENSMUSG00000059659 | Gm10069 | 2.84019438 | 0.00080329 | 0.02958451 |
| ENSMUSG00000011179 | Odc1 | -2.9117941 | 0.00080803 | 0.02971572 |
| ENSMUSG00000000805 | Car4 | -18.058854 | 0.00081035 | 0.02975321 |
| ENSMUSG00000038156 | Spon1 | -3.5405347 | 0.00081139 | 0.02975321 |
| ENSMUSG00000011884 | Gltp | -1.9551403 | 0.00081334 | 0.02978198 |
| ENSMUSG00000022142 | Nup155 | 1.50132053 | 0.00081591 | 0.02981679 |
| ENSMUSG00000028427 | Aqp7 | -9.6054097 | 0.00081664 | 0.02981679 |
| ENSMUSG00000055373 | Fut9 | 3.69214469 | 0.00082519 | 0.03007183 |
| ENSMUSG00000009654 | Oit3 | -5.4035597 | 0.00082599 | 0.03007183 |
| ENSMUSG00000035158 | Mitf | -1.7981305 | 0.00082798 | 0.03010092 |
| ENSMUSG00000031101 | Sash3 | -1.7207147 | 0.00083387 | 0.03025594 |
| ENSMUSG00000027208 | Fgf7 | -2.819201 | 0.00083462 | 0.03025594 |
| ENSMUSG00000024590 | Lmnb1 | 1.88139105 | 0.00083821 | 0.0303429 |
| ENSMUSG00000053293 | Pom121 | 1.42364769 | 0.00084399 | 0.03047145 |
| ENSMUSG00000028438 | Kif24 | 1.74960223 | 0.00084416 | 0.03047145 |
| ENSMUSG00000006611 | Hfe | -1.7339563 | 0.00084999 | 0.03063814 |
| ENSMUSG00000087141 | Plcxd2 | -3.1427644 | 0.00085911 | 0.03092301 |
| ENSMUSG00000063659 | Zbtb18 | 1.38360414 | 0.00086063 | 0.0309339 |
| ENSMUSG00000104667 | Gm4961 | 6.35337842 | 0.00087138 | 0.03127601 |
| ENSMUSG00000039145 | Camk1d | -1.7844694 | 0.00087774 | 0.03146006 |
| ENSMUSG00000054013 | Tmem179 | -73.894724 | 0.00089879 | 0.03216543 |
| ENSMUSG00000006763 | Saal1 | 1.37676648 | 0.00089995 | 0.03216543 |
| ENSMUSG00000066975 | Cryba4 | -35.499082 | 0.00090146 | 0.03217403 |
| ENSMUSG00000056531 | Ccdc18 | 2.00032593 | 0.00090442 | 0.03223433 |
| ENSMUSG00000027204 | Fbn1 | -2.377414 | 0.00090724 | 0.03228951 |
| ENSMUSG00000033467 | Crlf2 | -1.6712846 | 0.0009139 | 0.03238666 |
| ENSMUSG00000074874 | Ctla2b | -2.9978741 | 0.00091462 | 0.03238666 |
| ENSMUSG00000028633 | Ctps | 1.54046196 | 0.00091491 | 0.03238666 |
| ENSMUSG00000029119 | Man2b2 | -1.5195478 | 0.00091506 | 0.03238666 |

| | | | | |
|---|---|---|---|---|
| ENSMUSG00000043822 | Adamtsl5 | -2.1008683 | 0.00092085 | 0.03254604 |
| ENSMUSG00000013155 | Enkd1 | 1.56789215 | 0.00092317 | 0.03258279 |
| ENSMUSG00000036053 | Fmnl2 | -1.6820304 | 0.00092694 | 0.03267047 |
| ENSMUSG00000095687 | Rnaset2a | -1.5734651 | 0.00092882 | 0.03268068 |
| ENSMUSG00000030346 | Rad51ap1 | 1.83665244 | 0.0009298 | 0.03268068 |
| ENSMUSG00000022876 | Samsn1 | -2.1955123 | 0.00093374 | 0.03277389 |
| ENSMUSG00000041120 | Nbl1 | -2.1796249 | 0.00094606 | 0.03316036 |
| ENSMUSG00000039298 | Cdk5rap2 | 1.45690216 | 0.00094853 | 0.03317896 |
| ENSMUSG00000029408 | Abcb9 | 2.4969861 | 0.0009492 | 0.03317896 |
| ENSMUSG00000004933 | Matk | -3.0868094 | 0.00095332 | 0.03327721 |
| ENSMUSG00000019990 | Pde7b | -2.9519582 | 0.00095873 | 0.03342017 |
| ENSMUSG00000037108 | Zcwpw1 | 2.81937071 | 0.00096093 | 0.03345088 |
| ENSMUSG00000079592 | C1qtnf5 | -3.3087766 | 0.000964 | 0.03351185 |
| ENSMUSG00000058715 | Fcer1g | -1.9164856 | 0.00097406 | 0.03381522 |
| ENSMUSG00000017144 | Rnd3 | -1.6829143 | 0.00099392 | 0.03445773 |
| ENSMUSG00000050108 | Bpifc | -13.596288 | 0.00099631 | 0.03449351 |
| ENSMUSG00000106361 | Gm35066 | 39.9633169 | 0.00100978 | 0.03487636 |
| ENSMUSG00000068854 | H2bc21 | 1.52843764 | 0.00101011 | 0.03487636 |
| ENSMUSG00000025880 | Smad7 | -2.0484907 | 0.00101798 | 0.03510032 |
| ENSMUSG00000046731 | Kctd11 | -1.9037424 | 0.00102794 | 0.03539559 |
| ENSMUSG00000028678 | Kif2c | 1.96074976 | 0.00103176 | 0.03547919 |
| ENSMUSG00000027313 | Chac1 | -5.2082086 | 0.00103344 | 0.035489 |
| ENSMUSG00000029363 | Rfc5 | 1.5417498 | 0.00103748 | 0.03557938 |
| ENSMUSG00000028701 | Lurap1 | 1.71665686 | 0.00104113 | 0.03565653 |
| ENSMUSG00000032119 | Hinfp | 1.39567066 | 0.00104377 | 0.03569874 |
| ENSMUSG00000004296 | Il12b | -3.4027684 | 0.00104914 | 0.03583419 |
| ENSMUSG00000027353 | Mcm8 | 1.70854461 | 0.00106069 | 0.03615443 |
| ENSMUSG00000021177 | Tdp1 | 1.43942068 | 0.00106136 | 0.03615443 |
| ENSMUSG00000030657 | Xylt1 | -2.4673315 | 0.00106884 | 0.03631927 |
| ENSMUSG00000022305 | Lrp12 | -1.9121023 | 0.00107021 | 0.03631927 |
| ENSMUSG00000021377 | Dek | 1.49668806 | 0.00107048 | 0.03631927 |
| ENSMUSG00000025497 | Cdhr5 | 17.1265953 | 0.00107378 | 0.03636124 |
| ENSMUSG00000004263 | Atn1 | 1.26724687 | 0.0010753 | 0.03636124 |
| ENSMUSG00000037679 | Inf2 | -1.6515578 | 0.00107694 | 0.03636124 |
| ENSMUSG00000073434 | Wdr90 | 1.64681153 | 0.00107941 | 0.03636124 |
| ENSMUSG00000052248 | Zeb2os | -2.2638206 | 0.00108018 | 0.03636124 |
| ENSMUSG00000022584 | Ly6c2 | -3.0048765 | 0.00108149 | 0.03636124 |
| ENSMUSG00000074417 | Gm14548 | -4.3125071 | 0.00108174 | 0.03636124 |
| ENSMUSG00000046994 | Mars2 | 1.57536479 | 0.0010849 | 0.03641923 |
| ENSMUSG00000022514 | Il1rap | -1.6301615 | 0.00108864 | 0.03649661 |
| ENSMUSG00000022817 | Itgb5 | -1.4217541 | 0.00109499 | 0.03666126 |
| ENSMUSG00000107705 | Gm45062 | 412.898761 | 0.00109645 | 0.03666176 |
| ENSMUSG00000037621 | Atoh8 | -13.177353 | 0.00109926 | 0.03670752 |
| ENSMUSG00000021871 | Gm49342 | -1.7202558 | 0.00111214 | 0.03704872 |
| ENSMUSG00000046413 | Irx3os | 2.23165935 | 0.00111239 | 0.03704872 |

| | | | | |
|---|---|---|---|---|
| ENSMUSG00000033498 | Strc | 9.77572972 | 0.00111454 | 0.03707168 |
| ENSMUSG00000041064 | Pif1 | 2.17042554 | 0.00112271 | 0.03729437 |
| ENSMUSG00000022468 | Endou | -5.5260602 | 0.00112668 | 0.03737759 |
| ENSMUSG00000032267 | Usp28 | 1.55594386 | 0.00113407 | 0.03748433 |
| ENSMUSG00000078773 | Rad54b | 1.83722238 | 0.00113432 | 0.03748433 |
| ENSMUSG00000066278 | Vps37b | 1.46568576 | 0.00113444 | 0.03748433 |
| ENSMUSG00000028613 | Lrp8 | -3.4609563 | 0.00113698 | 0.03748433 |
| ENSMUSG00000024011 | Pi16 | -3.3492552 | 0.00113727 | 0.03748433 |
| ENSMUSG00000034708 | Grn | -1.4901047 | 0.00114027 | 0.03753432 |
| ENSMUSG00000005233 | Spc25 | 1.85119317 | 0.00114899 | 0.03777252 |
| ENSMUSG00000026942 | Traf2 | 1.42205365 | 0.00115145 | 0.03780449 |
| ENSMUSG00000028952 | Zbtb48 | 1.56841578 | 0.00115762 | 0.03794911 |
| ENSMUSG00000025133 | Ints4 | 1.36049263 | 0.00115884 | 0.03794911 |
| ENSMUSG00000035273 | Hpse | -2.3436355 | 0.00117592 | 0.03845877 |
| ENSMUSG00000022369 | Mtbp | 1.57605663 | 0.00118287 | 0.03857634 |
| ENSMUSG00000057541 | Pus7 | 1.58002927 | 0.0011834 | 0.03857634 |
| ENSMUSG00000069272 | H2ac8 | -3.4831967 | 0.00118407 | 0.03857634 |
| ENSMUSG00000026841 | Fibcd1 | -26.805306 | 0.00121067 | 0.0393925 |
| ENSMUSG00000094800 | Gm9780 | -2.2289615 | 0.00121782 | 0.03957457 |
| ENSMUSG00000089652 | Gm16025 | 15.555625 | 0.00122226 | 0.039668 |
| ENSMUSG00000024050 | Wiz | 1.32088064 | 0.00122507 | 0.03970855 |
| ENSMUSG00000059791 | Nrm | 1.62945837 | 0.00122844 | 0.03976698 |
| ENSMUSG00000021423 | Ly86 | -1.8480228 | 0.00123567 | 0.03992015 |
| ENSMUSG00000079259 | Trim71 | -8.9627281 | 0.00123754 | 0.03992015 |
| ENSMUSG00000022156 | Gzme | -39.589271 | 0.00123788 | 0.03992015 |
| ENSMUSG00000025225 | Nfkb2 | 1.64632647 | 0.00124931 | 0.04019681 |
| ENSMUSG00000048240 | Gng7 | 2.25367643 | 0.00124963 | 0.04019681 |
| ENSMUSG00000028212 | Ccne2 | 1.92749058 | 0.00125232 | 0.04023255 |
| ENSMUSG00000024791 | Cdca5 | 2.04874354 | 0.00126066 | 0.04044951 |
| ENSMUSG00000030707 | Coro1a | -1.7574102 | 0.00126573 | 0.04056084 |
| ENSMUSG00000028676 | Srsf10 | 1.31266714 | 0.00127123 | 0.04068582 |
| ENSMUSG00000029910 | Mad2l1 | 1.62330608 | 0.00127326 | 0.04069949 |
| ENSMUSG00000000632 | Sez6 | -35.001669 | 0.00127527 | 0.04070862 |
| ENSMUSG00000028874 | Fgr | -1.8031452 | 0.00127675 | 0.04070862 |
| ENSMUSG00000003166 | Dgcr2 | 1.27649268 | 0.00128038 | 0.04074377 |
| ENSMUSG00000038280 | Ostm1 | -1.3469366 | 0.00128106 | 0.04074377 |
| ENSMUSG00000017716 | Birc5 | 1.70022266 | 0.0012869 | 0.04087826 |
| ENSMUSG00000027331 | Knstrn | 1.76304732 | 0.00128972 | 0.04091678 |
| ENSMUSG00000020803 | Txndc17 | -1.5250492 | 0.00129277 | 0.04095415 |
| ENSMUSG00000017861 | Mybl2 | 2.0187952 | 0.00129412 | 0.04095415 |
| ENSMUSG00000026420 | Il24 | -23.330012 | 0.00130562 | 0.04122919 |
| ENSMUSG00000031591 | Asah1 | -1.2832476 | 0.00130606 | 0.04122919 |
| ENSMUSG00000017146 | Brca1 | 1.60476952 | 0.00131315 | 0.04140165 |
| ENSMUSG00000034652 | Cd300a | -2.0977242 | 0.00131629 | 0.04141464 |
| ENSMUSG00000028164 | Manba | -1.6866898 | 0.00131798 | 0.04141464 |

| | | | | |
|---|---|---|---|---|
| ENSMUSG00000094595 | Fsbp | 4.42965984 | 0.00131845 | 0.04141464 |
| ENSMUSG00000063354 | Slc39a4 | 3.4223053 | 0.0013238 | 0.04153125 |
| ENSMUSG00000030789 | Itgax | -2.0491839 | 0.00133116 | 0.04171065 |
| ENSMUSG00000024014 | Pim1 | -2.4191412 | 0.00133672 | 0.04183334 |
| ENSMUSG00000027845 | Dclre1b | 1.35563914 | 0.00133921 | 0.04185972 |
| ENSMUSG00000034247 | Plekhm1 | -1.3120447 | 0.00134796 | 0.04205677 |
| ENSMUSG00000034575 | Tent4a | 1.28522418 | 0.00134882 | 0.04205677 |
| ENSMUSG00000032555 | Topbp1 | 1.5183584 | 0.00135771 | 0.04223447 |
| ENSMUSG00000024395 | Lims2 | -2.6738194 | 0.00135785 | 0.04223447 |
| ENSMUSG00000039208 | Metrnl | -2.2596159 | 0.001361 | 0.04226175 |
| ENSMUSG00000002732 | Fkbp7 | -2.000838 | 0.00136205 | 0.04226175 |
| ENSMUSG00000071715 | Ncf4 | -1.724211 | 0.00136895 | 0.04242417 |
| ENSMUSG00000030484 | Lypd5 | -105.44916 | 0.0013733 | 0.04244152 |
| ENSMUSG00000046275 | Trarg1 | -3.1656219 | 0.00137606 | 0.04244152 |
| ENSMUSG00000001281 | Itgb7 | -2.2529615 | 0.00137639 | 0.04244152 |
| ENSMUSG00000026042 | Col5a2 | -3.026772 | 0.00137713 | 0.04244152 |
| ENSMUSG00000023484 | Prph | 9.47352135 | 0.00137786 | 0.04244152 |
| ENSMUSG00000027254 | Map1a | -2.5671763 | 0.00138546 | 0.04262387 |
| ENSMUSG00000005882 | Uqcc1 | 1.29455914 | 0.0013883 | 0.04264154 |
| ENSMUSG00000114608 | Gm36161 | -2.3183695 | 0.0013901 | 0.04264154 |
| ENSMUSG00000030528 | Blm | 1.60263147 | 0.00139107 | 0.04264154 |
| ENSMUSG00000022656 | Nectin3 | -2.4863677 | 0.00139598 | 0.04274041 |
| ENSMUSG00000026434 | Nucks1 | 1.37610924 | 0.00140708 | 0.04302842 |
| ENSMUSG00000098318 | Lockd | 1.85887315 | 0.00141153 | 0.04311283 |
| ENSMUSG00000019935 | Slc17a8 | -59.013878 | 0.00143693 | 0.04383572 |
| ENSMUSG00000027401 | Tgm3 | -159.39552 | 0.00144113 | 0.04391124 |
| ENSMUSG00000047534 | Mis18bp1 | 1.87659784 | 0.00144291 | 0.04391285 |
| ENSMUSG00000037872 | Ackr1 | -4.2228304 | 0.0014506 | 0.04409414 |
| ENSMUSG00000031250 | Tnmd | 34.6131329 | 0.00145594 | 0.04420345 |
| ENSMUSG00000053398 | Phgdh | 1.54678119 | 0.00146413 | 0.04439907 |
| ENSMUSG00000015355 | Cd48 | -1.8396644 | 0.00146829 | 0.0444723 |
| ENSMUSG00000103280 | Gm37277 | 3.70650055 | 0.0014716 | 0.04451954 |
| ENSMUSG00000048234 | Rnf149 | -1.6648631 | 0.00147396 | 0.04452482 |
| ENSMUSG00000041440 | Gk5 | 1.63605028 | 0.00147528 | 0.04452482 |
| ENSMUSG00000024561 | Mbd1 | 1.32046347 | 0.00147745 | 0.04453739 |
| ENSMUSG00000091649 | Phf11b | -2.2883148 | 0.00148022 | 0.04456807 |
| ENSMUSG00000015090 | Ptgds | 16.432363 | 0.00148431 | 0.04463824 |
| ENSMUSG00000001119 | Col6a1 | -2.2375836 | 0.00148651 | 0.04465153 |
| ENSMUSG00000062585 | Cnr2 | -2.395103 | 0.00149654 | 0.04488227 |
| ENSMUSG00000023467 | Tulp2 | 41.4989992 | 0.00149772 | 0.04488227 |
| ENSMUSG00000038244 | Mical2 | -1.4417592 | 0.00150139 | 0.04493904 |
| ENSMUSG00000020493 | Prr11 | 1.94396229 | 0.00151854 | 0.04539896 |
| ENSMUSG00000078521 | Aunip | 2.02232152 | 0.00152406 | 0.04541784 |
| ENSMUSG00000000555 | Itga5 | -2.1078563 | 0.00152445 | 0.04541784 |
| ENSMUSG00000115302 | Gm49394 | -1823165.1 | 0.00152453 | 0.04541784 |

| | | | | |
|---|---|---|---|---|
| ENSMUSG00000003948 | Mmd | -1.6702219 | 0.00153939 | 0.04578279 |
| ENSMUSG00000063931 | Pepd | -1.284238 | 0.00154115 | 0.04578279 |
| ENSMUSG00000037692 | Ahdc1 | 1.33956823 | 0.00154219 | 0.04578279 |
| ENSMUSG00000032020 | Ubash3b | -1.8123673 | 0.00154977 | 0.04589829 |
| ENSMUSG00000051650 | B3gnt2 | -1.7164877 | 0.0015502 | 0.04589829 |
| ENSMUSG00000026355 | Mcm6 | 1.51542073 | 0.0015515 | 0.04589829 |
| ENSMUSG00000045573 | Penk | -4.6748088 | 0.00155357 | 0.04590629 |
| ENSMUSG00000022416 | Cacna1i | -4.3024965 | 0.00155878 | 0.04600672 |
| ENSMUSG00000027559 | Car3 | -5.7269281 | 0.00158879 | 0.04679619 |
| ENSMUSG00000044197 | Gpr146 | -1.5966091 | 0.00158921 | 0.04679619 |
| ENSMUSG00000028822 | Tmem50a | -1.3145728 | 0.00159584 | 0.0469369 |
| ENSMUSG00000005824 | Tnfsf14 | -2.7514653 | 0.00159782 | 0.04694078 |
| ENSMUSG00000032221 | Mns1 | 1.67498383 | 0.00160292 | 0.04703636 |
| ENSMUSG00000037991 | Rmi2 | 2.27529978 | 0.00160647 | 0.04708607 |
| ENSMUSG00000025068 | Gsto1 | -1.6546841 | 0.00161694 | 0.0473384 |
| ENSMUSG00000026700 | Tnfsf4 | -14.805871 | 0.00163602 | 0.04784188 |
| ENSMUSG00000039697 | Ncoa7 | 1.69535958 | 0.00164192 | 0.04794623 |
| ENSMUSG00000036923 | Stox1 | 4.99771612 | 0.00164336 | 0.04794623 |
| ENSMUSG00000028459 | Cd72 | -2.1713985 | 0.00165187 | 0.0481042 |
| ENSMUSG00000020264 | Slc36a2 | -5.8089116 | 0.00165256 | 0.0481042 |
| ENSMUSG00000033014 | Trim33 | 1.39507696 | 0.00166517 | 0.04841562 |
| ENSMUSG00000027115 | Kif18a | 1.78545087 | 0.00166839 | 0.04845391 |
| ENSMUSG00000045763 | Basp1 | -1.882176 | 0.00168044 | 0.04874817 |
| ENSMUSG00000038976 | Ppp1r9b | -1.3745891 | 0.00168253 | 0.04875316 |
| ENSMUSG00000090231 | Cfb | -1.8969878 | 0.00171255 | 0.04956635 |
| ENSMUSG00000020898 | Ctc1 | 1.26417998 | 0.00173148 | 0.05005718 |
| ENSMUSG00000037628 | Cdkn3 | 2.17747956 | 0.00174345 | 0.05034609 |
| ENSMUSG00000029096 | Htra3 | -2.4459167 | 0.0017499 | 0.05042003 |
| ENSMUSG00000102748 | Pcdhgb2 | 4.63071692 | 0.00174998 | 0.05042003 |
| ENSMUSG00000030401 | Rtn2 | -2.8280387 | 0.00175212 | 0.05042446 |
| ENSMUSG00000029610 | Aimp2 | 1.46366991 | 0.00175572 | 0.05042877 |
| ENSMUSG00000060181 | Slc35e3 | 1.41947774 | 0.00175624 | 0.05042877 |
| ENSMUSG00000074146 | 4930579C12I | 5.31584644 | 0.00176404 | 0.05059558 |
| ENSMUSG00000112137 | Gm47865 | 5.49260093 | 0.00176698 | 0.0506228 |
| ENSMUSG00000028673 | Fuca1 | -1.5309865 | 0.00176966 | 0.05064248 |
| ENSMUSG00000058186 | Zfp980 | -58.113588 | 0.00177332 | 0.05069031 |
| ENSMUSG00000033222 | Ttf2 | 1.42133878 | 0.00178238 | 0.05089185 |
| ENSMUSG00000074994 | Qser1 | 1.54014982 | 0.00179689 | 0.05124864 |
| ENSMUSG00000020611 | Gna13 | -1.2671393 | 0.00180227 | 0.05134463 |
| ENSMUSG00000093587 | Gm20554 | 2.10576523 | 0.00181011 | 0.05151021 |
| ENSMUSG00000021591 | Glrx | -1.5978612 | 0.00181455 | 0.05157862 |
| ENSMUSG00000064289 | Tank | 1.346085 | 0.00184001 | 0.0521859 |
| ENSMUSG00000034484 | Snx2 | -1.216501 | 0.00184003 | 0.0521859 |
| ENSMUSG00000076523 | Igkv15-103 | -193.53338 | 0.00184207 | 0.0521859 |
| ENSMUSG00000031928 | Mre11a | 1.39975482 | 0.00184435 | 0.05219222 |

| | | | | |
|---|---|---|---|---|
| ENSMUSG00000025574 | Tk1 | 1.5354948 | 0.00185826 | 0.05252734 |
| ENSMUSG00000040204 | Pclaf | 1.82705588 | 0.00186534 | 0.05266898 |
| ENSMUSG00000028063 | Lmna | -1.3678059 | 0.00187099 | 0.05276992 |
| ENSMUSG00000040250 | Ints13 | 1.30865948 | 0.00188374 | 0.0530499 |
| ENSMUSG00000022790 | Igsf11 | -19.611301 | 0.0018851 | 0.0530499 |
| ENSMUSG00000031266 | Gla | -2.0680514 | 0.00189084 | 0.05315261 |
| ENSMUSG00000038623 | Tm6sf1 | -1.7240372 | 0.00189635 | 0.05321507 |
| ENSMUSG00000089809 | Rasgef1b | -2.5655696 | 0.00189725 | 0.05321507 |
| ENSMUSG00000029659 | Medag | -2.4923153 | 0.0019048 | 0.0533147 |
| ENSMUSG00000030671 | Pde3b | -2.0851135 | 0.001905 | 0.0533147 |
| ENSMUSG00000002297 | Dbf4 | 1.76171 | 0.00190983 | 0.05339119 |
| ENSMUSG00000029156 | Sgcb | -1.7579867 | 0.00191314 | 0.05342494 |
| ENSMUSG00000074039 | 4930520O04 | 3.54546928 | 0.00191779 | 0.05343936 |
| ENSMUSG00000113683 | Gm47123 | -112.97091 | 0.00191786 | 0.05343936 |
| ENSMUSG00000026482 | Rgl1 | -1.5709034 | 0.00194071 | 0.0540169 |
| ENSMUSG00000038871 | Bpgm | -1.8019404 | 0.00194687 | 0.0541289 |
| ENSMUSG00000028047 | Thbs3 | -2.4815591 | 0.00195029 | 0.05415491 |
| ENSMUSG00000053173 | Rpl18-ps2 | 3.48333542 | 0.00195207 | 0.05415491 |
| ENSMUSG00000000682 | Cd52 | -1.858244 | 0.00195637 | 0.05421514 |
| ENSMUSG00000037944 | Ccr7 | -3.5641672 | 0.00198362 | 0.05491033 |
| ENSMUSG00000054074 | Skida1 | 2.1232832 | 0.00198707 | 0.05494619 |
| ENSMUSG00000031949 | Adat1 | 1.66061646 | 0.00199647 | 0.0551461 |
| ENSMUSG00000043157 | Arl11 | -1.9609892 | 0.00200297 | 0.05521993 |
| ENSMUSG00000018822 | Sfrp5 | -9.5966272 | 0.00200349 | 0.05521993 |
| ENSMUSG00000025969 | Nrp2 | -1.990247 | 0.00200729 | 0.05524697 |
| ENSMUSG00000118181 | CAAA010661 | -1.6461838 | 0.00200882 | 0.05524697 |
| ENSMUSG00000026646 | Suv39h2 | 1.76625187 | 0.00202286 | 0.05557295 |
| ENSMUSG00000059173 | Pde1a | -2.8663334 | 0.00203 | 0.05568729 |
| ENSMUSG00000026955 | Sapcd2 | 2.17013691 | 0.00203141 | 0.05568729 |
| ENSMUSG00000063506 | Arhgap22 | -1.947224 | 0.00203792 | 0.05575368 |
| ENSMUSG00000050896 | Rtn4rl2 | -4.3345022 | 0.00203933 | 0.05575368 |
| ENSMUSG00000020583 | Matn3 | 4.9436776 | 0.00204041 | 0.05575368 |
| ENSMUSG00000066684 | Pilrb1 | -2.5822227 | 0.00204754 | 0.05588851 |
| ENSMUSG00000023072 | Cep89 | 1.40015967 | 0.0020508 | 0.05591724 |
| ENSMUSG00000038147 | Cd84 | -2.1380893 | 0.00205402 | 0.05594503 |
| ENSMUSG00000001227 | Sema6b | -1.8929674 | 0.00207442 | 0.05644021 |
| ENSMUSG00000063011 | Msln | 38.4219165 | 0.00210395 | 0.05718238 |
| ENSMUSG00000031697 | Orc6 | 1.47681372 | 0.00210975 | 0.0572284 |
| ENSMUSG00000001525 | Tubb5 | 1.41228067 | 0.00211062 | 0.0572284 |
| ENSMUSG00000021175 | Cdca7l | 1.48674571 | 0.0021124 | 0.0572284 |
| ENSMUSG00000003363 | Pld3 | -1.6947236 | 0.00211727 | 0.05729928 |
| ENSMUSG00000016087 | Fli1 | -1.6745305 | 0.00213821 | 0.05779189 |
| ENSMUSG00000041329 | Atp1b2 | -4.1380829 | 0.00214002 | 0.05779189 |
| ENSMUSG00000018574 | Acadvl | -1.4685661 | 0.00214789 | 0.05791348 |
| ENSMUSG00000041406 | BC055324 | 1.65791686 | 0.00215126 | 0.05791348 |

| | | | | |
|---|---|---|---|---|
| ENSMUSG00000051586 | Mical3 | 1.41431504 | 0.00215136 | 0.05791348 |
| ENSMUSG00000033763 | Mtss2 | 2.0346758 | 0.00216771 | 0.05829178 |
| ENSMUSG00000006649 | Nphs1 | 6.81416886 | 0.00218824 | 0.05878184 |
| ENSMUSG00000047443 | Erfe | 2.83221091 | 0.00220111 | 0.05906508 |
| ENSMUSG00000002332 | Dhrs1 | -1.4389041 | 0.00223278 | 0.05985171 |
| ENSMUSG00000050100 | Hmx2 | 16.5716166 | 0.0022511 | 0.06024289 |
| ENSMUSG00000025810 | Nrp1 | -2.0608775 | 0.00225211 | 0.06024289 |
| ENSMUSG00000037572 | Wdhd1 | 1.69719495 | 0.00226158 | 0.06043249 |
| ENSMUSG00000024737 | Slc15a3 | -2.1498779 | 0.0022737 | 0.06069263 |
| ENSMUSG00000021880 | Rnase6 | -2.3580627 | 0.00229667 | 0.06124144 |
| ENSMUSG00000042272 | Sestd1 | 1.33942543 | 0.00231761 | 0.06173492 |
| ENSMUSG00000025825 | Iscu | -1.2782823 | 0.00232136 | 0.06176993 |
| ENSMUSG00000030400 | Ercc2 | 1.32429554 | 0.00232742 | 0.06186663 |
| ENSMUSG00000055546 | Timd4 | -10.849834 | 0.00233091 | 0.06189462 |
| ENSMUSG00000047586 | Nccrp1 | -22.109406 | 0.00233734 | 0.06193177 |
| ENSMUSG00000073982 | Rhog | -1.4070297 | 0.00233931 | 0.06193177 |
| ENSMUSG00000039936 | Pik3cd | -1.4659384 | 0.00233962 | 0.06193177 |
| ENSMUSG00000052698 | Tln2 | 1.67041853 | 0.00234338 | 0.06196667 |
| ENSMUSG00000019214 | Chtf18 | 1.71299979 | 0.00234692 | 0.06199577 |
| ENSMUSG00000053113 | Socs3 | -1.8190265 | 0.00235575 | 0.06210693 |
| ENSMUSG00000017478 | Zc3h18 | 1.2351553 | 0.00235839 | 0.06210693 |
| ENSMUSG00000028927 | Padi2 | -2.465323 | 0.00235846 | 0.06210693 |
| ENSMUSG00000015013 | Trappc2l | -1.316122 | 0.00237098 | 0.06237196 |
| ENSMUSG00000047976 | Kcna1 | -56.941681 | 0.00237979 | 0.06253897 |
| ENSMUSG00000045273 | Cenph | 1.78597709 | 0.00238235 | 0.0625416 |
| ENSMUSG00000034317 | Trim59 | 1.57916403 | 0.00238843 | 0.06263644 |
| ENSMUSG00000002504 | Slc9a3r2 | 1.49524248 | 0.00239686 | 0.06273831 |
| ENSMUSG00000047757 | Fancb | 1.62174823 | 0.00239725 | 0.06273831 |
| ENSMUSG00000100254 | Trpc2 | 117.000226 | 0.00240422 | 0.06285593 |
| ENSMUSG00000031093 | Dock11 | -2.2146507 | 0.00241928 | 0.0631495 |
| ENSMUSG00000037295 | Ldlrap1 | -1.3779948 | 0.00242042 | 0.0631495 |
| ENSMUSG00000033149 | Phldb2 | -1.7590698 | 0.00243195 | 0.06335006 |
| ENSMUSG00000042213 | Zfand4 | 1.86842455 | 0.00243309 | 0.06335006 |
| ENSMUSG00000051517 | Arhgef39 | 2.01385579 | 0.0024406 | 0.06348071 |
| ENSMUSG00000022010 | Tsc22d1 | -1.5884583 | 0.00245053 | 0.06367336 |
| ENSMUSG00000040621 | Gemin8 | 1.61835642 | 0.00245302 | 0.06367336 |
| ENSMUSG00000030278 | Cidec | -5.1994075 | 0.00246043 | 0.06378718 |
| ENSMUSG00000041961 | Znrf3 | -2.1970176 | 0.00246243 | 0.06378718 |
| ENSMUSG00000033883 | Zfp267 | 1.74599217 | 0.00246928 | 0.06386351 |
| ENSMUSG00000026343 | Gpr39 | -4.3372281 | 0.0024704 | 0.06386351 |
| ENSMUSG00000026029 | Casp8 | -1.3293221 | 0.0024785 | 0.06400766 |
| ENSMUSG00000030177 | Ccdc77 | 1.49012818 | 0.00248808 | 0.0641876 |
| ENSMUSG00000032113 | Chek1 | 1.78771034 | 0.00249156 | 0.0641876 |
| ENSMUSG00000074604 | Mgst2 | 4.022477 | 0.00249304 | 0.0641876 |
| ENSMUSG00000029054 | Gabrd | 8.74050565 | 0.0024996 | 0.06419162 |

| | | | | |
|---|---|---|---|---|
| ENSMUSG00000024542 | Cep192 | 1.56722875 | 0.00249985 | 0.06419162 |
| ENSMUSG00000039456 | Morc3 | 1.28928296 | 0.00250283 | 0.06419162 |
| ENSMUSG00000020241 | Col6a2 | -2.3775211 | 0.0025033 | 0.06419162 |
| ENSMUSG00000027864 | Ptgfrn | -1.7711794 | 0.00251448 | 0.06441319 |
| ENSMUSG00000068101 | Cenpm | 1.79047516 | 0.00252198 | 0.06452481 |
| ENSMUSG00000046546 | Fam43a | -1.9150526 | 0.00252391 | 0.06452481 |
| ENSMUSG00000007655 | Cav1 | -1.8555401 | 0.0025281 | 0.06456696 |
| ENSMUSG00000055447 | Cd47 | -1.3216746 | 0.00253716 | 0.06473318 |
| ENSMUSG00000022070 | Bora | 1.7738099 | 0.00254561 | 0.06484546 |
| ENSMUSG00000021485 | Mxd3 | 1.72023148 | 0.00254666 | 0.06484546 |
| ENSMUSG00000113204 | Gm46430 | 2.24196632 | 0.00255678 | 0.06498461 |
| ENSMUSG00000029227 | Fip1l1 | 1.22152889 | 0.00255724 | 0.06498461 |
| ENSMUSG00000017781 | Pitpna | -1.2729905 | 0.00256182 | 0.06503597 |
| ENSMUSG00000086924 | Gm11766 | 4.01137303 | 0.00256961 | 0.06516867 |
| ENSMUSG00000022667 | Cd200r1 | -2.8158347 | 0.00257368 | 0.0652067 |
| ENSMUSG00000043183 | Simc1 | 1.49825622 | 0.00257853 | 0.06526456 |
| ENSMUSG00000015652 | Steap1 | -2.3078381 | 0.00259134 | 0.06552348 |
| ENSMUSG00000025534 | Gusb | -1.5072164 | 0.00259743 | 0.06561048 |
| ENSMUSG00000002741 | Ykt6 | -1.2459208 | 0.00260042 | 0.06561048 |
| ENSMUSG00000021303 | Gng4 | -51.243352 | 0.00260508 | 0.06561048 |
| ENSMUSG00000078886 | Gm2026 | 3.41681607 | 0.00260511 | 0.06561048 |
| ENSMUSG00000022971 | Ifnar2 | -1.285443 | 0.00261338 | 0.06575362 |
| ENSMUSG00000059060 | Rad51b | 2.59728524 | 0.00262267 | 0.06592205 |
| ENSMUSG00000021356 | Irf4 | -3.4165138 | 0.00262626 | 0.06594706 |
| ENSMUSG00000006205 | Htra1 | -2.1567555 | 0.0026328 | 0.06604614 |
| ENSMUSG00000066800 | Rnasel | -1.6865355 | 0.00265756 | 0.06660156 |
| ENSMUSG00000031442 | Mcf2l | 1.40117423 | 0.00266784 | 0.06678349 |
| ENSMUSG00000014329 | Bicc1 | -2.3697857 | 0.00267008 | 0.06678349 |
| ENSMUSG00000086454 | Platr14 | 14.9865732 | 0.00267993 | 0.0669397 |
| ENSMUSG00000026546 | Cfap45 | 2.10040605 | 0.00268159 | 0.0669397 |
| ENSMUSG00000023143 | Nagpa | -1.4228859 | 0.00270966 | 0.06752857 |
| ENSMUSG00000054293 | P2ry10b | -1.6999576 | 0.00271128 | 0.06752857 |
| ENSMUSG00000039405 | Prss23 | -2.1873481 | 0.00271315 | 0.06752857 |
| ENSMUSG00000005370 | Msh6 | 1.35294133 | 0.00271674 | 0.0675516 |
| ENSMUSG00000028060 | Khdc4 | 1.45460905 | 0.00272562 | 0.06770634 |
| ENSMUSG00000019303 | Psmc3ip | 1.55155926 | 0.00273524 | 0.06787883 |
| ENSMUSG00000058809 | Hspd1-ps3 | 2.23407488 | 0.00273966 | 0.06792223 |
| ENSMUSG00000039497 | Dse | -1.4071451 | 0.00275321 | 0.06819151 |
| ENSMUSG00000040473 | Cfap69 | -2.2854169 | 0.00276279 | 0.06836221 |
| ENSMUSG00000072812 | Ahnak2 | -1.8482267 | 0.00277386 | 0.06856934 |
| ENSMUSG00000046387 | Pcdhb17 | 1.98785944 | 0.0027977 | 0.06909155 |
| ENSMUSG00000024672 | Ms4a7 | -1.9336828 | 0.00281002 | 0.06932843 |
| ENSMUSG00000021458 | Aopep | -1.3186414 | 0.00281316 | 0.06933851 |
| ENSMUSG00000000278 | Scpep1 | -1.6066696 | 0.00282115 | 0.06942772 |
| ENSMUSG00000000392 | Fap | -2.96078 | 0.00282224 | 0.06942772 |

| | | | | |
|---|---|---|---|---|
| ENSMUSG00000031594 | Fgl1 | -8.435988 | 0.00282853 | 0.0695151 |
| ENSMUSG00000029238 | Clock | 1.47572582 | 0.00285195 | 0.0699922 |
| ENSMUSG00000024725 | Ostf1 | -1.3643565 | 0.00285601 | 0.0699922 |
| ENSMUSG00000073490 | Ifi207 | -1.7151668 | 0.0028586 | 0.0699922 |
| ENSMUSG00000031478 | Nek3 | 1.64291646 | 0.00286154 | 0.0699922 |
| ENSMUSG00000029082 | Bst1 | -2.5751374 | 0.00286171 | 0.0699922 |
| ENSMUSG00000028759 | Hp1bp3 | 1.24210092 | 0.00286552 | 0.07001796 |
| ENSMUSG00000001415 | Smg5 | 1.2371498 | 0.00289626 | 0.07070105 |
| ENSMUSG00000094915 | AC168977.2 | 8.78667191 | 0.0029011 | 0.0707512 |
| ENSMUSG00000002111 | Spi1 | -1.5794578 | 0.00291076 | 0.07091869 |
| ENSMUSG00000024268 | Celf4 | 1.55824579 | 0.00291485 | 0.0709504 |
| ENSMUSG00000030577 | Cd22 | -2.7387435 | 0.00294098 | 0.07151794 |
| ENSMUSG00000051506 | Wdfy4 | -1.8165445 | 0.00294744 | 0.07160634 |
| ENSMUSG00000067847 | Romo1 | -1.531259 | 0.00295592 | 0.07170864 |
| ENSMUSG00000018736 | Ndel1 | -1.390635 | 0.00295729 | 0.07170864 |
| ENSMUSG00000021248 | Tmed10 | -1.3248276 | 0.00298212 | 0.07224171 |
| ENSMUSG00000076545 | Igkv4-72 | -258.94848 | 0.00298609 | 0.07226906 |
| ENSMUSG00000054951 | 9130008F23I | 1.87146867 | 0.00299854 | 0.07250145 |
| ENSMUSG00000074731 | Zfp345 | 6.8914931 | 0.00300579 | 0.0726075 |
| ENSMUSG00000055531 | Cpsf6 | 1.34153253 | 0.00301947 | 0.0728654 |
| ENSMUSG00000040860 | Crocc | 1.49902263 | 0.0030222 | 0.0728654 |
| ENSMUSG00000097767 | Miat | -28.440741 | 0.0030526 | 0.07350138 |
| ENSMUSG00000021176 | Efcab11 | 1.94396944 | 0.00305436 | 0.07350138 |
| ENSMUSG00000021288 | Klc1 | -1.3082657 | 0.00307437 | 0.07391283 |
| ENSMUSG00000036469 | Marchf1 | -1.6425755 | 0.00308914 | 0.07419611 |
| ENSMUSG00000049409 | Prokr1 | -5.4858277 | 0.00309199 | 0.07419611 |
| ENSMUSG00000044811 | Cd300c2 | -1.6599946 | 0.00309899 | 0.07425217 |
| ENSMUSG00000021306 | Gpr137b | -2.2363597 | 0.0031017 | 0.07425217 |
| ENSMUSG00000070880 | Gad1 | 10.0750339 | 0.00310534 | 0.07425217 |
| ENSMUSG00000028111 | Ctsk | -3.600294 | 0.00310602 | 0.07425217 |
| ENSMUSG00000029718 | Pcolce | -1.9420445 | 0.00311513 | 0.07440008 |
| ENSMUSG00000005667 | Mthfd2 | 1.70185186 | 0.00312474 | 0.07455951 |
| ENSMUSG00000029071 | Dvl1 | 1.35037795 | 0.00313641 | 0.07476769 |
| ENSMUSG00000011148 | Adssl1 | -1.9321692 | 0.00315638 | 0.07517343 |
| ENSMUSG00000022607 | Ptk2 | 1.23671167 | 0.00318214 | 0.07561693 |
| ENSMUSG00000023827 | Agpat4 | -2.1580059 | 0.00318264 | 0.07561693 |
| ENSMUSG00000039384 | Dusp10 | -2.8206823 | 0.00318445 | 0.07561693 |
| ENSMUSG00000061130 | Ppm1b | 1.17916392 | 0.00318691 | 0.07561693 |
| ENSMUSG00000006403 | Adamts4 | -2.2820488 | 0.00319263 | 0.07568209 |
| ENSMUSG00000020524 | Gria1 | -18.730992 | 0.00320175 | 0.07582756 |
| ENSMUSG00000031165 | Was | -1.7001893 | 0.00321008 | 0.07595396 |
| ENSMUSG00000022018 | Rgcc | -2.0001665 | 0.00322179 | 0.07615271 |
| ENSMUSG00000002897 | Il17ra | -1.3620995 | 0.00322447 | 0.07615271 |
| ENSMUSG00000100642 | Gm28230 | -13.524218 | 0.0032407 | 0.07646499 |
| ENSMUSG00000013150 | Gfod2 | 1.38802853 | 0.00324817 | 0.07657007 |

| | | | | |
|---|---|---|---|---|
| ENSMUSG00000033186 | Mzt1 | 1.19733893 | 0.00327489 | 0.07712844 |
| ENSMUSG00000018906 | P4ha2 | -2.7650694 | 0.00328097 | 0.07720001 |
| ENSMUSG00000042684 | Npl | -2.2175211 | 0.00328609 | 0.07724905 |
| ENSMUSG00000037461 | Ints7 | 1.30609764 | 0.00329792 | 0.07740988 |
| ENSMUSG00000000794 | Kcnn3 | 7.03558209 | 0.00330142 | 0.07740988 |
| ENSMUSG00000079614 | Seh1l | 1.27837564 | 0.00330207 | 0.07740988 |
| ENSMUSG00000091345 | Col6a5 | -16.667512 | 0.00331673 | 0.07768176 |
| ENSMUSG00000026069 | Il1rl1 | -3.0049801 | 0.00333578 | 0.07805611 |
| ENSMUSG00000068758 | Il3ra | -2.0928894 | 0.00333975 | 0.07807701 |
| ENSMUSG00000027955 | Gask1b | -2.3294474 | 0.00334548 | 0.07808818 |
| ENSMUSG00000021388 | Aspn | -2.321836 | 0.00334637 | 0.07808818 |
| ENSMUSG00000032782 | Cntrob | 1.3696169 | 0.00335065 | 0.07811635 |
| ENSMUSG00000002055 | Spag5 | 1.52915968 | 0.00335609 | 0.0781714 |
| ENSMUSG00000049327 | Kmt5a | 1.45325997 | 0.00336597 | 0.07832981 |
| ENSMUSG00000028655 | Mfsd2a | -2.1109219 | 0.00337967 | 0.07857666 |
| ENSMUSG00000052406 | Rexo4 | 1.27965958 | 0.00338794 | 0.07869694 |
| ENSMUSG00000068859 | Sp9 | -18.086708 | 0.00340898 | 0.07905633 |
| ENSMUSG00000036172 | Cd200r3 | -9.2019197 | 0.00341027 | 0.07905633 |
| ENSMUSG00000020467 | Efemp1 | -2.9834021 | 0.00341275 | 0.07905633 |
| ENSMUSG00000048779 | P2ry6 | -1.5992944 | 0.00341592 | 0.07905781 |
| ENSMUSG00000033031 | Cip2a | 1.77509293 | 0.00342222 | 0.07913135 |
| ENSMUSG00000042029 | Ncapg2 | 1.58118203 | 0.00344002 | 0.07940624 |
| ENSMUSG00000049988 | Lrrc25 | -1.6089635 | 0.00344035 | 0.07940624 |
| ENSMUSG00000026395 | Ptprc | -1.7510546 | 0.00344566 | 0.07945652 |
| ENSMUSG00000085890 | Tnfsf13os | 3.160284 | 0.00345509 | 0.07957052 |
| ENSMUSG00000054484 | Tmem62 | -1.6064663 | 0.00345754 | 0.07957052 |
| ENSMUSG00000021965 | Ska3 | 1.80768518 | 0.00346 | 0.07957052 |
| ENSMUSG00000039509 | Nup133 | 1.35653466 | 0.00346573 | 0.07963025 |
| ENSMUSG00000030870 | Ubfd1 | 1.30206016 | 0.00347106 | 0.07968065 |
| ENSMUSG00000033596 | Rfwd3 | 1.32180651 | 0.00347468 | 0.07969174 |
| ENSMUSG00000036285 | Noa1 | 1.50785736 | 0.00348862 | 0.07993953 |
| ENSMUSG00000024803 | Ankrd1 | 6.30905979 | 0.00351071 | 0.08037313 |
| ENSMUSG00000031016 | Wee1 | 1.47495045 | 0.00351613 | 0.08042484 |
| ENSMUSG00000020774 | Aspa | 5.02843534 | 0.00352558 | 0.08056844 |
| ENSMUSG00000022265 | Ank | -2.0492043 | 0.00355745 | 0.08122372 |
| ENSMUSG00000070524 | Fcrlb | -4.3184698 | 0.00356285 | 0.08127384 |
| ENSMUSG00000061731 | Ext1 | -1.3649599 | 0.00356719 | 0.08129991 |
| ENSMUSG00000000561 | Wdr77 | 1.40500127 | 0.00357366 | 0.08137448 |
| ENSMUSG00000029757 | Dync1i1 | -3.9352995 | 0.00358626 | 0.08158825 |
| ENSMUSG00000020431 | Adcy1 | -22.540353 | 0.00359245 | 0.08165588 |
| ENSMUSG00000104453 | Gm37829 | 2.6795801 | 0.00361578 | 0.08211283 |
| ENSMUSG00000021322 | Aoah | -2.1734753 | 0.00362363 | 0.08221766 |
| ENSMUSG00000062248 | Cks2 | 1.75426555 | 0.0036317 | 0.08230524 |
| ENSMUSG00000089911 | Mfsd14a | 1.22801059 | 0.00363715 | 0.08230524 |
| ENSMUSG00000015340 | Cybb | -1.6157191 | 0.00363721 | 0.08230524 |

| | | | | |
|---|---|---|---|---|
| ENSMUSG00000009185 | Ccl8 | -2.5526889 | 0.00368632 | 0.08334229 |
| ENSMUSG00000102893 | Gm37855 | 7.17977255 | 0.00369396 | 0.08344073 |
| ENSMUSG00000047281 | Sfn | -2.4320708 | 0.00370546 | 0.08362627 |
| ENSMUSG00000022421 | Nptxr | 2.04021181 | 0.00372351 | 0.08391146 |
| ENSMUSG00000022848 | Slc49a4 | -1.3925542 | 0.0037247 | 0.08391146 |
| ENSMUSG00000042606 | Hirip3 | 1.51699078 | 0.00375416 | 0.08450016 |
| ENSMUSG00000036944 | Tmem71 | -1.8762098 | 0.00375867 | 0.08452673 |
| ENSMUSG00000024502 | Jakmip2 | -11.441455 | 0.00376518 | 0.08454492 |
| ENSMUSG00000025436 | Atp23 | 1.5456404 | 0.00376613 | 0.08454492 |
| ENSMUSG00000107383 | Gm4366 | 1.97906619 | 0.00377487 | 0.08466633 |
| ENSMUSG00000034203 | Chchd4 | 1.45681955 | 0.00378785 | 0.08472467 |
| ENSMUSG00000030092 | Cntn6 | -182.23205 | 0.00378812 | 0.08472467 |
| ENSMUSG00000009741 | Ubp1 | 1.34224013 | 0.00379248 | 0.08472467 |
| ENSMUSG00000066979 | Bub3 | 1.30863612 | 0.00379468 | 0.08472467 |
| ENSMUSG00000056529 | Ptafr | -1.8279507 | 0.00379535 | 0.08472467 |
| ENSMUSG00000050107 | Haspin | 1.70687308 | 0.00379747 | 0.08472467 |
| ENSMUSG00000047867 | Gimap6 | -1.9645281 | 0.00382814 | 0.08530671 |
| ENSMUSG00000026853 | Crat | -1.2502238 | 0.00383074 | 0.08530671 |
| ENSMUSG00000117818 | Gm50105 | 3.06949809 | 0.00383363 | 0.08530671 |
| ENSMUSG00000097327 | E030030I06R | 2.55853758 | 0.00386195 | 0.08586173 |
| ENSMUSG00000000248 | Clec2g | -9.0123507 | 0.00388988 | 0.08640709 |
| ENSMUSG00000040594 | Ranbp17 | 1.59759124 | 0.00390835 | 0.08674153 |
| ENSMUSG00000028876 | Epha10 | 23.4399044 | 0.0039175 | 0.0868611 |
| ENSMUSG00000044641 | Pard6b | 2.19565113 | 0.00392058 | 0.0868611 |
| ENSMUSG00000001687 | Ubl3 | -1.2090991 | 0.00392596 | 0.08690454 |
| ENSMUSG00000029311 | Hsd17b11 | -1.3261132 | 0.00393821 | 0.08709981 |
| ENSMUSG00000031253 | Srpx2 | -3.3178436 | 0.0039508 | 0.08730241 |
| ENSMUSG00000050150 | Slc9b1 | 8.31759308 | 0.00397223 | 0.08762564 |
| ENSMUSG00000033075 | Senp1 | 1.35560313 | 0.00397233 | 0.08762564 |
| ENSMUSG00000054517 | Trim65 | 1.51788808 | 0.00398068 | 0.0877054 |
| ENSMUSG00000024032 | Tff1 | -18.341599 | 0.00398284 | 0.0877054 |
| ENSMUSG00000089942 | Pira2 | -3.0212758 | 0.00399229 | 0.08783722 |
| ENSMUSG00000002068 | Ccne1 | 1.76664216 | 0.00400765 | 0.08809892 |
| ENSMUSG00000027933 | Ints3 | 1.19545395 | 0.00402015 | 0.08829727 |
| ENSMUSG00000105373 | Gm42429 | 3.95816048 | 0.00404168 | 0.08869367 |
| ENSMUSG00000073433 | Arhgdig | 1.7716542 | 0.00404731 | 0.08874042 |
| ENSMUSG00000023953 | Polh | 1.42531399 | 0.00405902 | 0.0889205 |
| ENSMUSG00000024151 | Msh2 | 1.28944846 | 0.004065 | 0.08897476 |
| ENSMUSG00000029664 | Tfpi2 | -2.945755 | 0.00407918 | 0.08920844 |
| ENSMUSG00000037395 | Rcor3 | 1.34762783 | 0.00410571 | 0.08971131 |
| ENSMUSG00000022020 | Naa16 | 1.49919758 | 0.00411209 | 0.08977349 |
| ENSMUSG00000028931 | Kcnab2 | -1.5375087 | 0.00413692 | 0.09023809 |
| ENSMUSG00000055567 | Unc80 | 8.25606403 | 0.00415201 | 0.09045029 |
| ENSMUSG00000050534 | Htr5b | -15.043469 | 0.00415377 | 0.09045029 |
| ENSMUSG00000037280 | Galnt6 | -3.1133049 | 0.00417623 | 0.09086163 |

| | | | | |
|---|---|---|---|---|
| ENSMUSG00000031901 | Dus2 | -1.6566904 | 0.0041944 | 0.09117883 |
| ENSMUSG00000040586 | Ofd1 | 1.52304343 | 0.00420536 | 0.09126158 |
| ENSMUSG00000023284 | Zfp605 | 1.67158281 | 0.00420733 | 0.09126158 |
| ENSMUSG00000116725 | Gm29686 | -17.982119 | 0.00420898 | 0.09126158 |
| ENSMUSG00000038151 | Prdm1 | -2.3979689 | 0.0042145 | 0.09130346 |
| ENSMUSG00000036860 | Mrpl55 | -1.4406982 | 0.00422054 | 0.0913564 |
| ENSMUSG00000042190 | Cmklr1 | -1.7334293 | 0.00423979 | 0.09169498 |
| ENSMUSG00000020346 | Mgat1 | -1.4102775 | 0.00425139 | 0.09186769 |
| ENSMUSG00000066607 | Insyn1 | -2.1732142 | 0.00425522 | 0.09187223 |
| ENSMUSG00000038301 | Snx10 | -1.3965579 | 0.00425977 | 0.09189246 |
| ENSMUSG00000073542 | Cep76 | 1.507809 | 0.00429058 | 0.09247847 |
| ENSMUSG00000079620 | Muc4 | 7.36454322 | 0.00429422 | 0.09247861 |
| ENSMUSG00000037643 | Prkci | 1.36331193 | 0.00432991 | 0.09316817 |
| ENSMUSG00000031482 | Slc25a15 | 1.37099727 | 0.00434531 | 0.09342046 |
| ENSMUSG00000005413 | Hmox1 | -2.576098 | 0.00435434 | 0.09353554 |
| ENSMUSG00000038697 | Taf5l | 1.31618743 | 0.00436729 | 0.09373361 |
| ENSMUSG00000001020 | S100a4 | -2.4813063 | 0.00437094 | 0.09373361 |
| ENSMUSG00000034714 | Ttyh2 | -1.486929 | 0.00438661 | 0.09399037 |
| ENSMUSG00000040663 | Clcf1 | -1.6902276 | 0.00439293 | 0.09404638 |
| ENSMUSG00000033970 | Rfc3 | 1.48540524 | 0.00441008 | 0.09433408 |
| ENSMUSG00000034993 | Vat1 | -1.5450196 | 0.0044186 | 0.0944368 |
| ENSMUSG00000041849 | Card6 | -1.6607001 | 0.00442419 | 0.09447649 |
| ENSMUSG00000015653 | Steap2 | -1.9277817 | 0.00443126 | 0.09447649 |
| ENSMUSG00000032407 | U2surp | 1.3159138 | 0.00444101 | 0.09447649 |
| ENSMUSG00000026618 | Iars2 | 1.18984072 | 0.00444116 | 0.09447649 |
| ENSMUSG00000028018 | Gstcd | 1.39935935 | 0.00444337 | 0.09447649 |
| ENSMUSG00000031095 | Cul4b | 1.27255711 | 0.00444346 | 0.09447649 |
| ENSMUSG00000024764 | Naa40 | 1.3310943 | 0.00444648 | 0.09447649 |
| ENSMUSG00000022314 | Rad21 | 1.35799961 | 0.00445316 | 0.09447841 |
| ENSMUSG00000094546 | Ighv1-26 | -46.400052 | 0.004454 | 0.09447841 |
| ENSMUSG00000030589 | Rasgrp4 | -2.771565 | 0.00446549 | 0.09462756 |
| ENSMUSG00000022296 | Baalc | -5.133386 | 0.00446848 | 0.09462756 |
| ENSMUSG00000029321 | Slc10a6 | -4.3149028 | 0.00448175 | 0.09482947 |
| ENSMUSG00000030970 | Ctbp2 | 1.53053832 | 0.00453813 | 0.09587764 |
| ENSMUSG00000010609 | Psen2 | -1.4577064 | 0.00453883 | 0.09587764 |
| ENSMUSG00000035969 | Rusc2 | -1.6960249 | 0.00454312 | 0.09588855 |
| ENSMUSG00000059336 | Slc14a1 | -2.8885107 | 0.00454786 | 0.09590879 |
| ENSMUSG00000055148 | Klf2 | -2.1178004 | 0.00456209 | 0.09607465 |
| ENSMUSG00000009090 | Ap1b1 | -1.2391602 | 0.00456328 | 0.09607465 |
| ENSMUSG00000030929 | Eri2 | 1.41331998 | 0.00458124 | 0.09630548 |
| ENSMUSG00000027424 | Mgme1 | 1.53858575 | 0.0045854 | 0.09630548 |
| ENSMUSG00000028337 | Coro2a | 1.81295759 | 0.00458561 | 0.09630548 |
| ENSMUSG00000030413 | Pglyrp1 | 3.0783647 | 0.00459085 | 0.09633588 |
| ENSMUSG00000000782 | Tcf7 | -2.6559945 | 0.00460076 | 0.09646421 |
| ENSMUSG00000006360 | Crip1 | -1.9335213 | 0.0046078 | 0.09648734 |

| | | | | |
|---|---|---|---|---|
| ENSMUSG00000037553 | Zdhhc18 | -1.2949543 | 0.00460946 | 0.09648734 |
| ENSMUSG00000044716 | Dok7 | -7.2236329 | 0.00463073 | 0.09685274 |
| ENSMUSG00000023067 | Cdkn1a | -2.1266168 | 0.00463719 | 0.09690815 |
| ENSMUSG00000029377 | Ereg | -6.4826102 | 0.0046471 | 0.09703548 |
| ENSMUSG00000040498 | Igsf23 | 29.6413405 | 0.00465792 | 0.09718145 |
| ENSMUSG00000036737 | Oxsr1 | 1.30561221 | 0.00467776 | 0.09745795 |
| ENSMUSG00000032253 | Phip | 1.51826039 | 0.00467884 | 0.09745795 |
| ENSMUSG00000025395 | Prim1 | 1.62032179 | 0.00469832 | 0.09778348 |
| ENSMUSG00000008690 | Ncaph2 | 1.26569625 | 0.00470498 | 0.09784206 |
| ENSMUSG00000071660 | Ttc9c | 1.33475582 | 0.00471629 | 0.09799705 |
| ENSMUSG00000027368 | Dusp2 | -2.3656813 | 0.00473182 | 0.09822991 |
| ENSMUSG00000030208 | Emp1 | -2.0489749 | 0.00473587 | 0.09822991 |
| ENSMUSG00000021638 | Ocln | 1.7013596 | 0.00473909 | 0.09822991 |
| ENSMUSG00000073616 | Cops9 | -1.622395 | 0.00475498 | 0.09847876 |
| ENSMUSG00000032291 | Crabp1 | -7.2331673 | 0.00476451 | 0.0985958 |
| ENSMUSG00000050967 | Creg2 | -2.2691693 | 0.0047841 | 0.09889435 |
| ENSMUSG00000002718 | Cse1l | 1.33625755 | 0.00478672 | 0.09889435 |
| ENSMUSG00000014932 | Yes1 | 1.34954317 | 0.00479617 | 0.09894278 |
| ENSMUSG00000025357 | Dgka | -1.7325446 | 0.00479685 | 0.09894278 |
| ENSMUSG00000024068 | Spast | 1.2443826 | 0.00480311 | 0.09899153 |
| ENSMUSG00000032786 | Alas1 | -1.4317292 | 0.00481847 | 0.09922761 |
| ENSMUSG00000021508 | Cxcl14 | -2.8453785 | 0.00483155 | 0.09941642 |
| ENSMUSG00000032512 | Wdr48 | 1.25463761 | 0.00485109 | 0.09972583 |
| ENSMUSG00000039911 | Spsb1 | -1.786912 | 0.00485443 | 0.09972583 |
| ENSMUSG00000021569 | Trip13 | 1.72175232 | 0.00485926 | 0.09973545 |
| ENSMUSG00000035530 | Eif1 | -1.2563576 | 0.00486275 | 0.09973545 |
| ENSMUSG00000118590 | AC167036.1 | 3.99500532 | 0.00486748 | 0.09975197 |
| ENSMUSG00000036564 | Ndrg4 | -2.015892 | 0.00488 | 0.09992786 |
| ENSMUSG00000048621 | Gm6377 | -2.0912241 | 0.00488779 | 0.10000684 |
| ENSMUSG00000071748 | Gm14698 | 1.75444952 | 0.00489271 | 0.10001549 |
| ENSMUSG00000021094 | Dhrs7 | -1.4342268 | 0.00489608 | 0.10001549 |
| ENSMUSG00000079092 | Prl2c2 | -45.515616 | 0.00490236 | 0.10006323 |
| ENSMUSG00000032332 | Col12a1 | -2.5600493 | 0.00491371 | 0.10021441 |
| ENSMUSG00000001065 | Zfp276 | 1.35747862 | 0.00494761 | 0.10082498 |
| ENSMUSG00000079293 | Clec7a | -2.0610944 | 0.00495584 | 0.1009118 |
| ENSMUSG00000022686 | B3gnt5 | -3.9269819 | 0.00498022 | 0.10129756 |
| ENSMUSG00000082082 | Gm13230 | 7.35266352 | 0.00498807 | 0.10129756 |
| ENSMUSG00000039783 | Kmo | -2.9891571 | 0.00499465 | 0.10129756 |
| ENSMUSG00000033257 | Ttll4 | 1.46057706 | 0.00499696 | 0.10129756 |
| ENSMUSG00000030114 | Klrg1 | -5.7030524 | 0.00500227 | 0.10129756 |
| ENSMUSG00000073705 | Cenps | 1.71626843 | 0.00501216 | 0.10129756 |
| ENSMUSG00000028863 | Meaf6 | 1.27539011 | 0.00501355 | 0.10129756 |
| ENSMUSG00000061544 | Zfp229 | 1.38385002 | 0.00501414 | 0.10129756 |
| ENSMUSG00000051314 | Ffar2 | -3.4267013 | 0.00501442 | 0.10129756 |
| ENSMUSG00000031938 | 4931406C07I | -2.0159772 | 0.00501465 | 0.10129756 |

| | | | | |
|---|---|---|---|---|
| ENSMUSG00000022901 | Cd86 | -1.6750458 | 0.00502614 | 0.10144906 |
| ENSMUSG00000021483 | Cdk20 | -1.4886467 | 0.00503411 | 0.10147281 |
| ENSMUSG00000054715 | Zscan22 | 1.44838825 | 0.00503531 | 0.10147281 |
| ENSMUSG00000031024 | Denn2b | 1.34684872 | 0.00504634 | 0.10161465 |
| ENSMUSG00000044201 | Cdc25c | 1.89583233 | 0.00505515 | 0.1017114 |
| ENSMUSG00000023913 | Pla2g7 | -1.7314849 | 0.0050594 | 0.10171634 |
| ENSMUSG00000019302 | Atp6v0a1 | -1.4112778 | 0.00509035 | 0.10225769 |
| ENSMUSG00000096740 | Lbhd1 | 1.60811042 | 0.00509653 | 0.10225814 |
| ENSMUSG00000027834 | Serpini1 | -3.4007058 | 0.00509862 | 0.10225814 |
| ENSMUSG00000035351 | Nup37 | 1.43292295 | 0.00510244 | 0.10225814 |
| ENSMUSG00000056121 | Fez2 | -1.4147493 | 0.00512784 | 0.10268601 |
| ENSMUSG00000031910 | Has3 | -4.4622175 | 0.0051433 | 0.10286816 |
| ENSMUSG00000075284 | Wipf1 | -1.5137664 | 0.00514736 | 0.10286816 |
| ENSMUSG00000001157 | Gmcl1 | 1.28343409 | 0.00514908 | 0.10286816 |
| ENSMUSG00000058099 | Nfam1 | -1.6795784 | 0.005201 | 0.1038239 |
| ENSMUSG00000034617 | Mtrr | 1.24687115 | 0.00521404 | 0.10400245 |
| ENSMUSG00000031264 | Btk | -1.9025621 | 0.00522267 | 0.10409299 |
| ENSMUSG00000021384 | Susd3 | -2.0553601 | 0.00524404 | 0.10438038 |
| ENSMUSG00000038838 | Vars2 | 1.35563965 | 0.00524531 | 0.10438038 |
| ENSMUSG00000027790 | Sis | 21.683738 | 0.00525267 | 0.10443985 |
| ENSMUSG00000014956 | Ppp1cb | 1.33926232 | 0.00525867 | 0.10443985 |
| ENSMUSG00000036916 | Zfp280c | 1.38168967 | 0.00526063 | 0.10443985 |
| ENSMUSG00000032409 | Atr | 1.4024535 | 0.00528358 | 0.10474918 |
| ENSMUSG00000111546 | Gm47050 | -13.134815 | 0.00528445 | 0.10474918 |
| ENSMUSG00000023992 | Trem2 | -2.9066414 | 0.00532334 | 0.1053579 |
| ENSMUSG00000006273 | Atp6v1b2 | -1.5093438 | 0.00532345 | 0.1053579 |
| ENSMUSG00000048981 | Krt31 | -344.88341 | 0.0053378 | 0.10551839 |
| ENSMUSG00000090290 | Tarbp1 | 1.68683655 | 0.00533987 | 0.10551839 |
| ENSMUSG00000032582 | Rbm6 | 1.25266962 | 0.00535742 | 0.10575689 |
| ENSMUSG00000028484 | Psip1 | 1.35822958 | 0.00536026 | 0.10575689 |
| ENSMUSG00000036745 | Ttll7 | -2.5538064 | 0.00537267 | 0.10591265 |
| ENSMUSG00000001663 | Gstt1 | 3.55663775 | 0.00537649 | 0.10591265 |
| ENSMUSG00000074227 | Spint2 | 1.51032774 | 0.00538648 | 0.10602734 |
| ENSMUSG00000038612 | Mcl1 | -1.214725 | 0.00539882 | 0.10617064 |
| ENSMUSG00000026158 | Ogfrl1 | 2.64223903 | 0.00540612 | 0.10617064 |
| ENSMUSG00000030047 | Arhgap25 | -1.8470385 | 0.0054063 | 0.10617064 |
| ENSMUSG00000047547 | Cltb | -1.5149313 | 0.00542763 | 0.10650719 |
| ENSMUSG00000033213 | AA467197 | -3.7758683 | 0.00547019 | 0.10725966 |
| ENSMUSG00000005465 | Il27ra | -2.2887543 | 0.00548646 | 0.10749573 |
| ENSMUSG00000023345 | Poc1a | 1.55334153 | 0.00549301 | 0.10754115 |
| ENSMUSG00000074024 | 4632427E13I | 2.47720457 | 0.00550826 | 0.1077463 |
| ENSMUSG00000022438 | Parvb | -1.8397467 | 0.00551197 | 0.1077463 |
| ENSMUSG00000002393 | Nr2f6 | 1.25707259 | 0.00559992 | 0.10933807 |
| ENSMUSG00000029204 | Rhoh | -1.9452832 | 0.00560201 | 0.10933807 |
| ENSMUSG00000029246 | Ppat | 1.40282181 | 0.00560828 | 0.10937652 |

| | | | | |
|---|---|---|---|---|
| ENSMUSG00000097073 | 9430037G07 | 4.99432359 | 0.00561399 | 0.10940397 |
| ENSMUSG00000026415 | Fcamr | 36.6152277 | 0.00564818 | 0.10998577 |
| ENSMUSG00000015944 | Castor2 | -1.4977707 | 0.00568393 | 0.1105861 |
| ENSMUSG00000115483 | Gm9732 | 8.05521091 | 0.00569336 | 0.1105861 |
| ENSMUSG00000038642 | Ctss | -1.765995 | 0.00569581 | 0.1105861 |
| ENSMUSG00000030579 | Tyrobp | -1.6776131 | 0.00569662 | 0.1105861 |
| ENSMUSG00000069607 | Cd300ld3 | -2.1588527 | 0.00570076 | 0.1105861 |
| ENSMUSG00000033166 | Dis3 | 1.48248736 | 0.00571119 | 0.11070393 |
| ENSMUSG00000031353 | Rbbp7 | 1.31108063 | 0.00572762 | 0.11093767 |
| ENSMUSG00000000031 | H19 | -5.9521295 | 0.00573436 | 0.11098362 |
| ENSMUSG00000033233 | Trim45 | 1.51486036 | 0.0057426 | 0.11105859 |
| ENSMUSG00000038811 | Gngt2 | -1.9174606 | 0.00578239 | 0.11174305 |
| ENSMUSG00000026796 | Fam129b | -1.3025814 | 0.00578921 | 0.1117898 |
| ENSMUSG00000031207 | Msn | -1.5300828 | 0.00580764 | 0.11194088 |
| ENSMUSG00000086942 | Gm15489 | 24.2703319 | 0.00580867 | 0.11194088 |
| ENSMUSG00000029735 | Tpk1 | 1.38765234 | 0.00581085 | 0.11194088 |
| ENSMUSG00000118633 | CT868690.1 | -6.5510865 | 0.00581667 | 0.11194088 |
| ENSMUSG00000037325 | Bbs7 | 1.47339804 | 0.00581906 | 0.11194088 |
| ENSMUSG00000036636 | Clcn7 | -1.3769574 | 0.00582402 | 0.11195168 |
| ENSMUSG00000102543 | Pcdhgc5 | -2.4453738 | 0.00583269 | 0.11196269 |
| ENSMUSG00000029730 | Mcm7 | 1.47750266 | 0.00583341 | 0.11196269 |
| ENSMUSG00000100954 | Gm10138 | 2.11142632 | 0.00588684 | 0.11290302 |
| ENSMUSG00000066037 | Hnrnpr | 1.30684363 | 0.00590985 | 0.11320573 |
| ENSMUSG00000036553 | Sh3tc1 | -1.6740896 | 0.00591154 | 0.11320573 |
| ENSMUSG00000024987 | Cyp26a1 | -5.2202746 | 0.00594118 | 0.11363284 |
| ENSMUSG00000056656 | Apol8 | -13.464327 | 0.00594278 | 0.11363284 |
| ENSMUSG00000110411 | Gm45457 | 4.14809457 | 0.0059479 | 0.11364219 |
| ENSMUSG00000033066 | Gas7 | -2.2055667 | 0.00596174 | 0.11364219 |
| ENSMUSG00000041852 | Tcf20 | 1.42983038 | 0.00596684 | 0.11364219 |
| ENSMUSG00000049103 | Ccr2 | -2.186822 | 0.00596698 | 0.11364219 |
| ENSMUSG00000003559 | As3mt | -1.4120096 | 0.00597133 | 0.11364219 |
| ENSMUSG00000007872 | Id3 | -2.6556454 | 0.00597186 | 0.11364219 |
| ENSMUSG00000022466 | Rpap3 | 1.25761982 | 0.00597458 | 0.11364219 |
| ENSMUSG00000004642 | Slbp | 1.37779277 | 0.00598936 | 0.11383817 |
| ENSMUSG00000032667 | Pon2 | -1.4639577 | 0.00601441 | 0.11422883 |
| ENSMUSG00000042608 | Stk40 | 1.46004275 | 0.00604003 | 0.11460404 |
| ENSMUSG00000069135 | Fgfr1op | 1.35979689 | 0.00604318 | 0.11460404 |
| ENSMUSG00000021367 | Edn1 | 2.34402083 | 0.00607117 | 0.11504884 |
| ENSMUSG00000028096 | Gpr89 | 1.3135891 | 0.00609315 | 0.11537932 |
| ENSMUSG00000044703 | Phf11a | -2.549023 | 0.00611573 | 0.1155776 |
| ENSMUSG00000015165 | Hnrnpl | 1.18607446 | 0.00611785 | 0.1155776 |
| ENSMUSG00000010307 | Tmem86a | -1.8152471 | 0.0061189 | 0.1155776 |
| ENSMUSG00000035376 | Hacd2 | 1.47710241 | 0.00612181 | 0.1155776 |
| ENSMUSG00000011382 | Dhdh | -1.8718313 | 0.00613348 | 0.11571194 |
| ENSMUSG00000051278 | Zgrf1 | 1.80577678 | 0.00614126 | 0.11577284 |

| | | | | |
|---|---|---|---|---|
| ENSMUSG00000020868 | Xylt2 | -1.3275119 | 0.00616281 | 0.11609282 |
| ENSMUSG00000001657 | Hoxc8 | -3.6118769 | 0.00616935 | 0.11613 |
| ENSMUSG00000051768 | Xrcc1 | 1.330715 | 0.0061777 | 0.11620107 |
| ENSMUSG00000018927 | Ccl6 | -2.2959659 | 0.00618319 | 0.11621837 |
| ENSMUSG00000040364 | Sec1 | 9.72518879 | 0.00619927 | 0.11643443 |
| ENSMUSG00000024251 | Thada | 1.39492886 | 0.00621102 | 0.11656892 |
| ENSMUSG00000022449 | Adamts20 | 3.43148038 | 0.00621952 | 0.11658031 |
| ENSMUSG00000027722 | Spata5 | 1.31626582 | 0.0062208 | 0.11658031 |
| ENSMUSG00000052102 | Gnpda1 | -1.270009 | 0.0062367 | 0.11679226 |
| ENSMUSG00000029427 | Zcchc8 | 1.28147489 | 0.00624254 | 0.11681545 |
| ENSMUSG00000000759 | Tubgcp3 | 1.32659934 | 0.00628625 | 0.11754458 |
| ENSMUSG00000012017 | Scarf2 | -1.5779142 | 0.00629518 | 0.11754458 |
| ENSMUSG00000039630 | Hnrnpu | 1.2228928 | 0.00629538 | 0.11754458 |
| ENSMUSG00000069227 | Gprin1 | 2.34971463 | 0.00630826 | 0.11769853 |
| ENSMUSG00000055805 | Fmnl1 | -1.5864883 | 0.00633231 | 0.11806071 |
| ENSMUSG00000027668 | Mfn1 | 1.24293613 | 0.00634316 | 0.11817631 |
| ENSMUSG00000095609 | Gm21188 | -2.1791021 | 0.00634924 | 0.11820289 |
| ENSMUSG00000097180 | 2700038G22 | 2.33232869 | 0.00635581 | 0.11823849 |
| ENSMUSG00000048170 | Mcmbp | 1.24970164 | 0.00637241 | 0.11839918 |
| ENSMUSG00000029161 | Cgref1 | -2.2967753 | 0.0063766 | 0.11839918 |
| ENSMUSG00000028657 | Ppt1 | -1.2449631 | 0.00637842 | 0.11839918 |
| ENSMUSG00000054555 | Adam12 | -2.1551557 | 0.00639156 | 0.11855648 |
| ENSMUSG00000032118 | Fez1 | -5.4158732 | 0.00640612 | 0.11866079 |
| ENSMUSG00000078202 | Nrarp | -1.6722464 | 0.00640652 | 0.11866079 |
| ENSMUSG00000022325 | Pop1 | 1.47161491 | 0.00641264 | 0.11868746 |
| ENSMUSG00000075470 | Alg10b | 1.31108604 | 0.00642011 | 0.11873935 |
| ENSMUSG00000105954 | Gm42793 | -10.642122 | 0.00642898 | 0.11881694 |
| ENSMUSG00000022391 | Rangap1 | 1.29155125 | 0.00644437 | 0.11901477 |
| ENSMUSG00000036908 | Unc93b1 | -1.3541169 | 0.00646347 | 0.11921671 |
| ENSMUSG00000022575 | Gsdmd | -1.6260811 | 0.00647081 | 0.11921671 |
| ENSMUSG00000034595 | Ppp1r18 | -1.4947898 | 0.00647316 | 0.11921671 |
| ENSMUSG00000043262 | Uevld | 1.37533007 | 0.00647407 | 0.11921671 |
| ENSMUSG00000024067 | Dpy30 | 1.25761884 | 0.00649158 | 0.11945258 |
| ENSMUSG00000108389 | Gm45206 | 6.58591707 | 0.00650142 | 0.11954714 |
| ENSMUSG00000028127 | Abcd3 | 1.31944696 | 0.00651911 | 0.11978577 |
| ENSMUSG00000048047 | Zbtb33 | 1.3346056 | 0.00653406 | 0.11995575 |
| ENSMUSG00000034796 | Cpne7 | 5.23404752 | 0.00653781 | 0.11995575 |
| ENSMUSG00000020993 | Trappc6b | -1.2784191 | 0.00655027 | 0.12009783 |
| ENSMUSG00000052160 | Pld4 | -1.6976828 | 0.00658214 | 0.12059503 |
| ENSMUSG00000002885 | Adgre5 | -1.6165525 | 0.0065909 | 0.1206686 |
| ENSMUSG00000041112 | Elmo1 | -1.5753117 | 0.00660817 | 0.12089757 |
| ENSMUSG00000024962 | Vegfb | 1.50641411 | 0.00661628 | 0.12095893 |
| ENSMUSG00000036905 | C1qb | -1.63521 | 0.00662483 | 0.12100703 |
| ENSMUSG00000030816 | Rnf40 | 1.21434644 | 0.00662843 | 0.12100703 |
| ENSMUSG00000024442 | Dele1 | 1.28168958 | 0.00665095 | 0.12127475 |

| | | | | |
|---|---|---|---|---|
| ENSMUSG00000075702 | Selenom | -1.7548213 | 0.00665264 | 0.12127475 |
| ENSMUSG00000066361 | Serpina3c | -7.0395447 | 0.00665846 | 0.12129382 |
| ENSMUSG00000030610 | Det1 | 1.35318413 | 0.00667684 | 0.12154138 |
| ENSMUSG00000005836 | Gata6 | -6.7560759 | 0.00670995 | 0.12205677 |
| ENSMUSG00000026880 | Stom | -1.7599498 | 0.00672491 | 0.12217657 |
| ENSMUSG00000038417 | Fig4 | -1.243583 | 0.00672666 | 0.12217657 |
| ENSMUSG00000048677 | Tpcn2 | -1.3624399 | 0.00673182 | 0.12217657 |
| ENSMUSG00000036943 | Rab8b | -1.4815317 | 0.00673577 | 0.12217657 |
| ENSMUSG00000022425 | Enpp2 | -4.8195313 | 0.00674762 | 0.12225946 |
| ENSMUSG00000068417 | Pnp2 | -2.2167722 | 0.00675208 | 0.12225946 |
| ENSMUSG00000024513 | Mbd2 | -1.2686592 | 0.00675477 | 0.12225946 |
| ENSMUSG00000079588 | Tmem182 | -4.3287701 | 0.0067807 | 0.12264142 |
| ENSMUSG00000033491 | Prss35 | -7.0133545 | 0.00679091 | 0.12273873 |
| ENSMUSG00000015134 | Aldh1a3 | -2.8928303 | 0.00679585 | 0.12274072 |
| ENSMUSG00000036894 | Rap2b | -1.316541 | 0.00680259 | 0.12274571 |
| ENSMUSG00000057069 | Ero1lb | -1.2845681 | 0.00680579 | 0.12274571 |
| ENSMUSG00000031125 | 3830403N18 | -15.664546 | 0.00683209 | 0.12308828 |
| ENSMUSG00000033520 | Idi2 | -183.18477 | 0.00683447 | 0.12308828 |
| ENSMUSG00000029819 | Npy | -89.555111 | 0.00684524 | 0.12311995 |
| ENSMUSG00000013846 | St3gal1 | -1.5130942 | 0.00684592 | 0.12311995 |
| ENSMUSG00000026491 | Ahctf1 | 1.36249508 | 0.00686433 | 0.12336376 |
| ENSMUSG00000023391 | Dlx2 | -27.493047 | 0.00688211 | 0.12359596 |
| ENSMUSG00000029672 | Fam3c | -1.5049503 | 0.00689381 | 0.12371858 |
| ENSMUSG00000039115 | Itga9 | -1.8504413 | 0.00690887 | 0.12390128 |
| ENSMUSG00000063334 | Krr1 | 1.23346991 | 0.00691952 | 0.12395875 |
| ENSMUSG00000025451 | Paip1 | 1.16063925 | 0.0069246 | 0.12395875 |
| ENSMUSG00000076609 | Igkc | -13.710314 | 0.0069267 | 0.12395875 |
| ENSMUSG00000024300 | Myo1f | -1.551902 | 0.00693431 | 0.12400754 |
| ENSMUSG00000037049 | Smpd1 | -1.4644242 | 0.00693973 | 0.12401716 |
| ENSMUSG00000020612 | Prkar1a | -1.1693055 | 0.00697732 | 0.12458103 |
| ENSMUSG00000022816 | Fstl1 | -1.8796707 | 0.00698109 | 0.12458103 |
| ENSMUSG00000046541 | Zfp526 | 1.36891653 | 0.00699826 | 0.12479984 |
| ENSMUSG00000036606 | Plxnb2 | 1.23069765 | 0.00700536 | 0.12483887 |
| ENSMUSG00000001510 | Dlx3 | -9.8558596 | 0.00701546 | 0.12486242 |
| ENSMUSG00000020121 | Srgap1 | -2.0513361 | 0.00701651 | 0.12486242 |
| ENSMUSG00000029029 | Wrap73 | 1.32494038 | 0.00703386 | 0.12508351 |
| ENSMUSG00000063856 | Gpx1 | -1.3997526 | 0.00704331 | 0.12512408 |
| ENSMUSG00000029110 | Rnf4 | 1.17231748 | 0.00704598 | 0.12512408 |
| ENSMUSG00000025283 | Sat1 | -1.6048579 | 0.00707196 | 0.12549763 |
| ENSMUSG00000028034 | Fubp1 | 1.55570921 | 0.00707701 | 0.1254996 |
| ENSMUSG00000030641 | Ddias | 1.68430815 | 0.00709249 | 0.12568643 |
| ENSMUSG00000020709 | Adap2 | -1.6182698 | 0.00711152 | 0.12593586 |
| ENSMUSG00000011751 | Sptbn4 | 2.8310132 | 0.00712446 | 0.12607709 |
| ENSMUSG00000047641 | Krt87 | -117.2697 | 0.00719232 | 0.12718953 |
| ENSMUSG00000045551 | Fpr1 | -3.0044586 | 0.00720905 | 0.12731119 |

| | | | | |
|---|---|---|---|---|
| ENSMUSG00000003420 | Fcgrt | -1.557028 | 0.00720922 | 0.12731119 |
| ENSMUSG00000034867 | Ankrd27 | 1.19686907 | 0.00722979 | 0.12758576 |
| ENSMUSG00000037017 | Zscan21 | 1.32089392 | 0.00732282 | 0.12913772 |
| ENSMUSG00000005142 | Man2b1 | -1.3717693 | 0.00733143 | 0.12919994 |
| ENSMUSG00000063651 | Cnfn | -52.579261 | 0.00734745 | 0.12928593 |
| ENSMUSG00000028755 | Cda | 1.8841318 | 0.00735024 | 0.12928593 |
| ENSMUSG00000026283 | Ing5 | 1.38418987 | 0.00735305 | 0.12928593 |
| ENSMUSG00000043939 | A530064D06 | -3.4728348 | 0.00735801 | 0.12928593 |
| ENSMUSG00000020290 | Xpo1 | 1.25580833 | 0.00736175 | 0.12928593 |
| ENSMUSG00000032403 | 2300009A05 | 1.38188841 | 0.00737368 | 0.12940597 |
| ENSMUSG00000036968 | Cnpy4 | -1.3363856 | 0.00739529 | 0.12961559 |
| ENSMUSG00000014030 | Pax5 | -5.1229143 | 0.00739582 | 0.12961559 |
| ENSMUSG00000016984 | Etaa1 | 1.36842494 | 0.00740253 | 0.12964369 |
| ENSMUSG00000028373 | Astn2 | -6.2989056 | 0.00742641 | 0.12993204 |
| ENSMUSG00000056167 | Cnot10 | 1.32925317 | 0.00742922 | 0.12993204 |
| ENSMUSG00000030148 | Clec4a2 | -1.8984456 | 0.00743997 | 0.13003069 |
| ENSMUSG00000049764 | Zfp280b | 1.61459315 | 0.00746377 | 0.13035695 |
| ENSMUSG00000005682 | Pan2 | 1.60563249 | 0.00746993 | 0.13037494 |
| ENSMUSG00000027669 | Gnb4 | -1.4628465 | 0.00747708 | 0.13037576 |
| ENSMUSG00000042590 | Ipo11 | 1.28879087 | 0.00748024 | 0.13037576 |
| ENSMUSG00000022221 | Ripk3 | -2.0199358 | 0.00748896 | 0.13043831 |
| ENSMUSG00000022665 | Ccdc80 | -2.9947629 | 0.0075105 | 0.13072378 |
| ENSMUSG00000029822 | Osbpl3 | 1.79267098 | 0.00752823 | 0.13094276 |
| ENSMUSG00000046111 | Cep295 | 1.43527836 | 0.00755127 | 0.1312536 |
| ENSMUSG00000026669 | Mcm10 | 1.59868747 | 0.00756629 | 0.13142485 |
| ENSMUSG00000026843 | Fubp3 | 1.20350202 | 0.00758873 | 0.13165734 |
| ENSMUSG00000024529 | Lox | -2.6222809 | 0.00759004 | 0.13165734 |
| ENSMUSG00000033860 | Fgg | -15.751845 | 0.00762769 | 0.13217069 |
| ENSMUSG00000063015 | Ccni | 1.27072848 | 0.00763003 | 0.13217069 |
| ENSMUSG00000055172 | C1ra | -1.7130158 | 0.00763526 | 0.13217114 |
| ENSMUSG00000011267 | Zfp296 | 1.63661798 | 0.00766955 | 0.13259561 |
| ENSMUSG00000022534 | Mefv | -3.4589403 | 0.00767022 | 0.13259561 |
| ENSMUSG00000028405 | Aco1 | 1.30726304 | 0.00768084 | 0.13268901 |
| ENSMUSG00000022148 | Fyb | -1.5241498 | 0.00770408 | 0.13289329 |
| ENSMUSG00000113536 | Gm49327 | 2.04333228 | 0.00770777 | 0.13289329 |
| ENSMUSG00000015023 | Ddx19a | 1.2839443 | 0.00771133 | 0.13289329 |
| ENSMUSG00000020091 | Eif4ebp2 | 1.20797886 | 0.00771358 | 0.13289329 |
| ENSMUSG00000031595 | Pdgfrl | -2.4969424 | 0.00772097 | 0.13293044 |
| ENSMUSG00000021611 | Tert | 1.83929985 | 0.0077393 | 0.13298555 |
| ENSMUSG00000076937 | Iglc2 | -19.898833 | 0.00773958 | 0.13298555 |
| ENSMUSG00000031626 | Sorbs2 | 1.75723437 | 0.00773987 | 0.13298555 |
| ENSMUSG00000031826 | Usp10 | 1.28276545 | 0.00774656 | 0.13301052 |
| ENSMUSG00000103037 | Pcdhgb1 | -2.688493 | 0.00779828 | 0.13380817 |
| ENSMUSG00000029661 | Col1a2 | -2.7078567 | 0.00780696 | 0.13381061 |
| ENSMUSG00000024404 | Riok3 | -1.2549163 | 0.00780895 | 0.13381061 |

| | | | | |
|---|---|---|---|---|
| ENSMUSG00000019876 | Pkib | -1.7556496 | 0.00785583 | 0.13452314 |
| ENSMUSG00000007050 | Lsm2 | 1.49781256 | 0.00787896 | 0.13482829 |
| ENSMUSG00000000731 | Aire | 5.7587367 | 0.0078872 | 0.13487857 |
| ENSMUSG00000040327 | Cul9 | -1.5564728 | 0.00789297 | 0.1348865 |
| ENSMUSG00000038816 | Ctnnal1 | 1.47402977 | 0.00790644 | 0.13494115 |
| ENSMUSG00000090031 | 4732440D04 | 1.92673292 | 0.00790679 | 0.13494115 |
| ENSMUSG00000058427 | Cxcl2 | -4.8855028 | 0.00791762 | 0.13501864 |
| ENSMUSG00000073412 | Lst1 | -1.7323908 | 0.00792196 | 0.13501864 |
| ENSMUSG00000071181 | 3830408C21I | 3.3408337 | 0.00793612 | 0.13516932 |
| ENSMUSG00000027102 | Hoxd8 | -2.0262375 | 0.00794882 | 0.1352949 |
| ENSMUSG00000056498 | Tmem154 | -1.8123115 | 0.00797743 | 0.13569115 |
| ENSMUSG00000065987 | Cd209b | -7.0830248 | 0.00799184 | 0.13584523 |
| ENSMUSG00000037913 | Tmem156 | -2.4374728 | 0.00800154 | 0.13590358 |
| ENSMUSG00000025764 | Jade1 | 1.40713296 | 0.0080154 | 0.13590358 |
| ENSMUSG00000047562 | Mmp10 | -19.952263 | 0.00801635 | 0.13590358 |
| ENSMUSG00000086015 | 4833417C18I | 4.73156245 | 0.00802029 | 0.13590358 |
| ENSMUSG00000043144 | Aqp6 | 86.7877294 | 0.00802201 | 0.13590358 |
| ENSMUSG00000042292 | Mrtfa | 1.25666403 | 0.00802975 | 0.13594406 |
| ENSMUSG00000029655 | N4bp2l2 | 1.24369674 | 0.0080474 | 0.13611187 |
| ENSMUSG00000056427 | Slit3 | -2.7408451 | 0.00805038 | 0.13611187 |
| ENSMUSG00000015748 | Prpf3 | 1.41154996 | 0.00806754 | 0.1361718 |
| ENSMUSG00000008604 | Ubqln4 | 1.35250199 | 0.00807161 | 0.1361718 |
| ENSMUSG00000051855 | Mest | -3.715578 | 0.00807164 | 0.1361718 |
| ENSMUSG00000045838 | Ccdc9b | -1.5442218 | 0.00807594 | 0.1361718 |
| ENSMUSG00000027173 | Depdc7 | -2.3332778 | 0.00808071 | 0.1361718 |
| ENSMUSG00000024812 | Tjp2 | 1.39321603 | 0.00808985 | 0.13623548 |
| ENSMUSG00000044165 | Bcl2l15 | 5.30337207 | 0.00811102 | 0.13644897 |
| ENSMUSG00000030894 | Tpp1 | -1.239355 | 0.00811744 | 0.13644897 |
| ENSMUSG00000030468 | Siglecg | -4.9813677 | 0.00812593 | 0.13644897 |
| ENSMUSG00000021178 | Psmc1 | -1.256628 | 0.0081283 | 0.13644897 |
| ENSMUSG00000045045 | Lrfn4 | 1.47458921 | 0.00812938 | 0.13644897 |
| ENSMUSG00000031737 | Irx5 | 1.44574751 | 0.00816256 | 0.1369155 |
| ENSMUSG00000066000 | Zfp979 | -3.4047505 | 0.00817902 | 0.1371011 |
| ENSMUSG00000112808 | Gm4739 | 2.05577931 | 0.00818731 | 0.1371496 |
| ENSMUSG00000036109 | Mbnl3 | 1.74098429 | 0.008221 | 0.13762316 |
| ENSMUSG00000042918 | Mamstr | 4.5938405 | 0.0082287 | 0.13766147 |
| ENSMUSG00000033770 | Clcnka | 6.16350774 | 0.00823823 | 0.13773012 |
| ENSMUSG00000021702 | Thbs4 | -2.7896098 | 0.00824544 | 0.13775158 |
| ENSMUSG00000027534 | Snx16 | 1.34666876 | 0.00825035 | 0.13775158 |
| ENSMUSG00000027905 | Ddx20 | 1.23350398 | 0.00825734 | 0.13777775 |
| ENSMUSG00000034205 | Loxl2 | -2.2631603 | 0.00828403 | 0.13813246 |
| ENSMUSG00000032011 | Thy1 | -2.0972742 | 0.00831049 | 0.1384675 |
| ENSMUSG00000043460 | Elfn2 | -12.401389 | 0.00831502 | 0.1384675 |
| ENSMUSG00000118607 | AC147806.2 | 2.48908473 | 0.0083553 | 0.13904702 |
| ENSMUSG00000027968 | Larp7 | 1.32714429 | 0.00836476 | 0.13911333 |

| | | | | |
|---|---|---|---|---|
| ENSMUSG00000068373 | D430041D05 | -6.9323905 | 0.00838587 | 0.13937334 |
| ENSMUSG00000038692 | Hoxb4 | -1.7913867 | 0.0084214 | 0.13978929 |
| ENSMUSG00000110790 | Gm47079 | 22.7290188 | 0.0084219 | 0.13978929 |
| ENSMUSG00000027848 | Olfml3 | -1.51194 | 0.00842839 | 0.13980562 |
| ENSMUSG00000036887 | C1qa | -1.629073 | 0.00845527 | 0.14016 |
| ENSMUSG00000026581 | Sell | -4.0101426 | 0.00846184 | 0.14017753 |
| ENSMUSG00000019986 | Ahi1 | -1.6009422 | 0.0084727 | 0.14026591 |
| ENSMUSG00000024978 | Gpam | -1.5961365 | 0.00848155 | 0.1403211 |
| ENSMUSG00000030095 | Tmem43 | -1.4513235 | 0.00849148 | 0.14039399 |
| ENSMUSG00000028743 | Akr7a5 | 1.36319056 | 0.00850305 | 0.14048746 |
| ENSMUSG00000040447 | Spns2 | 1.91934394 | 0.00850819 | 0.14048746 |
| ENSMUSG00000041859 | Mcm3 | 1.44584209 | 0.00851435 | 0.14049788 |
| ENSMUSG00000079012 | Serpina3m | -26.140505 | 0.00852597 | 0.14059822 |
| ENSMUSG00000016524 | Il19 | -18.074122 | 0.00860106 | 0.14174466 |
| ENSMUSG00000049823 | Zbtb12 | 1.47080656 | 0.00862225 | 0.14195225 |
| ENSMUSG00000026817 | Ak1 | -1.7301316 | 0.00862483 | 0.14195225 |
| ENSMUSG00000110216 | Gm36325 | 6.78504558 | 0.00863473 | 0.14197019 |
| ENSMUSG00000098188 | Sowahc | -1.4400949 | 0.0086371 | 0.14197019 |
| ENSMUSG00000045665 | Mfsd5 | -1.2383047 | 0.00865641 | 0.14210461 |
| ENSMUSG00000079018 | Ly6c1 | -1.9202703 | 0.00865646 | 0.14210461 |
| ENSMUSG00000028826 | Maco1 | 1.2341278 | 0.00866405 | 0.14210975 |
| ENSMUSG00000050195 | Scd4 | 9.05726259 | 0.0086707 | 0.14210975 |
| ENSMUSG00000097392 | Thoc2l | 1.62284515 | 0.00867355 | 0.14210975 |
| ENSMUSG00000032178 | Ilf3 | 1.23656498 | 0.00868848 | 0.14226198 |
| ENSMUSG00000036208 | Nepro | 1.32939109 | 0.00869404 | 0.14226198 |
| ENSMUSG00000020871 | Dlx4 | -2.8507879 | 0.00872059 | 0.14256126 |
| ENSMUSG00000000544 | Gpa33 | 6.70535014 | 0.00872355 | 0.14256126 |
| ENSMUSG00000031283 | Chrdl1 | -3.4195583 | 0.008731 | 0.14259131 |
| ENSMUSG00000024245 | Tmem178 | -10.376388 | 0.00875024 | 0.14281375 |
| ENSMUSG00000025269 | Apex2 | 1.49002855 | 0.00876261 | 0.1429239 |
| ENSMUSG00000052296 | Ppp6r1 | 1.17859874 | 0.00877515 | 0.14297001 |
| ENSMUSG00000058297 | Spock2 | -2.5748428 | 0.00878036 | 0.14297001 |
| ENSMUSG00000024037 | Wdr4 | 1.27103376 | 0.00878231 | 0.14297001 |
| ENSMUSG00000087651 | 1500009L16F | -2.7522693 | 0.00880153 | 0.14319107 |
| ENSMUSG00000105388 | Rpl36a-ps2 | 2.06007181 | 0.00882949 | 0.14355404 |
| ENSMUSG00000038172 | Ttc39b | -1.6702174 | 0.00886745 | 0.14397144 |
| ENSMUSG00000021710 | Nln | 1.23723672 | 0.00887114 | 0.14397144 |
| ENSMUSG00000033577 | Myo6 | 1.53554544 | 0.00887614 | 0.14397144 |
| ENSMUSG00000030498 | Gas2 | 1.63356106 | 0.00887782 | 0.14397144 |
| ENSMUSG00000074677 | Sirpb1c | -2.0413932 | 0.00890045 | 0.14424639 |
| ENSMUSG00000021798 | Ldb3 | 2.96939179 | 0.0089132 | 0.14433806 |
| ENSMUSG00000022471 | Xrcc6 | 1.2319623 | 0.00891747 | 0.14433806 |
| ENSMUSG00000042988 | Notum | -100.08425 | 0.00892956 | 0.14441573 |
| ENSMUSG00000047044 | D030056L22I | 1.39919271 | 0.00893364 | 0.14441573 |
| ENSMUSG00000060063 | Alox5ap | -1.9400296 | 0.00897596 | 0.14500762 |

| | | | | |
|---|---|---|---|---|
| ENSMUSG00000054640 | Slc8a1 | -1.7937268 | 0.00901323 | 0.14551727 |
| ENSMUSG00000021707 | Dhfr | 1.52143088 | 0.00902604 | 0.14563163 |
| ENSMUSG00000026640 | Plxna2 | -1.4836448 | 0.00904987 | 0.14589091 |
| ENSMUSG00000032047 | Acat1 | 1.33611404 | 0.0090536 | 0.14589091 |
| ENSMUSG00000042363 | Lgalsl | -1.3893704 | 0.00906288 | 0.14594791 |
| ENSMUSG00000020492 | Ska2 | 1.54876869 | 0.00908145 | 0.14605569 |
| ENSMUSG00000033991 | Ttc37 | 1.31203831 | 0.00908679 | 0.14605569 |
| ENSMUSG00000030111 | A2m | -7.6636963 | 0.00909226 | 0.14605569 |
| ENSMUSG00000025555 | Farp1 | 1.38184182 | 0.00909373 | 0.14605569 |
| ENSMUSG00000063804 | Lin28b | 15.7698179 | 0.00909831 | 0.14605569 |
| ENSMUSG00000007682 | Dio2 | -4.3304554 | 0.00910838 | 0.14612505 |
| ENSMUSG00000039982 | Dtx4 | 1.33713183 | 0.00913651 | 0.14637935 |
| ENSMUSG00000060508 | Nlrp9b | 8.84894804 | 0.00914017 | 0.14637935 |
| ENSMUSG00000062082 | Cd200r4 | -2.5776754 | 0.00914423 | 0.14637935 |
| ENSMUSG00000046667 | Rbm12b1 | 1.45608086 | 0.00914754 | 0.14637935 |
| ENSMUSG00000075122 | Cd80 | -2.2745419 | 0.00915948 | 0.14637935 |
| ENSMUSG00000023348 | Trip6 | 1.49066681 | 0.00916039 | 0.14637935 |
| ENSMUSG00000026032 | Ndufb3 | -1.2742721 | 0.00916471 | 0.14637935 |
| ENSMUSG00000039981 | Zc3h12d | -1.8852381 | 0.00917031 | 0.14637935 |
| ENSMUSG00000028455 | Stoml2 | 1.21955116 | 0.00921184 | 0.1469499 |
| ENSMUSG00000079003 | Samd1 | 1.47317871 | 0.0092319 | 0.14712978 |
| ENSMUSG00000033721 | Vav3 | -1.6421567 | 0.00923469 | 0.14712978 |
| ENSMUSG00000029283 | Cdc7 | 2.46443812 | 0.00926028 | 0.14744496 |
| ENSMUSG00000018930 | Ccl4 | -2.554369 | 0.00926967 | 0.14750208 |
| ENSMUSG00000050350 | Gpr18 | -2.2863659 | 0.00928244 | 0.14754617 |
| ENSMUSG00000042105 | Inpp5f | -1.4059176 | 0.00928405 | 0.14754617 |
| ENSMUSG00000002983 | Relb | 1.42086957 | 0.00929219 | 0.14758316 |
| ENSMUSG00000035131 | Brinp3 | -10.00532 | 0.00931153 | 0.14772835 |
| ENSMUSG00000031725 | Ces1f | -8.5381881 | 0.00931295 | 0.14772835 |
| ENSMUSG00000037253 | Mex3c | 1.25155941 | 0.00933151 | 0.1479304 |
| ENSMUSG00000049744 | Arhgap15 | -1.8141858 | 0.00933808 | 0.14794224 |
| ENSMUSG00000029154 | Cwh43 | -8.3036925 | 0.00934497 | 0.14795906 |
| ENSMUSG00000037185 | Krt80 | -3.4736113 | 0.00935364 | 0.1479937 |
| ENSMUSG00000004637 | Wwox | 1.50905341 | 0.0093588 | 0.1479937 |
| ENSMUSG00000036949 | Slc39a12 | -21.888537 | 0.00937232 | 0.14811321 |
| ENSMUSG00000005339 | Fcer1a | -7.3691201 | 0.00937856 | 0.14811321 |
| ENSMUSG00000027298 | Tyro3 | 1.63300588 | 0.00938769 | 0.14811321 |
| ENSMUSG00000039205 | Ciz1 | 1.28365116 | 0.00938967 | 0.14811321 |
| ENSMUSG00000035024 | Ncapd3 | 1.45056891 | 0.00940568 | 0.14827372 |
| ENSMUSG00000070003 | Ssbp4 | -1.3799634 | 0.00942483 | 0.14848342 |
| ENSMUSG00000022871 | Fetub | -4.5968585 | 0.00944938 | 0.14877799 |
| ENSMUSG00000005813 | Metap1 | 1.24351585 | 0.00946266 | 0.14888591 |
| ENSMUSG00000021624 | Cd180 | -1.5946916 | 0.00946795 | 0.14888591 |
| ENSMUSG00000074071 | Fam169b | -3.2697558 | 0.00947836 | 0.14895732 |
| ENSMUSG00000027224 | Duoxa1 | -5.4673773 | 0.00948874 | 0.14902832 |

| | | | | |
|---|---|---|---|---|
| ENSMUSG00000039634 | Zfp189 | 1.57915613 | 0.0095086 | 0.14924801 |
| ENSMUSG00000031506 | Ptpn7 | -2.1116198 | 0.00952799 | 0.14946002 |
| ENSMUSG00000056211 | R3hdm1 | 1.29711183 | 0.00953837 | 0.1495306 |
| ENSMUSG00000010660 | Plcd1 | -1.6704981 | 0.00955721 | 0.14973354 |
| ENSMUSG00000048142 | Nat8l | -8.5915709 | 0.0095863 | 0.15009682 |
| ENSMUSG00000026043 | Col3a1 | -2.3743265 | 0.00960338 | 0.15027165 |
| ENSMUSG00000044791 | Setd2 | 1.30523849 | 0.00961248 | 0.15032139 |
| ENSMUSG00000020473 | Aebp1 | -1.9917758 | 0.00966715 | 0.15092831 |
| ENSMUSG00000014074 | Rnf168 | 1.217457 | 0.00968215 | 0.15092831 |
| ENSMUSG00000110751 | C230053D17 | 12.6826677 | 0.00968369 | 0.15092831 |
| ENSMUSG00000002459 | Rgs20 | 2.1092233 | 0.00968743 | 0.15092831 |
| ENSMUSG00000026288 | Inpp5d | -1.4712377 | 0.00968887 | 0.15092831 |
| ENSMUSG00000025894 | Aasdhppt | 1.22105321 | 0.00969664 | 0.15092831 |
| ENSMUSG00000023191 | P3h3 | -1.5162647 | 0.00970399 | 0.15092831 |
| ENSMUSG00000030880 | Polr3e | 1.26299576 | 0.00970964 | 0.15092831 |
| ENSMUSG00000026473 | Glul | -2.0695755 | 0.00971487 | 0.15092831 |
| ENSMUSG00000010205 | Raver1 | 1.2926043 | 0.00971598 | 0.15092831 |
| ENSMUSG00000044162 | Tnip3 | -2.8522458 | 0.00971662 | 0.15092831 |
| ENSMUSG00000103276 | Gm37917 | -3.2458409 | 0.00974821 | 0.15126915 |
| ENSMUSG00000035448 | Ccr3 | -4.6925307 | 0.00975047 | 0.15126915 |
| ENSMUSG00000086067 | Gm16183 | 30.2918506 | 0.00976889 | 0.15146253 |
| ENSMUSG00000031485 | Plpbp | 1.17450347 | 0.00977953 | 0.15149188 |
| ENSMUSG00000058729 | Lin9 | 1.463573 | 0.00978271 | 0.15149188 |
| ENSMUSG00000037379 | Spon2 | 3.92830253 | 0.00980957 | 0.15174014 |
| ENSMUSG00000024675 | Ms4a4c | -2.1243962 | 0.00981068 | 0.15174014 |
| ENSMUSG00000066406 | Akap13 | -1.489435 | 0.00982447 | 0.15186096 |
| ENSMUSG00000022092 | Ppp3cc | -1.4025171 | 0.00983901 | 0.1519419 |
| ENSMUSG00000027540 | Ptpn1 | -1.2787568 | 0.00984166 | 0.1519419 |
| ENSMUSG00000021572 | Cep72 | 1.42025136 | 0.00986785 | 0.15225361 |
| ENSMUSG00000104217 | Gm37988 | -32.306286 | 0.00988324 | 0.15239861 |
| ENSMUSG00000004846 | Plod3 | -1.2950172 | 0.00994738 | 0.15316779 |
| ENSMUSG00000022075 | Rhobtb2 | 1.36247618 | 0.00995152 | 0.15316779 |
| ENSMUSG00000007033 | Hspa1l | 2.19972741 | 0.00996574 | 0.15316779 |
| ENSMUSG00000006307 | Kmt2b | 1.27261075 | 0.00996862 | 0.15316779 |
| ENSMUSG00000024993 | Fam45a | -1.303782 | 0.0099688 | 0.15316779 |
| ENSMUSG00000024727 | Trpm6 | 2.38403764 | 0.00996929 | 0.15316779 |
| ENSMUSG00000116812 | Gm2792 | 4.60146606 | 0.0099801 | 0.15324125 |
| ENSMUSG00000030036 | Mogs | 1.26426585 | 0.00999421 | 0.15336528 |
| ENSMUSG00000074419 | Gm15448 | -3.2876047 | 0.01001251 | 0.1535533 |
| ENSMUSG00000117465 | Gm49980 | 2.72592738 | 0.01002257 | 0.15357041 |
| ENSMUSG00000034486 | Gbx2 | -7.8118382 | 0.01002743 | 0.15357041 |
| ENSMUSG00000069920 | B3gnt9 | -1.9340834 | 0.01003175 | 0.15357041 |
| ENSMUSG00000057137 | Tmem140 | -1.6538507 | 0.01006879 | 0.15404462 |
| ENSMUSG00000058152 | Chsy3 | -3.0767477 | 0.01008462 | 0.15419396 |
| ENSMUSG00000055436 | Srsf11 | 1.35062604 | 0.01010748 | 0.15445056 |

| | | | | |
|---|---|---|---|---|
| ENSMUSG00000023156 | Rpp14 | 1.23236501 | 0.01013617 | 0.15471572 |
| ENSMUSG00000028843 | Sh3bgrl3 | -1.4817221 | 0.01014093 | 0.15471572 |
| ENSMUSG00000028483 | Snapc3 | 1.45297793 | 0.0101431 | 0.15471572 |
| ENSMUSG00000030019 | Fbxl14 | -1.1966908 | 0.01016123 | 0.15487756 |
| ENSMUSG00000008496 | Pou2f2 | -1.7507561 | 0.0101659 | 0.15487756 |
| ENSMUSG00000029925 | Tbxas1 | -2.0750943 | 0.01017764 | 0.15489571 |
| ENSMUSG00000056537 | Rlim | 1.24403336 | 0.01017928 | 0.15489571 |
| ENSMUSG00000039521 | Foxp3 | -3.2496018 | 0.0102096 | 0.15514038 |
| ENSMUSG00000020225 | Tmbim4 | -1.3997922 | 0.01021434 | 0.15514038 |
| ENSMUSG00000020262 | Adarb1 | -1.5923701 | 0.01021903 | 0.15514038 |
| ENSMUSG00000059447 | Hadhb | -1.1606301 | 0.01021978 | 0.15514038 |
| ENSMUSG00000003031 | Cdkn1b | 1.30278652 | 0.01023799 | 0.15530908 |
| ENSMUSG00000022892 | App | 1.47938175 | 0.01024553 | 0.15530908 |
| ENSMUSG00000043122 | A530016L24I | -12.666944 | 0.01025234 | 0.15530908 |
| ENSMUSG00000074102 | Rbm15b | 1.27193062 | 0.01025752 | 0.15530908 |
| ENSMUSG00000030316 | Tamm41 | 1.48318896 | 0.01026266 | 0.15530908 |
| ENSMUSG00000030043 | Tacr1 | -3.3590217 | 0.01026756 | 0.15530908 |
| ENSMUSG00000029036 | Atad3a | 1.44705435 | 0.01031035 | 0.15584931 |
| ENSMUSG00000031385 | Plxnb3 | -4.7072792 | 0.01031554 | 0.15584931 |
| ENSMUSG00000033644 | Piwil2 | 1.9380857 | 0.01035375 | 0.15632232 |
| ENSMUSG00000051498 | Tlr6 | -1.6931932 | 0.01035915 | 0.15632232 |
| ENSMUSG00000092627 | D130058E05 | -3.2875244 | 0.01038388 | 0.15648675 |
| ENSMUSG00000020265 | Sumo3 | 1.22262779 | 0.01038609 | 0.15648675 |
| ENSMUSG00000069609 | Cd300ld4 | -2.4256367 | 0.0103893 | 0.15648675 |
| ENSMUSG00000018169 | Mfng | -1.7564144 | 0.01039682 | 0.15648675 |
| ENSMUSG00000071037 | Camkmt | 1.54120936 | 0.01040084 | 0.15648675 |
| ENSMUSG00000056919 | Cep162 | 1.43842356 | 0.01041772 | 0.15664803 |
| ENSMUSG00000079597 | Cstdc4 | -48.746708 | 0.01043305 | 0.15678572 |
| ENSMUSG00000056917 | Sipa1 | -1.222745 | 0.01044248 | 0.15682697 |
| ENSMUSG00000043542 | Zc2hc1a | 1.35829996 | 0.01044814 | 0.15682697 |
| ENSMUSG00000035161 | Ints6 | 1.27788729 | 0.01046155 | 0.15693566 |
| ENSMUSG00000032243 | Itga11 | -3.5263073 | 0.0104685 | 0.15694725 |
| ENSMUSG00000055452 | Gm7353 | 2.70116882 | 0.01050265 | 0.15730411 |
| ENSMUSG00000030017 | Reg3g | -6.96605 | 0.01050469 | 0.15730411 |
| ENSMUSG00000032398 | Snapc5 | -1.5168647 | 0.01053545 | 0.15767184 |
| ENSMUSG00000006585 | Cdt1 | 1.44312991 | 0.01054784 | 0.15776446 |
| ENSMUSG00000058291 | Zfp68 | 1.17326376 | 0.01056935 | 0.15791446 |
| ENSMUSG00000030149 | Klrk1 | -2.2756598 | 0.01057563 | 0.15791446 |
| ENSMUSG00000039660 | Spout1 | 1.220221 | 0.01057652 | 0.15791446 |
| ENSMUSG00000019432 | Ddx39b | 1.23516422 | 0.01058321 | 0.15792158 |
| ENSMUSG00000040907 | Atp1a3 | -2.8794673 | 0.0106362 | 0.15861916 |
| ENSMUSG00000024053 | Emilin2 | -2.5638863 | 0.01064354 | 0.15863552 |
| ENSMUSG00000035189 | Ano4 | 4.14181122 | 0.01065158 | 0.15866239 |
| ENSMUSG00000039231 | Suv39h1 | 1.43931441 | 0.0106627 | 0.15873499 |
| ENSMUSG00000020131 | Pcsk4 | 1.72000508 | 0.01067359 | 0.15880402 |

| | | | | |
|---|---|---|---|---|
| ENSMUSG00000001435 | Col18a1 | -1.831867 | 0.01070285 | 0.15914616 |
| ENSMUSG00000060798 | Intu | 1.32212389 | 0.0107415 | 0.15962743 |
| ENSMUSG00000000753 | Serpinf1 | -1.9127617 | 0.01077238 | 0.15999278 |
| ENSMUSG00000021876 | Rnase4 | -1.8550895 | 0.01079753 | 0.16027264 |
| ENSMUSG00000062075 | Lmnb2 | 1.30751512 | 0.01082189 | 0.16054049 |
| ENSMUSG00000022179 | 4931414P19I | 1.39511155 | 0.0108487 | 0.16075076 |
| ENSMUSG00000058793 | Cds2 | -1.237243 | 0.01084872 | 0.16075076 |
| ENSMUSG00000040964 | Arhgef10l | 1.3016259 | 0.01085633 | 0.1607699 |
| ENSMUSG00000066877 | Nck2 | 1.25015692 | 0.01087759 | 0.16094632 |
| ENSMUSG00000008435 | Rdh13 | 1.35458643 | 0.01088358 | 0.16094632 |
| ENSMUSG00000027199 | Gatm | -1.651359 | 0.01088725 | 0.16094632 |
| ENSMUSG00000020592 | Sdc1 | -1.6142766 | 0.01090897 | 0.16117372 |
| ENSMUSG00000001632 | Brpf1 | 1.22306071 | 0.01091662 | 0.161193 |
| ENSMUSG00000028121 | Bcar3 | -2.243861 | 0.01093255 | 0.16133444 |
| ENSMUSG00000029314 | Gpat3 | -1.8899075 | 0.01096437 | 0.16170961 |
| ENSMUSG00000024640 | Psat1 | 2.03016501 | 0.01097324 | 0.16170961 |
| ENSMUSG00000018585 | Atox1 | -1.3505894 | 0.01097706 | 0.16170961 |
| ENSMUSG00000020891 | Alox8 | -4.5977961 | 0.01101905 | 0.16223416 |
| ENSMUSG00000059834 | Sclt1 | 1.42856318 | 0.01108518 | 0.16311328 |
| ENSMUSG00000026960 | Arl6ip6 | 1.34648002 | 0.01112883 | 0.16366081 |
| ENSMUSG00000028700 | Pomgnt1 | 1.28194901 | 0.01114498 | 0.16372773 |
| ENSMUSG00000093938 | Evi2b | -1.5637719 | 0.01114627 | 0.16372773 |
| ENSMUSG00000028466 | Creb3 | -1.3524741 | 0.01118836 | 0.16416541 |
| ENSMUSG00000024241 | Sos1 | 1.29423006 | 0.01118898 | 0.16416541 |
| ENSMUSG00000044294 | Krt84 | -168.47755 | 0.01119654 | 0.16418142 |
| ENSMUSG00000022186 | Oxct1 | 1.48783204 | 0.01123284 | 0.16445879 |
| ENSMUSG00000014782 | Plekhg4 | -2.9932129 | 0.01123319 | 0.16445879 |
| ENSMUSG00000026958 | Dpp7 | -1.579199 | 0.01123848 | 0.16445879 |
| ENSMUSG00000006050 | Sra1 | -1.321376 | 0.01124677 | 0.16445879 |
| ENSMUSG00000082925 | Gm13135 | -13.569165 | 0.01124781 | 0.16445879 |
| ENSMUSG00000026389 | Steap3 | -1.6672133 | 0.01126466 | 0.16461042 |
| ENSMUSG00000060261 | Gtf2i | 1.24017181 | 0.01127499 | 0.16462983 |
| ENSMUSG00000020196 | Cabin1 | 1.20612754 | 0.01127894 | 0.16462983 |
| ENSMUSG00000004591 | Pkn2 | 1.26193693 | 0.01132384 | 0.16519029 |
| ENSMUSG00000025017 | Pik3ap1 | -1.6867144 | 0.01136711 | 0.1657264 |
| ENSMUSG00000005687 | Bcas2 | 1.27760472 | 0.01138443 | 0.16588369 |
| ENSMUSG00000027030 | Stk39 | 1.56185739 | 0.01139616 | 0.16595943 |
| ENSMUSG00000036438 | Calm2 | 1.24821982 | 0.01142099 | 0.16622576 |
| ENSMUSG00000030780 | BC017158 | 1.28764984 | 0.01145506 | 0.16662625 |
| ENSMUSG00000039899 | Fgl2 | -1.6958719 | 0.01146514 | 0.16665576 |
| ENSMUSG00000059256 | Gzmd | -14.680064 | 0.01147766 | 0.16665576 |
| ENSMUSG00000105504 | Gbp5 | -1.8612843 | 0.0114897 | 0.16665576 |
| ENSMUSG00000034833 | Tespa1 | -3.9548309 | 0.01149356 | 0.16665576 |
| ENSMUSG00000021759 | Plpp1 | -1.6943765 | 0.01149609 | 0.16665576 |
| ENSMUSG00000008136 | Fhl2 | -1.5558622 | 0.01150225 | 0.16665576 |

| | | | | |
|---|---|---|---|---|
| ENSMUSG00000086740 | Gm17029 | 3.22181725 | 0.01150715 | 0.16665576 |
| ENSMUSG00000025982 | Sf3b1 | 1.2169754 | 0.0115116 | 0.16665576 |
| ENSMUSG00000026989 | Dapl1 | -11.865854 | 0.01153112 | 0.16665576 |
| ENSMUSG00000110720 | Gm46223 | -5.9393753 | 0.01153164 | 0.16665576 |
| ENSMUSG00000078762 | Haus5 | 1.36886914 | 0.0115396 | 0.16665576 |
| ENSMUSG00000084973 | Gm13848 | 5.80304431 | 0.0115426 | 0.16665576 |
| ENSMUSG00000001964 | Emd | 1.44300732 | 0.01154523 | 0.16665576 |
| ENSMUSG00000023031 | Cela1 | -2.0939339 | 0.01154891 | 0.16665576 |
| ENSMUSG00000025036 | Sfxn2 | 1.58819443 | 0.0115886 | 0.16713359 |
| ENSMUSG00000029291 | Rufy3 | 1.41277892 | 0.01166404 | 0.16808682 |
| ENSMUSG00000038607 | Gng10 | -1.4616596 | 0.01167143 | 0.16808682 |
| ENSMUSG00000021981 | Cab39l | 1.2421181 | 0.01167453 | 0.16808682 |
| ENSMUSG00000021214 | Akr1c18 | -4.1557697 | 0.0117123 | 0.16853507 |
| ENSMUSG00000004936 | Map2k1 | -1.3216766 | 0.01173807 | 0.16881035 |
| ENSMUSG00000028708 | Mknk1 | -1.2470382 | 0.01175408 | 0.16891542 |
| ENSMUSG00000050359 | Sprr1a | -4.7890278 | 0.01176221 | 0.16891542 |
| ENSMUSG00000035472 | Slc25a21 | -3.2733097 | 0.01176802 | 0.16891542 |
| ENSMUSG00000029084 | Cd38 | -1.7524773 | 0.01177197 | 0.16891542 |
| ENSMUSG00000029373 | Pf4 | -2.5488086 | 0.01177919 | 0.16892367 |
| ENSMUSG00000003418 | St8sia6 | 2.4880835 | 0.0118078 | 0.16923848 |
| ENSMUSG00000028751 | Pla2g2e | -59.596306 | 0.01183164 | 0.16947852 |
| ENSMUSG00000034329 | Brip1 | 1.47803478 | 0.01183789 | 0.16947852 |
| ENSMUSG00000038482 | Tfdp1 | 1.37103547 | 0.01185219 | 0.16958443 |
| ENSMUSG00000032502 | Stac | -6.6622801 | 0.01186096 | 0.16958443 |
| ENSMUSG00000025856 | Pdgfa | 1.53375819 | 0.01187098 | 0.16958443 |
| ENSMUSG00000026090 | 2010300C02I | 2.41313237 | 0.01187198 | 0.16958443 |
| ENSMUSG00000042312 | S100a13 | -1.4974227 | 0.01189818 | 0.16986319 |
| ENSMUSG00000025511 | Tspan4 | -1.4045073 | 0.01191747 | 0.17004314 |
| ENSMUSG00000037608 | Bclaf1 | 1.33935702 | 0.01194035 | 0.17027395 |
| ENSMUSG00000054364 | Rhob | -1.601469 | 0.01195353 | 0.17033702 |
| ENSMUSG00000022546 | Gpt | 2.750326 | 0.01195818 | 0.17033702 |
| ENSMUSG00000036501 | Fam13b | 1.56731361 | 0.01196854 | 0.17038913 |
| ENSMUSG00000027496 | Aurka | 1.61020601 | 0.01202318 | 0.17107006 |
| ENSMUSG00000028270 | Gbp2 | -1.9610887 | 0.01202984 | 0.17107006 |
| ENSMUSG00000098975 | Gm27177 | 3.27774942 | 0.01208978 | 0.17182635 |
| ENSMUSG00000062939 | Stat4 | -2.5247341 | 0.01209801 | 0.17184583 |
| ENSMUSG00000001943 | Vsig2 | 6.06823566 | 0.01210468 | 0.17184583 |
| ENSMUSG00000044576 | Garem2 | -6.6387878 | 0.01214325 | 0.17229715 |
| ENSMUSG00000051950 | B3glct | 1.5900559 | 0.01215402 | 0.17235382 |
| ENSMUSG00000099083 | Atf7 | 1.39353916 | 0.01216997 | 0.17248371 |
| ENSMUSG00000101609 | Kcnq1ot1 | 2.05855709 | 0.01223941 | 0.17337123 |
| ENSMUSG00000002748 | Baz1b | 1.29056727 | 0.01226145 | 0.17351059 |
| ENSMUSG00000022377 | Asap1 | -1.570066 | 0.01226291 | 0.17351059 |
| ENSMUSG00000027884 | Clcc1 | 1.18363857 | 0.01227815 | 0.17354987 |
| ENSMUSG00000042524 | Sun2 | 1.3612661 | 0.01228702 | 0.17354987 |

| | | | | |
|---|---|---|---|---|
| ENSMUSG00000032369 | Plscr1 | 1.60157328 | 0.01229033 | 0.17354987 |
| ENSMUSG00000071005 | Ccl19 | -9.0029898 | 0.012293 | 0.17354987 |
| ENSMUSG00000020697 | Lig3 | 1.32889204 | 0.0123226 | 0.17387114 |
| ENSMUSG00000089872 | Rps6kc1 | -1.2529952 | 0.01233246 | 0.17391367 |
| ENSMUSG00000042462 | Dctpp1 | 1.61742461 | 0.01237629 | 0.17420133 |
| ENSMUSG00000028945 | Rheb | 1.14852872 | 0.01238026 | 0.17420133 |
| ENSMUSG00000051344 | Plekhm3 | -1.5482008 | 0.0123816 | 0.17420133 |
| ENSMUSG00000021546 | Hnrnpk | 1.11358652 | 0.0123825 | 0.17420133 |
| ENSMUSG00000025328 | Padi3 | -9.6392023 | 0.01238713 | 0.17420133 |
| ENSMUSG00000081600 | Gm12286 | -19.250343 | 0.01240361 | 0.17431104 |
| ENSMUSG00000027963 | Extl2 | -1.3533809 | 0.01240865 | 0.17431104 |
| ENSMUSG00000090210 | Itga10 | 2.14100368 | 0.0124352 | 0.17456445 |
| ENSMUSG00000085440 | Sorbs2os | 12.916128 | 0.01244163 | 0.17456445 |
| ENSMUSG00000029376 | Mthfd2l | 1.37286308 | 0.01245206 | 0.17456445 |
| ENSMUSG00000027006 | Dnajc10 | 1.71226642 | 0.01245417 | 0.17456445 |
| ENSMUSG00000071552 | Tigit | -2.7186986 | 0.01251517 | 0.17524132 |
| ENSMUSG00000020184 | Mdm2 | -1.2446983 | 0.01252031 | 0.17524132 |
| ENSMUSG00000019877 | Serinc1 | -1.1982888 | 0.01252315 | 0.17524132 |
| ENSMUSG00000085237 | Gm15406 | 14.365133 | 0.01253149 | 0.17526158 |
| ENSMUSG00000004558 | Ndrg2 | -2.1045163 | 0.01255582 | 0.17548803 |
| ENSMUSG00000001518 | Itfg2 | -1.2688931 | 0.0125615 | 0.17548803 |
| ENSMUSG00000031066 | Usp11 | -1.8037699 | 0.01258597 | 0.17567266 |
| ENSMUSG00000028988 | Ctnnbip1 | 1.41917072 | 0.01258854 | 0.17567266 |
| ENSMUSG00000103144 | Pcdhga1 | 3.29789722 | 0.01262512 | 0.17608376 |
| ENSMUSG00000074622 | Mafb | -1.6246021 | 0.01263597 | 0.17608376 |
| ENSMUSG00000057551 | Zfp317 | 1.33937878 | 0.01263878 | 0.17608376 |
| ENSMUSG00000026917 | Wdr5 | 1.30849673 | 0.01266241 | 0.1763163 |
| ENSMUSG00000025899 | Alkbh8 | 1.39479386 | 0.01267287 | 0.17636531 |
| ENSMUSG00000024378 | Stard4 | 1.43522454 | 0.01272572 | 0.17700377 |
| ENSMUSG00000054932 | Afp | -7.3194832 | 0.01277891 | 0.17764645 |
| ENSMUSG00000022364 | Tbc1d31 | 1.52155517 | 0.0127996 | 0.17783681 |
| ENSMUSG00000030862 | Cpxm2 | -2.8299466 | 0.01280859 | 0.17786359 |
| ENSMUSG00000000058 | Cav2 | -1.3307267 | 0.01281553 | 0.17786359 |
| ENSMUSG00000051790 | Nlgn2 | -1.4256649 | 0.01284084 | 0.17811756 |
| ENSMUSG00000096336 | Igkv1-135 | -31.178944 | 0.01285005 | 0.17814804 |
| ENSMUSG00000026315 | Serpinb8 | -2.2080526 | 0.01287978 | 0.17842404 |
| ENSMUSG00000062300 | Nectin2 | 1.36016261 | 0.01289976 | 0.17842404 |
| ENSMUSG00000026166 | Ccl20 | 14.3127115 | 0.01290309 | 0.17842404 |
| ENSMUSG00000026833 | Olfm1 | -1.6565866 | 0.01291368 | 0.17842404 |
| ENSMUSG00000006304 | Arpc2 | -1.1868256 | 0.01291431 | 0.17842404 |
| ENSMUSG00000038181 | Chpf2 | -1.3492848 | 0.0129168 | 0.17842404 |
| ENSMUSG00000090247 | Bloc1s1 | -1.3670219 | 0.0129191 | 0.17842404 |
| ENSMUSG00000027398 | Il1b | -2.3691743 | 0.01292943 | 0.17846963 |
| ENSMUSG00000026974 | Zmynd19 | 1.47502347 | 0.0129478 | 0.17862626 |
| ENSMUSG00000020798 | Spns3 | 2.28135172 | 0.01296783 | 0.17871275 |

| | | | | |
|---|---|---|---|---|
| ENSMUSG00000044700 | Tmem201 | 1.51562594 | 0.01296987 | 0.17871275 |
| ENSMUSG00000016918 | Sulf1 | -1.8515325 | 0.01297517 | 0.17871275 |
| ENSMUSG00000096960 | A230028O05 | -10.393199 | 0.01298421 | 0.17874042 |
| ENSMUSG00000029602 | Rasal1 | 2.5023466 | 0.01300407 | 0.17884643 |
| ENSMUSG00000038552 | Fndc4 | 1.71897641 | 0.01300599 | 0.17884643 |
| ENSMUSG00000037946 | Fgd3 | -1.5063168 | 0.01302024 | 0.17894561 |
| ENSMUSG00000025821 | Zfp282 | 1.27869723 | 0.0131539 | 0.18068485 |
| ENSMUSG00000097636 | Mirt1 | -2.8621993 | 0.0132018 | 0.18124487 |
| ENSMUSG00000067786 | Nnat | -3.4867788 | 0.01323058 | 0.1815418 |
| ENSMUSG00000037902 | Sirpa | -1.6050222 | 0.0132434 | 0.18161971 |
| ENSMUSG00000015133 | Lrrk1 | -1.4937996 | 0.01328884 | 0.18214458 |
| ENSMUSG00000012889 | Podnl1 | -2.1749525 | 0.01331419 | 0.18235921 |
| ENSMUSG00000041130 | Zfp598 | 1.22393784 | 0.01331885 | 0.18235921 |
| ENSMUSG00000032601 | Prkar2a | 1.29268362 | 0.01334883 | 0.18259352 |
| ENSMUSG00000040331 | Nsmce4a | 1.3220204 | 0.01335034 | 0.18259352 |
| ENSMUSG00000038267 | Slc22a23 | 1.60977106 | 0.01335794 | 0.18259919 |
| ENSMUSG00000032126 | Hmbs | 1.30894386 | 0.01336999 | 0.18266574 |
| ENSMUSG00000044469 | Tnfaip8l1 | 1.55323173 | 0.0133805 | 0.18271102 |
| ENSMUSG00000024212 | Mllt1 | 1.27592419 | 0.01338879 | 0.18272611 |
| ENSMUSG00000002059 | Rab34 | -1.3875556 | 0.01339775 | 0.18275022 |
| ENSMUSG00000036896 | C1qc | -1.5813723 | 0.01340859 | 0.1827999 |
| ENSMUSG00000027641 | Rbl1 | 1.39352583 | 0.01341651 | 0.18280988 |
| ENSMUSG00000032231 | Anxa2 | -1.3595541 | 0.01344592 | 0.18301836 |
| ENSMUSG00000019874 | Fabp7 | -17.126698 | 0.01344622 | 0.18301836 |
| ENSMUSG00000079419 | Ms4a6c | -1.5052627 | 0.01346186 | 0.18313317 |
| ENSMUSG00000035861 | Tmprss11b | -21.008661 | 0.01348838 | 0.18339577 |
| ENSMUSG00000035949 | Fbxw2 | 1.16768237 | 0.01350514 | 0.18340818 |
| ENSMUSG00000074269 | Rec114 | 2.0582367 | 0.01350808 | 0.18340818 |
| ENSMUSG00000103567 | Pcdhga5 | -3.3738856 | 0.01351541 | 0.18340818 |
| ENSMUSG00000031486 | Adgra2 | -1.9658573 | 0.01351816 | 0.18340818 |
| ENSMUSG00000060441 | Trim5 | -2.3282819 | 0.01356755 | 0.18385298 |
| ENSMUSG00000028433 | Ubap2 | 1.23755381 | 0.01356925 | 0.18385298 |
| ENSMUSG00000022800 | Fyttd1 | 1.16013405 | 0.01357265 | 0.18385298 |
| ENSMUSG00000027860 | Vangl1 | 1.28384122 | 0.01359621 | 0.18397578 |
| ENSMUSG00000072872 | Rybp | 1.28362948 | 0.0136044 | 0.18397578 |
| ENSMUSG00000050555 | Hyls1 | 1.41581556 | 0.01360792 | 0.18397578 |
| ENSMUSG00000050890 | Pdik1l | 1.34941245 | 0.01361067 | 0.18397578 |
| ENSMUSG00000045679 | Pqlc3 | -1.4513866 | 0.01366661 | 0.18463362 |
| ENSMUSG00000043336 | Filip1l | -1.451118 | 0.01368388 | 0.18476867 |
| ENSMUSG00000048285 | Frmd6 | -1.5503271 | 0.0137201 | 0.18515935 |
| ENSMUSG00000055065 | Ddx17 | 1.37972857 | 0.01372786 | 0.18516584 |
| ENSMUSG00000039990 | Edrf1 | 1.35908439 | 0.01375964 | 0.18547008 |
| ENSMUSG00000041530 | Ago1 | 1.2957544 | 0.01376502 | 0.18547008 |
| ENSMUSG00000041390 | Mdfic | -1.4052096 | 0.01384546 | 0.18645519 |
| ENSMUSG00000025358 | Cdk2 | 1.38856577 | 0.01385683 | 0.18650945 |

| | | | | |
|---|---|---|---|---|
| ENSMUSG00000026418 | Tnni1 | -3.3371725 | 0.01387047 | 0.18659413 |
| ENSMUSG00000026743 | Mllt10 | 1.22103576 | 0.01392039 | 0.18716658 |
| ENSMUSG00000021456 | Fbp2 | -4.3015216 | 0.01395257 | 0.18750017 |
| ENSMUSG00000004892 | Bcan | -9.1308218 | 0.01400825 | 0.1881489 |
| ENSMUSG00000020672 | Sntg2 | -3.5709896 | 0.01404331 | 0.18852019 |
| ENSMUSG00000028399 | Ptprd | -6.0837606 | 0.01408393 | 0.18896565 |
| ENSMUSG00000020886 | Dlg4 | -1.8320159 | 0.01409471 | 0.18897303 |
| ENSMUSG00000035382 | Pcsk7 | 1.29555544 | 0.01409936 | 0.18897303 |
| ENSMUSG00000058354 | Krt6a | -9.5275644 | 0.01412528 | 0.18922067 |
| ENSMUSG00000035279 | Ssc5d | -2.650406 | 0.01415108 | 0.1894149 |
| ENSMUSG00000003123 | Lipe | -2.106807 | 0.01416075 | 0.1894149 |
| ENSMUSG00000063810 | Alms1 | 1.62959329 | 0.01416378 | 0.1894149 |
| ENSMUSG00000023047 | Amhr2 | 2.84949549 | 0.01416959 | 0.1894149 |
| ENSMUSG00000075585 | 6330403L08F | 1.37234567 | 0.01417732 | 0.18941847 |
| ENSMUSG00000028948 | Nol9 | 1.32241311 | 0.01418826 | 0.18946511 |
| ENSMUSG00000034891 | Sncb | 10.4899677 | 0.01419762 | 0.1894905 |
| ENSMUSG00000040738 | Ints8 | 1.228048 | 0.01422915 | 0.18981169 |
| ENSMUSG00000030924 | Rexo5 | 1.79120797 | 0.01425651 | 0.18999765 |
| ENSMUSG00000026663 | Atf6 | -1.2710242 | 0.01426138 | 0.18999765 |
| ENSMUSG00000020212 | Mdm1 | 1.62521232 | 0.01426552 | 0.18999765 |
| ENSMUSG00000076490 | Trbc1 | -2.5140147 | 0.01431514 | 0.19055853 |
| ENSMUSG00000028487 | Bnc2 | -3.331417 | 0.01433792 | 0.1907619 |
| ENSMUSG00000097828 | 6430562O15 | 2.73836185 | 0.01434739 | 0.19078801 |
| ENSMUSG00000056025 | Clca3a1 | 3.78531709 | 0.01437159 | 0.19100987 |
| ENSMUSG00000020361 | Hspa4 | -1.1873538 | 0.01440325 | 0.19133062 |
| ENSMUSG00000058748 | Zfp958 | 1.37075662 | 0.01444477 | 0.1916329 |
| ENSMUSG00000032479 | Map4 | -1.2667463 | 0.01445351 | 0.1916329 |
| ENSMUSG00000039153 | Runx2 | -1.561598 | 0.01445862 | 0.1916329 |
| ENSMUSG00000038611 | Phrf1 | 1.20303836 | 0.01445972 | 0.1916329 |
| ENSMUSG00000043518 | Rai2 | -2.9301882 | 0.01446678 | 0.1916329 |
| ENSMUSG00000052477 | C130026I21R | 3.1037524 | 0.01447125 | 0.1916329 |
| ENSMUSG00000052013 | Btla | -4.1195485 | 0.01447927 | 0.19163914 |
| ENSMUSG00000024041 | Cryaa | -11.02362 | 0.01449106 | 0.19169541 |
| ENSMUSG00000003746 | Man1a | -1.5013121 | 0.01450438 | 0.19177175 |
| ENSMUSG00000009575 | Cbx5 | 1.38145779 | 0.01451208 | 0.19177376 |
| ENSMUSG00000032423 | Syncrip | 1.29945487 | 0.01455402 | 0.19218578 |
| ENSMUSG00000118378 | Gm50393 | 22.6651481 | 0.01456403 | 0.19218578 |
| ENSMUSG00000026271 | Gpr35 | -1.6951656 | 0.01456595 | 0.19218578 |
| ENSMUSG00000079455 | Gm16026 | 5.32882472 | 0.01458157 | 0.19229205 |
| ENSMUSG00000028066 | Pmf1 | 1.41611599 | 0.01459571 | 0.19237869 |
| ENSMUSG00000117042 | 2700054A10 | 2.77351161 | 0.01461947 | 0.19250034 |
| ENSMUSG00000042790 | Rnf214 | 1.29124591 | 0.0146233 | 0.19250034 |
| ENSMUSG00000027018 | Hat1 | 1.32234974 | 0.01462766 | 0.19250034 |
| ENSMUSG00000062980 | Cped1 | -1.9185456 | 0.01464302 | 0.19260264 |
| ENSMUSG00000029648 | Flt1 | -1.6324949 | 0.0146803 | 0.19297728 |

| | | | | |
|---|---|---|---|---|
| ENSMUSG00000002064 | Sdf2 | -1.3192302 | 0.01468669 | 0.19297728 |
| ENSMUSG00000048534 | Jaml | -2.1285462 | 0.01469667 | 0.19298895 |
| ENSMUSG00000076613 | Ighg2b | -22.750808 | 0.01470276 | 0.19298895 |
| ENSMUSG00000085338 | 2410004I01R | 12.1462153 | 0.01476067 | 0.19362087 |
| ENSMUSG00000060862 | Zbtb40 | 1.36171643 | 0.0147679 | 0.19362087 |
| ENSMUSG00000021597 | Slf1 | 1.34029688 | 0.01477755 | 0.19362087 |
| ENSMUSG00000097537 | 2610020C07I | 2.3825424 | 0.01478138 | 0.19362087 |
| ENSMUSG00000086233 | Gm11816 | -7.4217592 | 0.01484499 | 0.19435386 |
| ENSMUSG00000076934 | Iglv1 | -25.054226 | 0.01485316 | 0.19436065 |
| ENSMUSG00000027552 | E2f5 | 1.38073226 | 0.01486636 | 0.19443327 |
| ENSMUSG00000033510 | Otud7a | -4.7130082 | 0.01489127 | 0.19465895 |
| ENSMUSG00000021922 | Itih4 | -10.50882 | 0.01491127 | 0.19482015 |
| ENSMUSG00000019872 | Smpdl3a | -1.5967093 | 0.01492117 | 0.19484934 |
| ENSMUSG00000058755 | Osm | -3.2989476 | 0.01493678 | 0.19493524 |
| ENSMUSG00000047412 | Zbtb44 | 1.28252288 | 0.01494309 | 0.19493524 |
| ENSMUSG00000027502 | Rtf2 | -1.2349697 | 0.01495966 | 0.1949788 |
| ENSMUSG00000024833 | Pola2 | 1.43374895 | 0.01496178 | 0.1949788 |
| ENSMUSG00000042549 | Map2k3os | 1.69481061 | 0.01497042 | 0.19499148 |
| ENSMUSG00000029127 | Zbtb49 | 1.39773547 | 0.01498228 | 0.195046 |
| ENSMUSG00000050875 | Minar2 | -6.8129278 | 0.01502093 | 0.19544904 |
| ENSMUSG00000040818 | Dennd6a | 1.23283708 | 0.01504931 | 0.19571809 |
| ENSMUSG00000008193 | Spib | 8.96087124 | 0.0151013 | 0.19629367 |
| ENSMUSG00000024986 | Hhex | -1.6436282 | 0.01511193 | 0.19633146 |
| ENSMUSG00000042328 | Hps4 | 1.28747601 | 0.01511974 | 0.19633254 |
| ENSMUSG00000048668 | Rhno1 | 1.39038928 | 0.01516491 | 0.19678207 |
| ENSMUSG00000112980 | D430020J02F | 1.71183717 | 0.01516984 | 0.19678207 |
| ENSMUSG00000078300 | Gm2606 | 5.95651077 | 0.01518503 | 0.19687862 |
| ENSMUSG00000024965 | Fermt3 | -1.5609324 | 0.0152136 | 0.19714838 |
| ENSMUSG00000025465 | Echs1 | 1.26657628 | 0.01523207 | 0.19715698 |
| ENSMUSG00000026944 | Abca2 | 1.349508 | 0.01523661 | 0.19715698 |
| ENSMUSG00000073988 | Ttpa | 2.31114876 | 0.01524386 | 0.19715698 |
| ENSMUSG00000026339 | Ccdc93 | 1.27360478 | 0.01524746 | 0.19715698 |
| ENSMUSG00000059921 | Unc5c | -5.5870617 | 0.01525623 | 0.19715698 |
| ENSMUSG00000116618 | Gm49719 | -18.52514 | 0.01526081 | 0.19715698 |
| ENSMUSG00000031714 | Gab1 | 1.4510891 | 0.01529395 | 0.19748464 |
| ENSMUSG00000051675 | Trim32 | 1.33996696 | 0.01533441 | 0.19790656 |
| ENSMUSG00000031258 | Xkrx | -6.0233007 | 0.01535675 | 0.19809422 |
| ENSMUSG00000031538 | Plat | -1.9236586 | 0.01537409 | 0.1982173 |
| ENSMUSG00000039523 | Cep104 | 1.23430857 | 0.01542205 | 0.1987349 |
| ENSMUSG00000007646 | Rad51c | 1.64492273 | 0.01543955 | 0.19885953 |
| ENSMUSG00000042638 | Gucy2c | 4.93610458 | 0.01546261 | 0.19903482 |
| ENSMUSG00000079469 | Pigb | 1.24381251 | 0.01546882 | 0.19903482 |
| ENSMUSG00000027202 | Slc12a1 | -10.636878 | 0.01552946 | 0.19971384 |
| ENSMUSG00000046490 | Rnf222 | -53.426196 | 0.01558586 | 0.20033777 |
| ENSMUSG00000026014 | Raph1 | 1.67783252 | 0.01559437 | 0.20034593 |

| | | | | |
|---|---|---|---|---|
| ENSMUSG00000034171 | Faah | 1.65166991 | 0.01561443 | 0.20037803 |
| ENSMUSG00000025287 | Acot9 | -1.3731352 | 0.01561862 | 0.20037803 |
| ENSMUSG00000079477 | Rab7 | -1.1802243 | 0.01562487 | 0.20037803 |
| ENSMUSG00000037860 | Aim2 | -1.5415789 | 0.01563746 | 0.20037803 |
| ENSMUSG00000037020 | Wdr62 | 1.50184713 | 0.01563916 | 0.20037803 |
| ENSMUSG00000057191 | AB124611 | -1.9568433 | 0.01564418 | 0.20037803 |
| ENSMUSG00000030271 | Ogg1 | 1.38400971 | 0.01567133 | 0.20057226 |
| ENSMUSG00000074141 | Il4i1 | 3.92572573 | 0.01568034 | 0.20057226 |
| ENSMUSG00000092490 | Gm20482 | 6.43332612 | 0.01568977 | 0.20057226 |
| ENSMUSG00000022803 | Popdc2 | 6.3340569 | 0.01569092 | 0.20057226 |
| ENSMUSG00000020844 | Nxn | -1.4913259 | 0.01570272 | 0.20062218 |
| ENSMUSG00000028884 | Rpa2 | 1.60677412 | 0.01571383 | 0.20066321 |
| ENSMUSG00000089715 | Cbx6 | 1.51130744 | 0.01579987 | 0.20154204 |
| ENSMUSG00000061607 | Mdc1 | 1.39837476 | 0.0158059 | 0.20154204 |
| ENSMUSG00000074934 | Grem1 | -6.6317767 | 0.01580644 | 0.20154204 |
| ENSMUSG00000068614 | Actc1 | 17.8130559 | 0.01586836 | 0.20223002 |
| ENSMUSG00000029338 | Antxr2 | -1.484156 | 0.01589122 | 0.20241985 |
| ENSMUSG00000099481 | Xndc1 | 1.4424737 | 0.01594115 | 0.20292413 |
| ENSMUSG00000020260 | Pofut2 | -1.2997672 | 0.01594678 | 0.20292413 |
| ENSMUSG00000041801 | Phlda3 | -1.4486839 | 0.0159563 | 0.20294369 |
| ENSMUSG00000039047 | Pigk | -1.2529267 | 0.01605412 | 0.20390005 |
| ENSMUSG00000039176 | Polg | 1.30432326 | 0.0160553 | 0.20390005 |
| ENSMUSG00000024970 | Spindoc | -1.3016307 | 0.01605556 | 0.20390005 |
| ENSMUSG00000024097 | Srsf7 | 1.35687028 | 0.01607002 | 0.20394191 |
| ENSMUSG00000022978 | Mis18a | 1.47283191 | 0.01607491 | 0.20394191 |
| ENSMUSG00000024909 | Efemp2 | -1.3280088 | 0.01609796 | 0.20410964 |
| ENSMUSG00000022658 | Tagln3 | -22.229048 | 0.0161042 | 0.20410964 |
| ENSMUSG00000022491 | Glycam1 | -277.83285 | 0.01611593 | 0.20415647 |
| ENSMUSG00000056458 | Mok | 2.09453116 | 0.0161298 | 0.20423037 |
| ENSMUSG00000000982 | Ccl3 | -2.6981836 | 0.01615173 | 0.20424157 |
| ENSMUSG00000020988 | L2hgdh | 1.40238341 | 0.01615863 | 0.20424157 |
| ENSMUSG00000029344 | Tpst2 | -1.3047867 | 0.01616075 | 0.20424157 |
| ENSMUSG00000003039 | Fam32a | 1.32484074 | 0.01616604 | 0.20424157 |
| ENSMUSG00000038264 | Sema7a | -2.0362391 | 0.01617659 | 0.20424157 |
| ENSMUSG00000025403 | Shmt2 | 1.24206565 | 0.0161789 | 0.20424157 |
| ENSMUSG00000020152 | Actr2 | -1.213904 | 0.01623484 | 0.20484598 |
| ENSMUSG00000113902 | Ndufb1-ps | -1.3556447 | 0.01626997 | 0.20518735 |
| ENSMUSG00000084990 | Gm14549 | -3.8489628 | 0.01630135 | 0.20548106 |
| ENSMUSG00000001785 | Pwp1 | 1.26621319 | 0.01634827 | 0.20597043 |
| ENSMUSG00000030428 | Ttyh1 | -3.6751954 | 0.01637045 | 0.20614765 |
| ENSMUSG00000026238 | Ptma | 1.21696303 | 0.01637877 | 0.20615016 |
| ENSMUSG00000020649 | Rrm2 | 1.41114957 | 0.01643924 | 0.2068089 |
| ENSMUSG00000026832 | Cytip | 2.27276391 | 0.01644932 | 0.20683336 |
| ENSMUSG00000027698 | Nceh1 | -1.4772543 | 0.01647644 | 0.20706408 |
| ENSMUSG00000024253 | Dync2li1 | 1.39632356 | 0.01648397 | 0.20706408 |

| | | | | |
|---|---|---|---|---|
| ENSMUSG00000028845 | Tekt2 | 4.5656052 | 0.01650237 | 0.20719284 |
| ENSMUSG00000107227 | Gm42559 | 3.3920914 | 0.01651885 | 0.20723889 |
| ENSMUSG00000061068 | Mcpt4 | -2.8976147 | 0.01652787 | 0.20723889 |
| ENSMUSG00000040260 | Daam2 | -2.4489003 | 0.01653051 | 0.20723889 |
| ENSMUSG00000030806 | Stx1b | 4.0592574 | 0.01654412 | 0.20730724 |
| ENSMUSG00000036199 | Ndufa13 | -1.255811 | 0.0165584 | 0.20738395 |
| ENSMUSG00000028180 | Zranb2 | 1.28623445 | 0.01657129 | 0.20744312 |
| ENSMUSG00000102929 | Gm37154 | 4.66483697 | 0.01658581 | 0.20752275 |
| ENSMUSG00000028517 | Plpp3 | -1.4015978 | 0.01660661 | 0.20768064 |
| ENSMUSG00000026258 | Snorc | 6.4104505 | 0.01661534 | 0.20768767 |
| ENSMUSG00000033762 | Recql4 | 1.62673404 | 0.01662886 | 0.20775448 |
| ENSMUSG00000022723 | Crybg3 | 1.51718058 | 0.01667173 | 0.20814208 |
| ENSMUSG00000087543 | Gm16576 | 2.06322632 | 0.01667627 | 0.20814208 |
| ENSMUSG00000036242 | Armh4 | -3.6774046 | 0.01671829 | 0.20856415 |
| ENSMUSG00000087679 | Tmem250-ps | 1.25038801 | 0.01672781 | 0.20858052 |
| ENSMUSG00000020263 | Appl2 | 1.59420129 | 0.01674023 | 0.20863298 |
| ENSMUSG00000021713 | Ppwd1 | 1.31830977 | 0.01676183 | 0.20876051 |
| ENSMUSG00000102840 | Gm38037 | 3.05606856 | 0.01676689 | 0.20876051 |
| ENSMUSG00000037013 | Ss18 | 1.21387467 | 0.01682104 | 0.20925521 |
| ENSMUSG00000049606 | Zfp644 | 1.26941576 | 0.0168292 | 0.20925521 |
| ENSMUSG00000051557 | Pusl1 | 1.35589234 | 0.01683133 | 0.20925521 |
| ENSMUSG00000057135 | Scimp | -2.0130785 | 0.01687088 | 0.20964439 |
| ENSMUSG00000030134 | Rasgef1a | -4.0133483 | 0.01692215 | 0.21017872 |
| ENSMUSG00000105906 | Iglc1 | -16.711708 | 0.01693858 | 0.21018009 |
| ENSMUSG00000047261 | Gap43 | -6.0802921 | 0.01693894 | 0.21018009 |
| ENSMUSG00000103464 | Gm38246 | 12.0965385 | 0.01695402 | 0.21018009 |
| ENSMUSG00000064127 | Med14 | 1.38206153 | 0.01695534 | 0.21018009 |
| ENSMUSG00000031532 | Saraf | -1.2986304 | 0.01698565 | 0.21043541 |
| ENSMUSG00000078653 | Cntd1 | 4.35145723 | 0.0169925 | 0.21043541 |
| ENSMUSG00000043252 | Tmem64 | 1.38199104 | 0.01701263 | 0.21058201 |
| ENSMUSG00000031706 | Rfx1 | 1.30355516 | 0.01702307 | 0.21060874 |
| ENSMUSG00000033054 | Npat | 1.38904098 | 0.01711546 | 0.211561 |
| ENSMUSG00000036155 | Mgat5 | -1.4070959 | 0.01711669 | 0.211561 |
| ENSMUSG00000022422 | Dscc1 | 1.67360486 | 0.01714001 | 0.2117462 |
| ENSMUSG00000026668 | Ucma | 43.2236915 | 0.01723421 | 0.21280653 |
| ENSMUSG00000027375 | Mal | -4.9684524 | 0.01727 | 0.21296363 |
| ENSMUSG00000041479 | Syt15 | -2.588959 | 0.01727152 | 0.21296363 |
| ENSMUSG00000069727 | Zfp975 | 1.43821762 | 0.01728774 | 0.21296363 |
| ENSMUSG00000037808 | Fam76b | 1.27895692 | 0.01728988 | 0.21296363 |
| ENSMUSG00000094437 | Gm9830 | 39.7984887 | 0.01729045 | 0.21296363 |
| ENSMUSG00000042289 | Hsd3b7 | -1.4755915 | 0.01730042 | 0.21296363 |
| ENSMUSG00000052384 | Nrros | -1.6184792 | 0.01731196 | 0.21296363 |
| ENSMUSG00000020717 | Pecam1 | -1.5278978 | 0.01731729 | 0.21296363 |
| ENSMUSG00000035237 | Lcat | -3.9149043 | 0.0173296 | 0.21296363 |
| ENSMUSG00000026429 | Ube2t | 1.46196508 | 0.01733117 | 0.21296363 |

| | | | | |
|---|---|---|---|---|
| ENSMUSG00000027478 | Dnmt3b | 1.70486238 | 0.01733912 | 0.21296363 |
| ENSMUSG00000020393 | Kremen1 | -1.6741247 | 0.01740305 | 0.21364555 |
| ENSMUSG00000043456 | Zfp536 | -8.6055982 | 0.01741409 | 0.21367781 |
| ENSMUSG00000001741 | Il16 | -1.7008536 | 0.01742694 | 0.21373233 |
| ENSMUSG00000049130 | C5ar1 | -2.153812 | 0.01744954 | 0.21390626 |
| ENSMUSG00000022295 | Atp6v1c1 | -1.2884506 | 0.01746117 | 0.21394564 |
| ENSMUSG00000037703 | Lzts3 | 1.56731118 | 0.01747282 | 0.21398523 |
| ENSMUSG00000058163 | Gm5431 | -1.9143253 | 0.01749917 | 0.21420472 |
| ENSMUSG00000030162 | Olr1 | -4.7534312 | 0.0175243 | 0.21440903 |
| ENSMUSG00000044702 | Palb2 | 1.49745104 | 0.01754863 | 0.21460335 |
| ENSMUSG00000038560 | Sp6 | -4.9272817 | 0.01759501 | 0.21506711 |
| ENSMUSG00000026020 | Nop58 | 1.40269492 | 0.01762011 | 0.21517506 |
| ENSMUSG00000074006 | Omp | 2.94445774 | 0.01762078 | 0.21517506 |
| ENSMUSG00000002831 | Plin4 | -2.9900788 | 0.01763027 | 0.21518749 |
| ENSMUSG00000043843 | Tmem145 | -20.259099 | 0.01768493 | 0.21560597 |
| ENSMUSG00000034613 | Ppm1h | 1.48678863 | 0.01769639 | 0.21560597 |
| ENSMUSG00000036752 | Tubb4b | 1.33999101 | 0.01769869 | 0.21560597 |
| ENSMUSG00000026255 | Efhd1 | -4.9982479 | 0.01770864 | 0.21560597 |
| ENSMUSG00000026305 | Lrrfip1 | -1.3209549 | 0.01770948 | 0.21560597 |
| ENSMUSG00000042211 | Fbxo38 | -1.1482044 | 0.01771546 | 0.21560597 |
| ENSMUSG00000039960 | Rhou | 1.41068267 | 0.01773726 | 0.21576798 |
| ENSMUSG00000037731 | Themis2 | -1.5472894 | 0.01777546 | 0.21583588 |
| ENSMUSG00000025132 | Arhgdia | -1.2890216 | 0.01778195 | 0.21583588 |
| ENSMUSG00000024560 | Cxxc1 | 1.20320772 | 0.01778305 | 0.21583588 |
| ENSMUSG00000044017 | Adgrd1 | -2.6842946 | 0.01778319 | 0.21583588 |
| ENSMUSG00000005803 | Sqor | -1.5214602 | 0.01778531 | 0.21583588 |
| ENSMUSG00000019779 | Frk | 1.76953365 | 0.01780275 | 0.21594444 |
| ENSMUSG00000033278 | Ptprm | -1.6941159 | 0.01783098 | 0.21605645 |
| ENSMUSG00000089722 | Cd300ld5 | -2.4063543 | 0.0178371 | 0.21605645 |
| ENSMUSG00000029094 | Afap1 | 1.40375842 | 0.01783749 | 0.21605645 |
| ENSMUSG00000014602 | Kif1a | -2.6274953 | 0.01785404 | 0.2160762 |
| ENSMUSG00000035258 | Abi3bp | -2.7105286 | 0.01785613 | 0.2160762 |
| ENSMUSG00000028194 | Ddah1 | -2.1489141 | 0.01787617 | 0.21621566 |
| ENSMUSG00000026358 | Rgs1 | -3.1548022 | 0.01789276 | 0.21631341 |
| ENSMUSG00000048965 | Mrgpre | -1.9192712 | 0.01790681 | 0.21638028 |
| ENSMUSG00000023918 | Adgrf4 | -8.3824342 | 0.01792759 | 0.21652848 |
| ENSMUSG00000031562 | Dctd | 1.77379589 | 0.01795474 | 0.21675339 |
| ENSMUSG00000026365 | Cfh | -2.3302099 | 0.01797972 | 0.21695189 |
| ENSMUSG00000050272 | Dscam | -5.9414339 | 0.01799683 | 0.21705522 |
| ENSMUSG00000048126 | Col6a3 | -2.1960818 | 0.01800577 | 0.21706006 |
| ENSMUSG00000059439 | Bcas3 | -1.3386279 | 0.01801739 | 0.21709714 |
| ENSMUSG00000017679 | Ttpal | -1.1883362 | 0.01803507 | 0.21720722 |
| ENSMUSG00000031872 | Bean1 | -4.0464929 | 0.01805364 | 0.21732781 |
| ENSMUSG00000030521 | Mphosph10 | 1.37782203 | 0.01806524 | 0.21736455 |
| ENSMUSG00000022102 | Dok2 | -2.0799972 | 0.01816136 | 0.21841761 |

| | | | | |
|---|---|---|---|---|
| ENSMUSG00000016427 | Ndufa1 | -1.3453606 | 0.01818139 | 0.21850806 |
| ENSMUSG00000022181 | C6 | -3.1685029 | 0.01818608 | 0.21850806 |
| ENSMUSG00000027636 | Sla2 | -2.1208307 | 0.01819524 | 0.21851489 |
| ENSMUSG00000039055 | Eme1 | 1.57650782 | 0.01821063 | 0.2185782 |
| ENSMUSG00000079654 | Prrt4 | -8.69076 | 0.01821772 | 0.2185782 |
| ENSMUSG00000032519 | Slc25a38 | 1.29784073 | 0.01825669 | 0.21894246 |
| ENSMUSG00000020218 | Wif1 | -11.009387 | 0.01827768 | 0.21909071 |
| ENSMUSG00000051065 | Mb21d2 | -1.4262855 | 0.01832002 | 0.21944809 |
| ENSMUSG00000031662 | Snx20 | -1.5267152 | 0.01832846 | 0.21944809 |
| ENSMUSG00000045896 | Paip2b | 1.25857441 | 0.0183334 | 0.21944809 |
| ENSMUSG00000038400 | Pmepa1 | -1.9501625 | 0.01836238 | 0.21969158 |
| ENSMUSG00000028394 | Pole3 | 1.25132193 | 0.01838508 | 0.21985964 |
| ENSMUSG00000090124 | Ugt1a7c | -1.6538537 | 0.01839403 | 0.21986316 |
| ENSMUSG00000027950 | Chrnb2 | 1.5460862 | 0.01840451 | 0.21988496 |
| ENSMUSG00000048582 | Gja3 | -72.160516 | 0.01842064 | 0.21997431 |
| ENSMUSG00000028920 | Fbxo42 | 1.19331776 | 0.0184681 | 0.2204375 |
| ENSMUSG00000000126 | Wnt9a | -1.9617197 | 0.01853614 | 0.22114576 |
| ENSMUSG00000080712 | H2bu1-ps | 20.9510974 | 0.01858436 | 0.22161693 |
| ENSMUSG00000048755 | Mcat | 1.2468055 | 0.01862573 | 0.22200614 |
| ENSMUSG00000036617 | Etl4 | 1.62030491 | 0.01865976 | 0.22226835 |
| ENSMUSG00000032352 | Lrrc1 | 1.52602091 | 0.01867011 | 0.22226835 |
| ENSMUSG00000020227 | Irak3 | -1.5137476 | 0.01867397 | 0.22226835 |
| ENSMUSG00000037747 | Phyhipl | -8.5801956 | 0.01870527 | 0.22253662 |
| ENSMUSG00000047613 | A430005L14I | 1.30216063 | 0.01874191 | 0.22286824 |
| ENSMUSG00000100980 | Gm29100 | -2.8806942 | 0.01877974 | 0.22321367 |
| ENSMUSG00000027284 | Cdan1 | 1.41073212 | 0.01883646 | 0.22378318 |
| ENSMUSG00000029195 | Klb | -5.6221966 | 0.01885882 | 0.2239441 |
| ENSMUSG00000014077 | Chp1 | -1.2117258 | 0.01888166 | 0.22411052 |
| ENSMUSG00000059326 | Csf2ra | -1.5403296 | 0.0188988 | 0.22420523 |
| ENSMUSG00000027671 | Actl6a | 1.2332242 | 0.01890728 | 0.22420523 |
| ENSMUSG00000110218 | Gm20219 | 1.85284672 | 0.01893134 | 0.2243858 |
| ENSMUSG00000021716 | Srek1ip1 | 1.19890637 | 0.01896093 | 0.22463176 |
| ENSMUSG00000005615 | Pcyt1a | -1.2752825 | 0.01897428 | 0.22468524 |
| ENSMUSG00000079602 | 9230102O04 | -10.584437 | 0.01900643 | 0.22496101 |
| ENSMUSG00000074037 | Mc1r | -27.659246 | 0.0190298 | 0.22513278 |
| ENSMUSG00000055013 | Agap1 | -1.307625 | 0.01906244 | 0.22541408 |
| ENSMUSG00000025044 | Msr1 | -2.1199657 | 0.01908153 | 0.22553479 |
| ENSMUSG00000027203 | Dut | 1.32088878 | 0.019107 | 0.22573082 |
| ENSMUSG00000050627 | Gpd1l | 1.44118585 | 0.019122 | 0.22575411 |
| ENSMUSG00000031527 | Eri1 | 1.23596391 | 0.01912674 | 0.22575411 |
| ENSMUSG00000075269 | Bex6 | -6.059577 | 0.0191586 | 0.22602519 |
| ENSMUSG00000033790 | Tubgcp5 | 1.28386007 | 0.01919467 | 0.22634569 |
| ENSMUSG00000096422 | Igkv12-44 | -24.430023 | 0.01921612 | 0.22649354 |
| ENSMUSG00000040549 | Ckap5 | 1.30949703 | 0.01922726 | 0.22651973 |
| ENSMUSG00000024600 | Slc27a6 | -4.6670762 | 0.01927171 | 0.22693822 |

| | | | | |
|---|---|---|---|---|
| ENSMUSG00000054321 | Taf4b | 1.44022529 | 0.01929849 | 0.22714829 |
| ENSMUSG00000018648 | Dusp14 | -2.0376482 | 0.01940703 | 0.22822468 |
| ENSMUSG00000035958 | Tdp2 | 1.2009581 | 0.01940939 | 0.22822468 |
| ENSMUSG00000027293 | Ehd4 | -1.3414203 | 0.01941688 | 0.22822468 |
| ENSMUSG00000027010 | Slc25a12 | -1.3208575 | 0.01943073 | 0.22828193 |
| ENSMUSG00000004460 | Dnajb11 | -1.2695825 | 0.01944658 | 0.22836247 |
| ENSMUSG00000026854 | Usp20 | 1.33348605 | 0.01947914 | 0.22863923 |
| ENSMUSG00000026249 | Serpine2 | -1.9928225 | 0.01948852 | 0.2286437 |
| ENSMUSG00000039952 | Dag1 | 1.50765479 | 0.01950257 | 0.22870292 |
| ENSMUSG00000052584 | Serp2 | -2.8483934 | 0.01953893 | 0.22891006 |
| ENSMUSG00000022346 | Myc | 1.6368973 | 0.01954861 | 0.22891006 |
| ENSMUSG00000014039 | Prdm15 | 1.40424752 | 0.01955052 | 0.22891006 |
| ENSMUSG00000034285 | Nipsnap1 | -1.36255 | 0.01955626 | 0.22891006 |
| ENSMUSG00000023078 | Cxcl13 | -4.1566873 | 0.0195936 | 0.22919694 |
| ENSMUSG00000027583 | Zbtb46 | 1.48964733 | 0.01959881 | 0.22919694 |
| ENSMUSG00000022246 | Rai14 | -1.4906545 | 0.01962149 | 0.22935667 |
| ENSMUSG00000104876 | Trdc | -3.9751676 | 0.01967271 | 0.22984956 |
| ENSMUSG00000024173 | Tpsab1 | -11.792107 | 0.01970163 | 0.23008167 |
| ENSMUSG00000095079 | Igha | -6.2837731 | 0.01976358 | 0.23069918 |
| ENSMUSG00000111212 | Gm47087 | 5.0747582 | 0.01978955 | 0.2308962 |
| ENSMUSG00000032089 | Il10ra | -1.520921 | 0.01987953 | 0.23179091 |
| ENSMUSG00000057329 | Bcl2 | -1.523186 | 0.01988447 | 0.23179091 |
| ENSMUSG00000047658 | Gal3st3 | 42.5151581 | 0.01989918 | 0.23185606 |
| ENSMUSG00000029028 | Lrrc47 | 1.2653112 | 0.01994347 | 0.23216257 |
| ENSMUSG00000079492 | Gm11127 | -10.124997 | 0.01995158 | 0.23216257 |
| ENSMUSG00000026600 | Soat1 | -1.4518902 | 0.01995711 | 0.23216257 |
| ENSMUSG00000042834 | Nrep | 1.70781822 | 0.01996203 | 0.23216257 |
| ENSMUSG00000023883 | Phf10 | 1.22194942 | 0.01999537 | 0.23234003 |
| ENSMUSG00000022280 | Rnf19a | 1.35499678 | 0.01999558 | 0.23234003 |
| ENSMUSG00000027763 | Mbnl1 | -1.339263 | 0.02005091 | 0.23287652 |
| ENSMUSG00000069806 | Cacng7 | 3.10203844 | 0.02007106 | 0.23300401 |
| ENSMUSG00000049115 | Agtr1a | 2.11377252 | 0.02013391 | 0.23362686 |
| ENSMUSG00000089728 | Clec2f | 3.5924564 | 0.02015202 | 0.23373036 |
| ENSMUSG00000004044 | Cavin1 | -1.5053489 | 0.02020292 | 0.23412741 |
| ENSMUSG00000021266 | Wars | -1.2529071 | 0.02021703 | 0.23412741 |
| ENSMUSG00000024897 | Apba1 | -1.8323966 | 0.02022502 | 0.23412741 |
| ENSMUSG00000068744 | Psrc1 | 1.94243448 | 0.02023588 | 0.23412741 |
| ENSMUSG00000006218 | Fam131c | 2.08472453 | 0.02023736 | 0.23412741 |
| ENSMUSG00000036591 | Arhgap21 | 1.28566508 | 0.02024154 | 0.23412741 |
| ENSMUSG00000046844 | Vat1l | -4.2733853 | 0.0202873 | 0.23445207 |
| ENSMUSG00000032745 | Gpbp1 | 1.20038166 | 0.02028937 | 0.23445207 |
| ENSMUSG00000026344 | Lypd1 | -5.7370776 | 0.02030326 | 0.23445207 |
| ENSMUSG00000038009 | Dnajc22 | 2.58320028 | 0.02030651 | 0.23445207 |
| ENSMUSG00000046245 | Pilra | -1.6414077 | 0.02033015 | 0.23453831 |
| ENSMUSG00000050199 | Lgr4 | -1.4246063 | 0.02033244 | 0.23453831 |

| | | | | |
|---|---|---|---|---|
| ENSMUSG00000059493 | Nhs | -2.4884419 | 0.02035153 | 0.23465207 |
| ENSMUSG00000000957 | Mmp14 | -1.4904716 | 0.02038225 | 0.23488101 |
| ENSMUSG00000030523 | Trpm1 | -8.7088312 | 0.02039335 | 0.23488101 |
| ENSMUSG00000022225 | Cma1 | -2.7997091 | 0.02040763 | 0.23488101 |
| ENSMUSG00000110462 | Gm35572 | 3.77284803 | 0.02040836 | 0.23488101 |
| ENSMUSG00000070610 | Gm13127 | 1.93613403 | 0.02041797 | 0.23488517 |
| ENSMUSG00000054920 | Klhl5 | -1.4319058 | 0.02043848 | 0.2350075 |
| ENSMUSG00000052415 | Tchh | -10.852621 | 0.0204471 | 0.2350075 |
| ENSMUSG00000023905 | Tnfrsf12a | -1.6621911 | 0.02050483 | 0.23556455 |
| ENSMUSG00000037411 | Serpine1 | -3.3554532 | 0.02051971 | 0.23562897 |
| ENSMUSG00000026548 | Slamf9 | -1.7978868 | 0.02054369 | 0.23570158 |
| ENSMUSG00000026933 | Camsap1 | 1.30551002 | 0.02054459 | 0.23570158 |
| ENSMUSG00000027858 | Tspan2 | 1.57870519 | 0.02055716 | 0.23573614 |
| ENSMUSG00000074358 | Ccdc61 | 1.34407174 | 0.02056615 | 0.23573614 |
| ENSMUSG00000008575 | Nfib | 1.40609898 | 0.02058163 | 0.23580723 |
| ENSMUSG00000095351 | Igkv3-2 | -371.13641 | 0.02060015 | 0.23591308 |
| ENSMUSG00000001025 | S100a6 | -1.647103 | 0.02069842 | 0.23683095 |
| ENSMUSG00000031950 | Gabarapl2 | -1.2219877 | 0.02069894 | 0.23683095 |
| ENSMUSG00000040354 | Mars1 | 1.18855052 | 0.02071921 | 0.23684483 |
| ENSMUSG00000016256 | Ctsz | -1.3598067 | 0.02072627 | 0.23684483 |
| ENSMUSG00000031601 | Cnot7 | 1.24123076 | 0.02072812 | 0.23684483 |
| ENSMUSG00000037710 | Cisd1 | 1.29796788 | 0.02074205 | 0.23689748 |
| ENSMUSG00000092178 | Gm45351 | 2.12278458 | 0.02076742 | 0.23708069 |
| ENSMUSG00000026193 | Fn1 | -3.0487965 | 0.02082644 | 0.23764777 |
| ENSMUSG00000031561 | Tenm3 | -5.269145 | 0.02083838 | 0.2376773 |
| ENSMUSG00000045775 | Slc16a5 | -2.2942621 | 0.02086747 | 0.23785821 |
| ENSMUSG00000029177 | Cenpa | 1.60916962 | 0.02087548 | 0.23785821 |
| ENSMUSG00000054302 | Eapp | -1.2767893 | 0.0208884 | 0.23785821 |
| ENSMUSG00000026678 | Rgs5 | -1.7419518 | 0.02089169 | 0.23785821 |
| ENSMUSG00000030214 | Plbd1 | -1.4695023 | 0.0209205 | 0.23807959 |
| ENSMUSG00000045176 | Borcs6 | -1.3727531 | 0.02093355 | 0.23812145 |
| ENSMUSG00000021635 | Rad17 | 1.17621412 | 0.02098558 | 0.23860656 |
| ENSMUSG00000030613 | Ccdc90b | 1.32837541 | 0.02100593 | 0.23867722 |
| ENSMUSG00000023075 | Akirin1 | -1.1992322 | 0.02101058 | 0.23867722 |
| ENSMUSG00000026773 | Pfkfb3 | -1.8646541 | 0.0210207 | 0.23868548 |
| ENSMUSG00000097467 | Gm26737 | 3.58041154 | 0.02103011 | 0.23868562 |
| ENSMUSG00000071724 | Smpd5 | -1.8589503 | 0.02107266 | 0.23906185 |
| ENSMUSG00000058589 | Anks1b | -4.6245695 | 0.02109249 | 0.23918004 |
| ENSMUSG00000044224 | Dnajc21 | 1.20075515 | 0.02112669 | 0.2394609 |
| ENSMUSG00000024135 | Srbd1 | 1.21518138 | 0.02114719 | 0.2395864 |
| ENSMUSG00000031483 | Erlin2 | 1.20893121 | 0.02116001 | 0.23961806 |
| ENSMUSG00000062783 | Csprs | 2.50260773 | 0.02116884 | 0.23961806 |
| ENSMUSG00000019194 | Scn1b | -1.5750113 | 0.02118168 | 0.23964374 |
| ENSMUSG00000034826 | Nup54 | 1.27472997 | 0.02120148 | 0.23964374 |
| ENSMUSG00000023906 | Cldn6 | 6.37894917 | 0.02120422 | 0.23964374 |

| | | | | |
|---|---|---|---|---|
| ENSMUSG00000028480 | Glipr2 | -1.604877 | 0.02120883 | 0.23964374 |
| ENSMUSG00000020255 | D10Wsu102e | 1.27739262 | 0.02125446 | 0.24005255 |
| ENSMUSG00000076508 | Igkv17-127 | -25.4621 | 0.02127259 | 0.24010011 |
| ENSMUSG00000029513 | Prkab1 | 1.37466952 | 0.02127756 | 0.24010011 |
| ENSMUSG00000064023 | Klk8 | -3.1583907 | 0.02130119 | 0.24025998 |
| ENSMUSG00000025348 | Itga7 | -1.9052339 | 0.02132214 | 0.2403153 |
| ENSMUSG00000059409 | Ppp2r5d | 1.21431616 | 0.021325 | 0.2403153 |
| ENSMUSG00000035704 | Alg8 | 1.3578413 | 0.02134067 | 0.24038527 |
| ENSMUSG00000045410 | Akr1e1 | 1.35358936 | 0.02136532 | 0.24055625 |
| ENSMUSG00000038319 | Kcnh2 | -2.1216947 | 0.02137688 | 0.24057989 |
| ENSMUSG00000055184 | Fam72a | 1.81516191 | 0.02140213 | 0.24065236 |
| ENSMUSG00000002996 | Hbp1 | -1.2290946 | 0.02140226 | 0.24065236 |
| ENSMUSG00000084883 | Ccdc85c | 1.28481563 | 0.02144745 | 0.24094931 |
| ENSMUSG00000047248 | C2cd3 | 1.25237456 | 0.02145379 | 0.24094931 |
| ENSMUSG00000003778 | Brd8 | 1.31237758 | 0.02146507 | 0.24094931 |
| ENSMUSG00000009356 | Lpo | 11.2030442 | 0.02146678 | 0.24094931 |
| ENSMUSG00000113836 | Gm3325 | 1.78036277 | 0.02148673 | 0.24094931 |
| ENSMUSG00000094724 | Rnaset2b | -1.3650834 | 0.02148724 | 0.24094931 |
| ENSMUSG00000035799 | Twist1 | -1.8133272 | 0.02149505 | 0.24094931 |
| ENSMUSG00000037286 | Stag1 | 1.21560155 | 0.02155422 | 0.2415061 |
| ENSMUSG00000039738 | Slx4 | 1.27092951 | 0.02157339 | 0.24161441 |
| ENSMUSG00000053289 | Ddx10 | 1.23870796 | 0.02162341 | 0.24204747 |
| ENSMUSG00000037224 | Zfyve28 | -2.7692816 | 0.02163111 | 0.24204747 |
| ENSMUSG00000069910 | Spdl1 | 1.46935378 | 0.02168741 | 0.24257062 |
| ENSMUSG00000025557 | Slc15a1 | 4.55977142 | 0.02173665 | 0.24292809 |
| ENSMUSG00000027984 | Hadh | 1.25363419 | 0.02173849 | 0.24292809 |
| ENSMUSG00000007827 | Ankrd26 | 1.44824178 | 0.02178947 | 0.2433908 |
| ENSMUSG00000047434 | Xxylt1 | -1.2244628 | 0.02185839 | 0.24391807 |
| ENSMUSG00000028469 | Npr2 | -1.9881792 | 0.0218607 | 0.24391807 |
| ENSMUSG00000115426 | Gm19510 | -18.540414 | 0.02187877 | 0.24391807 |
| ENSMUSG00000026662 | Sephs1 | 1.20472407 | 0.02188249 | 0.24391807 |
| ENSMUSG00000025049 | Taf5 | 1.41064348 | 0.02188967 | 0.24391807 |
| ENSMUSG00000032172 | Olfm2 | -4.1058537 | 0.02189591 | 0.24391807 |
| ENSMUSG00000038695 | Josd2 | 1.44729814 | 0.02191784 | 0.24391807 |
| ENSMUSG00000039307 | Hexdc | 1.42306197 | 0.02192237 | 0.24391807 |
| ENSMUSG00000009549 | Srp14 | -1.1917006 | 0.02192306 | 0.24391807 |
| ENSMUSG00000021916 | Glt8d1 | -1.2470037 | 0.02195034 | 0.24406915 |
| ENSMUSG00000085667 | Gm12992 | 2.07288023 | 0.0219653 | 0.24406915 |
| ENSMUSG00000027863 | Cd2 | -2.5319532 | 0.02196546 | 0.24406915 |
| ENSMUSG00000030203 | Dusp16 | 1.37239347 | 0.02199731 | 0.2443163 |
| ENSMUSG00000117869 | Snhg4 | 1.77726251 | 0.02205814 | 0.24487555 |
| ENSMUSG00000016253 | Nelfcd | 1.21234239 | 0.02206694 | 0.24487555 |
| ENSMUSG00000032525 | Nktr | 1.53306649 | 0.02214829 | 0.245671 |
| ENSMUSG00000030584 | Dpf1 | -2.0187996 | 0.02217369 | 0.24584542 |
| ENSMUSG00000081723 | Gm15931 | -2.7647948 | 0.02221565 | 0.2461731 |

| | | | | |
|---|---|---|---|---|
| ENSMUSG00000004233 | Wars2 | 1.39537926 | 0.02222261 | 0.2461731 |
| ENSMUSG00000039037 | St6galnac5 | -7.2959538 | 0.02232558 | 0.24720589 |
| ENSMUSG00000027509 | Rae1 | 1.23448293 | 0.02234363 | 0.24729801 |
| ENSMUSG00000034973 | Dop1a | 1.42890992 | 0.0223738 | 0.24752422 |
| ENSMUSG00000020311 | Erlec1 | -1.2851327 | 0.02240109 | 0.24762327 |
| ENSMUSG00000060206 | Zfp462 | 1.4296011 | 0.02240225 | 0.24762327 |
| ENSMUSG00000041459 | Tardbp | 1.32374419 | 0.02242499 | 0.24776686 |
| ENSMUSG00000058444 | Map2k5 | 1.24242404 | 0.02246426 | 0.24809285 |
| ENSMUSG00000021124 | Vti1b | -1.2715588 | 0.02250048 | 0.24838491 |
| ENSMUSG00000013707 | Tnfaip8l2 | -1.549382 | 0.02255088 | 0.24877071 |
| ENSMUSG00000113637 | Gm7049 | -13.744809 | 0.022555 | 0.24877071 |
| ENSMUSG00000078606 | Gm4070 | -1.6580553 | 0.02258072 | 0.24888392 |
| ENSMUSG00000030621 | Me3 | -2.2394626 | 0.02260213 | 0.24888392 |
| ENSMUSG00000109511 | Nup62 | 1.34925404 | 0.02261285 | 0.24888392 |
| ENSMUSG00000032803 | Cdv3 | 1.18632074 | 0.02262543 | 0.24888392 |
| ENSMUSG00000062421 | Arf2 | -1.2429064 | 0.02262695 | 0.24888392 |
| ENSMUSG00000039086 | Ss18l1 | 1.48190256 | 0.02263323 | 0.24888392 |
| ENSMUSG00000024084 | Qpct | -1.659489 | 0.02264296 | 0.24888392 |
| ENSMUSG00000031988 | Vps26b | 1.20915719 | 0.02265396 | 0.24888392 |
| ENSMUSG00000020946 | Gosr2 | -1.1943965 | 0.02266153 | 0.24888392 |
| ENSMUSG00000014852 | Adamts13 | 135.229398 | 0.02267943 | 0.24888392 |
| ENSMUSG00000029034 | Ints11 | 1.28527061 | 0.02268172 | 0.24888392 |
| ENSMUSG00000060671 | Atp8b2 | -1.2839522 | 0.02268279 | 0.24888392 |
| ENSMUSG00000001018 | Snapin | -1.208579 | 0.02275135 | 0.2495284 |
| ENSMUSG00000023328 | Ache | -4.7088724 | 0.02280701 | 0.24998337 |
| ENSMUSG00000022765 | Snap29 | -1.1661924 | 0.02281699 | 0.24998337 |
| ENSMUSG00000022489 | Pde1b | -1.9781779 | 0.02282234 | 0.24998337 |
| ENSMUSG00000030218 | Mgp | -2.183793 | 0.02285819 | 0.25018527 |
| ENSMUSG00000031060 | Rbm10 | 1.1980972 | 0.02286047 | 0.25018527 |
| ENSMUSG00000102145 | Gm38056 | -3.5624877 | 0.02289947 | 0.25036758 |
| ENSMUSG00000045658 | Pid1 | -1.7364951 | 0.0229116 | 0.25036758 |
| ENSMUSG00000074527 | Gm14296 | 2.53723265 | 0.02292314 | 0.25036758 |
| ENSMUSG00000039321 | Uts2r | 19.8462768 | 0.02292525 | 0.25036758 |
| ENSMUSG00000019737 | Syne4 | 2.5078393 | 0.02292639 | 0.25036758 |
| ENSMUSG00000015890 | Amdhd1 | -4.4206119 | 0.02299288 | 0.25090778 |
| ENSMUSG00000048106 | 4632415L05F | 1.29280649 | 0.02301478 | 0.25090778 |
| ENSMUSG00000049608 | Gpr55 | -2.1143035 | 0.02301497 | 0.25090778 |
| ENSMUSG00000024169 | Ift140 | 1.20306315 | 0.02302234 | 0.25090778 |
| ENSMUSG00000034187 | Nsf | -1.2470118 | 0.02302522 | 0.25090778 |
| ENSMUSG00000036002 | Fam214b | -1.5202158 | 0.0230837 | 0.25143715 |
| ENSMUSG00000026842 | Abl1 | 1.25114474 | 0.02315633 | 0.25212024 |
| ENSMUSG00000020368 | Canx | -1.1627703 | 0.02317577 | 0.25222388 |
| ENSMUSG00000113679 | Gm10432 | -4.7356231 | 0.02319876 | 0.25236596 |
| ENSMUSG00000050994 | Adgb | 4.18410351 | 0.0232153 | 0.25241295 |
| ENSMUSG00000058638 | Zfp110 | 1.1708649 | 0.02322664 | 0.25241295 |

| | | | | |
|---|---|---|---|---|
| ENSMUSG00000024538 | Ppic | -1.4583374 | 0.02323288 | 0.25241295 |
| ENSMUSG00000071073 | Lrrc73 | 1.8169914 | 0.0232795 | 0.2528114 |
| ENSMUSG00000042439 | Zfp532 | 1.31694511 | 0.02338177 | 0.25381357 |
| ENSMUSG00000063382 | Bcl9l | 1.32864685 | 0.02340223 | 0.25392715 |
| ENSMUSG00000017754 | Pltp | -1.4005099 | 0.02343413 | 0.25416481 |
| ENSMUSG00000056071 | S100a9 | -6.3123751 | 0.02349307 | 0.25469539 |
| ENSMUSG00000040554 | Aipl1 | -3.6355937 | 0.02350922 | 0.25476177 |
| ENSMUSG00000035020 | Epgn | -4.3664685 | 0.02353429 | 0.25492475 |
| ENSMUSG00000051639 | Fbl-ps2 | 3.92722922 | 0.02359485 | 0.25531424 |
| ENSMUSG00000092544 | Gm20422 | 19.2844336 | 0.02359928 | 0.25531424 |
| ENSMUSG00000051786 | Tubgcp6 | 1.25239436 | 0.02360039 | 0.25531424 |
| ENSMUSG00000040118 | Cacna2d1 | -1.8060937 | 0.02364547 | 0.25569305 |
| ENSMUSG00000022887 | Masp1 | -2.8712915 | 0.02367152 | 0.25585382 |
| ENSMUSG00000106665 | Gm43389 | 5.65558191 | 0.02368539 | 0.25585382 |
| ENSMUSG00000001506 | Col1a1 | -2.8540408 | 0.02369054 | 0.25585382 |
| ENSMUSG00000079429 | Mroh2a | 1.812414 | 0.02371073 | 0.25596303 |
| ENSMUSG00000051041 | Olfml1 | -4.1817111 | 0.0237282 | 0.25597915 |
| ENSMUSG00000000142 | Axin2 | -2.8358733 | 0.02373237 | 0.25597915 |
| ENSMUSG00000117499 | Gm50010 | 1.75229449 | 0.02374687 | 0.25602695 |
| ENSMUSG00000037098 | Rab11fip3 | 1.22923311 | 0.02378292 | 0.25630688 |
| ENSMUSG00000042305 | Tmem183a | 1.2406139 | 0.02379828 | 0.25636365 |
| ENSMUSG00000034593 | Myo5a | -1.5404289 | 0.02381155 | 0.25639795 |
| ENSMUSG00000032724 | Abtb2 | 1.44337636 | 0.02386032 | 0.25681423 |
| ENSMUSG00000027080 | Med19 | 1.20267522 | 0.02388515 | 0.25696069 |
| ENSMUSG00000032327 | Stra6 | -6.0625493 | 0.02389802 | 0.25696069 |
| ENSMUSG00000057147 | Dph6 | 1.20184639 | 0.02390766 | 0.25696069 |
| ENSMUSG00000095620 | Csta2 | 15.968747 | 0.02391437 | 0.25696069 |
| ENSMUSG00000094420 | Igkv10-96 | -24.414179 | 0.02399031 | 0.2575077 |
| ENSMUSG00000031586 | Rbpms | 1.86498272 | 0.0239905 | 0.2575077 |
| ENSMUSG00000029504 | Ddx51 | 1.19224282 | 0.02399968 | 0.2575077 |
| ENSMUSG00000031447 | Lamp1 | -1.1585285 | 0.02400581 | 0.2575077 |
| ENSMUSG00000026708 | Cenpl | 1.39732751 | 0.02402778 | 0.25763453 |
| ENSMUSG00000045534 | Kcna5 | -3.6957606 | 0.02403995 | 0.25765635 |
| ENSMUSG00000027423 | Snx5 | -1.1646278 | 0.02408108 | 0.25798839 |
| ENSMUSG00000024030 | Abcg1 | -1.6387717 | 0.02411876 | 0.2582832 |
| ENSMUSG00000015087 | Rabl6 | 1.15868346 | 0.02423864 | 0.25942657 |
| ENSMUSG00000061613 | U2af1 | 1.23444862 | 0.02425418 | 0.25942657 |
| ENSMUSG00000021193 | Pitrm1 | 1.24027712 | 0.02425616 | 0.25942657 |
| ENSMUSG00000026893 | Gca | 1.760594 | 0.02433051 | 0.26001322 |
| ENSMUSG00000046157 | Tmem229b | -1.4386425 | 0.02433148 | 0.26001322 |
| ENSMUSG00000026068 | Il18rap | -1.8324224 | 0.02436318 | 0.26024255 |
| ENSMUSG00000017724 | Etv4 | -2.1972812 | 0.02437692 | 0.26027992 |
| ENSMUSG00000035696 | Rnf38 | 1.28629012 | 0.02441378 | 0.26052052 |
| ENSMUSG00000018287 | Spag7 | -1.2287665 | 0.02441995 | 0.26052052 |
| ENSMUSG00000021903 | Galnt15 | -2.2251762 | 0.02447249 | 0.26097141 |

| | | | | |
|---|---|---|---|---|
| ENSMUSG00000100009 | Gm7967 | 5.37043214 | 0.02449999 | 0.26115508 |
| ENSMUSG00000003279 | Dlgap1 | -4.4621104 | 0.02451433 | 0.26119837 |
| ENSMUSG00000058914 | C1qtnf3 | -2.4751726 | 0.02454783 | 0.26137447 |
| ENSMUSG00000031400 | G6pdx | -1.3097735 | 0.02455143 | 0.26137447 |
| ENSMUSG00000021760 | Gpx8 | -1.463797 | 0.02459991 | 0.26178094 |
| ENSMUSG00000021532 | Fastkd3 | 1.29985419 | 0.02471873 | 0.26293527 |
| ENSMUSG00000029309 | Sparcl1 | -1.9357194 | 0.02473499 | 0.26298433 |
| ENSMUSG00000018507 | Trpv2 | -1.51018 | 0.02474404 | 0.26298433 |
| ENSMUSG00000074357 | AA386476 | 2.95231674 | 0.02475552 | 0.26299635 |
| ENSMUSG00000107008 | Gm2762 | 5.18769316 | 0.02477224 | 0.26306399 |
| ENSMUSG00000048696 | Mex3d | 1.35454201 | 0.02479798 | 0.26322739 |
| ENSMUSG00000047798 | Cd300lf | -1.7534594 | 0.02483712 | 0.26353273 |
| ENSMUSG00000023048 | Prr13 | -1.3129599 | 0.02484887 | 0.26353752 |
| ENSMUSG00000047719 | Ubiad1 | 1.23135206 | 0.02485831 | 0.26353752 |
| ENSMUSG00000040848 | Sft2d2 | 1.2639972 | 0.02490619 | 0.26378697 |
| ENSMUSG00000040321 | Zfp770 | 1.39966495 | 0.0249071 | 0.26378697 |
| ENSMUSG00000052031 | Tagap1 | 1.4450385 | 0.02491298 | 0.26378697 |
| ENSMUSG00000038759 | Nup205 | 1.37669308 | 0.02494317 | 0.2638798 |
| ENSMUSG00000052833 | Sae1 | 1.24029617 | 0.02495451 | 0.2638798 |
| ENSMUSG00000030602 | Pak4 | 1.26032327 | 0.02496696 | 0.2638798 |
| ENSMUSG00000076498 | Trbc2 | -2.3037682 | 0.02496948 | 0.2638798 |
| ENSMUSG00000024771 | Lipk | 20.8883465 | 0.02497367 | 0.2638798 |
| ENSMUSG00000015854 | Cd5l | 4.62279948 | 0.02502131 | 0.26427333 |
| ENSMUSG00000053040 | Aph1c | -1.9022554 | 0.02507811 | 0.26476318 |
| ENSMUSG00000072653 | Zfp783 | 4.57751323 | 0.02514803 | 0.26536786 |
| ENSMUSG00000022754 | Tmem45a | -2.7361817 | 0.02515627 | 0.26536786 |
| ENSMUSG00000001665 | Gstt3 | 1.83549859 | 0.02522865 | 0.26597183 |
| ENSMUSG00000028224 | Nbn | 1.22895988 | 0.02523446 | 0.26597183 |
| ENSMUSG00000040681 | Hmgn1 | 1.18895673 | 0.02525409 | 0.26606841 |
| ENSMUSG00000028195 | Ccn1 | -2.1786467 | 0.02530823 | 0.26641821 |
| ENSMUSG00000070547 | Mrgprb1 | -3.9636246 | 0.02530826 | 0.26641821 |
| ENSMUSG00000026770 | Il2ra | -2.6401023 | 0.02532369 | 0.26647027 |
| ENSMUSG00000020785 | Camkk1 | 1.9339497 | 0.02535247 | 0.26666269 |
| ENSMUSG00000025237 | Parp6 | 1.29694479 | 0.02539106 | 0.26693084 |
| ENSMUSG00000097081 | Gm10425 | 2.23420829 | 0.02539898 | 0.26693084 |
| ENSMUSG00000062905 | Vmn1r32 | -89.44136 | 0.02543667 | 0.26721651 |
| ENSMUSG00000028551 | Cdkn2c | 1.50895433 | 0.02549072 | 0.26767361 |
| ENSMUSG00000020793 | Galr2 | -2.4926188 | 0.02550836 | 0.26774821 |
| ENSMUSG00000022390 | Zc3h7b | 1.32512465 | 0.02554147 | 0.26798508 |
| ENSMUSG00000023349 | Clec4n | -1.8555319 | 0.02564672 | 0.26897836 |
| ENSMUSG00000061589 | Dot1l | 1.32741845 | 0.02570382 | 0.269466 |
| ENSMUSG00000008035 | Mid1ip1 | -1.4766587 | 0.02583207 | 0.27069882 |
| ENSMUSG00000037437 | Adam32 | -18.973842 | 0.02585562 | 0.27071088 |
| ENSMUSG00000018379 | Srsf1 | 1.23486793 | 0.02587007 | 0.27071088 |
| ENSMUSG00000022206 | Npr3 | -2.9100838 | 0.02588436 | 0.27071088 |

| | | | | |
|---|---|---|---|---|
| ENSMUSG00000034269 | Setd5 | 1.2783305 | 0.0258891 | 0.27071088 |
| ENSMUSG00000019845 | Tube1 | 1.61223363 | 0.02589146 | 0.27071088 |
| ENSMUSG00000118346 | Tmem179b | -1.3426506 | 0.02589714 | 0.27071088 |
| ENSMUSG00000042737 | Dpm3 | -1.3582361 | 0.02595503 | 0.27120444 |
| ENSMUSG00000070424 | Art5 | 4.25266414 | 0.02601261 | 0.27169443 |
| ENSMUSG00000038421 | Fcrla | -7.5073934 | 0.02606281 | 0.27206158 |
| ENSMUSG00000097119 | B230354K17 | 1.40712632 | 0.02606918 | 0.27206158 |
| ENSMUSG00000001864 | Aif1l | 2.73208753 | 0.02608791 | 0.27214528 |
| ENSMUSG00000032396 | Dis3l | 1.17709057 | 0.02615086 | 0.27269002 |
| ENSMUSG00000036983 | Tfb1m | 1.35427262 | 0.02617757 | 0.27285659 |
| ENSMUSG00000049241 | Hcar1 | -4.1131765 | 0.02618937 | 0.27286766 |
| ENSMUSG00000006442 | Srm | 1.52331334 | 0.02621474 | 0.27302008 |
| ENSMUSG00000110344 | Gm45716 | -2.4560912 | 0.02623753 | 0.27308349 |
| ENSMUSG00000118495 | AC161757.1 | -7.339797 | 0.02624232 | 0.27308349 |
| ENSMUSG00000034460 | Six4 | 1.53248573 | 0.02630892 | 0.27366446 |
| ENSMUSG00000050335 | Lgals3 | -2.0487609 | 0.02634777 | 0.27367858 |
| ENSMUSG00000020151 | Ptprr | -17.29258 | 0.02635562 | 0.27367858 |
| ENSMUSG00000024277 | Mapre2 | -1.691394 | 0.02635572 | 0.27367858 |
| ENSMUSG00000000320 | Alox12 | -4.5481499 | 0.02635744 | 0.27367858 |
| ENSMUSG00000021810 | Ecd | 1.18817167 | 0.02637704 | 0.27367858 |
| ENSMUSG00000030560 | Ctsc | -1.4754647 | 0.02639239 | 0.27367858 |
| ENSMUSG00000038121 | Fam210a | 1.28622153 | 0.02639421 | 0.27367858 |
| ENSMUSG00000030643 | Rab30 | 1.60279826 | 0.02639644 | 0.27367858 |
| ENSMUSG00000046785 | Epm2aip1 | 1.49350272 | 0.02643027 | 0.2739176 |
| ENSMUSG00000024855 | Pacs1 | 1.4041717 | 0.02646865 | 0.27420359 |
| ENSMUSG00000025257 | Ribc1 | 1.74387955 | 0.0265176 | 0.27459876 |
| ENSMUSG00000005225 | Plekha8 | 1.47628163 | 0.02652842 | 0.27459891 |
| ENSMUSG00000027347 | Rasgrp1 | -1.7357661 | 0.02657197 | 0.27477777 |
| ENSMUSG00000021794 | Glud1 | -1.2171269 | 0.0265751 | 0.27477777 |
| ENSMUSG00000110630 | K230015D01 | -2.3806362 | 0.02657814 | 0.27477777 |
| ENSMUSG00000078954 | Arhgap8 | 1.7846009 | 0.02659351 | 0.27482481 |
| ENSMUSG00000074151 | Nlrc5 | 1.41098498 | 0.0266187 | 0.2749425 |
| ENSMUSG00000045374 | Wdr81 | -1.2646027 | 0.02662653 | 0.2749425 |
| ENSMUSG00000034807 | Colgalt1 | 1.23082522 | 0.0266553 | 0.27501838 |
| ENSMUSG00000097211 | BC065403 | -5.7729963 | 0.02666109 | 0.27501838 |
| ENSMUSG00000041688 | Amot | 1.97776548 | 0.02666635 | 0.27501838 |
| ENSMUSG00000021996 | Esd | -1.3296033 | 0.02672016 | 0.27546159 |
| ENSMUSG00000056271 | Lman1l | 15.1814977 | 0.02674304 | 0.27558561 |
| ENSMUSG00000040312 | Cchcr1 | 1.34421384 | 0.02675729 | 0.27562071 |
| ENSMUSG00000030168 | Adipor2 | -1.2615425 | 0.02677004 | 0.27564031 |
| ENSMUSG00000042351 | Grap2 | -1.7113099 | 0.02678246 | 0.27565648 |
| ENSMUSG00000035305 | Ror1 | 1.85542713 | 0.02679343 | 0.27565779 |
| ENSMUSG00000036022 | Fam122b | 1.47346727 | 0.02680806 | 0.27569669 |
| ENSMUSG00000029076 | Sdf4 | -1.2411747 | 0.02683429 | 0.27585472 |
| ENSMUSG00000095432 | Zfp748 | 1.29149614 | 0.0268801 | 0.27621395 |

| | | | | |
|---|---|---|---|---|
| ENSMUSG00000038010 | Ccdc138 | 1.58421485 | 0.02689734 | 0.27627937 |
| ENSMUSG00000078616 | Trim30c | -2.5735362 | 0.02695281 | 0.27673728 |
| ENSMUSG00000039768 | Dnajc11 | 1.23270685 | 0.02696587 | 0.27675956 |
| ENSMUSG00000078921 | Tgtp2 | -1.8401802 | 0.02697732 | 0.27676529 |
| ENSMUSG00000036309 | Skp1a | -1.2380059 | 0.02699242 | 0.27680843 |
| ENSMUSG00000043531 | Sorcs1 | -3.9624014 | 0.02705331 | 0.277321 |
| ENSMUSG00000039680 | Mrps6 | 1.33651916 | 0.0271015 | 0.27770296 |
| ENSMUSG00000021978 | Extl3 | -1.2324891 | 0.02711757 | 0.27773303 |
| ENSMUSG00000026072 | Il1r1 | -1.768871 | 0.02712629 | 0.27773303 |
| ENSMUSG00000007279 | Scube2 | -3.1862907 | 0.02717347 | 0.27802388 |
| ENSMUSG00000036202 | Rif1 | 1.41875726 | 0.02718037 | 0.27802388 |
| ENSMUSG00000013663 | Pten | -1.3433925 | 0.02718752 | 0.27802388 |
| ENSMUSG00000022477 | Aco2 | 1.15409193 | 0.02722964 | 0.27834252 |
| ENSMUSG00000015869 | Prpsap1 | 1.2387267 | 0.02731262 | 0.27907855 |
| ENSMUSG00000017718 | Afmid | -1.4813402 | 0.02737668 | 0.27962065 |
| ENSMUSG00000030357 | Fkbp4 | 1.35569563 | 0.02741002 | 0.27984874 |
| ENSMUSG00000039704 | Lmbrd2 | 1.3443347 | 0.02744047 | 0.27999455 |
| ENSMUSG00000050708 | Ftl1 | -1.4760789 | 0.02745582 | 0.27999455 |
| ENSMUSG00000005374 | Tbl2 | 1.23879275 | 0.02745736 | 0.27999455 |
| ENSMUSG00000060733 | Ipmk | 1.19296162 | 0.02747848 | 0.28009749 |
| ENSMUSG00000026442 | Nfasc | -6.1119535 | 0.02749476 | 0.28015111 |
| ENSMUSG00000012483 | Rpa3 | 1.41539115 | 0.02752766 | 0.28029546 |
| ENSMUSG00000048186 | Bend7 | 1.97926903 | 0.02754092 | 0.28029546 |
| ENSMUSG00000004085 | Map3k20 | 1.19314776 | 0.02754202 | 0.28029546 |
| ENSMUSG00000042453 | Reln | 4.8716546 | 0.02757785 | 0.28054778 |
| ENSMUSG00000026820 | Ptges2 | 1.2505453 | 0.02762395 | 0.28090427 |
| ENSMUSG00000037762 | Slc16a9 | -3.4327146 | 0.02764329 | 0.28098853 |
| ENSMUSG00000021096 | Ppm1a | -1.169481 | 0.02768891 | 0.28133967 |
| ENSMUSG00000026170 | Cyp27a1 | -1.5082715 | 0.02771658 | 0.28150827 |
| ENSMUSG00000023994 | Nfya | 1.30232456 | 0.02773902 | 0.28151686 |
| ENSMUSG00000002147 | Stat6 | -1.1544172 | 0.02773958 | 0.28151686 |
| ENSMUSG00000106044 | Gm42860 | 2.23180411 | 0.0277773 | 0.28172019 |
| ENSMUSG00000004730 | Adgre1 | -1.5247293 | 0.02778369 | 0.28172019 |
| ENSMUSG00000085795 | Zfp703 | -1.5017144 | 0.02779287 | 0.28172019 |
| ENSMUSG00000006456 | Rbm14 | 1.33289884 | 0.02782029 | 0.2818856 |
| ENSMUSG00000052798 | Nup107 | 1.34117075 | 0.02784185 | 0.28198686 |
| ENSMUSG00000021993 | Mipep | 1.25576566 | 0.02785247 | 0.28198686 |
| ENSMUSG00000061451 | Tmem151a | -2.6797671 | 0.02802258 | 0.28359612 |
| ENSMUSG00000028587 | Orc1 | 1.66066724 | 0.02806412 | 0.28380228 |
| ENSMUSG00000070501 | Ifi214 | -2.8680091 | 0.02806529 | 0.28380228 |
| ENSMUSG00000086043 | Gm12473 | -5.0487715 | 0.0281316 | 0.28429925 |
| ENSMUSG00000040167 | Ikzf5 | 1.26403289 | 0.02813681 | 0.28429925 |
| ENSMUSG00000022797 | Tfrc | 1.67773639 | 0.02820129 | 0.28477015 |
| ENSMUSG00000102882 | Gm2065 | 4.65709601 | 0.02820791 | 0.28477015 |
| ENSMUSG00000116298 | 4833412C15I | 22.5286631 | 0.02821703 | 0.28477015 |

| | | | | |
|---|---|---|---|---|
| ENSMUSG00000078864 | Gm14322 | 1.87853785 | 0.02828964 | 0.28516074 |
| ENSMUSG00000043342 | Hoxd9 | -2.1755925 | 0.02829857 | 0.28516074 |
| ENSMUSG00000031149 | Praf2 | -1.35924 | 0.02830131 | 0.28516074 |
| ENSMUSG00000042228 | Lyn | -1.2801839 | 0.02831485 | 0.28516074 |
| ENSMUSG00000036218 | Pdzrn4 | -14.923782 | 0.02832061 | 0.28516074 |
| ENSMUSG00000026792 | Lrsam1 | 1.27456943 | 0.02832867 | 0.28516074 |
| ENSMUSG00000000420 | Galnt1 | -1.2223468 | 0.02835447 | 0.28516074 |
| ENSMUSG00000034525 | Ice1 | 1.28959914 | 0.02836287 | 0.28516074 |
| ENSMUSG00000021282 | Eif5 | -1.1581509 | 0.02837142 | 0.28516074 |
| ENSMUSG00000056078 | Lipm | 13.8572591 | 0.02837256 | 0.28516074 |
| ENSMUSG00000029070 | Mxra8 | -1.6721127 | 0.02837961 | 0.28516074 |
| ENSMUSG00000062526 | Mppe1 | -1.3527492 | 0.02839039 | 0.28516074 |
| ENSMUSG00000025607 | Copg2 | 1.16190663 | 0.02841953 | 0.28534061 |
| ENSMUSG00000050222 | Il17d | 2.43281445 | 0.02844089 | 0.28544226 |
| ENSMUSG00000073400 | Trim10 | 4.95196014 | 0.02846267 | 0.28551856 |
| ENSMUSG00000039004 | Bmp6 | -2.3777406 | 0.02847096 | 0.28551856 |
| ENSMUSG00000032046 | Abhd12 | -1.2808444 | 0.02850897 | 0.28576028 |
| ENSMUSG00000030772 | Dkk3 | -1.9197923 | 0.02851755 | 0.28576028 |
| ENSMUSG00000015467 | Egfl8 | -3.3981159 | 0.02853011 | 0.28577345 |
| ENSMUSG00000038685 | Rtel1 | 1.32435183 | 0.02856636 | 0.28601133 |
| ENSMUSG00000029797 | Sspo | 6.60266087 | 0.02858353 | 0.28601133 |
| ENSMUSG00000021957 | Tkt | 1.29258832 | 0.02858763 | 0.28601133 |
| ENSMUSG00000039395 | Mreg | -1.862193 | 0.02860244 | 0.28604695 |
| ENSMUSG00000037070 | Rbmxl1 | 1.24272167 | 0.02862851 | 0.28619497 |
| ENSMUSG00000074818 | Pdzd7 | 2.2609784 | 0.0287048 | 0.28684484 |
| ENSMUSG00000080989 | Gm14048 | 85.9647648 | 0.02895879 | 0.28896395 |
| ENSMUSG00000054675 | Tmem119 | -1.7080337 | 0.0289611 | 0.28896395 |
| ENSMUSG00000021262 | Evl | -1.4599642 | 0.02896682 | 0.28896395 |
| ENSMUSG00000030753 | Thap12 | 1.20798316 | 0.028983 | 0.28896395 |
| ENSMUSG00000019768 | Esr1 | -2.0219907 | 0.02898343 | 0.28896395 |
| ENSMUSG00000020032 | Nuak1 | -1.4468498 | 0.02899213 | 0.28896395 |
| ENSMUSG00000002105 | Slc39a13 | -1.2955962 | 0.02900518 | 0.28896395 |
| ENSMUSG00000021772 | Nkiras1 | -1.5641795 | 0.02900783 | 0.28896395 |
| ENSMUSG00000009376 | Met | 1.51433871 | 0.02902283 | 0.28900006 |
| ENSMUSG00000025915 | Sgk3 | -1.5286019 | 0.02906445 | 0.28930114 |
| ENSMUSG00000036452 | Arhgap26 | 1.48467142 | 0.0290945 | 0.28948463 |
| ENSMUSG00000045659 | Plekha7 | 1.42327125 | 0.02910567 | 0.28948463 |
| ENSMUSG00000018999 | Slc35b4 | -1.1754362 | 0.02912189 | 0.28953267 |
| ENSMUSG00000038848 | Ythdf1 | 1.14503771 | 0.02915555 | 0.28968345 |
| ENSMUSG00000039741 | Bahcc1 | 1.60270883 | 0.02915986 | 0.28968345 |
| ENSMUSG00000033904 | Ccp110 | 1.42920251 | 0.02920108 | 0.28994825 |
| ENSMUSG00000029334 | Prkg2 | -3.5839726 | 0.02920933 | 0.28994825 |
| ENSMUSG00000057716 | Tmem178b | -27.643046 | 0.02926697 | 0.29015204 |
| ENSMUSG00000052928 | Ctif | -1.4237803 | 0.02926969 | 0.29015204 |
| ENSMUSG00000051306 | Usp42 | 1.21548355 | 0.02927237 | 0.29015204 |

| | | | | |
|---|---|---|---|---|
| ENSMUSG00000025050 | Pcgf6 | 1.33202374 | 0.02927834 | 0.29015204 |
| ENSMUSG00000014547 | Wdfy2 | 1.32659328 | 0.02928695 | 0.29015204 |
| ENSMUSG00000019854 | Reps1 | 1.14475988 | 0.02931653 | 0.2902561 |
| ENSMUSG00000028362 | Tnfsf8 | -1.9438147 | 0.02933003 | 0.2902561 |
| ENSMUSG00000060509 | Xcr1 | -2.3550425 | 0.02933172 | 0.2902561 |
| ENSMUSG00000051879 | Krt71 | 4.87807983 | 0.0293664 | 0.29042 |
| ENSMUSG00000048174 | Tmem81 | -1.955385 | 0.02937114 | 0.29042 |
| ENSMUSG00000029368 | Alb | -31.93899 | 0.02940666 | 0.29065812 |
| ENSMUSG00000042308 | Setd1a | 1.19178415 | 0.02945675 | 0.29091208 |
| ENSMUSG00000063286 | Gm8995 | -1.6329168 | 0.02946068 | 0.29091208 |
| ENSMUSG00000035762 | Tmem161b | 1.27500177 | 0.0294667 | 0.29091208 |
| ENSMUSG00000022678 | Nde1 | 1.23393808 | 0.02947902 | 0.29092073 |
| ENSMUSG00000036299 | BC031181 | -1.2766763 | 0.02951344 | 0.29093675 |
| ENSMUSG00000040891 | Foxa3 | 9.12168274 | 0.02953058 | 0.29093675 |
| ENSMUSG00000020257 | Wdr82 | 1.20916579 | 0.02954243 | 0.29093675 |
| ENSMUSG00000051726 | Kcnf1 | -8.5340544 | 0.02954629 | 0.29093675 |
| ENSMUSG00000032411 | Tfdp2 | 1.31129442 | 0.02954884 | 0.29093675 |
| ENSMUSG00000097715 | Gpr137b-ps | -1.7165413 | 0.02954934 | 0.29093675 |
| ENSMUSG00000040569 | Slc26a7 | 4.33450951 | 0.02956756 | 0.29100339 |
| ENSMUSG00000021906 | Oxnad1 | 1.2543588 | 0.02959228 | 0.291134 |
| ENSMUSG00000024974 | Smc3 | 1.31791687 | 0.02961914 | 0.29128545 |
| ENSMUSG00000031996 | Aplp2 | 1.2299067 | 0.02966108 | 0.29158503 |
| ENSMUSG00000040591 | 1110051M20 | -1.2990418 | 0.02970636 | 0.29190525 |
| ENSMUSG00000044583 | Tlr7 | -1.5658431 | 0.02973368 | 0.29190525 |
| ENSMUSG00000045826 | Ptprcap | -2.1091686 | 0.02975461 | 0.29190525 |
| ENSMUSG00000045092 | S1pr1 | -1.7327488 | 0.02975554 | 0.29190525 |
| ENSMUSG00000003644 | Rps6ka1 | 1.39527524 | 0.0297604 | 0.29190525 |
| ENSMUSG00000023000 | Dhh | -2.792982 | 0.02976995 | 0.29190525 |
| ENSMUSG00000042215 | Bag2 | 1.47496358 | 0.02977406 | 0.29190525 |
| ENSMUSG00000024525 | Impa2 | 1.32183709 | 0.02979926 | 0.29193956 |
| ENSMUSG00000038173 | Enpp6 | -3.5250116 | 0.02980054 | 0.29193956 |
| ENSMUSG00000110424 | 1700012D14 | 2.20789306 | 0.02983039 | 0.29211936 |
| ENSMUSG00000118618 | AC123856.1 | 5.81840081 | 0.02986313 | 0.29231926 |
| ENSMUSG00000027238 | Frmd5 | -3.4544058 | 0.0298738 | 0.29231926 |
| ENSMUSG00000021904 | Sema3g | -1.9846614 | 0.02990296 | 0.29249191 |
| ENSMUSG00000025375 | Aatk | -2.180275 | 0.02995821 | 0.29291957 |
| ENSMUSG00000063320 | 1190007I07R | 1.53470636 | 0.02998253 | 0.29294667 |
| ENSMUSG00000061535 | C1qtnf7 | -4.0087779 | 0.02998935 | 0.29294667 |
| ENSMUSG00000020042 | Btbd11 | 1.58778856 | 0.02999556 | 0.29294667 |
| ENSMUSG00000045333 | Zfp423 | -2.3716014 | 0.03004208 | 0.2932191 |
| ENSMUSG00000021994 | Wnt5a | -1.6218013 | 0.0300542 | 0.2932191 |
| ENSMUSG00000064147 | Rab44 | -2.4511232 | 0.03005807 | 0.2932191 |
| ENSMUSG00000031304 | Il2rg | -1.342276 | 0.03017778 | 0.29398099 |
| ENSMUSG00000035206 | Sppl2b | 1.22450002 | 0.030178 | 0.29398099 |
| ENSMUSG00000027067 | Ssrp1 | 1.1851453 | 0.03017895 | 0.29398099 |

| | | | | |
|---|---|---|---|---|
| ENSMUSG00000033228 | Scaf11 | 1.20466352 | 0.03018245 | 0.29398099 |
| ENSMUSG00000073000 | Gm10451 | 2.20055971 | 0.03020261 | 0.29404452 |
| ENSMUSG00000051396 | Gm45902 | 1.56966236 | 0.03023581 | 0.29404452 |
| ENSMUSG00000027230 | Creb3l1 | -2.0393397 | 0.03024042 | 0.29404452 |
| ENSMUSG00000061911 | Myt1l | 2.37251434 | 0.03024732 | 0.29404452 |
| ENSMUSG00000013787 | Ehmt2 | 1.16049796 | 0.03025288 | 0.29404452 |
| ENSMUSG00000057766 | Ankrd29 | -2.2223853 | 0.0302584 | 0.29404452 |
| ENSMUSG00000049985 | Ankrd55 | -5.2895767 | 0.03030553 | 0.29438754 |
| ENSMUSG00000064043 | Trerf1 | -1.5757631 | 0.03031687 | 0.29438754 |
| ENSMUSG00000001270 | Ckb | -2.078613 | 0.03034447 | 0.29440438 |
| ENSMUSG00000027708 | Dcun1d1 | 1.22790297 | 0.03035239 | 0.29440438 |
| ENSMUSG00000095990 | Zfp97 | 1.27312442 | 0.03035932 | 0.29440438 |
| ENSMUSG00000040945 | Rcc2 | 1.32678861 | 0.03036494 | 0.29440438 |
| ENSMUSG00000030725 | Lipt2 | 1.62396715 | 0.0304372 | 0.29499245 |
| ENSMUSG00000005057 | Sh2b2 | -1.5317239 | 0.03046102 | 0.29511077 |
| ENSMUSG00000026175 | Vil1 | 5.04682619 | 0.03047868 | 0.2951693 |
| ENSMUSG00000055240 | Zfp101 | 1.55035199 | 0.03049574 | 0.295188 |
| ENSMUSG00000066721 | Zfp575 | 2.00537057 | 0.03050384 | 0.295188 |
| ENSMUSG00000001751 | Naglu | -1.4307706 | 0.03052266 | 0.29525768 |
| ENSMUSG00000051177 | Plcb1 | -2.0561393 | 0.03055248 | 0.29543363 |
| ENSMUSG00000018217 | Pmp22 | -1.5767388 | 0.03059217 | 0.29570496 |
| ENSMUSG00000027931 | Npr1 | 2.15840564 | 0.03062065 | 0.29586769 |
| ENSMUSG00000044737 | Klk14 | -23.987915 | 0.03064233 | 0.29596459 |
| ENSMUSG00000056130 | Ticam2 | -1.6621779 | 0.03066618 | 0.29608245 |
| ENSMUSG00000027981 | Rnpc3 | 1.41652462 | 0.03070039 | 0.2962341 |
| ENSMUSG00000032382 | Snx1 | -1.3299717 | 0.0307052 | 0.2962341 |
| ENSMUSG00000026011 | Ctla4 | -3.1093779 | 0.03071812 | 0.29624625 |
| ENSMUSG00000007610 | Gtpbp3 | 1.27375792 | 0.03073965 | 0.29634138 |
| ENSMUSG00000035834 | Polr3g | -1.4163179 | 0.03075857 | 0.29641137 |
| ENSMUSG00000031749 | St3gal2 | -1.4313509 | 0.03082551 | 0.29694381 |
| ENSMUSG00000036867 | Smad6 | -2.2088997 | 0.03089128 | 0.29745101 |
| ENSMUSG00000044827 | Tlr1 | -1.6487122 | 0.03090157 | 0.29745101 |
| ENSMUSG00000052749 | Trim30b | -1.8523412 | 0.03092117 | 0.29752699 |
| ENSMUSG00000028161 | Ppp3ca | -1.2676772 | 0.03096745 | 0.29785956 |
| ENSMUSG00000040325 | Dcaf1 | 1.25183425 | 0.03099018 | 0.29796535 |
| ENSMUSG00000112833 | Gm36595 | -20.739586 | 0.03103631 | 0.29829606 |
| ENSMUSG00000034007 | Scaper | 1.22666325 | 0.03107383 | 0.29854369 |
| ENSMUSG00000026322 | Htr4 | -3.6797675 | 0.03109468 | 0.29857419 |
| ENSMUSG00000048429 | Timm29 | 1.19778116 | 0.0311005 | 0.29857419 |
| ENSMUSG00000028015 | Ctso | -1.4132148 | 0.03118255 | 0.29906501 |
| ENSMUSG00000021943 | Gdf10 | -4.7622514 | 0.0311846 | 0.29906501 |
| ENSMUSG00000030474 | Siglece | -1.7871431 | 0.03118693 | 0.29906501 |
| ENSMUSG00000031568 | Rwdd4a | 1.18691683 | 0.03121767 | 0.29917307 |
| ENSMUSG00000097129 | 4930507D05 | 7.65163851 | 0.03122174 | 0.29917307 |
| ENSMUSG00000039601 | Rcan2 | -3.2308754 | 0.03129307 | 0.29964302 |

| | | | | |
|---|---|---|---|---|
| ENSMUSG00000032452 | Clstn2 | -4.4800492 | 0.03129437 | 0.29964302 |
| ENSMUSG00000071256 | Zfp213 | 1.2693285 | 0.03131594 | 0.29973663 |
| ENSMUSG00000040040 | Ift88 | 1.27966125 | 0.03135704 | 0.30001702 |
| ENSMUSG00000031971 | Ccsap | 1.4152185 | 0.03140624 | 0.30037462 |
| ENSMUSG00000060860 | Ube2s | 1.37345619 | 0.03143611 | 0.30054718 |
| ENSMUSG00000091803 | Cox16 | -1.3441126 | 0.03145487 | 0.30061347 |
| ENSMUSG00000042766 | Trim46 | 1.71645253 | 0.03147295 | 0.30062332 |
| ENSMUSG00000032397 | Tipin | 1.38627233 | 0.03147956 | 0.30062332 |
| ENSMUSG00000053604 | Rpia | 1.41553941 | 0.03150029 | 0.30067618 |
| ENSMUSG00000021870 | Slmap | 1.33842863 | 0.03150876 | 0.30067618 |
| ENSMUSG00000078129 | Actl10 | 13.9729749 | 0.0315751 | 0.30107787 |
| ENSMUSG00000025743 | Sdc3 | -1.5709557 | 0.03158234 | 0.30107787 |
| ENSMUSG00000024018 | Ccdc167 | 1.41594313 | 0.0315864 | 0.30107787 |
| ENSMUSG00000007805 | Twist2 | -2.6794985 | 0.03171097 | 0.30215191 |
| ENSMUSG00000026227 | 2810459M11 | 5.32998923 | 0.0317248 | 0.30217038 |
| ENSMUSG00000044894 | Uqcrq | -1.3382296 | 0.03177535 | 0.30243656 |
| ENSMUSG00000041995 | Zbed3 | 1.37431779 | 0.03177655 | 0.30243656 |
| ENSMUSG00000063903 | Klk1 | 8.08887432 | 0.0317888 | 0.30243988 |
| ENSMUSG00000001999 | Blvra | -1.3179277 | 0.03183885 | 0.30280269 |
| ENSMUSG00000000276 | Dgke | -1.310525 | 0.03186795 | 0.30296606 |
| ENSMUSG00000032033 | Barx2 | 2.72724326 | 0.03189917 | 0.30311868 |
| ENSMUSG00000022208 | Jph4 | -2.1427882 | 0.0319086 | 0.30311868 |
| ENSMUSG00000044646 | Zbtb7c | -1.3773927 | 0.03192406 | 0.30311868 |
| ENSMUSG00000031728 | Zfp821 | 1.2732522 | 0.03193171 | 0.30311868 |
| ENSMUSG00000019461 | Plscr3 | -1.1633763 | 0.03196562 | 0.3033272 |
| ENSMUSG00000029500 | Pgam5 | 1.2166192 | 0.03199687 | 0.30351044 |
| ENSMUSG00000026516 | Nvl | 1.28960016 | 0.03211776 | 0.30450429 |
| ENSMUSG00000020409 | Slu7 | -1.178345 | 0.03212561 | 0.30450429 |
| ENSMUSG00000034681 | Rnps1 | 1.22666303 | 0.03216988 | 0.30475761 |
| ENSMUSG00000020963 | Tshr | -4.1400509 | 0.03217632 | 0.30475761 |
| ENSMUSG00000030465 | Psd3 | -1.8938783 | 0.03222206 | 0.30507713 |
| ENSMUSG00000093861 | Igkv1-110 | -10.690082 | 0.03225476 | 0.30527295 |
| ENSMUSG00000037894 | H2az1 | 1.36836241 | 0.03228331 | 0.30534637 |
| ENSMUSG00000099375 | Gm28187 | 2.49761599 | 0.03229296 | 0.30534637 |
| ENSMUSG00000021500 | Ddx46 | 1.23274805 | 0.03229856 | 0.30534637 |
| ENSMUSG00000032481 | Smarcc1 | 1.35987411 | 0.03231064 | 0.30534701 |
| ENSMUSG00000044674 | Fzd1 | -1.7813248 | 0.03234666 | 0.30557376 |
| ENSMUSG00000045751 | Mms22l | 1.3436319 | 0.03237767 | 0.30575299 |
| ENSMUSG00000079190 | AC133103.1 | 6.13646755 | 0.03240068 | 0.30585665 |
| ENSMUSG00000028134 | Ptbp2 | 1.44774813 | 0.03246669 | 0.30636592 |
| ENSMUSG00000045231 | BC106179 | -2.9357396 | 0.03254367 | 0.30684966 |
| ENSMUSG00000030660 | Pik3c2a | 1.29720961 | 0.03255108 | 0.30684966 |
| ENSMUSG00000016018 | Mtrex | 1.20356238 | 0.03255417 | 0.30684966 |
| ENSMUSG00000097848 | Gm807 | -3.8689033 | 0.03260207 | 0.30709566 |
| ENSMUSG00000058655 | Eif4b | 1.21344859 | 0.03260444 | 0.30709566 |

| | | | | |
|---|---|---|---|---|
| ENSMUSG00000033327 | Tnxb | -2.2457753 | 0.0326458 | 0.3073713 |
| ENSMUSG00000061769 | Klra6 | 11.5466698 | 0.0326622 | 0.30741182 |
| ENSMUSG00000085058 | 8030453O22 | 3.56074741 | 0.0326994 | 0.30764795 |
| ENSMUSG00000016028 | Celsr1 | 1.73982626 | 0.03276375 | 0.30808349 |
| ENSMUSG00000025722 | Wdr73 | 1.24696183 | 0.03276994 | 0.30808349 |
| ENSMUSG00000006958 | Chrd | -2.0515427 | 0.03279912 | 0.30824384 |
| ENSMUSG00000073627 | C130036L24I | 1.88705903 | 0.03287218 | 0.30881622 |
| ENSMUSG00000041653 | Pnpla3 | -7.7680929 | 0.03290226 | 0.3089846 |
| ENSMUSG00000023094 | Msrb2 | -2.0281983 | 0.03294735 | 0.30929367 |
| ENSMUSG00000027339 | Rassf2 | -1.5737967 | 0.03299385 | 0.30961586 |
| ENSMUSG00000093989 | Rnasek | -1.4455982 | 0.03301063 | 0.30965896 |
| ENSMUSG00000029875 | Ccdc184 | 3.71282066 | 0.03302479 | 0.30967745 |
| ENSMUSG00000085492 | Trmt61b | 1.46887201 | 0.0330674 | 0.30996268 |
| ENSMUSG00000024163 | Mapk8ip3 | 1.46847164 | 0.03310608 | 0.31021077 |
| ENSMUSG00000098022 | Zfp82 | 1.48677705 | 0.03313652 | 0.31036313 |
| ENSMUSG00000042662 | Dusp15 | -7.1472695 | 0.03314676 | 0.31036313 |
| ENSMUSG00000033998 | Kcnk1 | 1.60591479 | 0.03323464 | 0.31100806 |
| ENSMUSG00000036572 | Upf3b | 1.24650928 | 0.03324012 | 0.31100806 |
| ENSMUSG00000046675 | Tmem251 | -1.2993003 | 0.03328887 | 0.31134958 |
| ENSMUSG00000007589 | Tinf2 | 1.2356044 | 0.03331015 | 0.31143395 |
| ENSMUSG00000054619 | Mettl7a1 | -1.7798802 | 0.03343269 | 0.31246473 |
| ENSMUSG00000071054 | Safb | 1.22765401 | 0.03346986 | 0.31258222 |
| ENSMUSG00000087075 | Lbhd2 | 4.34243576 | 0.03346987 | 0.31258222 |
| ENSMUSG00000068264 | Ap5s1 | -1.4295004 | 0.03351577 | 0.31280811 |
| ENSMUSG00000040372 | Gpr63 | -2.7469599 | 0.0335392 | 0.31280811 |
| ENSMUSG00000031669 | Gins3 | 1.28194677 | 0.03354136 | 0.31280811 |
| ENSMUSG00000024206 | Rfx2 | 1.43272526 | 0.03354329 | 0.31280811 |
| ENSMUSG00000096054 | Syne1 | -1.5544699 | 0.03376066 | 0.31457683 |
| ENSMUSG00000053581 | Zfand2a | -1.4072036 | 0.03376759 | 0.31457683 |
| ENSMUSG00000031543 | Ank1 | -2.0979979 | 0.03377205 | 0.31457683 |
| ENSMUSG00000051341 | Zfp52 | -1.3241 | 0.03378247 | 0.31457683 |
| ENSMUSG00000036275 | 9530068E07I | -1.2204079 | 0.03380592 | 0.31467988 |
| ENSMUSG00000032496 | Ltf | 4.85577243 | 0.03383379 | 0.31473337 |
| ENSMUSG00000080237 | Gm14239 | 29.2104143 | 0.03383644 | 0.31473337 |
| ENSMUSG00000068923 | Syt11 | -1.6058459 | 0.03385944 | 0.31483211 |
| ENSMUSG00000043411 | Usp48 | 1.32706887 | 0.03389255 | 0.31502467 |
| ENSMUSG00000047141 | Zfp654 | 1.27092543 | 0.03391874 | 0.31515287 |
| ENSMUSG00000020903 | Stx8 | -1.314174 | 0.03398271 | 0.3156318 |
| ENSMUSG00000024943 | Smc5 | 1.33391283 | 0.03401682 | 0.3158332 |
| ENSMUSG00000021048 | Mthfd1 | 1.18756112 | 0.03403368 | 0.31587435 |
| ENSMUSG00000040797 | Iqsec3 | -48.884706 | 0.03405172 | 0.31592633 |
| ENSMUSG00000022365 | Derl1 | -1.2118242 | 0.03408452 | 0.31611523 |
| ENSMUSG00000044033 | Ccdc141 | 2.4371041 | 0.03412436 | 0.3162899 |
| ENSMUSG00000018363 | Smurf2 | -1.2367422 | 0.03412824 | 0.3162899 |
| ENSMUSG00000044375 | Pcare | 31.025413 | 0.03421056 | 0.31686968 |

| | | | | |
|---|---|---|---|---|
| ENSMUSG00000001901 | Kcnh6 | 54.267257 | 0.03423052 | 0.31686968 |
| ENSMUSG00000114871 | Gm21370 | -9.1097793 | 0.03423879 | 0.31686968 |
| ENSMUSG00000030165 | Klrd1 | -1.6351197 | 0.03426313 | 0.31686968 |
| ENSMUSG00000036932 | Aifm1 | 1.19145251 | 0.03427843 | 0.31686968 |
| ENSMUSG00000024695 | Zfp91 | 1.1838455 | 0.03429159 | 0.31686968 |
| ENSMUSG00000024817 | Uhrf2 | 1.3519529 | 0.03430297 | 0.31686968 |
| ENSMUSG00000027882 | Stxbp3 | 1.23739161 | 0.03432432 | 0.31686968 |
| ENSMUSG00000091474 | 2610021A01 | 1.31598567 | 0.03432986 | 0.31686968 |
| ENSMUSG00000053916 | Nanp | 1.33315847 | 0.03434059 | 0.31686968 |
| ENSMUSG00000026904 | Slc4a10 | 26.5396091 | 0.03434198 | 0.31686968 |
| ENSMUSG00000032024 | Clmp | -1.8605467 | 0.03434896 | 0.31686968 |
| ENSMUSG00000027184 | Caprin1 | 1.18940011 | 0.0343529 | 0.31686968 |
| ENSMUSG00000001910 | Nacc1 | 1.21579177 | 0.03439313 | 0.31712563 |
| ENSMUSG00000021929 | Kpna3 | 1.23182247 | 0.03442051 | 0.31721098 |
| ENSMUSG00000002668 | Dennd1c | -1.4602018 | 0.03445137 | 0.31721098 |
| ENSMUSG00000041343 | Ankrd42 | 2.05541479 | 0.03446383 | 0.31721098 |
| ENSMUSG00000053338 | Tarm1 | -3.8998378 | 0.03447035 | 0.31721098 |
| ENSMUSG00000062232 | Rapgef2 | -1.2535324 | 0.03447247 | 0.31721098 |
| ENSMUSG00000037851 | Iars | 1.29637927 | 0.03447728 | 0.31721098 |
| ENSMUSG00000032726 | Bmp8a | -18.188088 | 0.03452131 | 0.31739836 |
| ENSMUSG00000039476 | Prrx2 | -2.9632397 | 0.03452467 | 0.31739836 |
| ENSMUSG00000034947 | Tmem106a | -1.4052766 | 0.03453512 | 0.31739836 |
| ENSMUSG00000020074 | Ccar1 | 1.22847725 | 0.03460822 | 0.31795518 |
| ENSMUSG00000085101 | Platr16 | 14.5724552 | 0.03462706 | 0.31801331 |
| ENSMUSG00000018995 | Nars2 | 1.32107756 | 0.03466103 | 0.31821028 |
| ENSMUSG00000040612 | Ildr2 | -2.6081992 | 0.03469296 | 0.31833317 |
| ENSMUSG00000028962 | Slc4a2 | 1.13548811 | 0.03469947 | 0.31833317 |
| ENSMUSG00000078867 | Gm14418 | 2.01891019 | 0.0347646 | 0.31865646 |
| ENSMUSG00000019320 | Noxo1 | 1.90179234 | 0.03477341 | 0.31865646 |
| ENSMUSG00000039242 | B3galnt2 | 1.33106956 | 0.03478281 | 0.31865646 |
| ENSMUSG00000004043 | Stat5a | -1.4540127 | 0.03478487 | 0.31865646 |
| ENSMUSG00000033960 | Jcad | -1.5645228 | 0.03481711 | 0.31883691 |
| ENSMUSG00000034917 | Tjp3 | 1.8605628 | 0.03485802 | 0.31909652 |
| ENSMUSG00000029401 | Rilpl2 | -1.3018828 | 0.03488771 | 0.31920301 |
| ENSMUSG00000032038 | St3gal4 | -1.3423962 | 0.03489477 | 0.31920301 |
| ENSMUSG00000020630 | Rnaseh1 | -1.2507436 | 0.03493269 | 0.31943491 |
| ENSMUSG00000021548 | Ccnh | 1.18360556 | 0.03495482 | 0.31952226 |
| ENSMUSG00000026134 | Prim2 | 1.26048334 | 0.03500005 | 0.31975806 |
| ENSMUSG00000048445 | Ccdc57 | 1.53253767 | 0.03500578 | 0.31975806 |
| ENSMUSG00000029108 | Pcdh7 | -2.3322848 | 0.03510746 | 0.3205661 |
| ENSMUSG00000029032 | Arhgef16 | 1.62135184 | 0.03512813 | 0.3205661 |
| ENSMUSG00000029166 | Mapre3 | -1.4486606 | 0.03513209 | 0.3205661 |
| ENSMUSG00000006310 | Zbtb32 | 2.56085505 | 0.03515527 | 0.32066251 |
| ENSMUSG00000037849 | Ifi206 | -2.1545318 | 0.03517829 | 0.32075731 |
| ENSMUSG00000031613 | Hpgd | -2.7557277 | 0.03523875 | 0.32119341 |

| | | | | |
|---|---|---|---|---|
| ENSMUSG00000032174 | Icam5 | 2.2716545 | 0.03526199 | 0.32128996 |
| ENSMUSG00000005078 | Jkamp | -1.2651064 | 0.03531178 | 0.32145087 |
| ENSMUSG00000006301 | Tmbim1 | -1.3138158 | 0.03531704 | 0.32145087 |
| ENSMUSG00000097519 | 4930558J18F | 2.13501726 | 0.0353176 | 0.32145087 |
| ENSMUSG00000028622 | Mrpl37 | 1.20850056 | 0.03536114 | 0.32173196 |
| ENSMUSG00000029247 | Paics | 1.23573076 | 0.03538222 | 0.32180849 |
| ENSMUSG00000069237 | Fam8a1 | 1.18783682 | 0.03548641 | 0.32255956 |
| ENSMUSG00000028417 | Tal2 | -6.2900602 | 0.03549018 | 0.32255956 |
| ENSMUSG00000047747 | Rnf150 | -1.6935269 | 0.03552489 | 0.3227596 |
| ENSMUSG00000024393 | Prrc2a | 1.17596154 | 0.03554764 | 0.32285081 |
| ENSMUSG00000000204 | Slfn4 | 3.82705543 | 0.03557938 | 0.32302368 |
| ENSMUSG00000024511 | Rab27b | -1.6338217 | 0.03559224 | 0.32302503 |
| ENSMUSG00000021374 | Nup153 | 1.24567339 | 0.03564603 | 0.32339765 |
| ENSMUSG00000083307 | AA414768 | -2.102013 | 0.03566822 | 0.32348348 |
| ENSMUSG00000037060 | Cavin3 | -1.6265507 | 0.03574141 | 0.32403163 |
| ENSMUSG00000071478 | H2ac7 | -4.4921729 | 0.0357822 | 0.32428573 |
| ENSMUSG00000042476 | Abcb4 | -2.7110433 | 0.03585016 | 0.32468243 |
| ENSMUSG00000092021 | Gbp11 | -9.8961573 | 0.03586182 | 0.32468243 |
| ENSMUSG00000041912 | Tdrkh | 2.33713748 | 0.0358643 | 0.32468243 |
| ENSMUSG00000042817 | Flt3 | -2.4445056 | 0.0359259 | 0.32503679 |
| ENSMUSG00000020399 | Havcr2 | -1.478561 | 0.03593501 | 0.32503679 |
| ENSMUSG00000002731 | Prkra | 1.21714133 | 0.03595337 | 0.32503679 |
| ENSMUSG00000035004 | Igsf6 | -1.4977866 | 0.03595461 | 0.32503679 |
| ENSMUSG00000067586 | S1pr3 | -2.2188973 | 0.03599501 | 0.32528634 |
| ENSMUSG00000106961 | Gm43128 | 4.94934833 | 0.03602801 | 0.32546885 |
| ENSMUSG00000060216 | Arrb2 | -1.431442 | 0.0361014 | 0.32601589 |
| ENSMUSG00000062310 | Glrp1 | 3.05602774 | 0.03611923 | 0.32606105 |
| ENSMUSG00000031351 | Zfp185 | 1.6751482 | 0.03618836 | 0.32644981 |
| ENSMUSG00000086124 | A530076I17F | -37.452395 | 0.03618857 | 0.32644981 |
| ENSMUSG00000029469 | Ift81 | 1.27544899 | 0.03621415 | 0.32644981 |
| ENSMUSG00000022883 | Robo1 | -2.223699 | 0.03622082 | 0.32644981 |
| ENSMUSG00000028476 | Reck | -1.6337009 | 0.03623138 | 0.32644981 |
| ENSMUSG00000018339 | Gpx3 | -3.7284149 | 0.03624912 | 0.32644981 |
| ENSMUSG00000052997 | Uba2 | 1.16923766 | 0.03625222 | 0.32644981 |
| ENSMUSG00000044005 | Gls2 | 2.11854142 | 0.03627743 | 0.32646341 |
| ENSMUSG00000038260 | Trpm4 | 1.41720981 | 0.03627942 | 0.32646341 |
| ENSMUSG00000021671 | Poc5 | 1.21982886 | 0.03629944 | 0.32652788 |
| ENSMUSG00000032185 | Carm1 | 1.25038013 | 0.03635192 | 0.32671875 |
| ENSMUSG00000015016 | Acsf3 | 1.29737064 | 0.03635823 | 0.32671875 |
| ENSMUSG00000003228 | Grk5 | 1.46036972 | 0.03635923 | 0.32671875 |
| ENSMUSG00000015468 | Notch4 | -1.5842024 | 0.03647897 | 0.32767886 |
| ENSMUSG00000039671 | Zmynd8 | 1.36332056 | 0.03652247 | 0.32788098 |
| ENSMUSG00000006599 | Gtf2h1 | 1.19638766 | 0.03653645 | 0.32788098 |
| ENSMUSG00000033684 | Qsox1 | -1.4337943 | 0.03654018 | 0.32788098 |
| ENSMUSG00000057193 | Slc44a2 | -1.1816752 | 0.03660964 | 0.32823165 |

| | | | | |
|---|---|---|---|---|
| ENSMUSG00000066755 | Tnfsf18 | -3.4823719 | 0.03661087 | 0.32823165 |
| ENSMUSG00000034854 | Mfsd12 | -1.4843972 | 0.036618 | 0.32823165 |
| ENSMUSG00000000409 | Lck | -1.6094413 | 0.03663725 | 0.32828836 |
| ENSMUSG00000106239 | Gm9260 | -10.796859 | 0.03668662 | 0.32851726 |
| ENSMUSG00000037905 | Bri3bp | 1.44294681 | 0.03668865 | 0.32851726 |
| ENSMUSG00000099398 | Ms4a14 | -2.2113374 | 0.03670756 | 0.3285708 |
| ENSMUSG00000031508 | Ankrd10 | 1.28113622 | 0.0367459 | 0.32879819 |
| ENSMUSG00000034974 | Dapk3 | -1.5760014 | 0.03679252 | 0.32909948 |
| ENSMUSG00000029675 | Eln | -2.3993816 | 0.03683442 | 0.32929495 |
| ENSMUSG00000042133 | Ppig | 1.1730883 | 0.03684029 | 0.32929495 |
| ENSMUSG00000096199 | Ptrhd1 | -1.3607242 | 0.03686901 | 0.32941033 |
| ENSMUSG00000040669 | Phc1 | 1.20156314 | 0.03688883 | 0.32941033 |
| ENSMUSG00000041161 | Otud3 | 1.35155106 | 0.03689209 | 0.32941033 |
| ENSMUSG00000081219 | Bambi-ps1 | -3.9993198 | 0.03695548 | 0.32986048 |
| ENSMUSG00000117333 | Gm16386 | 2.34874089 | 0.036993 | 0.33007939 |
| ENSMUSG00000019173 | Rab5c | -1.2486039 | 0.03707578 | 0.33064365 |
| ENSMUSG00000068323 | Slc4a5 | 2.69088214 | 0.03708226 | 0.33064365 |
| ENSMUSG00000029265 | Dr1 | 1.17923507 | 0.0371014 | 0.33069829 |
| ENSMUSG00000043987 | Cep164 | 1.30055384 | 0.03713088 | 0.33084498 |
| ENSMUSG00000031730 | Dhodh | 1.23090159 | 0.03717303 | 0.33110447 |
| ENSMUSG00000030884 | Uqcrc2 | 1.12636642 | 0.03723807 | 0.33156756 |
| ENSMUSG00000022394 | L3mbtl2 | 1.20551328 | 0.03732545 | 0.3320004 |
| ENSMUSG00000044317 | Gpr4 | -1.7734765 | 0.03733361 | 0.3320004 |
| ENSMUSG00000097979 | Gm4691 | -7.9508954 | 0.0373344 | 0.3320004 |
| ENSMUSG00000025964 | Adam23 | -3.1580305 | 0.03734434 | 0.3320004 |
| ENSMUSG00000107362 | Gm40309 | 5.86608705 | 0.03735663 | 0.3320004 |
| ENSMUSG00000049672 | Zbtb14 | 1.27144517 | 0.03736507 | 0.3320004 |
| ENSMUSG00000031584 | Gsr | -1.401565 | 0.03740911 | 0.33209828 |
| ENSMUSG00000061887 | Ssbp3 | 1.20753275 | 0.03741059 | 0.33209828 |
| ENSMUSG00000044792 | Isca1 | 1.19264329 | 0.0374168 | 0.33209828 |
| ENSMUSG00000030747 | Dgat2 | -2.6461269 | 0.0374366 | 0.33209828 |
| ENSMUSG00000075588 | Hoxb2 | -1.7310453 | 0.03745138 | 0.33209828 |
| ENSMUSG00000024174 | Pot1b | 1.42263082 | 0.03746514 | 0.33209828 |
| ENSMUSG00000028820 | Sfpq | 1.21865511 | 0.03746959 | 0.33209828 |
| ENSMUSG00000049553 | Polr1a | 1.35008691 | 0.03749007 | 0.33209828 |
| ENSMUSG00000021133 | Susd6 | -1.3820233 | 0.03750194 | 0.33209828 |
| ENSMUSG00000040339 | Fam102b | -1.4754674 | 0.03750677 | 0.33209828 |
| ENSMUSG00000051329 | Nup160 | 1.28863746 | 0.03752672 | 0.33215915 |
| ENSMUSG00000037458 | Azin1 | -1.2625795 | 0.03763351 | 0.3327528 |
| ENSMUSG00000037337 | Map4k1 | -1.4639896 | 0.03764129 | 0.3327528 |
| ENSMUSG00000022673 | Mcm4 | 1.29494137 | 0.03765091 | 0.3327528 |
| ENSMUSG00000059027 | 9630013D21 | 2.67565464 | 0.03766323 | 0.3327528 |
| ENSMUSG00000040724 | Kcna2 | -5.6465874 | 0.03767635 | 0.3327528 |
| ENSMUSG00000046095 | Krt32 | -49.369396 | 0.03769479 | 0.3327528 |
| ENSMUSG00000033669 | Zfp7 | 1.40798055 | 0.03771008 | 0.3327528 |

| | | | | |
|---|---|---|---|---|
| ENSMUSG00000056536 | Pign | 1.29692736 | 0.03771157 | 0.3327528 |
| ENSMUSG00000022236 | Ropn1l | -2.1587206 | 0.03773417 | 0.3327528 |
| ENSMUSG00000019916 | P4ha1 | -1.7478361 | 0.03774313 | 0.3327528 |
| ENSMUSG00000025898 | Cwf19l2 | 1.19489582 | 0.03774902 | 0.3327528 |
| ENSMUSG00000097604 | Gm17322 | -7.706256 | 0.03775092 | 0.3327528 |
| ENSMUSG00000040025 | Ythdf2 | 1.18151216 | 0.0377652 | 0.33276324 |
| ENSMUSG00000016498 | Pdcd1lg2 | -3.2151381 | 0.03778125 | 0.33278933 |
| ENSMUSG00000112627 | 4933412E12I | -1.6953683 | 0.03787182 | 0.33347147 |
| ENSMUSG00000114993 | Gm6363 | 22.2327232 | 0.03793044 | 0.33383899 |
| ENSMUSG00000024424 | Ttc39c | -1.3930523 | 0.03794714 | 0.33383899 |
| ENSMUSG00000074715 | Ccl28 | 6.28813906 | 0.03795297 | 0.33383899 |
| ENSMUSG00000026094 | Stk17b | -1.3385743 | 0.03799146 | 0.33404658 |
| ENSMUSG00000097589 | Dleu2 | 1.57349889 | 0.03800286 | 0.33404658 |
| ENSMUSG00000040247 | Tbc1d10c | -2.1315998 | 0.03805239 | 0.33436633 |
| ENSMUSG00000034021 | Pds5b | 1.31082291 | 0.03812158 | 0.33470721 |
| ENSMUSG00000110616 | Gm36879 | 4.12295527 | 0.03812762 | 0.33470721 |
| ENSMUSG00000062382 | Ftl1-ps1 | -1.6279178 | 0.03813622 | 0.33470721 |
| ENSMUSG00000042532 | Golga7b | -17.477414 | 0.03814387 | 0.33470721 |
| ENSMUSG00000052139 | Babam2 | -1.2856554 | 0.03816173 | 0.33474829 |
| ENSMUSG00000025583 | Rptor | -1.1914033 | 0.0381767 | 0.33476405 |
| ENSMUSG00000026098 | Pms1 | 1.31184871 | 0.03819891 | 0.33479046 |
| ENSMUSG00000097736 | 9530059O14 | -9.8158185 | 0.03820606 | 0.33479046 |
| ENSMUSG00000017264 | Exosc10 | 1.18418434 | 0.03821964 | 0.33479403 |
| ENSMUSG00000026185 | Igfbp5 | -2.6075064 | 0.03826269 | 0.33505563 |
| ENSMUSG00000110266 | Gm32742 | 5.34884017 | 0.03831321 | 0.33538247 |
| ENSMUSG00000104693 | Gm42941 | 6.08868853 | 0.0383455 | 0.3355495 |
| ENSMUSG00000028385 | Snx30 | -1.4335604 | 0.03841413 | 0.33602492 |
| ENSMUSG00000019970 | Sgk1 | -1.8941376 | 0.03842627 | 0.33602492 |
| ENSMUSG00000030279 | C2cd5 | 1.26851221 | 0.03853052 | 0.33682061 |
| ENSMUSG00000026688 | Mgst3 | -1.6318747 | 0.03859376 | 0.33716197 |
| ENSMUSG00000074971 | Fibin | -2.6169191 | 0.03862634 | 0.33716197 |
| ENSMUSG00000031788 | Kifc3 | 1.25345105 | 0.03862977 | 0.33716197 |
| ENSMUSG00000074199 | Krtdap | -28.10987 | 0.0386308 | 0.33716197 |
| ENSMUSG00000025920 | Stau2 | 1.26620002 | 0.03863591 | 0.33716197 |
| ENSMUSG00000048856 | Slc25a47 | 1.7385237 | 0.03871854 | 0.33755897 |
| ENSMUSG00000033326 | Kdm4a | 1.20710228 | 0.03872867 | 0.33755897 |
| ENSMUSG00000030309 | Caprin2 | 1.5450633 | 0.03872911 | 0.33755897 |
| ENSMUSG00000024276 | Zfp397 | 1.39763733 | 0.03873453 | 0.33755897 |
| ENSMUSG00000050473 | Slc35d3 | -8.1482488 | 0.03876584 | 0.33761034 |
| ENSMUSG00000035898 | Uba6 | 1.27958264 | 0.03877385 | 0.33761034 |
| ENSMUSG00000048440 | Cyp4f16 | -1.2885245 | 0.03878125 | 0.33761034 |
| ENSMUSG00000029111 | Nelfa | 1.20881447 | 0.03879357 | 0.33761034 |
| ENSMUSG00000026544 | Dusp23 | -1.5430273 | 0.0388083 | 0.33762295 |
| ENSMUSG00000027175 | Tcp11l1 | -1.5822765 | 0.03884453 | 0.33771014 |
| ENSMUSG00000022718 | Dgcr8 | 1.36464777 | 0.03886474 | 0.33771014 |

| | | | | |
|---|---|---|---|---|
| ENSMUSG00000108655 | Gm44949 | 3.3047876 | 0.03887046 | 0.33771014 |
| ENSMUSG00000063077 | Kif1b | 1.27959598 | 0.03887148 | 0.33771014 |
| ENSMUSG00000047371 | Zfp768 | 1.2625044 | 0.03895635 | 0.33833176 |
| ENSMUSG00000027680 | Fxr1 | 1.14249366 | 0.03906355 | 0.3391469 |
| ENSMUSG00000006221 | Hspb7 | -3.553 | 0.0391024 | 0.33931392 |
| ENSMUSG00000074749 | Kiz | 1.27544084 | 0.0391289 | 0.33931392 |
| ENSMUSG00000069266 | H4c2 | 5.16265217 | 0.03913405 | 0.33931392 |
| ENSMUSG00000039956 | Mrap | -2.6696757 | 0.0391362 | 0.33931392 |
| ENSMUSG00000036120 | Rfxank | 1.20776927 | 0.03915646 | 0.33937376 |
| ENSMUSG00000022358 | Fbxo32 | 2.17415629 | 0.03921599 | 0.33977383 |
| ENSMUSG00000024800 | Rpp30 | 1.38816033 | 0.03923591 | 0.33977741 |
| ENSMUSG00000030232 | Aebp2 | 1.19968513 | 0.03925752 | 0.33977741 |
| ENSMUSG00000029208 | Guf1 | 1.41875519 | 0.03927207 | 0.33977741 |
| ENSMUSG00000037418 | Best1 | 2.27391472 | 0.03927653 | 0.33977741 |
| ENSMUSG00000021750 | Fam107a | -9.5370384 | 0.03928865 | 0.33977741 |
| ENSMUSG00000114456 | H2bc9 | 2.98773946 | 0.03930852 | 0.33977741 |
| ENSMUSG00000076437 | Selenoh | 1.29689305 | 0.0393383 | 0.33977741 |
| ENSMUSG00000082163 | Gm14276 | 1.81146527 | 0.03934524 | 0.33977741 |
| ENSMUSG00000051451 | Crebzf | 1.39155836 | 0.03934634 | 0.33977741 |
| ENSMUSG00000021770 | Samd8 | -1.3228246 | 0.03935011 | 0.33977741 |
| ENSMUSG00000022026 | Olfm4 | -6.0505637 | 0.03936871 | 0.33982254 |
| ENSMUSG00000019518 | Ap4m1 | 1.267254 | 0.03940943 | 0.34005858 |
| ENSMUSG00000032377 | Plscr4 | -2.0552373 | 0.03945727 | 0.34035579 |
| ENSMUSG00000112932 | Gm48308 | 3.79817488 | 0.03951588 | 0.34070748 |
| ENSMUSG00000043872 | Zmym1 | 1.35919486 | 0.03953098 | 0.34070748 |
| ENSMUSG00000031285 | Dcx | 17.1986044 | 0.03953826 | 0.34070748 |
| ENSMUSG00000014776 | Nol3 | -1.8203234 | 0.03957335 | 0.34086418 |
| ENSMUSG00000071253 | Slc25a16 | 1.2700821 | 0.03959308 | 0.34086418 |
| ENSMUSG00000021065 | Fut8 | -1.7255667 | 0.03959669 | 0.34086418 |
| ENSMUSG00000032897 | Nfyc | 1.21196702 | 0.03965428 | 0.34114265 |
| ENSMUSG00000032454 | Rbp2 | -12.952476 | 0.03965589 | 0.34114265 |
| ENSMUSG00000027859 | Ngf | -2.1958546 | 0.03969908 | 0.341286 |
| ENSMUSG00000028419 | Chmp5 | -1.1829819 | 0.03969941 | 0.341286 |
| ENSMUSG00000026228 | Htr2b | -7.077224 | 0.03971429 | 0.34129846 |
| ENSMUSG00000026357 | Rgs18 | -2.7000545 | 0.03976887 | 0.34165201 |
| ENSMUSG00000031849 | Comp | -3.5188888 | 0.03983058 | 0.34206646 |
| ENSMUSG00000003660 | Snrnp200 | 1.31736825 | 0.03984574 | 0.34208103 |
| ENSMUSG00000056608 | Chd9 | 1.49405568 | 0.03986427 | 0.34212452 |
| ENSMUSG00000020654 | Adcy3 | -1.4963685 | 0.03988153 | 0.34215711 |
| ENSMUSG00000022680 | Pdxdc1 | 1.19273093 | 0.03995497 | 0.34267149 |
| ENSMUSG00000036615 | Rfxap | 1.20555481 | 0.04000504 | 0.34298518 |
| ENSMUSG00000098221 | Gm27030 | -20.678424 | 0.04003025 | 0.34308554 |
| ENSMUSG00000027667 | Zfp639 | 1.1984335 | 0.04005065 | 0.34314471 |
| ENSMUSG00000020889 | Nr1d1 | -1.6373808 | 0.04010019 | 0.34320476 |
| ENSMUSG00000070392 | Gm20634 | -2.0743863 | 0.04010505 | 0.34320476 |

| | | | | |
|---|---|---|---|---|
| ENSMUSG00000054604 | Cggbp1 | 1.14202917 | 0.04011236 | 0.34320476 |
| ENSMUSG00000029725 | Ppp1r35 | -1.239379 | 0.04012295 | 0.34320476 |
| ENSMUSG00000048574 | Ccnb1-ps | 7.40153167 | 0.04012519 | 0.34320476 |
| ENSMUSG00000070348 | Ccnd1 | 1.34473968 | 0.04016751 | 0.34345111 |
| ENSMUSG00000052837 | Junb | -1.6521063 | 0.04023021 | 0.34387154 |
| ENSMUSG00000024238 | Zeb1 | -1.7565705 | 0.04038492 | 0.34497415 |
| ENSMUSG00000049807 | Arhgap23 | -1.3134436 | 0.04039335 | 0.34497415 |
| ENSMUSG00000039782 | Cpeb2 | -1.6042462 | 0.04041336 | 0.34497415 |
| ENSMUSG00000021589 | Rhobtb3 | 1.28444129 | 0.04041351 | 0.34497415 |
| ENSMUSG00000029364 | Wsb2 | -1.2526124 | 0.04044875 | 0.34500951 |
| ENSMUSG00000039994 | Timeless | 1.24877145 | 0.04046757 | 0.34500951 |
| ENSMUSG00000094777 | Hist1h2ap | 12.8689998 | 0.04047019 | 0.34500951 |
| ENSMUSG00000026276 | Septin2 | 1.25462397 | 0.04047196 | 0.34500951 |
| ENSMUSG00000086742 | Gm16201 | -2.4701308 | 0.0405191 | 0.34517164 |
| ENSMUSG00000032946 | Rasgrp2 | -2.193987 | 0.04052929 | 0.34517164 |
| ENSMUSG00000117891 | Gm41764 | 25.7806662 | 0.04054575 | 0.34517164 |
| ENSMUSG00000095028 | Sirpb1b | -1.9242996 | 0.04055566 | 0.34517164 |
| ENSMUSG00000027330 | Cdc25b | 1.4553915 | 0.04055889 | 0.34517164 |
| ENSMUSG00000046574 | Prr12 | 1.31307812 | 0.04059333 | 0.34534904 |
| ENSMUSG00000056228 | Cars2 | 1.43168415 | 0.04064501 | 0.3455595 |
| ENSMUSG00000087403 | Kantr | 1.41920408 | 0.04064886 | 0.3455595 |
| ENSMUSG00000018821 | Avpi1 | -1.485902 | 0.04068257 | 0.3455595 |
| ENSMUSG00000033863 | Klf9 | -1.3589235 | 0.04068524 | 0.3455595 |
| ENSMUSG00000035551 | Igfbpl1 | -69.546175 | 0.04068606 | 0.3455595 |
| ENSMUSG00000028894 | Inpp5b | -1.3997886 | 0.04070359 | 0.34559296 |
| ENSMUSG00000083083 | Gm15382 | -12.654324 | 0.04080994 | 0.34638013 |
| ENSMUSG00000115311 | Gm35823 | 8.19966263 | 0.04085991 | 0.34658444 |
| ENSMUSG00000003541 | Ier3 | -1.9971137 | 0.04086128 | 0.34658444 |
| ENSMUSG00000002797 | Ggct | -1.4490795 | 0.04088801 | 0.34669543 |
| ENSMUSG00000024922 | Ovol1 | -2.312464 | 0.04090786 | 0.34674799 |
| ENSMUSG00000037958 | Nsrp1 | -1.1963104 | 0.04093447 | 0.34685785 |
| ENSMUSG00000029474 | Rnf34 | -1.1669291 | 0.04095093 | 0.34688164 |
| ENSMUSG00000061175 | Fnip2 | -1.5002209 | 0.04101479 | 0.34730688 |
| ENSMUSG00000026816 | Gtf3c5 | 1.20935263 | 0.04110015 | 0.34790368 |
| ENSMUSG00000048007 | Timm8a1 | 1.46184596 | 0.04111265 | 0.34790368 |
| ENSMUSG00000037316 | Bag4 | 1.22479438 | 0.04117386 | 0.34830565 |
| ENSMUSG00000070323 | Mmp27 | -2.8880167 | 0.04120188 | 0.34832725 |
| ENSMUSG00000037428 | Vgf | -6.3428956 | 0.04120383 | 0.34832725 |
| ENSMUSG00000022286 | Grhl2 | 1.32168987 | 0.04124001 | 0.34842129 |
| ENSMUSG00000004266 | Ptpn6 | -1.3970088 | 0.04124237 | 0.34842129 |
| ENSMUSG00000019158 | Tmem160 | -1.4439138 | 0.04126965 | 0.34850655 |
| ENSMUSG00000032264 | Zw10 | 1.22135826 | 0.0412799 | 0.34850655 |
| ENSMUSG00000025423 | Pias2 | 1.21338344 | 0.04129697 | 0.3485349 |
| ENSMUSG00000107043 | Gm42849 | 3.47564815 | 0.04138173 | 0.34900142 |
| ENSMUSG00000062312 | Erbb2 | 1.33394982 | 0.04139187 | 0.34900142 |

| | | | | |
|---|---|---|---|---|
| ENSMUSG00000061411 | Nol4l | 1.45454992 | 0.04139345 | 0.34900142 |
| ENSMUSG00000047417 | Rexo1 | 1.23576254 | 0.04141032 | 0.34902785 |
| ENSMUSG00000066057 | Gm1976 | 1.42962705 | 0.04146821 | 0.34930473 |
| ENSMUSG00000028062 | Lamtor2 | -1.2272021 | 0.04147066 | 0.34930473 |
| ENSMUSG00000058503 | Fam133b | 1.1898933 | 0.04150244 | 0.34945663 |
| ENSMUSG00000060803 | Gstp1 | -1.2949847 | 0.0415232 | 0.34951556 |
| ENSMUSG00000030306 | Tmtc1 | -2.1326799 | 0.0416461 | 0.35043402 |
| ENSMUSG00000026739 | Bmi1 | 1.22024421 | 0.04170016 | 0.35077272 |
| ENSMUSG00000038781 | Stap2 | 1.67869728 | 0.04174889 | 0.35105366 |
| ENSMUSG00000029136 | Rbks | -1.6779135 | 0.04176927 | 0.35105366 |
| ENSMUSG00000050390 | C77080 | 1.274318 | 0.041775 | 0.35105366 |
| ENSMUSG00000029992 | Gfpt1 | 1.1981463 | 0.0419325 | 0.35226068 |
| ENSMUSG00000059201 | Lep | -3.4280936 | 0.04199159 | 0.35261638 |
| ENSMUSG00000032263 | Bckdhb | 1.39949397 | 0.04200259 | 0.35261638 |
| ENSMUSG00000029635 | Cdk8 | 1.2103413 | 0.04204995 | 0.35278633 |
| ENSMUSG00000026836 | Acvr1 | -1.3110547 | 0.04205194 | 0.35278633 |
| ENSMUSG00000046805 | Mpeg1 | -1.3753551 | 0.0420653 | 0.35278633 |
| ENSMUSG00000033825 | Tpsb2 | -2.8922953 | 0.04207836 | 0.35278633 |
| ENSMUSG00000025059 | Gk | 1.51411615 | 0.0421237 | 0.35304998 |
| ENSMUSG00000029330 | Cds1 | 2.4962015 | 0.0421625 | 0.35323182 |
| ENSMUSG00000034848 | Ttc21b | 1.25995441 | 0.0421732 | 0.35323182 |
| ENSMUSG00000054418 | 2900041M22 | -2.8603713 | 0.04219704 | 0.35331504 |
| ENSMUSG00000075217 | Fads2b | -12.59396 | 0.04221488 | 0.35334802 |
| ENSMUSG00000030495 | Slc7a10 | -4.9665991 | 0.04225648 | 0.35357972 |
| ENSMUSG00000117599 | Gm49971 | 5.64455727 | 0.04231063 | 0.3539163 |
| ENSMUSG00000115009 | G930009F23 | 2.22809234 | 0.04240289 | 0.35428962 |
| ENSMUSG00000032860 | P2ry2 | -1.5196595 | 0.0424072 | 0.35428962 |
| ENSMUSG00000036893 | Ehmt1 | 1.20785717 | 0.0424257 | 0.35428962 |
| ENSMUSG00000002968 | Med25 | 1.19065196 | 0.04242607 | 0.35428962 |
| ENSMUSG00000027883 | Gpsm2 | 1.36666632 | 0.04245015 | 0.35428962 |
| ENSMUSG00000105039 | Gm32585 | 5.15954535 | 0.04245023 | 0.35428962 |
| ENSMUSG00000028271 | Gtf2b | -1.2230395 | 0.04246216 | 0.35428962 |
| ENSMUSG00000058392 | Rrp1b | 1.39200141 | 0.04248281 | 0.35428962 |
| ENSMUSG00000007656 | Arpp19 | 1.22500452 | 0.04250225 | 0.35428962 |
| ENSMUSG00000019843 | Fyn | -1.4398512 | 0.04250345 | 0.35428962 |
| ENSMUSG00000029135 | Fosl2 | -1.7021256 | 0.04250913 | 0.35428962 |
| ENSMUSG00000034361 | Cpne2 | -1.4005789 | 0.04254776 | 0.35428962 |
| ENSMUSG00000029687 | Ezh2 | 1.30082067 | 0.04255739 | 0.35428962 |
| ENSMUSG00000041889 | Shisa4 | -1.9920798 | 0.0425608 | 0.35428962 |
| ENSMUSG00000067149 | Jchain | -11.896523 | 0.04256439 | 0.35428962 |
| ENSMUSG00000047388 | Atmin | 1.2487614 | 0.04262007 | 0.35463694 |
| ENSMUSG00000045394 | Epcam | 1.34322709 | 0.04264962 | 0.35476667 |
| ENSMUSG00000031292 | Cdkl5 | 1.61904976 | 0.04268648 | 0.35495709 |
| ENSMUSG00000040495 | Chrm4 | 6.45153755 | 0.04271766 | 0.35510015 |
| ENSMUSG00000024521 | Pmaip1 | -1.5487837 | 0.04279507 | 0.35559602 |

| | | | | |
|---|---|---|---|---|
| ENSMUSG00000040177 | 2310057M21 | 1.31187485 | 0.04281586 | 0.35559602 |
| ENSMUSG00000106688 | Gm42851 | 3.12840186 | 0.04281929 | 0.35559602 |
| ENSMUSG00000037373 | Ctbp1 | 1.17254877 | 0.04287272 | 0.35582849 |
| ENSMUSG00000001348 | Acp5 | -1.4247668 | 0.04287529 | 0.35582849 |
| ENSMUSG00000002833 | Hdgfl2 | 1.21494896 | 0.04290825 | 0.35598576 |
| ENSMUSG00000076615 | Ighg3 | -33.827355 | 0.04293125 | 0.35606034 |
| ENSMUSG00000031494 | Cd209a | -2.8458752 | 0.04295524 | 0.3561431 |
| ENSMUSG00000031377 | Bmx | -3.6238132 | 0.04301304 | 0.35650596 |
| ENSMUSG00000032537 | Ephb1 | 2.50619098 | 0.04304561 | 0.35657216 |
| ENSMUSG00000024304 | Cdh2 | -2.3697544 | 0.04304909 | 0.35657216 |
| ENSMUSG00000105076 | A930003O13 | -4.6318158 | 0.04308534 | 0.3567562 |
| ENSMUSG00000000902 | Smarcb1 | 1.13790089 | 0.04311303 | 0.35685645 |
| ENSMUSG00000033576 | Apol6 | -3.0336087 | 0.04312554 | 0.35685645 |
| ENSMUSG00000033365 | Ipo13 | 1.29882113 | 0.04315853 | 0.35701324 |
| ENSMUSG00000020873 | Slc35b1 | -1.2187728 | 0.04318698 | 0.35708136 |
| ENSMUSG00000047592 | Nxpe5 | -2.249658 | 0.04319487 | 0.35708136 |
| ENSMUSG00000032320 | Rcn2 | -1.2047516 | 0.04322962 | 0.35721168 |
| ENSMUSG00000028544 | Slc5a9 | 3.39121177 | 0.04324118 | 0.35721168 |
| ENSMUSG00000041842 | Fhdc1 | 1.2567332 | 0.04325281 | 0.35721168 |
| ENSMUSG00000032246 | Calml4 | -3.075433 | 0.04327578 | 0.3572853 |
| ENSMUSG00000028996 | Rbp7 | 2.51023574 | 0.0433412 | 0.35753653 |
| ENSMUSG00000026712 | Mrc1 | -2.0014679 | 0.04334666 | 0.35753653 |
| ENSMUSG00000108218 | Olfr1372-ps1 | 3.50298728 | 0.04334842 | 0.35753653 |
| ENSMUSG00000005054 | Cstb | -1.3309651 | 0.04338168 | 0.3576948 |
| ENSMUSG00000036832 | Lpar3 | -3.8251084 | 0.04343729 | 0.35803713 |
| ENSMUSG00000055435 | Maf | -1.5412217 | 0.04350026 | 0.3584399 |
| ENSMUSG00000048503 | Tlcd5 | -1.8414214 | 0.04353897 | 0.35864257 |
| ENSMUSG00000038047 | Haus6 | 1.33397343 | 0.0436455 | 0.35931496 |
| ENSMUSG00000091709 | Gm17189 | 3.99659368 | 0.04365437 | 0.35931496 |
| ENSMUSG00000017309 | Cd300lg | -1.6307952 | 0.04366469 | 0.35931496 |
| ENSMUSG00000028977 | Casz1 | 1.38489379 | 0.04367716 | 0.35931496 |
| ENSMUSG00000066682 | Pilrb2 | -2.024984 | 0.04369163 | 0.35931767 |
| ENSMUSG00000022519 | Srl | 1.4901089 | 0.04370593 | 0.35931905 |
| ENSMUSG00000086606 | Gm13205 | 2.22805299 | 0.04374408 | 0.35951637 |
| ENSMUSG00000036940 | Kdm1a | 1.13988282 | 0.04379467 | 0.35980437 |
| ENSMUSG00000006715 | Gmnn | 1.43254384 | 0.04380744 | 0.35980437 |
| ENSMUSG00000021192 | Golga5 | -1.2024822 | 0.0438426 | 0.3599768 |
| ENSMUSG00000027324 | Rpusd2 | 1.37076797 | 0.04386222 | 0.36002159 |
| ENSMUSG00000072494 | Ppp1r3e | 1.34967786 | 0.0439557 | 0.36067232 |
| ENSMUSG00000032691 | Nlrp3 | -1.6340402 | 0.04401499 | 0.36104225 |
| ENSMUSG00000021250 | Fos | -2.4938505 | 0.04405118 | 0.3612225 |
| ENSMUSG00000040405 | Havcr1 | -15.287643 | 0.04406894 | 0.36123044 |
| ENSMUSG00000038663 | Fsd2 | 2.90050748 | 0.04408058 | 0.36123044 |
| ENSMUSG00000103621 | Gm38366 | 3.10200739 | 0.04415075 | 0.36157694 |
| ENSMUSG00000044906 | 4930503L19F | -1.5211709 | 0.04415132 | 0.36157694 |

| | | | | |
|---|---|---|---|---|
| ENSMUSG00000026630 | Batf3 | -1.8527611 | 0.04421377 | 0.36197176 |
| ENSMUSG00000031605 | Klhl2 | -1.313707 | 0.04425595 | 0.36214932 |
| ENSMUSG00000026174 | Cnot9 | 1.22765748 | 0.04426396 | 0.36214932 |
| ENSMUSG00000024663 | Rab3il1 | -1.3831372 | 0.04429673 | 0.36230075 |
| ENSMUSG00000038349 | Plcl1 | -1.8867743 | 0.04437054 | 0.36278771 |
| ENSMUSG00000038375 | Trp53inp2 | -1.5327008 | 0.04441025 | 0.36299556 |
| ENSMUSG00000037600 | Kdf1 | 1.29410785 | 0.04444636 | 0.36317396 |
| ENSMUSG00000054978 | Kbtbd13 | 3.61139765 | 0.04452143 | 0.36367038 |
| ENSMUSG00000026341 | Actr3 | -1.1304696 | 0.04458584 | 0.36405868 |
| ENSMUSG00000034152 | Exoc3 | 1.15893997 | 0.04461077 | 0.36405868 |
| ENSMUSG00000024621 | Csf1r | -1.3881817 | 0.04461194 | 0.36405868 |
| ENSMUSG00000000127 | Fer | 1.20900929 | 0.04464248 | 0.36419096 |
| ENSMUSG00000050288 | Fzd2 | -1.3578139 | 0.04466841 | 0.36428555 |
| ENSMUSG00000097842 | 9330104G04 | 2.10974011 | 0.04471535 | 0.3645513 |
| ENSMUSG00000028300 | C9orf72 | -1.2486463 | 0.04476552 | 0.36482089 |
| ENSMUSG00000037994 | Slc9b2 | -2.5584659 | 0.04477713 | 0.36482089 |
| ENSMUSG00000039801 | Cplane1 | 1.33524775 | 0.0448149 | 0.36488494 |
| ENSMUSG00000038827 | Abitram | 1.23213436 | 0.04481824 | 0.36488494 |
| ENSMUSG00000020483 | Dynll2 | 1.25922189 | 0.04482836 | 0.36488494 |
| ENSMUSG00000040658 | Dnph1 | 1.59919597 | 0.04484242 | 0.36488494 |
| ENSMUSG00000024670 | Cd6 | -2.1883024 | 0.04489622 | 0.36515941 |
| ENSMUSG00000019984 | Med23 | 1.24266534 | 0.04490489 | 0.36515941 |
| ENSMUSG00000023066 | Rttn | 1.36818601 | 0.04493889 | 0.36531894 |
| ENSMUSG00000097795 | Gm7678 | 28.0574009 | 0.04497519 | 0.3654971 |
| ENSMUSG00000003438 | Timm50 | 1.25735398 | 0.04500707 | 0.36563932 |
| ENSMUSG00000004865 | Srpk1 | 1.23600497 | 0.045071 | 0.36599775 |
| ENSMUSG00000085945 | 2310014F06I | 2.74414721 | 0.04509265 | 0.36599775 |
| ENSMUSG00000022537 | Tmem44 | 1.97154501 | 0.0450944 | 0.36599775 |
| ENSMUSG00000023046 | Igfbp6 | -2.3509621 | 0.0451543 | 0.36633618 |
| ENSMUSG00000039257 | Vstm2b | -13.60065 | 0.04517855 | 0.36633618 |
| ENSMUSG00000055943 | Emc7 | -1.2165408 | 0.04520327 | 0.36633618 |
| ENSMUSG00000116673 | A630089N07 | 1.72043269 | 0.04520749 | 0.36633618 |
| ENSMUSG00000056069 | Otulinl | -1.4308381 | 0.04520818 | 0.36633618 |
| ENSMUSG00000089817 | Gm7162 | 3.45956533 | 0.04523514 | 0.3664378 |
| ENSMUSG00000027167 | Elp4 | 1.22030472 | 0.04533595 | 0.36713741 |
| ENSMUSG00000031984 | 2810004N23 | 1.17935425 | 0.04542613 | 0.3677505 |
| ENSMUSG00000054150 | Syne3 | -1.3135013 | 0.04545063 | 0.36783166 |
| ENSMUSG00000018654 | Ikzf1 | -1.4818114 | 0.04552449 | 0.36831211 |
| ENSMUSG00000026718 | Stam | 1.19625961 | 0.04558755 | 0.36860709 |
| ENSMUSG00000094796 | BC147527 | -1.8785934 | 0.04558996 | 0.36860709 |
| ENSMUSG00000040016 | Ptger3 | -2.7517283 | 0.04564936 | 0.36896999 |
| ENSMUSG00000031841 | Cdh13 | -1.5967293 | 0.04574341 | 0.3695132 |
| ENSMUSG00000096140 | Ankrd66 | -4.4216586 | 0.04574565 | 0.3695132 |
| ENSMUSG00000033633 | Clec18a | 22.0313818 | 0.04577072 | 0.36959818 |
| ENSMUSG00000009894 | Snap47 | -1.2846932 | 0.04582084 | 0.3696783 |

| | | | | |
|---|---|---|---|---|
| ENSMUSG00000037926 | Ssh2 | -1.3418122 | 0.04582851 | 0.3696783 |
| ENSMUSG00000054892 | Txk | -2.3099796 | 0.04582863 | 0.3696783 |
| ENSMUSG00000064264 | Zfp428 | 1.47775526 | 0.04583987 | 0.3696783 |
| ENSMUSG00000025955 | Akr1cl | -27.824332 | 0.04585338 | 0.3696783 |
| ENSMUSG00000069763 | Tmem100 | -2.3577232 | 0.04589739 | 0.36985479 |
| ENSMUSG00000022040 | Ephx2 | -2.0909924 | 0.04591671 | 0.36985479 |
| ENSMUSG00000007097 | Atp1a2 | -2.9831933 | 0.04593024 | 0.36985479 |
| ENSMUSG00000063663 | Brwd3 | 1.40232892 | 0.04593348 | 0.36985479 |
| ENSMUSG00000074238 | Ap1ar | 1.30492879 | 0.04597028 | 0.36990163 |
| ENSMUSG00000095766 | Gm21182 | 5.34540134 | 0.04598198 | 0.36990163 |
| ENSMUSG00000063480 | Snu13 | 1.19029898 | 0.04598297 | 0.36990163 |
| ENSMUSG00000036502 | Tmem255a | 1.91999632 | 0.04602174 | 0.37009635 |
| ENSMUSG00000024370 | Cdc23 | 1.15900696 | 0.04606332 | 0.37031351 |
| ENSMUSG00000081534 | Slc48a1 | -1.2156958 | 0.04611628 | 0.37058625 |
| ENSMUSG00000042279 | H1f8 | 41.9046206 | 0.04612654 | 0.37058625 |
| ENSMUSG00000026281 | Dtymk | 1.27975792 | 0.04615519 | 0.37058625 |
| ENSMUSG00000027544 | Nfatc2 | -1.7967352 | 0.0461689 | 0.37058625 |
| ENSMUSG00000067928 | Zfp760 | 1.4564309 | 0.04617397 | 0.37058625 |
| ENSMUSG00000027881 | Prpf38b | 1.27298867 | 0.04618965 | 0.37058625 |
| ENSMUSG00000034773 | Hrob | 1.50313601 | 0.04620447 | 0.37058625 |
| ENSMUSG00000030272 | Camk1 | 1.27098407 | 0.04622224 | 0.37058625 |
| ENSMUSG00000001418 | Glmp | -1.2888138 | 0.04622849 | 0.37058625 |
| ENSMUSG00000026941 | Mamdc4 | 2.74623952 | 0.04627475 | 0.37076792 |
| ENSMUSG00000035852 | Misp | 2.04272761 | 0.04628033 | 0.37076792 |
| ENSMUSG00000026784 | Pdss1 | 1.34065321 | 0.04630101 | 0.37081666 |
| ENSMUSG00000090145 | Ugt1a6b | -12.355255 | 0.04633191 | 0.37086137 |
| ENSMUSG00000003545 | Fosb | -3.2036634 | 0.04633578 | 0.37086137 |
| ENSMUSG00000027452 | Acss1 | 1.82420623 | 0.04640596 | 0.37117099 |
| ENSMUSG00000087413 | Gm11266 | 2.05871912 | 0.04641355 | 0.37117099 |
| ENSMUSG00000066568 | Lsm14a | 1.13635839 | 0.04641828 | 0.37117099 |
| ENSMUSG00000022119 | Rbm26 | 1.28035364 | 0.04645216 | 0.37132501 |
| ENSMUSG00000024613 | Tcof1 | 1.35049226 | 0.04648343 | 0.37145816 |
| ENSMUSG00000032615 | Nt5m | -1.3960808 | 0.04650291 | 0.37149697 |
| ENSMUSG00000032418 | Me1 | -2.1552201 | 0.04653841 | 0.37166376 |
| ENSMUSG00000100150 | Gm19585 | -3.6009731 | 0.04656187 | 0.37173429 |
| ENSMUSG00000022043 | Trim35 | -1.2828249 | 0.04662236 | 0.3721003 |
| ENSMUSG00000039988 | Ankrd13c | 1.17657884 | 0.04668365 | 0.37232446 |
| ENSMUSG00000046841 | Ckap4 | -1.5012523 | 0.04668748 | 0.37232446 |
| ENSMUSG00000023041 | Krt6b | -55.256489 | 0.0466944 | 0.37232446 |
| ENSMUSG00000024081 | Cebpz | 1.2571256 | 0.04671919 | 0.3724053 |
| ENSMUSG00000040522 | Tlr8 | -1.7881097 | 0.04676728 | 0.37251352 |
| ENSMUSG00000061759 | Armt1 | 1.22900034 | 0.04677257 | 0.37251352 |
| ENSMUSG00000034430 | Zxdc | 1.23245165 | 0.04678955 | 0.37251352 |
| ENSMUSG00000027983 | Cyp2u1 | -1.8060524 | 0.0467914 | 0.37251352 |
| ENSMUSG00000042605 | Atxn2 | 1.23933332 | 0.04682367 | 0.37265362 |

| | | | | |
|---|---|---|---|---|
| ENSMUSG00000022945 | Chaf1b | 1.41385242 | 0.04696365 | 0.37355876 |
| ENSMUSG00000038916 | Soga3 | 15.5568934 | 0.04697753 | 0.37355876 |
| ENSMUSG00000071001 | Hrct1 | -1.7976686 | 0.0469823 | 0.37355876 |
| ENSMUSG00000028792 | Ak2 | 1.22894665 | 0.0469962 | 0.37355876 |
| ENSMUSG00000022248 | Rad1 | 1.39358261 | 0.0470266 | 0.37368352 |
| ENSMUSG00000047632 | Fgfbp3 | 1.61378485 | 0.04714074 | 0.37440408 |
| ENSMUSG00000026510 | Trp53bp2 | 1.30160517 | 0.04714674 | 0.37440408 |
| ENSMUSG00000033740 | St18 | -4.4039781 | 0.04717659 | 0.37445922 |
| ENSMUSG00000020733 | Slc9a3r1 | 1.36173579 | 0.04718316 | 0.37445922 |
| ENSMUSG00000018425 | Dhx40 | -1.2282836 | 0.04721599 | 0.37460279 |
| ENSMUSG00000049792 | Bag5 | -1.2546498 | 0.04723717 | 0.37465389 |
| ENSMUSG00000018509 | Cenpv | -1.4144885 | 0.04731468 | 0.375061 |
| ENSMUSG00000028886 | Eya3 | 1.24204767 | 0.04731802 | 0.375061 |
| ENSMUSG00000012535 | Tnpo3 | 1.21151669 | 0.04733677 | 0.3750926 |
| ENSMUSG00000036450 | Hif1an | 1.19200722 | 0.04736499 | 0.37519923 |
| ENSMUSG00000031897 | Psmb10 | -1.4514185 | 0.04741963 | 0.37547614 |
| ENSMUSG00000062352 | Itgb1bp1 | -1.2633382 | 0.0474295 | 0.37547614 |
| ENSMUSG00000031138 | F9 | -5.4843243 | 0.04745479 | 0.37555934 |
| ENSMUSG00000033335 | Dnm2 | -1.1686346 | 0.04750432 | 0.37576479 |
| ENSMUSG00000034452 | Slc24a1 | 4.87978328 | 0.04751032 | 0.37576479 |
| ENSMUSG00000105843 | Gm19439 | 11.843609 | 0.04754724 | 0.37593979 |
| ENSMUSG00000031144 | Syp | -2.8625249 | 0.04756284 | 0.37594617 |
| ENSMUSG00000115855 | Gm34643 | 5.17034013 | 0.04758319 | 0.37599009 |
| ENSMUSG00000073485 | H3f3aos | 2.53502567 | 0.0476767 | 0.37657721 |
| ENSMUSG00000030347 | D6Wsu163e | 1.20091798 | 0.04768713 | 0.37657721 |
| ENSMUSG00000035900 | Gramd4 | 1.33324593 | 0.04778781 | 0.37725501 |
| ENSMUSG00000053687 | Dpep2 | -1.9355916 | 0.04783393 | 0.3773015 |
| ENSMUSG00000002103 | Acp2 | -1.1970281 | 0.04784948 | 0.3773015 |
| ENSMUSG00000003227 | Edar | -3.3108777 | 0.04785272 | 0.3773015 |
| ENSMUSG00000036570 | Fxyd1 | -2.9818783 | 0.04785309 | 0.3773015 |
| ENSMUSG00000039108 | Lsm14b | 1.22284951 | 0.04788187 | 0.37741128 |
| ENSMUSG00000029475 | Kdm2b | 1.27251504 | 0.0479724 | 0.37795068 |
| ENSMUSG00000047787 | Flrt1 | -5.2596624 | 0.04798004 | 0.37795068 |
| ENSMUSG00000020627 | Klhl29 | -1.7713994 | 0.0480204 | 0.37807665 |
| ENSMUSG00000097124 | A530020G20 | 2.19081436 | 0.04802579 | 0.37807665 |
| ENSMUSG00000095042 | Gm12537 | -22.166949 | 0.04806001 | 0.37813939 |
| ENSMUSG00000000531 | Grasp | -1.5503815 | 0.04806749 | 0.37813939 |
| ENSMUSG00000027342 | Pcna | 1.26682962 | 0.04808254 | 0.37813939 |
| ENSMUSG00000051124 | Gimap9 | -1.5227481 | 0.04810841 | 0.37813939 |
| ENSMUSG00000033411 | Ctdspl2 | 1.3037892 | 0.0481124 | 0.37813939 |
| ENSMUSG00000020363 | Gfpt2 | -1.83153 | 0.04812304 | 0.37813939 |
| ENSMUSG00000022227 | Mcpt1 | -74.464316 | 0.04821237 | 0.37872417 |
| ENSMUSG00000017715 | Pgs1 | -1.1949549 | 0.04834592 | 0.37965593 |
| ENSMUSG00000069270 | H2ac6 | 1.88635172 | 0.04836715 | 0.3796651 |
| ENSMUSG00000040562 | Gstm2 | -1.5895641 | 0.04837697 | 0.3796651 |

| | | | | |
|---|---|---|---|---|
| ENSMUSG00000024413 | Npc1 | -1.2900612 | 0.0484143 | 0.37974695 |
| ENSMUSG00000032440 | Tgfbr2 | -1.4612466 | 0.04841729 | 0.37974695 |
| ENSMUSG00000074158 | Zfp976 | 1.57768465 | 0.04853023 | 0.380351 |
| ENSMUSG00000069495 | Epc2 | 1.2678693 | 0.04854891 | 0.380351 |
| ENSMUSG00000014767 | Tbp | 1.25431157 | 0.04855182 | 0.380351 |
| ENSMUSG00000031362 | Xlr4c | -2.6537396 | 0.04855417 | 0.380351 |
| ENSMUSG00000026867 | Gapvd1 | 1.14485361 | 0.04861524 | 0.38059504 |
| ENSMUSG00000028032 | Papss1 | 1.23818141 | 0.04861528 | 0.38059504 |
| ENSMUSG00000026890 | Lhx6 | -1.6959788 | 0.04869109 | 0.38107116 |
| ENSMUSG00000036466 | Megf11 | -2.85562 | 0.04873444 | 0.38129297 |
| ENSMUSG00000041974 | Spidr | 1.4754784 | 0.04878706 | 0.38152457 |
| ENSMUSG00000089854 | Gm16133 | -11.602546 | 0.04881051 | 0.38152457 |
| ENSMUSG00000041737 | Tmem45b | -3.520759 | 0.04881784 | 0.38152457 |
| ENSMUSG00000042557 | Sin3a | 1.1852753 | 0.0488241 | 0.38152457 |
| ENSMUSG00000018378 | Cuedc1 | -1.3521019 | 0.04885327 | 0.38161097 |
| ENSMUSG00000025407 | Gli1 | -2.0025505 | 0.04886519 | 0.38161097 |
| ENSMUSG00000018012 | Rac3 | 1.62820473 | 0.04889407 | 0.38171924 |
| ENSMUSG00000029192 | Tbc1d14 | 1.12189125 | 0.04894683 | 0.38201374 |
| ENSMUSG00000020986 | Sec23a | -1.2374644 | 0.04896522 | 0.38204 |
| ENSMUSG00000060735 | Rxfp3 | -50.72553 | 0.04898992 | 0.38211533 |
| ENSMUSG00000086965 | Rtl10 | 1.94302791 | 0.04901071 | 0.38216024 |
| ENSMUSG00000027439 | Gzf1 | 1.14991021 | 0.04903619 | 0.38223777 |
| ENSMUSG00000036613 | Eipr1 | -1.4934044 | 0.04905074 | 0.38223777 |
| ENSMUSG00000055022 | Cntn1 | -7.7442944 | 0.04908983 | 0.38242516 |
| ENSMUSG00000030138 | Bms1 | 1.18625917 | 0.04913055 | 0.38247019 |
| ENSMUSG00000104903 | Gm43707 | 16.4668025 | 0.049132 | 0.38247019 |
| ENSMUSG00000000282 | Mnt | -1.2279534 | 0.04914506 | 0.38247019 |
| ENSMUSG00000021952 | Xpo4 | 1.31813177 | 0.04915582 | 0.38247019 |
| ENSMUSG00000022969 | Il10rb | -1.2988032 | 0.04922196 | 0.38286759 |
| ENSMUSG00000020228 | Helb | 1.29259508 | 0.04923784 | 0.38287389 |
| ENSMUSG00000082070 | Gm1866 | 9.35597822 | 0.04932898 | 0.38341434 |
| ENSMUSG00000052371 | Hoxd3os1 | -4.0007949 | 0.04933751 | 0.38341434 |
| ENSMUSG00000020009 | Ifngr1 | -1.2325936 | 0.04938978 | 0.3837032 |
| ENSMUSG00000082315 | Gm16523 | 17.9867615 | 0.04943493 | 0.38393654 |
| ENSMUSG00000029203 | Ube2k | 1.13912242 | 0.04946311 | 0.38403805 |
| ENSMUSG00000037669 | Ldah | -1.2496635 | 0.04947884 | 0.38404288 |
| ENSMUSG00000031358 | Msl3 | 1.2724101 | 0.04953935 | 0.38439513 |
| ENSMUSG00000002076 | Hsf2bp | 3.01492264 | 0.04958762 | 0.38461964 |
| ENSMUSG00000020303 | Stc2 | -2.3817104 | 0.04959856 | 0.38461964 |
| ENSMUSG00000068114 | Ccdc134 | 1.26013356 | 0.04968212 | 0.38515013 |
| ENSMUSG00000031878 | Nae1 | 1.18600345 | 0.04970014 | 0.38517233 |
| ENSMUSG00000026159 | Agfg1 | 1.1642027 | 0.04975419 | 0.38547363 |
| ENSMUSG00000041757 | Plekha6 | 1.73385822 | 0.04982767 | 0.38586234 |
| ENSMUSG00000062867 | Impdh2 | 1.31459863 | 0.04983751 | 0.38586234 |
| ENSMUSG00000042197 | Zfp451 | 1.23044747 | 0.04985301 | 0.38586234 |

| | | | | |
|---|---|---|---|---|
| ENSMUSG00000028766 | Alpl | 2.34273614 | 0.0498651 | 0.38586234 |
| ENSMUSG00000039865 | Slc44a3 | 1.53391372 | 0.04993219 | 0.38626389 |
| ENSMUSG00000051185 | Fam174a | -1.2159123 | 0.04999792 | 0.38655412 |
