## Supplemental Table 3-C for "Environmentally Induced Sperm RNAs Transmit Cancer Susceptibility to Offspring in a Mouse Model"

**Table S3.C-Differentially expressed genes in miR-10b offspring's mammary tumors**

| Name | gene_symbol | FC | pvalue | padj |
|---|---|---|---|---|
| ENSMUSG00000085541 | Gm16010 | -2.378E+12 | 4.90E-28 | 1.25E-23 |
| ENSMUSG00000044103 | Il1f9 | -15607479 | 8.45E-18 | 1.07E-13 |
| ENSMUSG00000044485 | Klk1b11 | -1403700.1 | 2.09E-12 | 1.77E-08 |
| ENSMUSG00000061723 | Tnnt3 | -243810.62 | 1.37E-08 | 8.73E-05 |
| ENSMUSG00000092329 | Galnt2l | -7.386E+10 | 3.89E-08 | 0.00019781 |
| ENSMUSG00000017300 | Tnnc2 | -111077.02 | 2.00E-07 | 0.00084816 |
| ENSMUSG00000048981 | Krt31 | -85764.192 | 1.02E-06 | 0.00370856 |
| ENSMUSG00000032807 | Alox12b | -1722840.7 | 1.26E-06 | 0.00386871 |
| ENSMUSG00000063661 | Krt73 | -107797.17 | 1.37E-06 | 0.00386871 |
| ENSMUSG00000056270 | Prr9 | -32230962 | 5.39E-06 | 0.01370262 |
| ENSMUSG00000023041 | Krt6b | 36159.9674 | 6.11E-06 | 0.01412665 |
| ENSMUSG00000005681 | Apoa2 | -40.247704 | 9.60E-05 | 0.20335042 |
| ENSMUSG00000016349 | Eef1a2 | -219419.68 | 0.00012587 | 0.23001288 |
| ENSMUSG00000036863 | Syde2 | -5.1900381 | 0.00012672 | 0.23001288 |
| ENSMUSG00000067147 | Rpl7a-ps11 | 33638878.2 | 0.00013795 | 0.23370278 |
| ENSMUSG00000098387 | Pet117 | 123.13819 | 0.00017571 | 0.25338504 |
| ENSMUSG00000005716 | Pvalb | -36458.232 | 0.00017682 | 0.25338504 |
| ENSMUSG00000045515 | Pou3f3 | -31586913 | 0.00017948 | 0.25338504 |
| ENSMUSG00000008843 | Cldn13 | -28424182 | 0.00020246 | 0.264199 |
| ENSMUSG00000046834 | Krt1 | -80.653079 | 0.00020793 | 0.264199 |
| ENSMUSG00000035305 | Ror1 | -3.9557089 | 0.00026284 | 0.31805727 |
| ENSMUSG00000090121 | Abhd12b | -13321616 | 0.00035643 | 0.40155462 |
| ENSMUSG00000014603 | Alx3 | -101.85481 | 0.00038048 | 0.40155462 |
| ENSMUSG00000085936 | 2610307P16I | 27.7465999 | 0.00038842 | 0.40155462 |
| ENSMUSG00000074195 | Clca4b | 12797240.2 | 0.00039538 | 0.40155462 |
| ENSMUSG00000022512 | Cldn1 | -6.4355761 | 0.00041085 | 0.40155462 |
| ENSMUSG00000026839 | Upp2 | -4.0868001 | 0.00043101 | 0.40565772 |
| ENSMUSG00000089661 | Mia | -29.698373 | 0.00047264 | 0.42895615 |
| ENSMUSG00000093954 | Gm16867 | -9.9572949 | 0.00049266 | 0.43170705 |
| ENSMUSG00000017588 | Krt27 | -8713261.4 | 0.0005266 | 0.44606779 |
| ENSMUSG00000029368 | Alb | -186.76353 | 0.00055328 | 0.45354299 |
| ENSMUSG00000029130 | Rnf32 | -24.130179 | 0.00058613 | 0.46545706 |
| ENSMUSG00000096953 | Gm26571 | 130.879306 | 0.00062919 | 0.48451776 |
| ENSMUSG00000020364 | Zfp354a | -24.499389 | 0.00065748 | 0.49140645 |
| ENSMUSG00000026405 | C4bp | -4173582.5 | 0.00084488 | 0.61343177 |
| ENSMUSG00000030873 | Scnn1b | -26.230409 | 0.00086918 | 0.61354722 |
| ENSMUSG00000046259 | Sprr2h | -3998434 | 0.00091205 | 0.62640278 |
| ENSMUSG00000086231 | Rapgef4os3 | -27.638897 | 0.00110761 | 0.74069821 |
| ENSMUSG00000011034 | Slc5a1 | -22.618052 | 0.00116372 | 0.7541876 |
| ENSMUSG00000030731 | Syt3 | 23.9146065 | 0.00119854 | 0.7541876 |
| ENSMUSG00000041782 | Lad1 | -3.213442 | 0.00121681 | 0.7541876 |
| ENSMUSG00000036687 | Tmem184a | -18.32367 | 0.0012524 | 0.75776369 |
| ENSMUSG00000089736 | Tgfbr3l | -6.4347705 | 0.00131165 | 0.76152603 |

| | | | | |
|---|---|---|---|---|
| ENSMUSG00000029544 | Cabp1 | 6.2941148 | 0.00131856 | 0.76152603 |
| ENSMUSG00000046761 | Fam83h | -2.7416612 | 0.00148025 | 0.82401339 |
| ENSMUSG00000029596 | Sdsl | -6.182259 | 0.00150526 | 0.82401339 |
| ENSMUSG00000031227 | Magee1 | -2.5788734 | 0.00152403 | 0.82401339 |
| ENSMUSG00000054181 | A930012O16 | 24.3637896 | 0.00160062 | 0.84739439 |
| ENSMUSG00000026327 | Serpinb11 | -37575.691 | 0.00178958 | 0.92809595 |
| ENSMUSG00000048013 | Krt35 | -1661409.6 | 0.00188781 | 0.95091951 |
| ENSMUSG00000061048 | Cdh3 | -31.622417 | 0.00190842 | 0.95091951 |
| ENSMUSG00000071858 | Gm94 | -1492971 | 0.00194683 | 0.95140044 |
| ENSMUSG00000104720 | Gm43138 | -40.434887 | 0.00222567 | 1 |
| ENSMUSG00000029032 | Arhgef16 | -16.429183 | 0.00228991 | 1 |
| ENSMUSG00000007279 | Scube2 | -21.212177 | 0.00237525 | 1 |
| ENSMUSG00000049036 | Tmem121 | -22.408519 | 0.00255071 | 1 |
| ENSMUSG00000032013 | Trim29 | -2.8447813 | 0.00260667 | 1 |
| ENSMUSG00000090257 | Gm4524 | -21.15765 | 0.00276503 | 1 |
| ENSMUSG00000081375 | Gm14686 | 43.8187688 | 0.00285957 | 1 |
| ENSMUSG00000025170 | Rab40b | -7.1613718 | 0.00308972 | 1 |
| ENSMUSG00000039676 | Capsl | -15.825774 | 0.00314993 | 1 |
| ENSMUSG00000089901 | Gm8113 | -19.402898 | 0.00321299 | 1 |
| ENSMUSG00000072672 | Duxbl3 | -50.146147 | 0.00342024 | 1 |
| ENSMUSG00000037016 | Frem2 | -17.624455 | 0.00344753 | 1 |
| ENSMUSG00000111348 | Gm19531 | -26.041208 | 0.00371101 | 1 |
| ENSMUSG00000020098 | Pcbd1 | 15.7885407 | 0.0037182 | 1 |
| ENSMUSG00000022203 | Efs | -2.1788159 | 0.00378759 | 1 |
| ENSMUSG00000002007 | Srpk3 | -43.70333 | 0.00384851 | 1 |
| ENSMUSG00000039760 | Il22ra2 | -137.19467 | 0.00388826 | 1 |
| ENSMUSG00000019761 | Krt10 | -10.224782 | 0.00401457 | 1 |
| ENSMUSG00000024935 | Slc1a1 | -18.717145 | 0.00411402 | 1 |
| ENSMUSG00000040525 | Cblc | -14.278967 | 0.00426952 | 1 |
| ENSMUSG00000036585 | Fgf1 | -3.9794253 | 0.00433842 | 1 |
| ENSMUSG00000043782 | Bicdl2 | 14.5811642 | 0.00454698 | 1 |
| ENSMUSG00000020916 | Krt36 | 68188.0009 | 0.00479725 | 1 |
| ENSMUSG00000035228 | Ccdc106 | -11.039788 | 0.00483836 | 1 |
| ENSMUSG00000091476 | Catspere2 | -23.462493 | 0.00489537 | 1 |
| ENSMUSG00000073778 | Faddos | -24.277981 | 0.00499905 | 1 |
| ENSMUSG00000085175 | Gm11423 | 4.45286799 | 0.00523791 | 1 |
| ENSMUSG00000103486 | Gm10657 | 5.62231893 | 0.00532793 | 1 |
| ENSMUSG00000037798 | Mat1a | -32.188398 | 0.00537564 | 1 |
| ENSMUSG00000009248 | Ascl2 | -20.70979 | 0.00544629 | 1 |
| ENSMUSG00000001504 | Irx2 | -3.0270198 | 0.00544914 | 1 |
| ENSMUSG00000039438 | Ttc36 | -23.608462 | 0.00570256 | 1 |
| ENSMUSG00000100455 | Gm29170 | -2.2392832 | 0.00589388 | 1 |
| ENSMUSG00000007591 | Tssk4 | -26.822352 | 0.00594139 | 1 |
| ENSMUSG00000100813 | Gm28874 | -26.721298 | 0.00596651 | 1 |
| ENSMUSG00000037855 | Zfp365 | -3.0548569 | 0.00624906 | 1 |

| | | | | |
|---|---|---|---|---|
| ENSMUSG00000097537 | 2610020C07I | -2.8156883 | 0.00628671 | 1 |
| ENSMUSG00000044461 | Shisa2 | -8.9113641 | 0.0064321 | 1 |
| ENSMUSG00000042195 | Slc35f2 | -18.15297 | 0.00658417 | 1 |
| ENSMUSG00000045545 | Krt14 | -5.3486479 | 0.00660581 | 1 |
| ENSMUSG00000032854 | Ugt8a | -24.491985 | 0.00686702 | 1 |
| ENSMUSG00000087178 | A230056P14I | -4.6855532 | 0.00687568 | 1 |
| ENSMUSG00000035557 | Krt17 | -4.185744 | 0.00707491 | 1 |
| ENSMUSG00000024548 | Setbp1 | -2.8403778 | 0.00712006 | 1 |
| ENSMUSG00000097330 | Gm26672 | 32.6504361 | 0.00724877 | 1 |
| ENSMUSG00000002996 | Hbp1 | 1.27145207 | 0.00736621 | 1 |
| ENSMUSG00000057886 | Cbx3-ps6 | 1.73832742 | 0.00753384 | 1 |
| ENSMUSG00000046085 | 4931422A03I | -23.274566 | 0.00760659 | 1 |
| ENSMUSG00000021509 | Slc25a48 | 15.4090557 | 0.0078476 | 1 |
| ENSMUSG00000065987 | Cd209b | -7.0882811 | 0.00833415 | 1 |
| ENSMUSG00000022218 | Tgm1 | -2.720093 | 0.00838994 | 1 |
| ENSMUSG00000044309 | Apol7c | -4.7561129 | 0.00842771 | 1 |
| ENSMUSG00000029602 | Rasal1 | -4.007569 | 0.00848776 | 1 |
| ENSMUSG00000020159 | Gabrp | -17.726956 | 0.00885818 | 1 |
| ENSMUSG00000016942 | Tmprss6 | -15.813024 | 0.00894287 | 1 |
| ENSMUSG00000025759 | Mfsd8 | -1.5757045 | 0.00913582 | 1 |
| ENSMUSG00000028518 | Prkaa2 | -13.288923 | 0.00918508 | 1 |
| ENSMUSG00000040606 | Kazn | -2.1161693 | 0.00924085 | 1 |
| ENSMUSG00000044667 | Plppr4 | -15.779968 | 0.00925411 | 1 |
| ENSMUSG00000096764 | Gm21985 | -129.78297 | 0.00944588 | 1 |
| ENSMUSG00000054793 | Cadm4 | 2.28180825 | 0.00945135 | 1 |
| ENSMUSG00000025930 | Msc | -8.8082305 | 0.00955538 | 1 |
| ENSMUSG00000031162 | Gata1 | -21.21515 | 0.00957605 | 1 |
| ENSMUSG00000022096 | Hr | -2.0317809 | 0.00961623 | 1 |
| ENSMUSG00000046999 | 1110032F04I | -13.492697 | 0.00974192 | 1 |
| ENSMUSG00000052415 | Tchh | -31.384956 | 0.00990365 | 1 |
| ENSMUSG00000079711 | Smok4a | -23.047693 | 0.00992476 | 1 |
| ENSMUSG00000032012 | Nectin1 | -1.5987492 | 0.00999345 | 1 |
| ENSMUSG00000034117 | Ptgdr2 | -21.910452 | 0.0100086 | 1 |
| ENSMUSG00000097587 | 4930578M01 | 24.5681119 | 0.01019334 | 1 |
| ENSMUSG00000052271 | Bhlha15 | -20.861198 | 0.01025196 | 1 |
| ENSMUSG00000034813 | Grip1 | 15.6190139 | 0.01026301 | 1 |
| ENSMUSG00000031791 | Tmem38a | -1.7030613 | 0.01045474 | 1 |
| ENSMUSG00000078880 | Gm14308 | -4.9383816 | 0.01046764 | 1 |
| ENSMUSG00000000308 | Ckmt1 | 12.6067628 | 0.0107004 | 1 |
| ENSMUSG00000092277 | Gm19684 | -23.328327 | 0.01080779 | 1 |
| ENSMUSG00000055546 | Timd4 | -7.3248325 | 0.01095631 | 1 |
| ENSMUSG00000023873 | 1700010I14R | 10.6926387 | 0.0110628 | 1 |
| ENSMUSG00000103570 | C630004M23 | 22.3543034 | 0.0111864 | 1 |
| ENSMUSG00000031137 | Fgf13 | -8.0134658 | 0.01122853 | 1 |
| ENSMUSG00000118295 | Gm8437 | -5.8880667 | 0.01129583 | 1 |

| | | | | |
|---|---|---|---|---|
| ENSMUSG00000030711 | Sult1a1 | -2.6621775 | 0.01130853 | 1 |
| ENSMUSG00000047150 | 1700001C19I | 13.8174439 | 0.01143418 | 1 |
| ENSMUSG00000003477 | Inmt | -22.351027 | 0.01145929 | 1 |
| ENSMUSG00000058748 | Zfp958 | 1.38267261 | 0.01153928 | 1 |
| ENSMUSG00000047986 | Palm3 | 4.7789578 | 0.0117477 | 1 |
| ENSMUSG00000033295 | Ptprf | -2.3051806 | 0.01213091 | 1 |
| ENSMUSG00000075588 | Hoxb2 | -2.4593084 | 0.01228381 | 1 |
| ENSMUSG00000001583 | Tnk1 | -3.447294 | 0.01228651 | 1 |
| ENSMUSG00000024793 | Tnfrsf25 | -10.451118 | 0.0123924 | 1 |
| ENSMUSG00000107502 | Gm44065 | 18.4869958 | 0.0124194 | 1 |
| ENSMUSG00000071658 | Gng3 | -8.194907 | 0.01243839 | 1 |
| ENSMUSG00000059832 | Kprp | -100624.85 | 0.01245504 | 1 |
| ENSMUSG00000097167 | Gm16740 | -3.0132299 | 0.01249189 | 1 |
| ENSMUSG00000091705 | H2-Q2 | -8.425088 | 0.01265378 | 1 |
| ENSMUSG00000033436 | Armcx2 | -1.9020638 | 0.01267548 | 1 |
| ENSMUSG00000028434 | Epb41l4b | -2.9073676 | 0.01268483 | 1 |
| ENSMUSG00000044393 | Dsg2 | -4.6294079 | 0.01273542 | 1 |
| ENSMUSG00000074595 | Wfdc6a | -23.675208 | 0.01281176 | 1 |
| ENSMUSG00000025993 | Slc40a1 | 2.21045187 | 0.01294886 | 1 |
| ENSMUSG00000033427 | Upb1 | -12.714247 | 0.01313217 | 1 |
| ENSMUSG00000027978 | Prss12 | -1.9863109 | 0.01328485 | 1 |
| ENSMUSG00000009145 | Dqx1 | -6.6298893 | 0.01329012 | 1 |
| ENSMUSG00000048455 | Sprr1b | -89922.732 | 0.01329788 | 1 |
| ENSMUSG00000063245 | Zfp993 | 2.36501415 | 0.01337038 | 1 |
| ENSMUSG00000022865 | Cxadr | -2.6246029 | 0.01349681 | 1 |
| ENSMUSG00000032719 | Sbspon | -11.972327 | 0.01351672 | 1 |
| ENSMUSG00000036805 | Noxa1 | -62446.14 | 0.01361231 | 1 |
| ENSMUSG00000060311 | Mucl1 | -20.89361 | 0.01367425 | 1 |
| ENSMUSG00000046329 | Slc25a23 | -2.2668932 | 0.01375468 | 1 |
| ENSMUSG00000094915 | AC168977.2 | -4.5295678 | 0.01412112 | 1 |
| ENSMUSG00000102813 | Gm37795 | -2.775831 | 0.01426164 | 1 |
| ENSMUSG00000048763 | Hoxb3 | -1.8450034 | 0.01449732 | 1 |
| ENSMUSG00000049670 | Morn4 | -9.4818006 | 0.0145389 | 1 |
| ENSMUSG00000073403 | Gm10499 | -43.399353 | 0.01456162 | 1 |
| ENSMUSG00000073427 | Gm4924 | 2.4145101 | 0.01468617 | 1 |
| ENSMUSG00000042268 | Slc26a9 | 13.6613246 | 0.0149515 | 1 |
| ENSMUSG00000034472 | Rasd2 | -19.02085 | 0.01532001 | 1 |
| ENSMUSG00000091119 | Ccdc152 | -15.698361 | 0.01544552 | 1 |
| ENSMUSG00000058656 | Samd12 | 5.83834532 | 0.01558537 | 1 |
| ENSMUSG00000038260 | Trpm4 | -1.5322598 | 0.01566013 | 1 |
| ENSMUSG00000022211 | Carmil3 | -18.078916 | 0.01569482 | 1 |
| ENSMUSG00000034275 | Igsf9b | -5.056545 | 0.01575625 | 1 |
| ENSMUSG00000096878 | Gm21083 | -22.146084 | 0.01576107 | 1 |
| ENSMUSG00000053821 | Gm9922 | -15.600208 | 0.01590857 | 1 |
| ENSMUSG00000028373 | Astn2 | -14.939189 | 0.01601238 | 1 |

| | | | | |
|---|---|---|---|---|
| ENSMUSG00000047511 | Olfr1396 | -16.757463 | 0.01613963 | 1 |
| ENSMUSG00000022132 | Cldn10 | -5.3702144 | 0.01614724 | 1 |
| ENSMUSG00000102919 | Gm37726 | -31.484085 | 0.01617965 | 1 |
| ENSMUSG00000114788 | Gm48899 | 12.4546095 | 0.01627467 | 1 |
| ENSMUSG00000017692 | Rhbdl3 | -8.7633588 | 0.01629242 | 1 |
| ENSMUSG00000085596 | Gm11476 | 8.53884127 | 0.01654125 | 1 |
| ENSMUSG00000111740 | Gm49783 | -4.222992 | 0.01656431 | 1 |
| ENSMUSG00000096221 | 1500002C15I | -2.077324 | 0.01686783 | 1 |
| ENSMUSG00000041075 | Fzd7 | -1.6829527 | 0.01692992 | 1 |
| ENSMUSG00000018893 | Mb | -4.8029834 | 0.01694975 | 1 |
| ENSMUSG00000058396 | Gpr182 | -3.6666984 | 0.01705908 | 1 |
| ENSMUSG00000104569 | Gm43054 | -7.8342085 | 0.01707079 | 1 |
| ENSMUSG00000020627 | Klhl29 | -2.8259004 | 0.017215 | 1 |
| ENSMUSG00000054161 | Fam83e | -9.4401788 | 0.01722353 | 1 |
| ENSMUSG00000025939 | Ube2w | 1.18369461 | 0.01722832 | 1 |
| ENSMUSG00000061397 | Krt79 | -14.629942 | 0.0172287 | 1 |
| ENSMUSG00000030446 | Zfp273 | 1.52580475 | 0.01742686 | 1 |
| ENSMUSG00000026142 | Rhbdd1 | 1.22273915 | 0.01747078 | 1 |
| ENSMUSG00000112895 | Gm47567 | -7.6719019 | 0.01753269 | 1 |
| ENSMUSG00000050640 | Tmem150c | -3.0689547 | 0.01762686 | 1 |
| ENSMUSG00000106296 | 4632404M16 | -4.5591895 | 0.01765411 | 1 |
| ENSMUSG00000005069 | Pex5 | 1.14825095 | 0.01772789 | 1 |
| ENSMUSG00000026729 | 4930562F07I | -24.081424 | 0.01774135 | 1 |
| ENSMUSG00000033831 | Fgb | -27.506227 | 0.0179569 | 1 |
| ENSMUSG00000022758 | P2rx6 | -9.589925 | 0.01802663 | 1 |
| ENSMUSG00000097350 | 4732491K20I | 3.53092467 | 0.01814497 | 1 |
| ENSMUSG00000058152 | Chsy3 | -5.2044754 | 0.01816577 | 1 |
| ENSMUSG00000006464 | Bbs1 | -2.3102563 | 0.0182263 | 1 |
| ENSMUSG00000020701 | Tmem132e | -21.979937 | 0.01865122 | 1 |
| ENSMUSG00000075012 | Fjx1 | -4.035134 | 0.01865509 | 1 |
| ENSMUSG00000040891 | Foxa3 | 21.1384574 | 0.01873067 | 1 |
| ENSMUSG00000103715 | 4933431K14I | -2.4909474 | 0.01882588 | 1 |
| ENSMUSG00000028289 | Epha7 | -10.087165 | 0.01886761 | 1 |
| ENSMUSG00000027403 | Tgm6 | -50844.471 | 0.01896445 | 1 |
| ENSMUSG00000062376 | Borcs7 | 1.36658056 | 0.01904142 | 1 |
| ENSMUSG00000037552 | Plekhg2 | -1.3445563 | 0.01905646 | 1 |
| ENSMUSG00000021636 | Marveld2 | 2.06816745 | 0.0191643 | 1 |
| ENSMUSG00000053358 | Gm9905 | 6.28109608 | 0.01939125 | 1 |
| ENSMUSG00000097804 | Gm16685 | -10.54871 | 0.01941475 | 1 |
| ENSMUSG00000029504 | Ddx51 | 1.19572672 | 0.019472 | 1 |
| ENSMUSG00000104781 | Gm43303 | 21.3460374 | 0.01987742 | 1 |
| ENSMUSG00000026869 | Psmd5 | 1.18683827 | 0.02008958 | 1 |
| ENSMUSG00000005225 | Plekha8 | -1.641918 | 0.02012481 | 1 |
| ENSMUSG00000037971 | 1110032A03I | -1.6133931 | 0.02020161 | 1 |
| ENSMUSG00000020083 | Fam241b | 9.85541631 | 0.02028046 | 1 |

| | | | | |
|---|---|---|---|---|
| ENSMUSG00000006720 | Zfp184 | 9.63567348 | 0.02030437 | 1 |
| ENSMUSG00000028713 | Cyp4b1 | -2.4234125 | 0.02057744 | 1 |
| ENSMUSG00000110580 | D830024N08 | 8.25943225 | 0.0206191 | 1 |
| ENSMUSG00000104214 | Gm36638 | 12.8462254 | 0.02063056 | 1 |
| ENSMUSG00000028088 | Fmo5 | -2.0515328 | 0.0209189 | 1 |
| ENSMUSG00000063646 | Jakmip1 | -4.3041903 | 0.02092503 | 1 |
| ENSMUSG00000043015 | Nemp2 | -1.4776057 | 0.02093483 | 1 |
| ENSMUSG00000022340 | Sybu | -4.0535114 | 0.02121398 | 1 |
| ENSMUSG00000118057 | B020010K11 | -3.8675387 | 0.02143446 | 1 |
| ENSMUSG00000021604 | Irx4 | -8.2655253 | 0.02156823 | 1 |
| ENSMUSG00000081274 | Gm15727 | -33.323649 | 0.02162882 | 1 |
| ENSMUSG00000001552 | Jup | -1.3759884 | 0.02172413 | 1 |
| ENSMUSG00000026988 | Wdsub1 | -1.2709127 | 0.02182719 | 1 |
| ENSMUSG00000116207 | Nnt | 361.348004 | 0.02230189 | 1 |
| ENSMUSG00000104154 | Gm38104 | 13.9381445 | 0.02258613 | 1 |
| ENSMUSG00000040003 | Magi2 | -1.9067161 | 0.0227071 | 1 |
| ENSMUSG00000031737 | Irx5 | -1.6152362 | 0.0227342 | 1 |
| ENSMUSG00000048728 | Zfp454 | 11.4398882 | 0.02288617 | 1 |
| ENSMUSG00000086894 | Gm15708 | 2.85855531 | 0.02302822 | 1 |
| ENSMUSG00000115344 | Gm49364 | -12.439789 | 0.0231317 | 1 |
| ENSMUSG00000003352 | Cacnb3 | -1.3730843 | 0.02328649 | 1 |
| ENSMUSG00000010721 | Lmbr1 | 1.33408757 | 0.02354991 | 1 |
| ENSMUSG00000062896 | Rpl31-ps11 | -8.415426 | 0.0235757 | 1 |
| ENSMUSG00000022510 | Trp63 | -4.4422812 | 0.02359867 | 1 |
| ENSMUSG00000028262 | Clca3a2 | -3.6526059 | 0.02368629 | 1 |
| ENSMUSG00000118077 | Gm50315 | 14.4925683 | 0.02410547 | 1 |
| ENSMUSG00000102577 | Gm37969 | -9.7228584 | 0.02410782 | 1 |
| ENSMUSG00000097287 | D130017N08 | -3.1466069 | 0.02418307 | 1 |
| ENSMUSG00000055313 | Pgbd1 | 6.54278137 | 0.02428914 | 1 |
| ENSMUSG00000112881 | Gm49344 | -29.630987 | 0.02445309 | 1 |
| ENSMUSG00000044903 | Psg22 | 3.50362615 | 0.02481486 | 1 |
| ENSMUSG00000074283 | Zfp109 | -3.2539692 | 0.02485238 | 1 |
| ENSMUSG00000034227 | Foxj1 | -13.41404 | 0.02487098 | 1 |
| ENSMUSG00000103839 | Gm37607 | 2.12557227 | 0.02501202 | 1 |
| ENSMUSG00000109564 | Muc16 | -21.497545 | 0.02516447 | 1 |
| ENSMUSG00000020672 | Sntg2 | -4.6334051 | 0.02535054 | 1 |
| ENSMUSG00000025813 | Homer2 | -3.7538461 | 0.02547796 | 1 |
| ENSMUSG00000103957 | Gm10766 | -18.863705 | 0.02552029 | 1 |
| ENSMUSG00000054662 | Ano9 | -8.5409254 | 0.02553935 | 1 |
| ENSMUSG00000099041 | Gm28035 | -33.273553 | 0.02566331 | 1 |
| ENSMUSG00000048521 | Cxcr6 | -1.9805214 | 0.02583857 | 1 |
| ENSMUSG00000118504 | 4933434E20 | -1.6329401 | 0.02590953 | 1 |
| ENSMUSG00000089945 | Pakap | -9.766659 | 0.02609289 | 1 |
| ENSMUSG00000057914 | Cacnb2 | 13.1524851 | 0.02645155 | 1 |
| ENSMUSG00000039545 | Flicr | -12.006973 | 0.02648644 | 1 |

| | | | | |
|---|---|---|---|---|
| ENSMUSG00000114934 | Gm48342 | -8.1007687 | 0.02654161 | 1 |
| ENSMUSG00000107388 | Gm42788 | -5.3458439 | 0.02656389 | 1 |
| ENSMUSG00000020932 | Gfap | -18.529312 | 0.02672972 | 1 |
| ENSMUSG00000026617 | Bpnt1 | 1.19931706 | 0.02675931 | 1 |
| ENSMUSG00000118053 | Gm50244 | -4.7849545 | 0.02678908 | 1 |
| ENSMUSG00000007034 | Slc44a4 | 4.10748932 | 0.02698486 | 1 |
| ENSMUSG00000043794 | D830025C05I | 21.0704047 | 0.0270084 | 1 |
| ENSMUSG00000113432 | 8430406P12I | 4.32451738 | 0.02702259 | 1 |
| ENSMUSG00000075593 | Gal3st4 | -14.327346 | 0.02728653 | 1 |
| ENSMUSG00000114255 | Gm10734 | 9.78041948 | 0.02730517 | 1 |
| ENSMUSG00000098041 | Gm26981 | 9.68072407 | 0.02767054 | 1 |
| ENSMUSG00000097239 | Gm27029 | 2.16826699 | 0.0279331 | 1 |
| ENSMUSG00000100594 | 2810414N06 | -2.9324478 | 0.02795588 | 1 |
| ENSMUSG00000107336 | Gm43461 | -5.8811288 | 0.02801193 | 1 |
| ENSMUSG00000109305 | Smim38 | -8.5094464 | 0.02801596 | 1 |
| ENSMUSG00000108854 | D830036C21I | 14.9811434 | 0.02804835 | 1 |
| ENSMUSG00000060402 | Chst8 | -21.474019 | 0.02805418 | 1 |
| ENSMUSG00000049303 | Syt12 | 3.1895841 | 0.02822568 | 1 |
| ENSMUSG00000112721 | Gm35608 | -13.012705 | 0.02830793 | 1 |
| ENSMUSG00000092526 | Gm17907 | -8.6196012 | 0.02853008 | 1 |
| ENSMUSG00000042686 | Jph1 | -7.2127807 | 0.02853265 | 1 |
| ENSMUSG00000087535 | Zmiz1os1 | -3.0666445 | 0.02877132 | 1 |
| ENSMUSG00000089849 | Runx2os2 | -17.277134 | 0.02925659 | 1 |
| ENSMUSG00000096923 | A730071L15I | 19.5578579 | 0.0296203 | 1 |
| ENSMUSG00000117628 | Gm50012 | 2.69267857 | 0.02973212 | 1 |
| ENSMUSG00000024228 | Nudt12 | -3.2693526 | 0.02983312 | 1 |
| ENSMUSG00000030669 | Calca | -16.818586 | 0.02983662 | 1 |
| ENSMUSG00000031568 | Rwdd4a | 1.18492641 | 0.02996031 | 1 |
| ENSMUSG00000057147 | Dph6 | 1.18487092 | 0.02998942 | 1 |
| ENSMUSG00000035504 | Reep6 | 1.5406042 | 0.03021509 | 1 |
| ENSMUSG00000061878 | Sphk1 | -1.9902969 | 0.03029995 | 1 |
| ENSMUSG00000028517 | Plpp3 | -1.3890233 | 0.03043209 | 1 |
| ENSMUSG00000111325 | Gm47140 | -7.3731571 | 0.03052507 | 1 |
| ENSMUSG00000079022 | Col22a1 | 11.0370472 | 0.03061166 | 1 |
| ENSMUSG00000062773 | Tex101 | 20.7670525 | 0.030842 | 1 |
| ENSMUSG00000115270 | 5430430K15I | -19.993007 | 0.03131167 | 1 |
| ENSMUSG00000058443 | Rpl10-ps3 | -1.5552182 | 0.03137482 | 1 |
| ENSMUSG00000021974 | Fgf9 | -6.6397927 | 0.03197469 | 1 |
| ENSMUSG00000100782 | Gm28231 | -23.97039 | 0.03217873 | 1 |
| ENSMUSG00000042385 | Gzmk | -6.0391446 | 0.03235372 | 1 |
| ENSMUSG00000091102 | 5830462I19R | -5.7712923 | 0.03240319 | 1 |
| ENSMUSG00000063887 | Nlgn1 | -15.242423 | 0.03240861 | 1 |
| ENSMUSG00000087042 | Gm11611 | -16.875096 | 0.03251329 | 1 |
| ENSMUSG00000041733 | Coq5 | 1.17464223 | 0.03280499 | 1 |
| ENSMUSG00000000365 | Rnf17 | 4.47562189 | 0.03295153 | 1 |

| | | | | |
|---|---|---|---|---|
| ENSMUSG00000027408 | Cpxm1 | -1.8080403 | 0.03333228 | 1 |
| ENSMUSG00000117809 | Gm50478 | 34.3113922 | 0.03337186 | 1 |
| ENSMUSG00000072494 | Ppp1r3e | -1.3724245 | 0.03352455 | 1 |
| ENSMUSG00000110547 | Gm29773 | -11.960435 | 0.0336389 | 1 |
| ENSMUSG00000074472 | Zfp872 | -9.8630716 | 0.03377257 | 1 |
| ENSMUSG00000051359 | Ncald | -3.827283 | 0.03380259 | 1 |
| ENSMUSG00000031520 | Vegfc | -2.0362701 | 0.03388628 | 1 |
| ENSMUSG00000038879 | Nipal2 | -2.7527003 | 0.03405032 | 1 |
| ENSMUSG00000027500 | Stmn2 | -11.427229 | 0.03407397 | 1 |
| ENSMUSG00000041556 | Fbxo2 | -10.340168 | 0.03421841 | 1 |
| ENSMUSG00000024055 | Cyp4f13 | -1.5283459 | 0.0343133 | 1 |
| ENSMUSG00000029154 | Cwh43 | 12.675987 | 0.03436247 | 1 |
| ENSMUSG00000038305 | Spats2l | -2.0458493 | 0.03444827 | 1 |
| ENSMUSG00000024176 | Sox8 | 4.68012053 | 0.03483153 | 1 |
| ENSMUSG00000046623 | Gjb4 | -6.254946 | 0.03527275 | 1 |
| ENSMUSG00000024936 | Kcnk7 | -6.6588231 | 0.03534498 | 1 |
| ENSMUSG00000019851 | Perp | -2.0984531 | 0.03541276 | 1 |
| ENSMUSG00000085024 | C230035I16R | -9.7789398 | 0.03541591 | 1 |
| ENSMUSG00000108308 | Gm45218 | 28.1513617 | 0.03541845 | 1 |
| ENSMUSG00000087354 | 4930404I05R | -6.3563732 | 0.03546656 | 1 |
| ENSMUSG00000031595 | Pdgfrl | -2.5153756 | 0.03551712 | 1 |
| ENSMUSG00000040247 | Tbc1d10c | -2.1862852 | 0.03571547 | 1 |
| ENSMUSG00000104633 | Gm42421 | -7.0373153 | 0.03590965 | 1 |
| ENSMUSG00000031119 | Gpc4 | -1.6975528 | 0.03608563 | 1 |
| ENSMUSG00000104011 | Gm32391 | 16.2119268 | 0.03632219 | 1 |
| ENSMUSG00000027305 | Ndufaf1 | 1.23139774 | 0.03642854 | 1 |
| ENSMUSG00000087382 | Ctcflos | 19.3514693 | 0.03644374 | 1 |
| ENSMUSG00000059824 | Dbp | -2.1772874 | 0.03655543 | 1 |
| ENSMUSG00000038276 | Asic3 | -12.881361 | 0.0368636 | 1 |
| ENSMUSG00000013367 | Iglon5 | -6.7558414 | 0.03697971 | 1 |
| ENSMUSG00000106218 | Gm43713 | -18.699913 | 0.0372137 | 1 |
| ENSMUSG00000072647 | Adam1a | 2.49846497 | 0.037798 | 1 |
| ENSMUSG00000062785 | Kcnc3 | -8.0567236 | 0.03786131 | 1 |
| ENSMUSG00000059742 | Kcnh7 | -4.7611717 | 0.03792731 | 1 |
| ENSMUSG00000074506 | Gm10705 | 4.04347032 | 0.03813674 | 1 |
| ENSMUSG00000033542 | Arhgef5 | -1.43343 | 0.03814241 | 1 |
| ENSMUSG00000000094 | Tbx4 | -1.3398655 | 0.03821479 | 1 |
| ENSMUSG00000030218 | Mgp | -2.1067867 | 0.03823146 | 1 |
| ENSMUSG00000054003 | Tdrd9 | -18.709087 | 0.03836759 | 1 |
| ENSMUSG00000033389 | Arhgap44 | -3.1178795 | 0.03892857 | 1 |
| ENSMUSG00000104876 | Trdc | -3.8130393 | 0.03896755 | 1 |
| ENSMUSG00000031273 | Col4a6 | -6.7733575 | 0.03898993 | 1 |
| ENSMUSG00000093769 | H3c14 | -3.400392 | 0.03901626 | 1 |
| ENSMUSG00000027239 | Mdk | 3.96549784 | 0.03925805 | 1 |
| ENSMUSG00000037990 | Sh3rf3 | -3.4614821 | 0.03932544 | 1 |

| | | | | |
|---|---|---|---|---|
| ENSMUSG00000020607 | Lratd1 | -8.123102 | 0.0393286 | 1 |
| ENSMUSG00000036169 | Sostdc1 | -8.0486156 | 0.03937921 | 1 |
| ENSMUSG00000086607 | 4930511M06 | 2.38803974 | 0.03967857 | 1 |
| ENSMUSG00000105021 | Gm8234 | -21.127066 | 0.03978469 | 1 |
| ENSMUSG00000047181 | Samd14 | -1.4665133 | 0.03983387 | 1 |
| ENSMUSG00000044505 | Lingo4 | -12.286132 | 0.03995935 | 1 |
| ENSMUSG00000020279 | Il9r | -5.2828367 | 0.03998932 | 1 |
| ENSMUSG00000000037 | Scml2 | 4.5194214 | 0.04004749 | 1 |
| ENSMUSG00000107205 | Gm42576 | -3.3840152 | 0.04012205 | 1 |
| ENSMUSG00000001542 | Ell2 | 1.35497578 | 0.04017144 | 1 |
| ENSMUSG00000054247 | Gm9939 | 25.3729237 | 0.04037586 | 1 |
| ENSMUSG00000059878 | Zfp422 | 1.50949767 | 0.04039653 | 1 |
| ENSMUSG00000092368 | A930015D03 | 3.14953475 | 0.04041963 | 1 |
| ENSMUSG00000107932 | Gm44432 | 11.9385052 | 0.04044789 | 1 |
| ENSMUSG00000003354 | Ccdc65 | -4.8255434 | 0.04048842 | 1 |
| ENSMUSG00000072966 | Gprasp2 | -10.473753 | 0.04064936 | 1 |
| ENSMUSG00000050623 | Catsperz | 4.39878068 | 0.04076085 | 1 |
| ENSMUSG00000059195 | Gm12715 | -1.5175583 | 0.04085397 | 1 |
| ENSMUSG00000048191 | Muc6 | -12.287317 | 0.04142689 | 1 |
| ENSMUSG00000074805 | Il1bos | -15.28734 | 0.04150561 | 1 |
| ENSMUSG00000012282 | Wnt8a | 16.5817321 | 0.04160586 | 1 |
| ENSMUSG00000057234 | Mettl15 | -1.420959 | 0.04168113 | 1 |
| ENSMUSG00000083773 | Gm13394 | -2.7398932 | 0.04197917 | 1 |
| ENSMUSG00000047797 | Gjb1 | -19.664149 | 0.04212416 | 1 |
| ENSMUSG00000047412 | Zbtb44 | 1.22706118 | 0.04212575 | 1 |
| ENSMUSG00000087006 | Gm13889 | -2.2293397 | 0.04232805 | 1 |
| ENSMUSG00000090812 | Samd15 | -8.3386443 | 0.0423721 | 1 |
| ENSMUSG00000040323 | Gm15429 | -2.6456848 | 0.0426757 | 1 |
| ENSMUSG00000065999 | Zfp985 | 1.30518196 | 0.04268638 | 1 |
| ENSMUSG00000047428 | Dlk2 | -12.491054 | 0.04269263 | 1 |
| ENSMUSG00000089662 | Gm14057 | -17.040856 | 0.0427843 | 1 |
| ENSMUSG00000087249 | Gm16062 | 1.8465797 | 0.04307467 | 1 |
| ENSMUSG00000090628 | Gm17083 | -10.304613 | 0.04309103 | 1 |
| ENSMUSG00000048450 | Msx1 | -2.3553743 | 0.0434742 | 1 |
| ENSMUSG00000022679 | Mpv17l | -1.8008368 | 0.04361674 | 1 |
| ENSMUSG00000053646 | Plxnb1 | -1.8479177 | 0.04375176 | 1 |
| ENSMUSG00000053588 | A730061H03 | 9.58453953 | 0.04378408 | 1 |
| ENSMUSG00000022151 | Ttc33 | 1.24787053 | 0.04402077 | 1 |
| ENSMUSG00000106508 | 4933425M03 | -5.1463459 | 0.04446058 | 1 |
| ENSMUSG00000057280 | Musk | -19.959766 | 0.04479973 | 1 |
| ENSMUSG00000115955 | 9530056E24I | -5.5686414 | 0.04486737 | 1 |
| ENSMUSG00000113637 | Gm7049 | 10.2076711 | 0.04499375 | 1 |
| ENSMUSG00000097890 | 4930547M16 | -11.906484 | 0.04527418 | 1 |
| ENSMUSG00000029361 | Nos1 | 2.51734209 | 0.04547013 | 1 |
| ENSMUSG00000097602 | 4930519P11I | -9.1962472 | 0.04551225 | 1 |

| | | | | |
|---|---|---|---|---|
| ENSMUSG00000071005 | Ccl19 | -6.0722556 | 0.04552267 | 1 |
| ENSMUSG00000031171 | Ftsj1 | 1.27672999 | 0.04564883 | 1 |
| ENSMUSG00000091050 | 9330020H09I | -2.0429535 | 0.04604946 | 1 |
| ENSMUSG00000097022 | BC001981 | -16.491134 | 0.0460502 | 1 |
| ENSMUSG00000031428 | Zcchc18 | -2.9638245 | 0.04611177 | 1 |
| ENSMUSG00000056917 | Sipa1 | -1.1652776 | 0.04640339 | 1 |
| ENSMUSG00000103966 | Gm37120 | -9.9465413 | 0.04644445 | 1 |
| ENSMUSG00000111116 | Gm48065 | 2.54148788 | 0.04676531 | 1 |
| ENSMUSG00000102151 | Gm37472 | -10.503549 | 0.04677244 | 1 |
| ENSMUSG00000103382 | Gm37755 | 6.60636154 | 0.04694514 | 1 |
| ENSMUSG00000035390 | Brsk1 | -2.482791 | 0.04725358 | 1 |
| ENSMUSG00000090622 | A930033H14 | -1.9442677 | 0.04738905 | 1 |
| ENSMUSG00000108753 | Gm45094 | -4.6739783 | 0.04738933 | 1 |
| ENSMUSG00000038242 | Aox4 | -11.536748 | 0.04760229 | 1 |
| ENSMUSG00000020891 | Alox8 | -6.8053001 | 0.0477414 | 1 |
| ENSMUSG00000004347 | Pde1c | -11.317128 | 0.04783781 | 1 |
| ENSMUSG00000031378 | Abcd1 | -1.2593947 | 0.04789851 | 1 |
| ENSMUSG00000020323 | Prss57 | -10.056091 | 0.0479328 | 1 |
| ENSMUSG00000028488 | Sh3gl2 | 11.1522477 | 0.04829527 | 1 |
| ENSMUSG00000028348 | Cavin4 | 3.45125414 | 0.04833102 | 1 |
| ENSMUSG00000041754 | Trem3 | 4.25586531 | 0.04844242 | 1 |
| ENSMUSG00000022594 | Lynx1 | -1.6110117 | 0.04847666 | 1 |
| ENSMUSG00000091255 | Speer4e | 13.7343815 | 0.04851934 | 1 |
| ENSMUSG00000001986 | Gria3 | -3.2787638 | 0.04872185 | 1 |
| ENSMUSG00000103901 | Gm37499 | 1.7495523 | 0.04879461 | 1 |
| ENSMUSG00000097462 | 9530026P05I | -11.802 | 0.04933937 | 1 |
| ENSMUSG00000039798 | 2600006K01I | -9.5015673 | 0.04944408 | 1 |
| ENSMUSG00000103869 | Gm37420 | -3.6522808 | 0.04958016 | 1 |
| ENSMUSG00000024778 | Fas | -1.6951941 | 0.04967381 | 1 |
| ENSMUSG00000030775 | Trat1 | -4.0037339 | 0.04981759 | 1 |
| ENSMUSG00000020072 | Pbld2 | -10.378673 | 0.04997241 | 1 |
