## Supplemental Table 4 for "Environmentally Induced Sperm RNAs Transmit Cancer Susceptibility to Offspring in a Mouse Model"

**Table S4** - List of Antibodies used for IMC

| **Antibody** | **Metal** | **Clone** | **Dilution** | **Supplier** | **Catalog number** |
|---|---|---|---|---|---|
| aSMA | 141 Pr | 1A4 | 1:100 | Fluidigm | 3141017D |
| CD19 | 142 Nd | 6OMP31 | 1:50 | Fluidigm | 3142014D |
| Vimentin | 143 Nd | D21H3 | 1:150 | Fluidigm | 3143027D |
| EpCAM/CD326 | 144 Nd | 9C4 | 1:50 | Fluidigm | 3144026D |
| panCK | 146 Nd | polyclonal | 1:50 | Agilent/Dako | Z0622 |
| CD45 | 147 Sm | 30-F11 | 1:50 | Fluidigm | 3147003B |
| CD11b | 149 Sm | EPR1344 | 1:150 | Fluidigm | 3149028D |
| CD31 | 152 Sm | D8V9E | 1:150 | Cell Signaling Technology | 77699BF |
| F4/80 | 154 Sm | D2S9R | 1:150 | Cell Signaling Technology | 70076BF |
| CD4 | 156 Gd | EPR6855 | 1:50 | Fluidigm | 3156033D |
| E-cadherin | 158 Gd | 24E10 | 1:100 | Fluidigm | 3158029D |
| CD68 | 159 Tb | polyclonal | 1:50 | ThermoFisher Scientific | PA5-78996 |
| CD11c | 160 Gd | D1V9Y | 1:50 | Cell Signaling Technology | 97585BF |
| CD8a | 162 Dy | D4W2Z | 1:50 | Cell Signaling Technology | 98941BF |
| Arginase-1 | 164 Dy | D4E3M | 1:50 | Fluidigm | 3164027D |
| FOXP3 | 165 Ho | FJK-16s | 1:50 | Fluidigm | 3165024A |
| PD-L1 | 166 Er | polyclonal | 1:50 | R&D Systems | AF1019 |

| | | | | | |
|---|---|---|---|---|---|
| Granzyme B | 167 Er | EPR20129-217 | 1:150 | Fluidigm | 3167021D |
| Ki67 | 168 Er | B56 | 1:150 | Fluidigm | 3168022D |
| Collagen type I | 169 Tm | polyclonal | 1:150 | Fluidigm | 3169023D |
| CD3 | 170 Er | polyclonal | 1:100 | Fluidigm | 3170019D |
| Cleaved caspase 3 | 172 Yb | 5A1E | 1:50 | Fluidigm | 3172027D |
