## Supplemental figures 1-6 for "Environmentally Induced Sperm RNAs Transmit Cancer Susceptibility to Offspring in a Mouse Model"

Supplementary Figure 1

a.

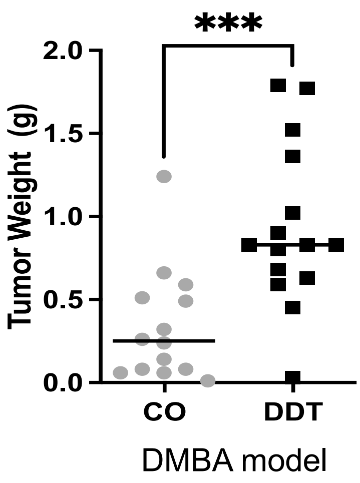


b.

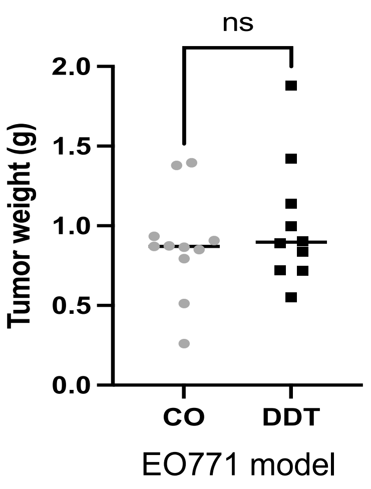

### Supplementary Figure 2

a.

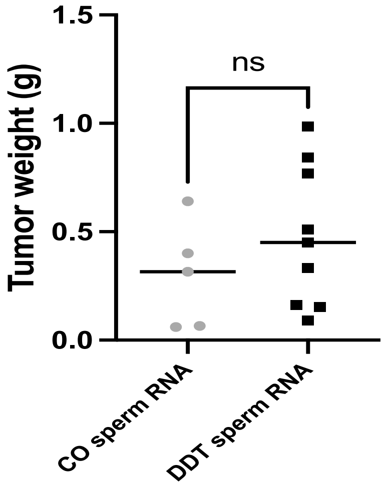


DMBA model

b.

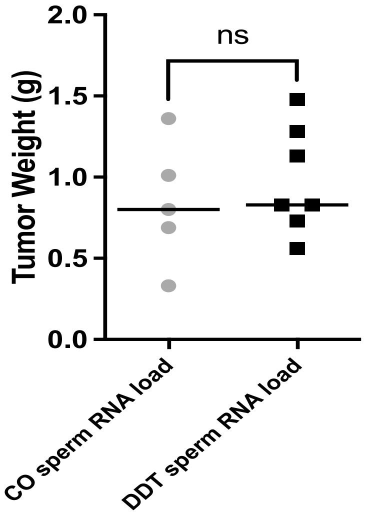


EO771 model

### Supplementary Figure 3

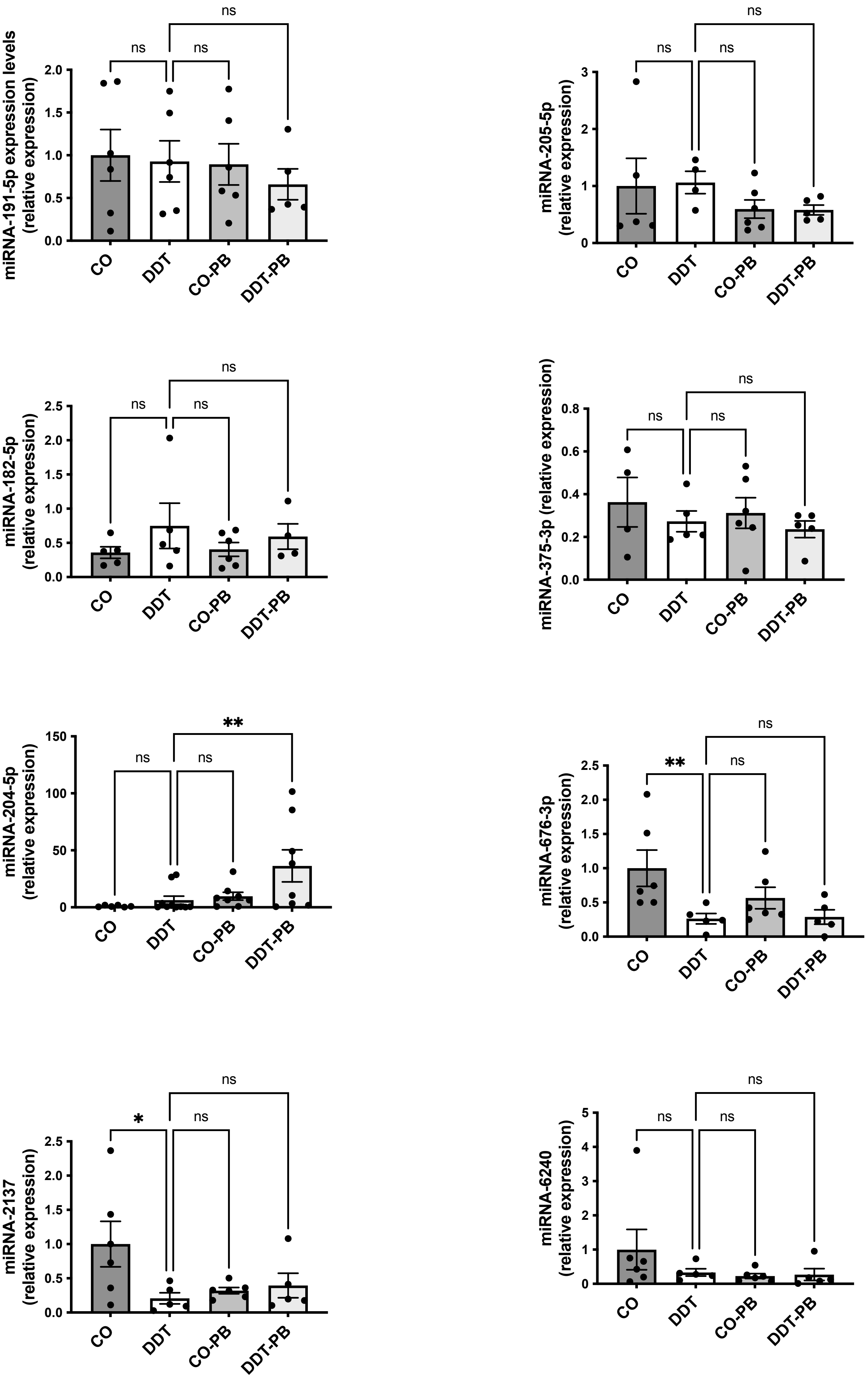

### Supplementary Figure 4

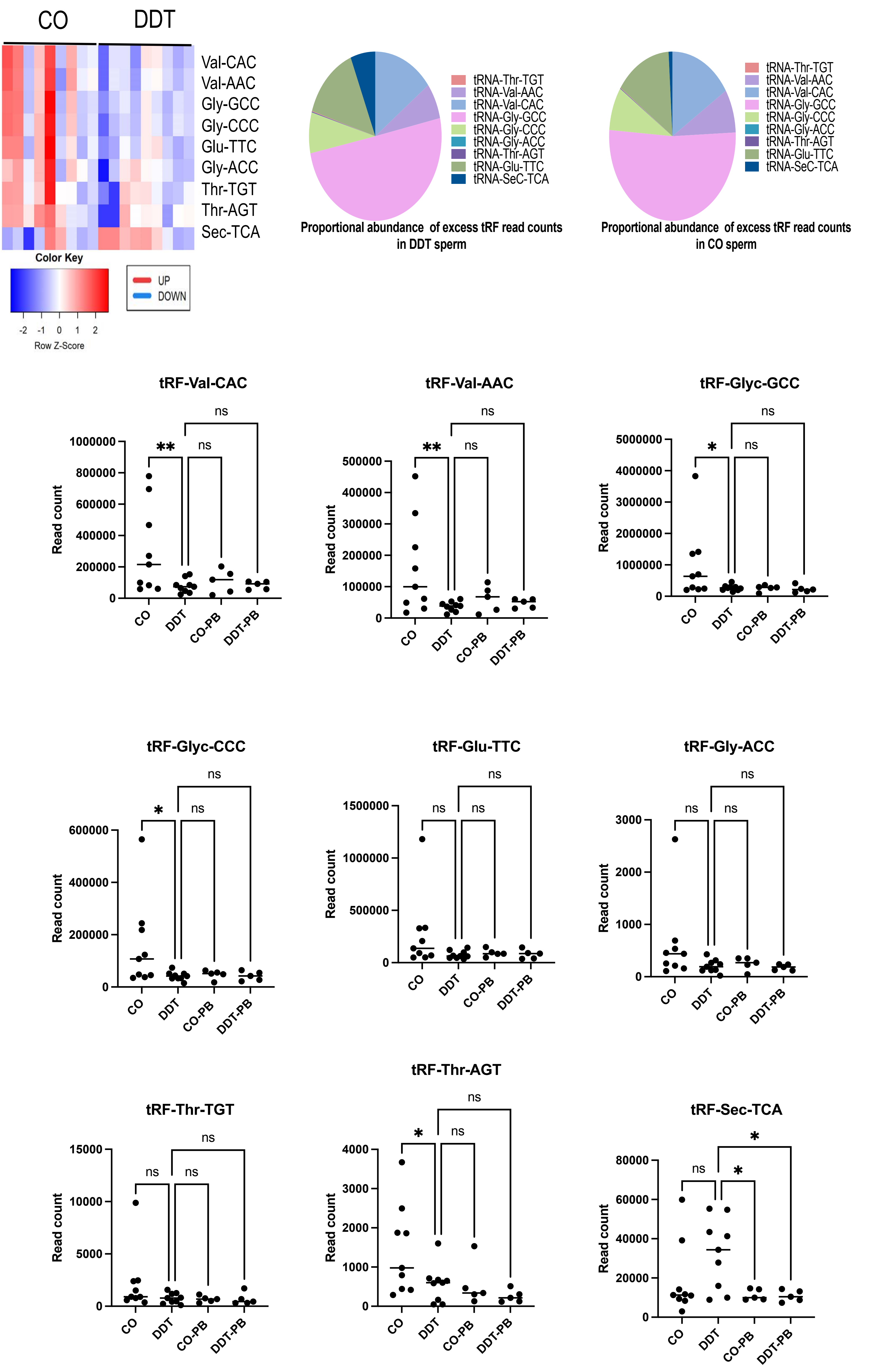

Supplementary Figure 5

a.

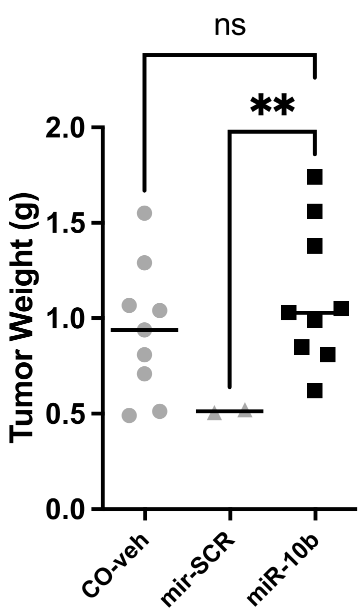

Supplementary Figure 6

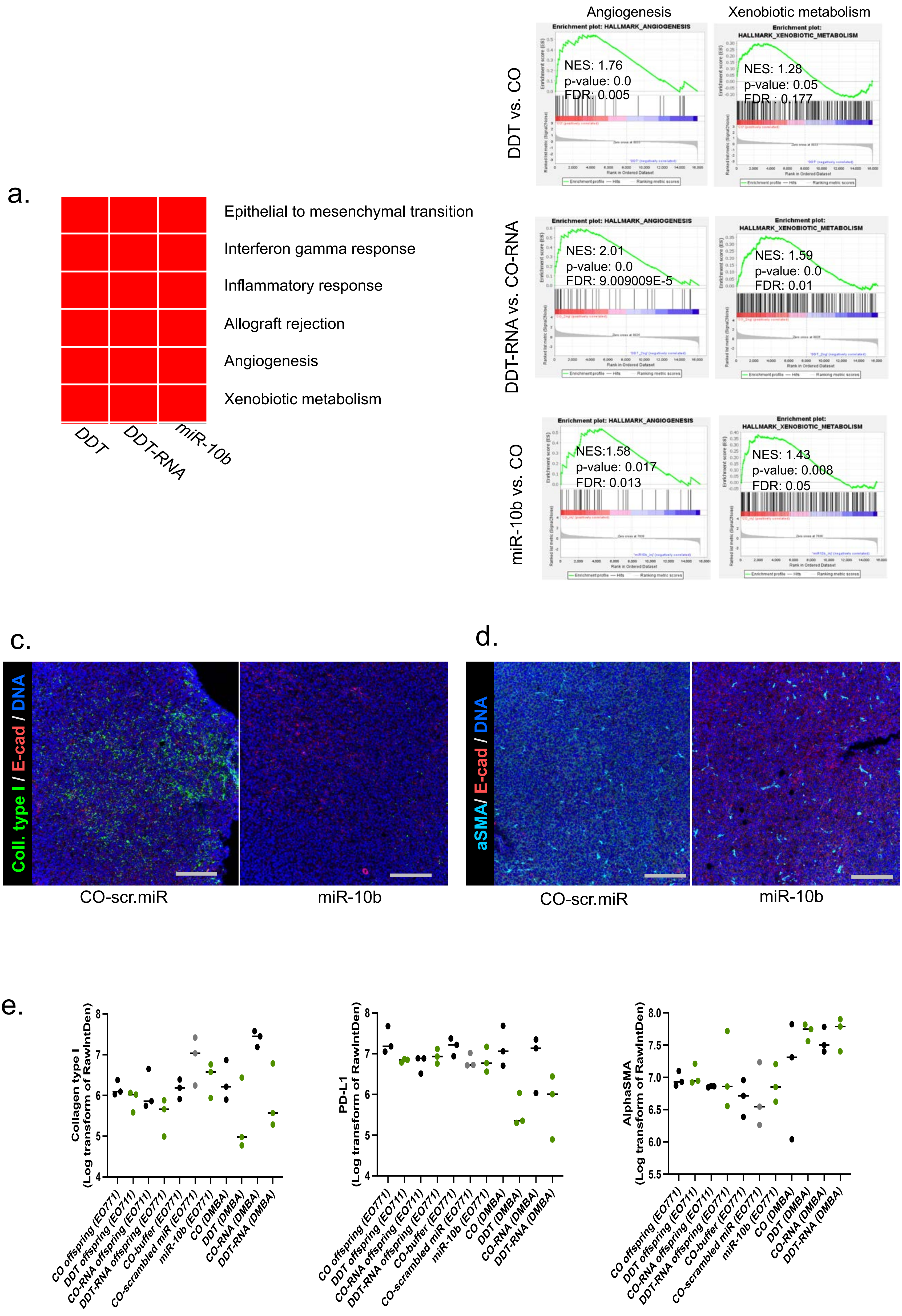

#### Supplementary Figure Legends

**Fig-S1. Mammary tumor weight in offspring of DDT-exposed males.**
End-point mammary tumor weight in carcinogen-induced (n=13-15) and orthotopic (EO771 cells, n=10-11) tumors in CO and DDT female offspring. Data includes both measurable and non-measurable tumors. Horizontal bars in scatter plots represent the mean. ***p<0.001; ns, non-significant by t-test.

**Fig-S2. Mammary tumor weight in offspring generated with sperm RNA load of DDT-exposed males .**
End-point mammary tumor weight in carcinogen-induced (n=5-9) and orthotopic (EO771 cells, n=5-7) tumors in CO-RNA and DDT-RNA female offspring. Data includes both measurable and non-measurable tumors. Horizontal bars in scatter plots represent the mean. ns, non-significant by t-test.

**Fig-S3. miRNA expression levels in sperm of DDT-exposed males after treatment with the hepatic enzyme inducer, phenobarbital.**
(a) Expression levels of miRNAs, assessed by q-PCR, in sperm of CO and DDT-exposed male mice treated with PB or vehicle injection (n=5-6). Expression levels of miRNAs are relative to miR-26a. Data are shown as mean± SEM; ns, non-significant, *p<0.05, **p<0.01 by one-way ANOVA.

**Fig-S4. DDT exposure reprograms the sperm tRNA fragments (tRFs), but treatment with phenobarbital does not revert their expression.**

(a) Heat-map showing differentially expressed tRFs in sperm of CO and DDT-exposed males (n=9).

(b-c) Proportional abundance of differentially expressed tRFs (DESeq2 normalized counts) in sperm of CO and DDT-exposed males (n=9).

(d) tRFs expression levels (DESeq2 normalized counts), assessed by RNA-seq, in sperm of CO and DDT-exposed male mice treated with PB or vehicle injection (n=5-9). Horizontal bars in scatter plots represent the mean. ns, non-significant, *p<0.05, **p<0.01 by one-way ANOVA.

**Fig-S5. Mammary tumor weight in offspring generated with synthetic miR-10b.**
End-point mammary tumor weight in orthotopic (EO771 cells, n=2-9) tumors in CO and miRNA-10b female offspring. Data includes both measurable and non-measurable tumors. Horizontal bars in scatter plots represent the mean. ns, non-significant, **p<0.01 by one-way ANOVA.

**Fig-S6. Tumors of DDT-derived offspring show distinct signaling programs compared to controls.**
(a-b) GSEA pathways in mammary tumors of DDT-derived offspring (DDT, DDT-RNA and miRNA-10b, n=3), showing (a) overlapping GSEA pathways and (b) additional significantly down-regulated pathways (Angiogenesis and Xenobiotic Metabolism) in tumors of DDT, DDT-RNA and miRNA-10b offspring.

(c)IMC staining for collagen type I (green), E-cadherin (red), DNA (blue, iridium intercalator) in orthotopic mammary tumors (EO771 cells, n=3-6) of miR-10b offspring. Scale bar equals to 200 μm.

(d)IMC staining for α-SMA (cyan), E-cadherin (red), DNA (blue, iridium intercalator) in orthotopic mammary tumors (EO771 cells, n=3-6) of miR-10b offspring. Scale bar equals to 200 μm.

(e)Scatter plot graphs showing individual group quantification from IMC data for collagen type I, α-SMA and PD-L1 in mammary tumors of DDT-derived offspring. Horizontal bars in scatter plots represent the mean.
